# Breaking new ground into RAD51–BRC repeats interplay in Homologous Recombination

**DOI:** 10.1101/2025.11.13.688182

**Authors:** Francesco Rinaldi, Pedro Franco, Marina Veronesi, Elisa Romeo, Veronica Bresciani, Giulia Varignani, Federico Catalano, Mattia Bernetti, Matteo Masetti, Julian D. Langer, Stefania Girotto, Andrea Cavalli

## Abstract

Homologous recombination (HR) is a critical repair pathway involving numerous proteins that ensure error-free DNA double-strand breaks (DSBs) repair. Dysfunction in HR components can compromise genome integrity. Despite advances, many aspects of HR remain poorly understood. Notably, even one of the earliest identified and most critical interactions, between RAD51 and BRCA2, remains incompletely characterized, mainly due to the lack of structural data. This study presents a comprehensive biophysical analysis of the RAD51–BRC repeats interaction, integrating computational and experimental approaches. Starting with assessing the correlation between the binding affinities of individual BRC repeats and their impact on RAD51 disassembly, our investigation extends to larger BRCA2 truncations, offering unprecedented insights into the molecular determinants of RAD51 recognition. As mutations in the BRC repeats impair RAD51 recruitment and are associated with cancer, these results provide a valuable framework for interpreting pathogenic variants and guiding precision medicine therapies.

**GRAPHICAL ABSTRACT:** 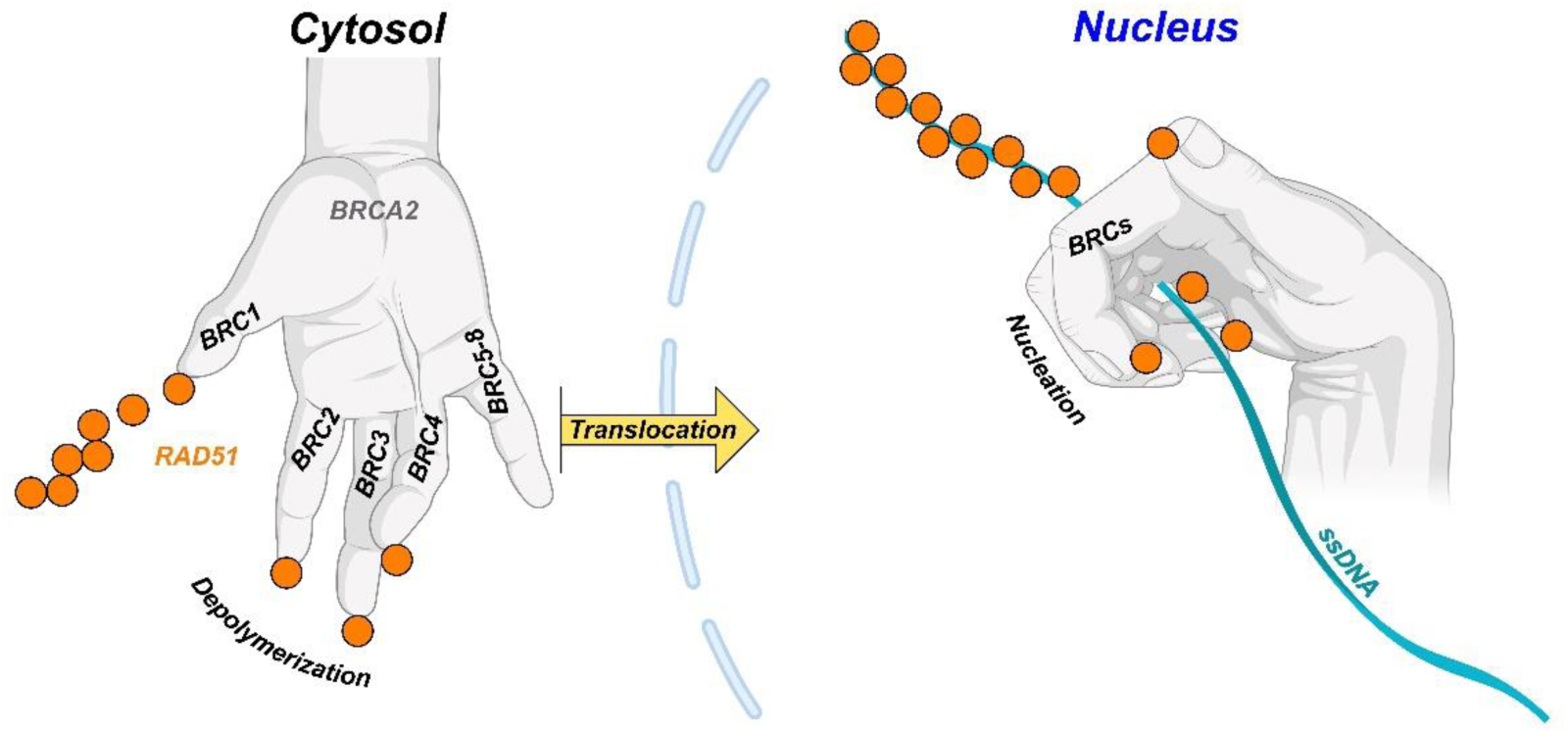

## INTRODUCTION

Homologous recombination (HR) is a high-fidelity DNA repair mechanism that cells use to fix potentially lethal DNA double-strand breaks (DSBs)(1–3). HR machinery must rely on the coordinated interaction of numerous factors to carry out this highly sophisticated and finely tuned repair process(1–4). Remarkably, single mutations in various components (e.g., RAD51, BRCA1, BRCA2, RAD51 paralogs, PALB2, to mention a few) of this system are strongly associated with increased cancer risk, underscoring the critical importance of the individual cofactors in maintaining the integrity of HR, and by extension, genome integrity(1). Although these critical pieces of the puzzle are gradually coming together, offering early glimpses into how the various players in this intricate machinery may interact as parts of a wider “recombinosome”, much remains to be elucidated within this complex and dynamic process(3,5). The high degree of structural flexibility and mobility among HR components adds to its intricacy, leaving many fundamental interactions poorly understood(3,6). Indeed, even one of the earliest identified and core interactions of HR, that between RAD51 and BRCA2, remains only partially characterized(7–10). After DNA-end resection, BRCA2 acts as a key player ensuring the cytosolic recruitment of the RAD51 recombinase and promoting its loading onto the generated ssDNA(1–3,5). Since RAD51 lacks the nuclear localization sequence (NLS), the interaction with BRCA2 is essential for cytosolic RAD51 recruitment and subsequent translocation into the nucleus(11). Furthermore, the formation of RAD51 filaments on ssDNA is particularly crucial in HR as it allows the identification and base-pairing of homologous sequences, which are then exploited for the high-fidelity repair of DNA(12). BRCA2 interacts with RAD51 through two distinct domains: the C-terminal region, which contributes to the stabilization of RAD51 oligomers during strand-exchange activity, and the central region, which engages RAD51 through eight so-called BRC repeats, each comprising 35-40 amino acids(3,7,8,13–15). Interestingly, the BRC repeats are conserved across eukaryotic organisms within the same class, although their number significantly changes among more distantly related species (**Figure S1**)(13,14). Notably, individual BRC repeats in humans exhibit limited sequence conservation among themselves (**Figure S1**)(13,14).

Despite the importance of this protein-protein interaction, only low-resolution information is available for full-length BRCA2 alone or in complex with RAD51(3,9,10,16–18). The only high-resolution structural insight currently available on the BRC repeat – RAD51 interaction was reported in 2002 when the crystal structure of an N-terminal truncated RAD51 in complex with BRC4, the fourth BRCA2 repeat, was solved(3,8,18). This study revealed that two motifs, FXXA and LFDE, are essential for BRC4 binding to the C-terminal domain of RAD51(8), but structural insights into the interactions between the other BRC-repeats and RAD51 are still lacking(7,8,18–25). To date, no direct biophysical investigation of the affinity of BRC-repeats with fully human RAD51 has been reported. However, indirect biochemical evidence suggests that the first four repeats can inhibit DNA binding, whereas BRC5-8 have the opposite effect, promoting RAD51-DNA interaction(26,27). This indicates distinct roles for different repeats during RAD51 nucleation on resected ssDNA(26,27). Nevertheless, no data are currently available on the individual contributions of the BRC repeats to RAD51 nuclear recruitment, a crucial step in HR. Such information would be invaluable for guiding the rational design of novel anticancer therapies targeting this critical interface, which has already been recognized as a validated target in “chemically induced synthetic lethality” strategies(3,18,28–30). This approach would enable extending the concept of synthetic lethality (SL) to non-BRCA2-mutated patients, a promising strategy to tackle aggressive cancer forms such as pancreatic cancer(3,18,28–30). Furthermore, recent evidence suggests that interaction patterns similar to those observed between RAD51 and BRCA2 also occur among other HR factors(31). Therefore, insights into the RAD51-BRCA2 interaction could elucidate the coordinated mechanisms underlying the interplay between BRC repeats and other components of the HR machinery, ultimately revealing novel potential targets for SL-based cancer therapies.

Here, we investigate the BRC-repeats-RAD51 interaction by combining computational methods, biophysical experiments, and integrative structural biology. We aimed to comprehensively characterize the interaction of each BRC repeat with RAD51 and to unveil novel structural insights into both individual BRC-repeats and larger BRCA2 truncations, which include multiple peptides and their connecting regions. Our findings reveal that peptides’ affinities are directly correlated to their ability to disassemble cytosolic RAD51 fibrils. Moreover, these data also shed light on the importance of the interaction between the BRC repeats and the N-terminal domain of RAD51, as well as the dynamic and multifaceted behavior of the regions linking the first four BRC repeats.

## MATERIAL AND METHODS

### AlphaFold predictions

Predictions reported in this work were generated through either AlphaFold 2.3.2 or Alphafold3 as Singularity containers on the IIT-HPC CPU/GPU Cluster (logical name Franklin). For Alphafold3, predictions were also generated by utilizing the web-server version. For Alphafold2, the model preset used for prediction was set to multimer, enabling Amber relaxation, which resolves remaining structural violations and clashes in the predicted structure(32), only for the highest-ranked model. Database preset was set to full database (full_dbs databases: bfd, mgnify (mgy_clusters_2022_05), pdb_mmcif, pdb_seqres, pdb70, UniRef30_2023_02, uniprot, uniref90). Competitive predictions were generated through Alphafold 2.3.2 by inserting in the FASTA file the sequences of Δ97-RAD51 and RAD51, BRC4, and another BRC repeat in line with a former report by Chang et al.(33,34) To assess which peptide was correctly modeled, all generated predictions were aligned to the crystal structure of the RAD51-BRC4 complex (PDB entry: 1n0w). The model of the RAD51-CAM833A complex was generated through Alphafold3, inputting its equivalent SMILE string. Graphs of multiple sequence alignment (MSA) coverages and sequence similarity predicted local distance difference test (pLDDT) and predicted aligned error (PAE) was generated through Python scripts adapted from https://raw.githubusercontent.com/jasperzuallaert/VIBFold/main/visualize_alphafold_resultr.py and https://raw.githubusercontent.com/busrasavas/AFanalysis/main/AFanalysis.py and https://github.com/flyark/AFM-LIS/blob/main/alphafold3_lis_v0.11.ipynb (35)

### Binding affinity predictions and molecular dynamics simulations

Starting from the structure of the RAD51-BRC4 complex (PDB Entry: 1N0W), a residue scanning procedure was conducted. Specifically, the BRC4 peptide sequence was mutated into other BRC repeats: BRC2, BRC4, BRC7, BRC5. Since mutation of BRC4 to BRC2 included a histidine residue, all possible states were considered for the histidine, namely the two delta (HID) and epsilon (HIE) tautomers of the neutral histidine and the protonated (HIP) state. The binding affinity predictions, given in kcal/mol, were calculated with Prime MM-GBSA in implicit solvent using default settings, as implemented in the Residue Scanning functionality of the BioLuminate module in Schrodinger’s Maestro(36), using the Schrodinger Suite version 2022-2 build 128. The first and last residues of the peptides, highly flexible and completely plumbed into the solvent, were excluded from the analysis. For statistical robustness, the analysis was performed on 100 structures from the MD simulation of the RAD51-BRC4 complex. Structures were extracted at regular intervals from the MD trajectory, with a 4 ns separation between consecutive frames to ensure even sampling. The statistical error was estimated as the standard error of the mean. Molecular Dynamics (MD) simulations were carried out for RAD51 in complex with BRC4, BRC7, BRC5 or BRC2. Using the crystal structure of the RAD51-BRC4 complex (PDB ID: 1N0W) as a template, the structure of RAD51 in complex with the other peptides were generated by manually mutating the BRC4 sequence to the sequences of BRC7, BRC5 and BRC2 through the Schrodinger’s Maestro interface. The four systems were prepared for MD using the Protein Preparation Wizard functionality in Schrodinger’s Maestro. Crystal ion and water molecules were retained, and termini of both RAD51 protein and the peptides were capped with acetyl (ACE) and n-methyl amide (NMA). A rhombic dodecahedron xy-square box, with edges distant 20 Å from the proteins, was used and filled with TIP3P waters. The systems were neutralized with Na^+^ ions, and NaCl at a concentration of 0.15 M was added to the buffer. The OPLS4 force field was used to model the system(37). The Minimization functionality in Schrodinger’s Maestro was employed, using default parameters, to energy minimize the system prior to equilibration. All systems were equilibrated for a total of 1 ns via a multi-step equilibration including three short simulations lasting 200 ps at 100, 200 and 300 K in the NVT ensemble using the Nose-Hoover chain thermostat and further 400 ps in NPT ensemble using the Martyna-Tobias-Klein barostat, using default parameters (38). A production run of plain MD in the NPT ensemble using the same barostat and lasting 400 ns was conducted for each system. Visualization of the MD trajectories was carried out with the Visual Molecular Dynamics (VMD) software version 1.9.4 (39). The Root-Mean-Square Fluctuation (RMSF) was calculated on α carbons of the peptides excluding the first and last residues. Statistical errors on the RMSF were estimated using bootstrapping, by dividing the trajectories into 5 blocks and performing 200 bootstrap iterations. All analyses of the MD trajectories were conducted with MDTraj version 1.9.7(40), NumPy and Matplotlib inside a Jupyter lab environment. All MD simulations were carried out with the Desmond software package(41) included in the Schrodinger Suite version 2022-2 build 128.

### Expression of labeled and unlabeled protein complexes

pET15b vector containing the coding sequence for His-TEV-RAD51[F86E, A89E], pET28a vectors with the coding sequences for His-TEV-BRC1, His-TEV-BRC2, His-TEV-BRC3, His-TEV-BRC4, His-TEV-BRC3-4, and pCOLA Duet containing both the His-TEV-BRC1-4 and His-TEV-RAD51[F86E, A89E] sequences were purchased from Genscript Biotech (Netherlands) B.V. All the inserts harbored an N-terminal His-tag cleavable through Tobacco Etch Virus (TEV) protease. Untagged Δ97-RAD51 (aminoacids 98-339) coding sequence was cloned into pET15b vector while untagged BRC1-2, BRC2-3, BRC2-4 sequences were cloned into pET28a vector. E. coli Rosetta™ 2(DE3) chemo-competent cells (71400, Novagen) were co-transformed with the following combination of plasmids pET15b Δ97-RAD51/pET28a His-TEV-BRC4, pET15b His-TEV-RAD51[F86E, A89E]/pET28a His-TEV-BRC1, pET15b His-TEV-RAD51[F86E, A89E]/pET28a His-TEV-BRC2, pET15b His-TEV-RAD51[F86E, A89E]/pET28a His-TEV-BRC3, pET15b His-TEV-RAD51[F86E, A89E]/pET28a His-TEV-BRC4, pET15b His-TEV-RAD51[F86E, A89E]/pET28a BRC1-2, pET15b His-TEV-RAD51[F86E, A89E]/pET28a BRC2-3, pET15b His-TEV-RAD51[F86E, A89E]/pET28a BRC3-4, pET15b His-TEV-RAD51[F86E, A89E]/pET28a BRC2-4 or pET15b His-TEV-RAD51[F86E, A89E]/pCOLA Duet His-TEV-BRC1-4/His-TEV-RAD51[F86E, A89E] according to the manufacturer protocols. Transformed bacteria were then plated onto LB Agar plates containing ampicillin (100 μg/mL) and kanamycin (50 μg/mL) and incubated overnight at 37°C to allow the colony growth. On the following day, a single colony was picked up from LB-Agar plates and inoculated in 2 mL of LB medium supplemented with ampicillin (100 μg/mL) and kanamycin (50 μg/mL) antibiotics in a 15 mL Falcon Tube. The following day, 1 mL of saturated cell culture was diluted in 500 mL of Terrific Broth (TB) supplemented with 1X 50/52 solution, 5 mM MgSO_4_ and antibiotics (ampicillin 110 µg/mL and kanamycin 50 μg/mL) and then incubated at 20°C in a 2L Erlenmeyer flask for 72 hours at 300-350 rpm. The initial volume utilized for the expression of Δ97-RAD51 and RAD51-BRC4 bacteria culture was 2L, while for all the other expressed complexes was 10L in order to obtain suitable amount of purified protein complexes for further characterization described in this work. To produce uniformly ^15^N labeled Δ97-RAD51 and RAD51-BRC4 1 mL of overnight saturated LB culture was inoculated in 500 mL of minimal auto-induction bacterial medium containing 50 mM Na_2_HPO_4_, 50 mM KH_2_PO_4_, 5 mM Na_2_SO_4_ 2 mM MgSO_4_, 0.1 mM FeCl_3_, 50 mM ^15^NH_4_Cl, trace metals, 1X 50/52, and antibiotics (110 µg/mL ampicillin and 50 μg/mL kanamycin). Bacteria cultures were incubated at 20°C in a 2L Erlenmeyer flask for 72 hours at 300-350 rpm. To obtain a suitable amount of protein for further NMR relaxation experiments the starting volume of bacteria cultures for protein expression of Δ97-RAD51-BRC4 and RAD51-BRC4 was 3L. After 72 hours, bacteria were centrifuged at 1500 rpm for 30 minutes at 4°C in a Beckman-Coulter Avanti J25 (rotor JLA-8.1000 The obtained pellet was washed with PBS and stored at -80 °C until used. Monomeric His-RAD51 [F86E, A89E] and His-RAD51 WT were expressed as previously described by our group (7,19)

### Protein complexes purification

Bacterial pellet was resuspended in an appropriate volume (10 mL per g of bacteria pellet) of Lysis Buffer (20 mM K_2_HPO_4_/KH_2_PO_4_ pH 8, 500 mM NaCl, 10 mM Imidazole, 10% Glycerol, 1X cOmplete™, EDTA-free Protease Inhibitor Cocktail, Roche) and lysed on ice through sonication (Branson Ultrasonics™ Sonifier™ SFX550 Digital Ultrasonic Cell Disruptors, 1/2” Tip or equivalent 60 seconds on, 30 seconds off for 10 minutes 50/60% Amplitude). The disrupted cell suspension was poured into polycarbonate tubes and centrifuged for 30 minutes at 20,000 rpm and 4°C, utilizing a JA 25.50 rotor (Beckman-Coulter Avanti J25). Supernatant fraction was filtered with a 0.20 μM filters to remove large aggregates prior to chromatography. All chromatographic steps were performed through a GE AKTA Purifier FPLC System (GE Healthcare) at room temperature with filtered (0.2 μm) and degassed buffers. Supernatant was applied through system pump onto a His-Trap HP (5 mL Cytiva) previously equilibrated with Buffer A (20 mM K_2_HPO_4_/KH_2_PO_4_ pH 8, 500 mM NaCl, 10 mM Imidazole, 10% Glycerol). Once all sample was loaded the column was washed at least with 7 column volumes (CVs) and then an additional 7 CVs washing step at 10% Buffer B (20 mM K_2_HPO_4_/KH_2_PO_4_ pH 8, 500 mM NaCl, 500 mM Imidazole, 10% Glycerol) was performed to remove a-specifically bound proteins. Δ97-RAD51 was eluted using a linear gradient from 10% to 100% of buffer B and pooling fractions eluted between 30% Buffer B and 70% Buffer B. For all other complexes two additional isocratic elution steps of 7 CVs each were performed corresponding to 16% (80 mM Imidazole) and 70% Buffer B (350 mM Imidazole). All these complexes eluted at 70% Buffer B. Then, fractions containing protein complexes were pooled and quantified with a Nanodrop spectrophotometer. His-TEV Protease (internally produced by adapting the protocol of Tropea and co-workers(42)) was added to obtain protease/protein ratio 1:30 (w/w). Samples mixed with His-TEV Protease were subsequently dialyzed overnight (O/N) at 4°C in a SnakeSkin™ Dialysis Membrane (MWCO 10 kDa, Thermofisher, 68100) against Buffer C (20 mM K_2_HPO_4_/KH_2_PO_4_ pH 8, 100 mM Na_2_SO_4_, 100 mM NaCl, 10 mM Imidazole, 5% Glycerol), without any agitation. On the following day, the sample was applied onto a HisTrap HP 1 mL (Cytiva), previously equilibrated Buffer C, through a 5 mL or 10 mL loop. Cleaved proteins eluted as unbound fraction, while His-Tagged components (His-Tag, un-cleaved protein and His-TEV Protease) were retained in the column and thus removed by washing with 5 CVs of Buffer D (20 mM K_2_HPO_4_/KH_2_PO_4_ pH 8, 100 mM NaCl, 100 mM Na_2_SO_4_, 500 mM Imidazole, 5% Glycerol). Flow-through was then concentrated to 1-0.5 mL using a Vivaspin 20 centrifugal concentrator - 10 kDa Molecular Weight Cut Off (MWCO, Sartorius) at 20°C, and applied onto different size exclusion chromatography columns depending on the purified complex (Δ97-RAD51/BRC4 with a Superdex 75 Increase 10/300 GL, Cytiva, RAD51/BRC4, RAD51 /BRC2, RAD51/BRC1-2, RAD51/BRC2-3, RAD51/BRC3-4 with a Superdex 200 Increase 10/300 GL, Cytiva while RAD51/BRC2-4 and RAD51/BRC1-4 with a Superose 6 Increase 10/300 GL, Cytiva) previously equilibrated with Buffer E (20 mM K_2_HPO_4_/KH_2_PO_4_ pH 8, 100 mM NaCl, 200 mM Li_2_SO_4_, 5% Glycerol, 1 mM DTT) and with a flow set to 0.5 mL/min. Samples were snap frozen in liquid nitrogen and finally stored at -80°C. For the purification of ^15^N Δ97-RAD51/BRC4 and ^15^N RAD51/BRC4 the composition of Buffer E was modified removing glycerol and DTT (final composition 20 mM K_2_HPO_4_/KH_2_PO_4_ pH 8, 100 mM NaCl, 200 mM Li_2_SO_4_) and eluted samples were directly utilized for either SLS, or NMR experiments, avoiding freezing in liquid nitrogen. Monomeric His-RAD51 [F86E, A89E] and His-RAD51 WT were purified as previously described by our group (7,19)

### Peptide and CAM833 synthesis

BRC repeat peptides BRC1, BRC2, BRC3, BRC4, BRC7 and BRC8 were synthesized by Thermofisher Scientific, while their biotinylated version harboring a Biotin-aminohexanoic acid moiety at the N-terminus, unmodified BRC5, BRC6 and fluorinated peptides were synthesized by Peptide Protein Research Ltd. Table **S20** report a summary of the peptide aminoacidic composition and design. All peptides were purified with HPLC to 95% purity. CAM833A was synthesized in house as already reported by Scott et al.(30) to 99 % purity as confirmed by UPLC-MS and NMR: ^1^H NMR (400 MHz, DMSO-d_6_) δ 8.96 (h, J = 5.7, 5.2 Hz, 1H), 8.81 (d, J = 7.7 Hz, 0.3H), 8.62 – 8.57 (m, 1H), 8.41 (d, J = 7.7 Hz, 0.7H), 8.28 – 8.14 (m, 2H), 7.94 (dd, J = 9.4, 2.9 Hz, 1H), 7.88 – 7.77 (m, 1H), 7.37 (d, J = 8.6 Hz, 0.3H), 7.30 (d, J = 8.6 Hz, 0.7H), 7.01 (d, J = 2.6 Hz, 0.3H), 6.96 (d, J = 2.6 Hz, 0.7H), 6.94 (d, J = 2.6 Hz, 0.3H), 6.87 (dd, J = 8.7, 2.7 Hz, 0.7H), 5.25 – 5.16 (m, 1H), 5.10 (q, J = 7.2 Hz, 0.7H), 4.61 (t, J = 7.5 Hz, 0.3H), 4.43 (t, J = 7.7 Hz, 0.7H), 4.35 (m, 1H), 4.27 (s, 0H), 4.23 – 4.12 (m, 1H), 3.88 (dd, J = 16.7, 5.5 Hz, 0.3H), 3.76 (s, 1H), 3.73 (s, 2H), 3.71 – 3.38 (m, 2H), 2.25 (s, 0,3H), 2.09 – 2.01 (m, 0.7H), 1.79 (ddd, J = 12.5, 7.2, 4.8 Hz, 1H), 1.39 (d, J = 6.9 Hz, 1H), 1.31 (d, J = 7.0 Hz, 2H) m/z 529.1 (M+H)+.

### Ligand interaction diagram and structural figures preparation

Ligand interaction diagram was generated through Maestro (Schrodinger suite 2022−2). Figures for protein structures have been prepared using PyMOL software. Following PDB entries 1n0w(8) and 6tw9(30) were utilized to prepare figures displayed in the supplementary material.

### Fluorescence Polarization (FP)

FP assays were performed as already described by our group(7). Experiments involving His-RAD51 [F86E, A89E] were performed with buffer containing 20 mM Hepes pH 8.00, 100 mM Na_2_SO_4_, 0.1 mM EDTA, Glycerol 5%. Fluorescence data were acquired at room temperature using a Tecan Infinite 200 (excitation wavelength 485 nm, excitation bandwidth 20 nm; emission wavelength 535, emission bandwidth 20 nm). Competitive binding experiments were carried out by incubating, for 30 minutes at room temperature, 10 nM Alexa Fluor 488-labelled BRC4 peptide and a constant concentration of His-RAD51 [F86E, A89E] to give approximately 80% – 90% saturation of binding in direct binding experiments (40 nM)(7). The competitor (unlabeled BRC repeat peptide) was then added to the protein - labelled peptide mixture and incubated for an additional 30 minutes at room temperature. All competitors were dissolved 100% DMSO to prepare an initial stock at 2.084 mM. The final DMSO concentration in all samples was kept constant at 5 % as well as in reference wells. The final highest competitors’ concentration tested for BRC1, BRC2, and BRC3 was 20 μM while for BRC6, BRC7 and BRC8 the final concentration for competition tests was 100 μM. Competitive binding curves were recorded and fitted with GraphPad Prism 10 using Ki equation. Each FP measurement was the average of three independently measured binding curves.

### Isothermal Titration Calorimetry

ITC experiments were carried out using a MicroCal PEAQ-ITC (Malvern, UK) at 25 °C. RAD51 [F86E, A89E] buffer was exchanged with a buffer containing 20 mM Hepes pH 8.00, 100 Na_2_SO_4_, 0.1 mM EDTA and 5% Glycerol by utilizing a Amicon Ultra – centrifugal filter - 10 kDa Molecular Weight Cut Off (MWCO). BRC7, BRC8 were directly dissolved in the same buffer protein buffer and titrated into RAD51 [F86E, A89E]. BRC6 was dissolved in MilliQ water (stock solution 2 mM) and then diluted in ITC buffer. The protein buffer was then adjusted to avoid heat exchange induced by buffer mismatch by adding MilliQ water to the protein. BRC1, BRC2, and BRC3 were initially dissolved in 100% DMSO (stock concentration BRC1 4.168 mM, BRC2/BRC3 2.084 mM) and then diluted with ITC buffer to obtain a final concentration of 4.8% DMSO. RAD51 [F86E, A89E] was buffer matched accordingly to reach the same DMSO concentration and avoid heat exchange induced by buffer mismatch. The peptides were titrated into RAD51 [F86E, A89E] with an initial 0.4 μL injection and 18 injections of 2 μL. Different peptides’ and RAD51 [F86E, A89E] monomer concentrations were utilized for binding experiments depending on the studied peptides as reported in every figure. Data were fitted to a single-site binding curve with the MicroCal PEAQ-ITC Analysis software and graphed using GraphPad Prism 10.

### Evaluation of defibrillation effect driven by BRC repeats on RAD51 WT fibrils

25 μM of RAD51 WT were incubated at 25°C for 1 hour on the thermoblock in the presence or absence of a 4-fold higher stoichiometric excess of BRC repeats peptides previously dissolved in 100% DMSO at 2.084 mM concentration (final concentration peptide concentration 100 μM, 4.8% DMSO). After incubation, samples were spun down at 13300 rpm at 4°C in a Thermo Scientific MicroCL 17R Microcentrifuge and applied onto a Superdex 200 Increase 10/300 GL, (Cytiva) previously equilibrated with 20 mM Hepes (pH 8.00), 250 mM KCl, EDTA 0.1 mM, glycerol 10%.

### Mass Photometry

Silicon buffer gaskets were attached to the standard microscope coverslips glass slides. Then, the slides were mounted on a TwoMP mass photometer (Refeyn) to carry out the experiments. Prior to each measurement the gasket was filled with 18 µL buffer and the focal plane was automatically estimated. For each measurement of the co-expressed and co-purified protein complexes intermediate dilution were prepared in the assay buffer (20 mM K₂HPO₄/KH₂PO₄ pH 8, 100 mM NaCl, 200 mM Li_2_SO_4_, 1 mM DTT) immediately before measurement to obtain a 10-fold higher concentrated stock. Then, proteins were diluted 1:10 into the assay buffer-filled gasket to reach the desired concentrations and carefully resuspended to homogenize the concentration in the drop. 60 second movies were then recorded through AcquireMP (Refeyn). Data were processed using the DiscoverMP Program. Mass distributions were plotted with DiscoverMP and mean mass peaks determined by Gaussian fitting. To evaluate the disassembly of different BRC repeats peptides 7.5 µM His-RAD51 WT were thoroughly mixed with 30 µM of each peptide (solubilized in either 100% DMSO for BRC1, BRC2, BRC3, BRC4, BRC7 or MilliQ water for BRC8 as 2.084 stock solutions, final peptide/protein stoichiometric ratio 4:1) and incubated for 30 minutes at 20 °C in assay buffer containing 20 mM Hepes pH 8.00, 250 mM KCl, Glycerol 5%. As a negative control His-RAD51 WT was incubated only with buffer containing DMSO. 2 µL of the prepared mixtures were diluted into 18 µL of the assay buffer-filled gasket and carefully resuspended to homogenize the concentration in the drop. 60 second movies were recorded through AcquireMP (Refeyn) with small field mode. Movies were processed using the DiscoverMP Program. Finally, to evaluate the concentration-dependent oligomeric states of His-RAD51 WT different 10X stocks were prepared at 3 µM, 1.5 µM, 0.75 µM, 0.375 µM, 0.188 µM, 0.093 µM in assay buffer containing 20 mM Hepes pH 8.00, 250 mM KCl, Glycerol 5% by performing a serial dilution. Measurements were performed by diluting 2 µL of the concentrated solutions into 18 µL of the same assay buffer. 60 second movies were recorded through AcquireMP (Refeyn) with regular field mode and processed using the DiscoverMP Program. For all reported experiments, contrast-to-mass calibration was performed with an internal mixture of standards including known-mass proteins (Bovine Serum Albumin (BSA), Apoferritin and Bovine Thyroglobulin).

### Microscale Thermophoresis (MST)

His-RAD51 WT was labelled with the Monolith His-Tag Labeling Kit RED-tris-NTA 2nd Generation kit (NanoTemper Technologies, München, Germany), which specifically recognizes the hexahistidine tail of the recombinant protein. A 4.2 µM dye solution was prepared by suspending the dye in 30 µL of PBS-T. The obtained solution was further diluted into 720 µL of labelling buffer (20 mM Hepes pH 8.00, 200 mM KCl, 5% Glycerol) to obtain a final 168 nM dye solution. His-RAD51 WT was then diluted to 200 nM, mixed 1:1 with the 168 nM dye solution and incubated for 30 minutes at room temperature in the dark. The obtained mixture was then centrifuged at 13300 rpm and 4 °C in a Thermo Scientific MicroCL 17R Microcentrifuge, to remove aggregates. BRC1, BRC2 and BRC3 peptides were dissolved in 100% DMSO to obtain 1 mM stock then serially diluted in 100% DMSO to obtain 16 concentrations. Each sample at different concentrations was then individually diluted 1:10 in assay buffer (containing 20 mM Hepes pH 8.00, 200 mM KCl, 5% Glycerol, 0.05% v/v Tween-20, 0.1% w/v PEG-8000, 0.5 mM Sodium Deoxycholate) to obtain 10% DMSO intermediate dilution. Finally, prior to each measurement each sample dilution was mixed at 1:1 with the labelled His-RAD51 WT thus allowing to obtain a constant 5% DMSO concentration in each sample. Samples were incubated for 5 minutes, spun down at 13300 rpm to remove eventual aggregates. The final highest peptides’ concentration tested was 50 μM. MST measurements were performed at 1% excitation and high power on 16 premium capillaries (NanoTemper Technologies, München, Germany) containing a constant concentration (50 nM) of labelled His-RAD51 WT and 16 different concentrations of the BRC1, BRC2, BRC3 peptides. Four replicates for each binding curve were fitted using the Affinity Analysis software of Nanotemper Technologies and analyzed at 2.5 seconds to obtain binding affinity data. Results were then re-graphed using GraphPad Prism 10.

### Negative Staining – Trasmission Electron Microscopy experiments

His-RAD51 WT at 2.6 µM was incubated at 4°C for 2 hours with 6 different stoichiometric ratios of BRC1, BRC2 and BRC3 (peptide/protein ratio 0:1, 0.4:1, 0.6:1, 0.8:1, 1:1, 2:1), as already described by our group(19). Each peptide was initially solubilized as a 1 mM stock in 100% DMSO and diluted to have a final 5% v/v DMSO concentration for all studied stoichiometric ratios. 10 μL of samples were adsorbed for 60 s on plasma-cleaned 150 mesh ultrathin carbon coated copper grids (Electron Microscopy Sciences, Hatfield, PA, USA). After a washing step with buffer 20 mM Hepes pH 8.00, 250 mM KCl, 0.1 mM EDTA, the grids were negatively stained with a solution of uranyl acetate (1% in distilled water) for 60 s. Finally, the grids were analyzed in bright field mode using a JEOL JEM-1011 (JEOL, Japan) transmission electron microscope equipped with a thermionic source (W filament) operating at an acceleration voltage of 100 kV and with a Gatan Orius SC1000 series charge coupled device (CCD) camera (4008 x 2672 active pixels). For each sample, the length of 300 imaged fibrils were measured and analyzed by ordinary one-way ANOVA using Graphpad Prism 10.

### Static Light Scattering

Static light scattering (SLS) analyses were performed on a Viscotek GPCmax/TDA (Malvern, UK) instrument, connected in tandem with a series of two TSKgel G3000PWxl size-exclusion chromatography columns (Tosoh Bioscience) as already described in (7). For all the experiments the system was equilibrated with buffer containing 20 mM K_2_HPO_4_/KH_2_PO_4_ pH 8, 100 mM NaCl, 200 mM Li_2_SO_4_, 1 mM DTT, 2% sucrose. Flow was set at 0.4 mL/min. Un-labelled samples were injected at the following concentrations Δ97-RAD51/BRC4 = 3.30 mg/mL, RAD51/BRC4 = 4.03 mg/mL, RAD51/BRC2 = 2.00 mg/mL, RAD51/BRC1-2 = 2 mg/mL, RAD51/BRC2-3 = 2.24 mg/mL RAD51/BRC3-4 = 2.73 mg/mL, RAD51/BRC2-4 = 3.03 mg/mL, RAD51/BRC1-4 = 1.43 mg/mL. ^15^N labelled Δ97-RAD51/BRC4 and RAD51/BRC4 samples were run at the same flow rate in buffer containing 20 mM K_2_HPO_4_/KH_2_PO_4_ pH 8, 100 mM NaCl, 200 mM Li_2_SO_4_ and injected suddenly after the SEC elution, prior to NMR relaxation experiments, at 7.35 mg/mL and 5.14 mg/mL, respectively. Data analysis was performed using Viscotek software, calibrating the instrument with Bovine Serum Albumin (BSA) at 5 mg/mL. Data were exported as .csv files and re-graphed using GraphPad Prism 10.

### Analytical Size Exclusion Chromatographies

Analytical SEC analyses samples were performed by exploiting an Akta Micro system with flow set at 0.1 mL/min and loading 50 µL of each sample onto a Superdex® 200 5/150 GL (Cytiva) equilibrated with buffer containing 20 mM K_2_HPO_4_/KH_2_PO_4_ pH 8, 100 mM NaCl, 200 mM Li_2_SO_4_, 1 mM DTT, 2% sucrose. The following concentrations of each sample Δ97-RAD51/BRC4 = 1 mg/mL, RAD51/BRC4 = 1 mg/mL, RAD51/BRC2 = 2.00 mg/mL, RAD51/BRC1-2 = 0.8 mg/mL, RAD51/BRC2-3 = 0.44 mg/mL, RAD51/BRC3-4 = 1.1 mg/mL, RAD51/BRC2-4 = 1 mg/mL, RAD51/BRC1-4 = 1.43 mg/mL. A standard solution of globular markers (BioRad, Gel Filtration Standards #1511901) was diluted 1:20 in the same SEC buffer and 50 µl were injected in the column to be utilized as a reference. Eluted species were monitored by absorbance at 215 nm.

### Biolayer Interferometry

Biolayer interferometry (BLI) experiments were performed utilizing an Octet K2 system (Sartorius). All BLI experiments were carried out in an assay buffer containing 20 mM Hepes pH 8.00, 100 mM Na_2_SO_4_, 5% glycerol, 0.05% Tween 20, 0.1% PEG8000 and 0.5 mM Sodium Deoxycholate. The following protocol was applied: 60 sec baseline, 240 sec loading, 240 sec baseline, 180 sec association, 180 sec dissociation. For every step, shaking at 1000 rpm was enabled and temperature set at 25°C. Streptavidin Octet biosensors (18-5019, Sartorius) were initially dipped into assay buffer to record an initial baseline of 60 seconds. BioBRC repeats (BRC1, BRC2, BRC3, BRC7, BRC8) were solubilized in 100% DMSO at a 2 mM concentration, diluted at a final 1 µM concentration in assay buffer and then immobilized to the Streptavidin sensor through 240 seconds loading step. After the loading stage, a second baseline of 240 seconds was recorded in wells containing only assay buffer to verify the stability of signal and remove unbound peptide. For each experiment, sensors were subsequently dipped for 180 seconds into wells containing His-RAD51 [F86E, A89E] (at two different concentrations 460 nM, 230 nM) to measure association signals and finally moved to wells containing only assay buffer to assess complex dissociation. Two replicates were run for each His-RAD51 [F86E, A89E] concentration. BLI experiments were carried out including a double reference: a reference well (where only immobilized the biotinylated peptide was present on the sensor and no analyte (0 nM) during association) and reference sensor (where no biotinylated peptide was immobilized on the sensor and His-RAD51 [F86E, A89E] concentration was matched during association). Data were analyzed using Octet Analysis Studio 12.2 subtracting to recorded sensorgrams the signals of both the reference well and reference sensors to remove signals due to a-specific binding. Recorded sensorgrams were corrected by aligning to the average of the second baseline steps, applying Savitzky–Golay filtering and inter-step correction based on the second baseline step. To calculate Rmax, the response at the end of the association phase (170-175 sec) was extrapolated. All data presented data were exported to .csv files and re-graphed using GraphPad Prism 10. Multiple comparisons between calculated Rmax were carried out through ordinary one-way Anova test.

### Cross-linking: sample preparation and reaction conditions set up

For crosslinking with bis(sulfosuccinimidyl)suberate (BS3), purified His-RAD51[F86E, A89E] concentration was adjusted to 24 µM (=0.98 mg/mL) and mixed with an equimolar concentration of biotinylated BRC repeat peptides (previously solubilized as a 2 mM stock in 100% DMSO) with a final DMSO concentration equal to 1.2 % (v/v). The mixture was incubated for 30 minutes at 20°C on a thermoblock to allow complex formation and then crosslinked with a final concentration of 1 mM BS3 (previously solubilized in MilliQ water as a 50 mM stock). After 30 minutes at the same temperature, 4x Laemmli Sample Buffer (BioRad #1610747) was added to crosslinked samples and boiled at 95°C for 5 minutes. For crosslinking with endogenously expressed complexes, the protein concentration was adjusted to 2 mg/mL and then crosslinked with a final BS3 concentration of 1 mM. 4x Laemmli Sample Buffer supplemented with 100 mM DTT was added to crosslinked samples and boiled at 95°C for 5 minutes.

### Coomassie Blue and Western Blot analyses

Protein samples were loaded onto 4–15% Criterion™ TGX™ Precast Midi Protein Gel (18 well, 30 µl #5671084) Precast gels and electrophoretic runs were performed in Tris-Glycine SDS Buffer. Gels were stained using Page Blue Protein Staining solution (Thermo Fisher, 24620) according to manufacturer protocols. Images were acquired using a Gel Doc EZ Imager (Biorad). For Western Blots (WB), proteins run on SDS-PAGE gels were then transferred to TransBlot Turbo Midi-size nitrocellulose membranes (Biorad) using a Transblot Turbo apparatus (Biorad, set at 25V, 1.0A, 30 minutes). Membranes were blocked for 1 h at room temperature in 5% milk PBS-T and incubated overnight at 4°C in the same buffer containing primary antibodies (1:200 dilution: mouse anti-His (Sigma-Aldrich, SAB4301134); 1:5000 dilution rabbit anti RAD51 (Abcam, ab63801)). Membranes were washed three times in PBS-T and incubated with secondary antibodies (1:10000 dilutions Goat Anti-Mouse HRP or Goat Anti-Rabbit HRP, Biorad) in blocking buffer for 1 h at room temperature. After three washes in PBS-T, chemiluminescence was detected using the Clarity Western ECL substrate (Biorad, 1705061). Images were recorded using a ChemiDoc MP Imaging System (Biorad). To evaluate the efficiency of cross-linking reactions membranes were incubated for 1 hour at room temperature with Streptavidin HRP to detect the biotin moiety of bioBRC repeats (1:3000 dilution in 3% BSA-TBS-T). After three washes in TBS-T, chemiluminescence was detected using the Clarity Western ECL substrate (Biorad, 1705061), and images were recorded using a ChemiDoc MP Imaging System (Biorad). Different amounts of RAD51 were loaded in the WBs due to different sensitivity of the antibodies.

### LC-MS analysis

Crosslinking samples in Laemmli Sample Buffer were digested following the S-Trap protocol (Protifi) with minor adaptations and purified using solid-phase extraction(43). Briefly, a sample containing 100 µg of proteins was diluted to a final volume of 100 µl with 50 mM ammonium bicarbonate (ABC) and proteins were reduced by addition of 4 µl TCEP for 30 min at 37°C with mild agitation. Cysteine alkylation was performed with 8 µl iodoacetamide (IAA) for 30 min in the dark with mild agitation. The samples were then acidified with 12 µl 12% phosphoric acid, diluted with 750 µl of the S-Trap binding buffer (1 M TEAB Buffer and Methanol (10:90)) and transferred to S-Trap mini columns (Protifi). SDS removal was conducted by three washing cycles with 400 µl S-Trap binding buffer prior to digestion. MS-grade Trypsin (Serva) was added in an enzyme to protein ratio 1:50 using 125 µl digestion buffer (50mM ABC, pH 8,5) as media and the column incubated overnight at 37°C with light agitation. Peptides were eluted stepwise in 80 µl 50mM ABC, 0,2% formic acid, 50% acetonitrile (ACN). The pooled fractions were diluted 1:1 with 0,1% trifluoroacetic acid (TFA) and desalted with C18-SPE cartridges (Biotage). After equilibration with 2 ml ACN, 1 ml 50% ACN/1% acetic acid and 2 ml 0,1% TFA the samples were loaded onto the cartridge, washed with 2 ml 0,1% TFA and eluted with 1 mL 80% ACN/0,1% TFA. The eluted fractions were dried using an Eppendorf concentrator (Eppendorf) and stored at -20°C before analysis. Dried peptides were reconstituted in 5% ACN with 0.1% formic acid (FA). Peptides were loaded onto an Acclaim PepMap C18 capillary trapping column (particle size 3 µm, L = 20 mm) and separated on a ReproSil C18-PepSep analytical column (particle size = 1,9 µm, ID = 75 µm, L = 25 cm, Bruker Coorporation, Billerica, USA) using a nano-HPLC (Dionex U3000 RSLCnano) at a temperature of 55°C. Trapping was carried out for 6 min with a flow rate of 6 μL/min using a loading buffer composed of 0.05% trifluoroacetic acid in H_2_O. Peptides were separated by a gradient of water (buffer A: 100% H_2_O and 0.1% FA) and acetonitrile (buffer B: 80% ACN, 20% H_2_O, and 0.1% FA) with a constant flow rate of 400 nL/min. The gradient went from 4% to 48% buffer B in 45 min. All solvents were LC-MS grade and purchased from Riedel-de Häen/Honeywell (Seelze, Germany). Eluting peptides were analyzed in data-dependent acquisition mode on an Orbitrap Eclipse mass spectrometer (Thermo Fisher Scientific) coupled to the nano-HPLC by a Nano Flex ESI source. MS1 survey scans were acquired over a scan range of 350–1400 mass-to-charge ratio (m/z) in the Orbitrap detector (resolution = 120k, automatic gain control (AGC) = 2e5, and maximum injection time: 50 ms). Sequence information was acquired by a “ddMS2 OT HCD” MS2 method with a fixed cycle time of 5 s for MS/MS scans. MS2 scans were generated from the most abundant precursors with a minimum intensity of 5e3 and charge states from two to eight. Selected precursors were isolated in the quadrupole using a 1.4 Da window and fragmented using higher-energy collisional dissociation (HCD) at 30% normalized collision energy. For Orbitrap MS2, an AGC of 5e4 and a maximum injection time of 54 ms were used (resolution = 30k). Dynamic exclusion was set to 30 s with a mass tolerance of 10 parts per million (ppm). Each sample was measured in duplicate LC-MS/MS runs. MS raw data were processed using the MaxQuant software (v2.5.2.0) with customized parameters for the Andromeda search engine(44). Spectra were matched to a FASTA file containing the BRCA2 and RAD51 sequences downloaded from UniProtKB (October 2023), a contaminant and decoy database. The RAD51 sequence was modified to include the His-Tag on the N-terminus (MGSSHHHHHHSSGLVPRGSHMLEDP-). A minimum tryptic peptide length of seven amino acids and a maximum of two missed cleavage sites were set. The crosslinker BS3 was chosen from the default list of available crosslinkers. Precursor mass tolerance was set to 4.5 ppm and fragment ion tolerance to 20 ppm, with a static modification (carbamidomethylation) for cysteine residues. Acetylation on the protein N-terminus and oxidation of methionine residues were included as variable modifications. A false discovery rate (FDR) below 1% was applied at crosslink, peptide, and modification levels. Search results were imported to xiVIEW for subsequent analysis. After stringent manual curation of the spectra, only inter-protein crosslinks with spectral scores above 30 were selected for fitting on the structures(45). Crosslinks were fitted on the models generated through Alphafold 2.3 or Alphafold 3, using xiVIEW(45).

### NMR experiments

All the NMR experiments were recorded at 298°K with a Bruker FT NMR Avance Neo 600-MHz spectrometer equipped with a 5-mm CryoProbe™ QCI ^1^H/^19^F-^13^C/^15^N-D-Z quadruple resonance with shielded z-gradient coil, and the automatic sample changer SampleJet™ with temperature control.

In ^19^F-NMR binding experiments, for all samples before the ^19^F NMR spectra, was recorded a 1D ^1^H experiment. The binding of the ^19^F-peptides was evaluated by ^19^F T_2_ filter NMR experiments(46,47), testing them at 50 μM, in the absence and the presence of 0.96 μM RAD51 [F86E, A89E] monomer in 20 mM Hepes pH 8, 100 mM Na_2_SO_4_, 2 mM DTT, 10% D_2_O (for the lock signal), 10 μM EDTA, 1% DMSO-*d_6_,* and 1% Glycerol. To confirm the binding site of the ^19^F-peptides to the RAD51 [F86E, A89E] the same ^19^F T_2_ filter experiments were run in competition manner using as competitor the two known RAD51 binders BRC4 and CAM833A. In these experiments the ^19^F peptides were tested at two different concentration (50 and 20 μM) in the absence and in the presence of 0.96 μM of RAD51 [F86E, A89E] and in the presence of RAD51 [F86E, A89E] + 1.92 μM BRC4 or + 10 μM CAM833A. As proposed by Dalvit and colleagues(48), knowing the dissociation constant (K_d_) of the competing molecule (I) and its concentration, it is possible to calculate the K_d_ of the different ^19^F-peptides, by the equation:

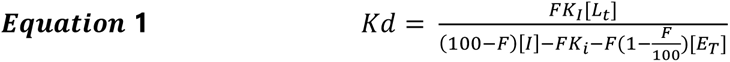

where K_d_ is the dissociation constant of the observed ^19^F-peptide (spy), F is its displacement value in the presence of a competing molecule I (BRC4 or CAM833A) expressed in %, [L_T_] is the spy concentration, [I] is the concentration of the competing molecule, K_I_ is the dissociation constant of I, and [E_T_] is the concentration of the protein (RAD51 [F86E, A89E]). The ^19^F competition binding experiments were carried out in triplicate. The % of displacement F has been calculated measuring the integral of the ^19^F signal of the ^19^F-peptides, alone (Int_L_) in the presence of protein (Int_L+E_) and in the presence of protein and competitor (Int_L+E+I_) using the following equation:

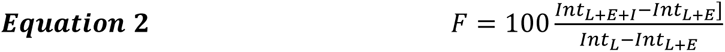

The K_d_ of the ^19^F-pept was obtained by the average of the K_d_s obtained using the two competitors. The 1D ^1^H experiments were recorded using the standard NOESY (nuclear Overhauser effect spectroscopy) presat “noesygppr1d” Bruker pulse sequence, with 64k data points, 30 ppm of spectral width (sw), 128 scans, 10 ms of mixing time (d8) and 1.835 s and 4s of acquisition time (aq) and relaxation delay (d1), respectively. The data were multiplied by an exponential function of 0.3 Hz prior to Fourier transformation. ^19^F T_2_ filter NMR experiments were recorded using the Bruker cpmgigsp sequence using d1 of 5 s, proton decoupling during the acquisition period and 128 scans and a spin-echo scheme (cpmg)(49,50) with total τ = 0, 0.8, 0.120 s for the samples containing 50 μM ^19^F-peptides, and 384 scans and τ = 0.064, 0.096,s for that with 20 uM of ^19^F-peptides. The data were multiplied by an exponential function of 1 Hz prior to Fourier transformation and the ^19^F chemical shift was referred to that of the CFCl_3_.

For NMR protein characterization by ^1^H -^15^N HSQC experiments, the ^15^N labeled RAD51/BRC4 and Δ97-RAD51/BRC4 were tested in 20 mM K_2_HPO_4_/KH_2_PO_4_ pH 8, 100 mM NaCl, 200 mM Li_2_SO_4_,10% D_2_O (for the lock signal). All the experiments were recorded at 298°K. For both proteins were recorded a 1D ^1^H spectrum (noesygppr1d) with 64 scan, d1 of 4s, sw of 30 ppm, aq of 1.835 s, 64k datapoints and d8 of 10 ms. The ^1^H-^15^N Heteronuclear Single Quantum Coherence (^1^H-^15^N HSQC)(51,52) spectra were recorded using the Bruker hsqcfpf3gpphwg sequence with 48 scans, a time domain size of 2048x256, d1 of 1.2 s, sw of 16 and 40 ppm for ^1^H and ^15^N respectively, CNST4 (J(YH)) of 90 Hz

### Size Exclusion Small Angle X-Ray Scattering, molecular modelling, and structural analysis

Batch and SEC SAXS experiments reported in this work were performed at the B21 beamline of Diamond Light Source Synchrotron (DLS, Didcot, UK)(53). Detailed data collection parameters (e.g. concentration of batch SAXS samples, buffers etc.) are listed in Tables **S11-S19**. Depending on the analyzed complex, samples were injected into a Superdex 200 3.2/300 or Superose 6 Increase 3.2/300 column, pre-equilibrated with running buffer and flow set to 0.075 mL/min to attenuate radiation damage. SEC-SAXS data were analyzed using Scatter(54). Scattering frames, corresponding to samples, were selected, averaged, and subtracted from averaged buffer frames. The buffer-subtracted, 1D scattering curves were then processed using BioXTAS RAW to compute the radius of gyration by Guinier analysis, the dimensionless Kratky Plot and to obtain molecular mass estimates (Tables **S11-S19**)(55). The Porod Volume, the pair distribution function (P(r)) and the maximum particle dimension (Dmax) were calculated using PRIMUS and the GNOM algorithm(56,57). Data were exported in .csv format and re-graphed using GraphPad Prism 10. To estimate data ambiguity and the last optimal point containing useful structural information of SAXS data the algorithms AMBIMETER and SHANUM, available within the ATSAS suite, were respectively applied(58,59). Based on the outputs of the Guinier analysis and of the SHANUM algorithm, SEC-SAXS curves were truncated accordingly to remove the points at low q-range and high q-ranges not containing useful structural information. Generated scattering curves, were utilized for subsequent atomistic modelling, exploiting models generated through AlphaFold(32,60). Single model agreement with experimental data was assessed through FoXS(61,62). To generate multistate models, flexibility was applied to residues of unstructured regions and modelled through MultiFoXS(61,62). Details on the input and output parameters are reported in Tables **S11-S19**. SAXS data and relative fittings of single and multi-state models were exported in .csv format and re-graphed using GraphPad Prism 10. Residuals were calculated for each fitting by applying the formula reported by Trewhella et al.(63). All data presented in this study have been deposited to SASBDB and are freely available(64,65).

## RESULTS

### Computational investigations of the BRC repeats-RAD51 interaction

To investigate the structural features of the BRC-repeats, we initially challenged Alphafold2.3 (AF2) and Alphafold3 (AF3) to predict the structure of the full-length BRCA2 (Figure **S2**). However, the resulting models suggested a largely disordered protein, with low confidence scores across most regions, except for the DNA-binding domain (Figure **S2**), which showed higher structural reliability. This result was most likely due to the limited availability of high-quality templates in the PDB and the poor evolutionary conservation of BRCA2 across species (Figure **S3**).

Notably, analysis of the BRC-repeat region consistently predicted it as fully unstructured with low confidence metrics (Figure **S2**). Interestingly, when simulations were repeated using only the isolated BRC1-8 region, an improvement in peptide folding was observed (Figure **S2**). However, despite this enhancement, the predicted BRC1-8 region still contained substantial unfolded linkers between individual repeats (Figure **S2**). These disordered interconnecting regions might hinder subsequent modeling of interactions with RAD51, as they are likely to lower interface-predicted template modeling (ipTM) scores(66,67). Given these results, we initially modeled the individual BRC repeats in complex with either an N-terminal truncated RAD51 (Δ97-RAD51), to compare predictions with the available X-ray crystal structure (PDB entry: 1n0w), or with full-length RAD51 (RAD51), to investigate potential interactions between the BRC-repeats and the RAD51 N-terminal domain (Figures **S4-S8**). In both settings, all BRC-repeats binding mode closely matched the experimentally determined structure of BRC4 bound to RAD51 (PDB entry: 1n0w, Figures **S4-S8**). Notably, in all predictions performed with RAD51, we consistently observed a re-arrangement of the RAD51 N-terminus to allow peptide binding (Figures **S6-S8**)(33,34). Moreover, additional simulations were also set up using AF2, in which the BRC4 peptide was modelled in competition with other BRC-repeats for binding to either Δ97-RAD51 or RAD51 (33,34). In most cases, BRC4 was observed to bind RAD51, consistent with the available crystal structure(8), whereas the other peptides were misplaced and associated with low-confidence predictions (Figure **1**, and Figures **S9-S37**). For instance, BRC5 and BRC6 were never modeled at the expected binding site (Figure **1**, Figures **S9, S16-S19**, **S30-S33**), and only a small number of predictions positioned BRC3, BRC7, and BRC8 at the putative site of interaction (Figure **1**, Figures **S9**, **S14**, **S15**, **S20**-**S23**, **S28-S29**, **S34-S37**). Interestingly, BRC2 was consistently modeled at the expected binding site in a higher number of predictions compared to BRC4 (Figure **1** and Figure **S9**, **S12-S13**, **S26-S27**). Additionally, the presence of the RAD51 N**-**terminal domain significantly improved the modeling accuracy for BRC1, suggesting its potential role in stabilizing this specific interaction (Figure **1** and Figure **S9**, **S10**, **S11**, **S24**, **S25**). Overall, these results position BRC1, BRC2, and BRC4 as higher-affinity binders to RAD51 than the other peptides.

**Figure 1.**
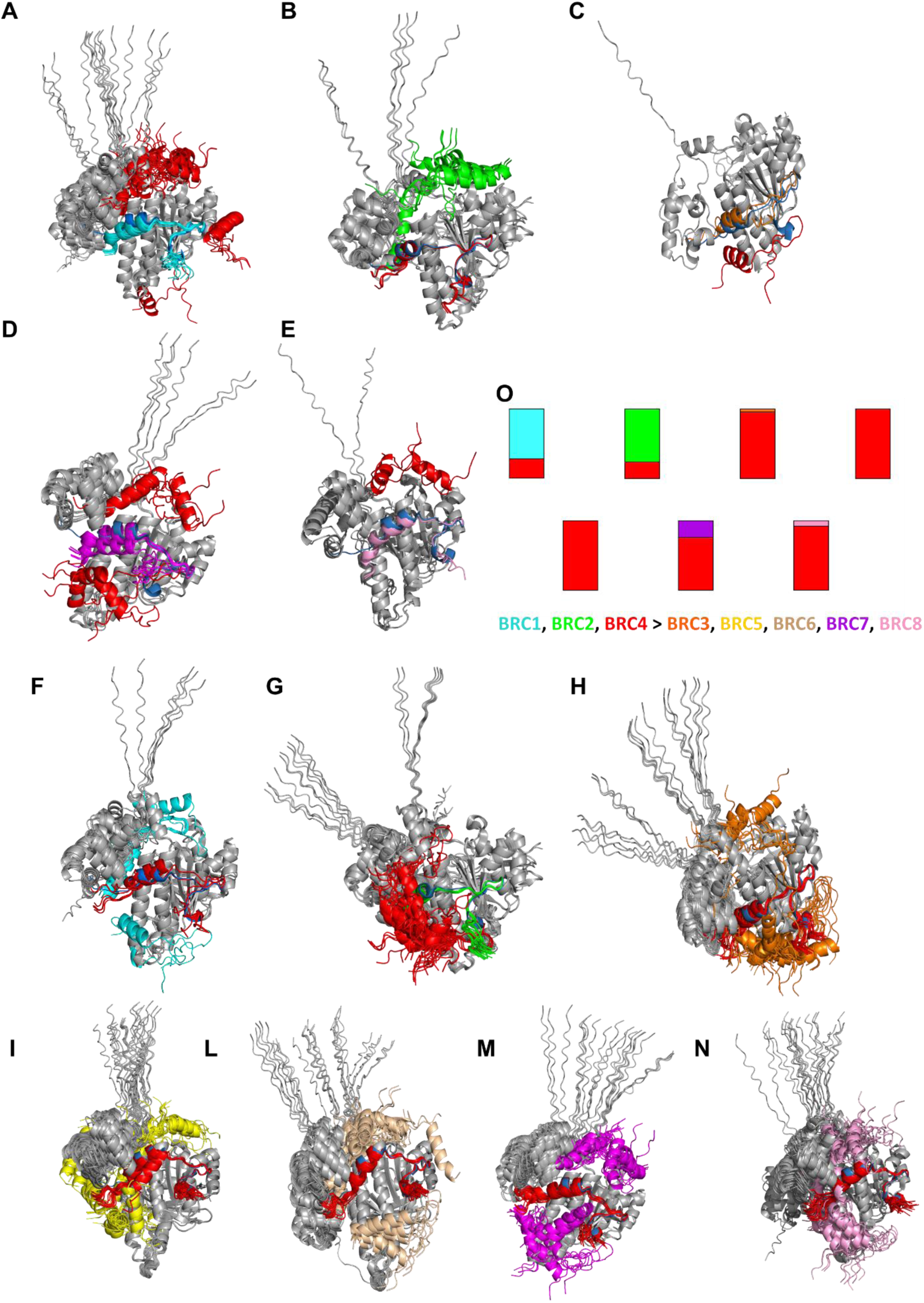
Competitive AlphaFold predictions of the BRC-repeats vs BRC4, using RAD51 FL as receptor. Simulations’ outputs were aligned to the available RAD51-BRC4 complex structure (PDB: 1n0w) in which the experimentally determined BRC4 peptide is shown in blue. Overlay of simulations using RAD51-FL where BRC4 (red) was misplaced while **A.** BRC1 (cyan) **B.** BRC2 (green) **C.** BRC3 (orange) **D.** BRC7 (violet) **E.** BRC8 (pink) were positioned in the BRC4 binding site. Overlay of simulations where the BRC4 (red) peptide is correctly modelled in its binding site and the other BRC-repeats: **F.** BRC1 (cyan) **G.** BRC2 (green) **H.** BRC3 (orange) **I.** BRC5 (yellow) **L.** BRC6 (wheat) **M.** BRC7 (violet) **N.** BRC8 (pink) display a different binding mode. In panel **O.** a summary graph reports the results of competitive simulations between BRC4 and the other BRC-repeats (25 models for each competitive simulation). Predicted Align Error and relative pLDDT graphs are reported in Figures **S10-S23**)

To further rationalize these findings, we employed complementary computational approaches to investigate the structural features underlying the differing affinities of the peptides. Based on the AF prediction, we hypothesized that BRC-peptides interact with RAD51 similarly to BRC4. Initially, we performed a residue scanning analysis (Figure **2A**) to assess the per-residue relative changes in binding affinity compared to BRC4. Three representative peptides were considered for this analysis based on the results of competitive AF2 simulations: BRC2, systematically placed in the correct binding site in a higher number of structures compared to BRC4; BRC7, accurately modeled in a few structures; and finally, BRC5 was never modeled at the expected binding site in any of the competitive simulations. In all these peptides, residue F1524 (FXXA domain), which is crucial for binding, was preserved while mutations at other positions within the FXXA domain appeared only moderately detrimental. The only exception was T1526, unfavorably mutated into S1526 in the BRC2 repeat. Conversely, mutations in the LFDE domain had a more significant impact, particularly around position 1546. Indeed, in BRC5, the mutation of residue F1546, which interacts with RAD51 by protruding into a hydrophobic cavity, was highly disfavored. This interaction was maintained in BRC7, although the mutation of the adjacent L1545 was disfavored. In BRC2, these residues were conserved, with no mutations occurring. However, the mutation of E1548D, with a shorter sidechain located at the end of the LFDE domain, resulted in loss of affinity. Notably, BRC2 displayed a mixture of favorable and unfavorable mutations throughout its sequence relative to BRC4. The mutation of S1528 into H1528 in BRC2 was also investigated, considering that histidine could adopt different tautomeric forms with distinct effects (**Figure S38**). Indeed, the δ tautomer H1528 might preserve the hydrogen bond with D187, like the interaction observed with serine in BRC4 (**Figure S38**). The binding was further strengthened in the protonated H1528 (positively charged), likely due to the formation of a salt bridge. Conversely, the ε tautomer, which could not preserve the hydrogen bond, was disfavored (**Figure S38**). Further insights were obtained by comparing these results with molecular dynamics (MD) simulations. We inspected the Root-Mean-Square Fluctuation (RMSF) of the peptide α carbons as a measurement of peptide flexibility in the simulations. Fluctuations at position 1533 were immediately evident from the plot (Figure **2B**). This residue harbored a sidechain exposed to the solvent, allowing it to explore a wide conformational space and occasionally establish transient contacts with RAD51 residues (Figure **2B**). This was indicated by the higher RMSF values and unfavorable mutations into residues with neutral side chains. At position 1528, almost all peptides displayed consistent behavior (Figure **2B**). Notably, in BRC4 and BRC7, a serine formed a hydrogen bond with RAD51’s D187, while in BRC2, the interaction was maintained by the histidine. BRC5 displayed a different RMSF profile in this region, likely due to the presence of a cysteine, instead of a serine, which determined the loss of a hydrogen bond and increased fluctuations of the residue. Interestingly, towards the end of the MD simulation (in the 400 ns timescale), we observed that BRC5 S1527 rearranged to recapitulate the hydrogen bond with D187. At position 1541, BRC4 and BRC2 carried a lysine, which did not provide any stable interaction with RAD51 (Figure **2B**). By contrast, BRC7 and BRC5 behaved differently. Indeed, BRC7 presented an asparagine in this position and often interacted through a hydrogen bond with the carbonylic oxygen of A1537. Moreover, BRC5 presented an aspartic acid residue, which formed a salt bridge with K1544. These observations suggested a local stabilization of the structure, affecting the adjacent LFDE domains of the BRC5 and BRC7 peptides, resulting in low RMSF profiles. Although the residue scanning analysis did not highlight any effect on the binding affinity, this local stabilization observed in the MD simulations might impact the interaction of the BRC peptides with RAD51 and specifically with its N-terminal domain. MD simulations for position 1528 of BRC2 were conducted only for the δ tautomer. In line with scanning analysis, the delta hydrogen of H1528 provided a hydrogen bond with the side chain of D187, which was maintained during the simulation. Overall, AF analyses suggest that all BRC-repeats can bind to the canonical BRC4 binding site on RAD51, albeit with different affinities. Residue-scanning analyses and MD simulations indicate that variations in aminoacidic composition in proximity of the LFDE domain play a crucial role in modulating the affinity of BRC repeats for RAD51.

**Figure 2.**
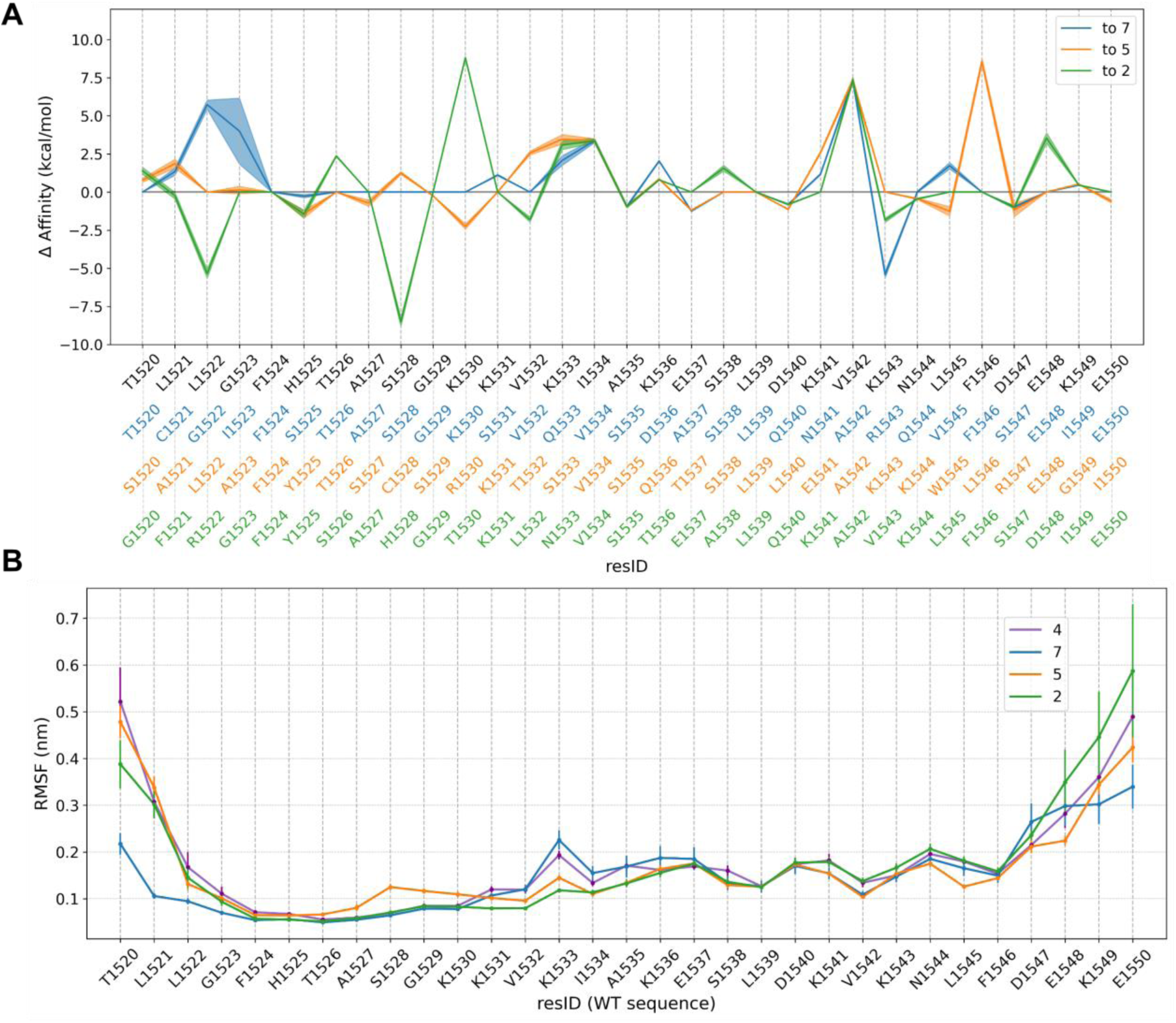
Residue Scanning Analysis and Molecular Dynamics simulations of BRC2, BRC5 and BRC7. **A.** The three curves report the residue-wise change in binding affinity due to mutating the BRC4 peptide into BRC7 (azure), BRC5 (orange), and BRC2 (green), respectively, with shaded areas representing the statistical error. The three mutated sequences are reported on the x-axis with consistent colors, along with the original sequence of BRC4 in black. Negative values indicate an improvement in the binding affinity, while positive values indicate unfavorable mutations. All peptides’ sequences have been aligned to the BRC4 sequence. **B.** Root-Mean-Square Fluctuation (RMSF) from atomistic MD simulations of four different peptides in complex with RAD51 (BRC4, BRC7, BRC5, and BRC2 in purple, azure, orange, and green, respectively). The RMSF is computed on peptide α-carbons, and error bars indicate statistical error.

### Biophysical evaluation of BRC repeats binding to monomeric and fibrillary RAD51

To validate our computational findings, we performed Fluorescence Polarization (FP) and Isothermal Titration Calorimetry (ITC) analyses of the BRC-repeats interaction with RAD51. RAD51 wild type (WT) exhibits an intrinsic tendency to self-assemble into oligomeric structures, which are depolymerized by BRC4(19,68). This behavior has previously posed a limitation for ITC experiments, as it could lead to simultaneous binding and unbinding events, thereby preventing reliable estimation of binding affinities(7,19). To overcome this issue, a monomeric form of RAD51(7) was used here, enabling reliable ITC measurements of RAD51-BRC repeat interactions. For consistency, the same RAD51 form was also employed in the competitive FP experiments (Figures **S39-S41** and Table **1**)(7). In line with previous in silico data, the first three

**Table 1.**
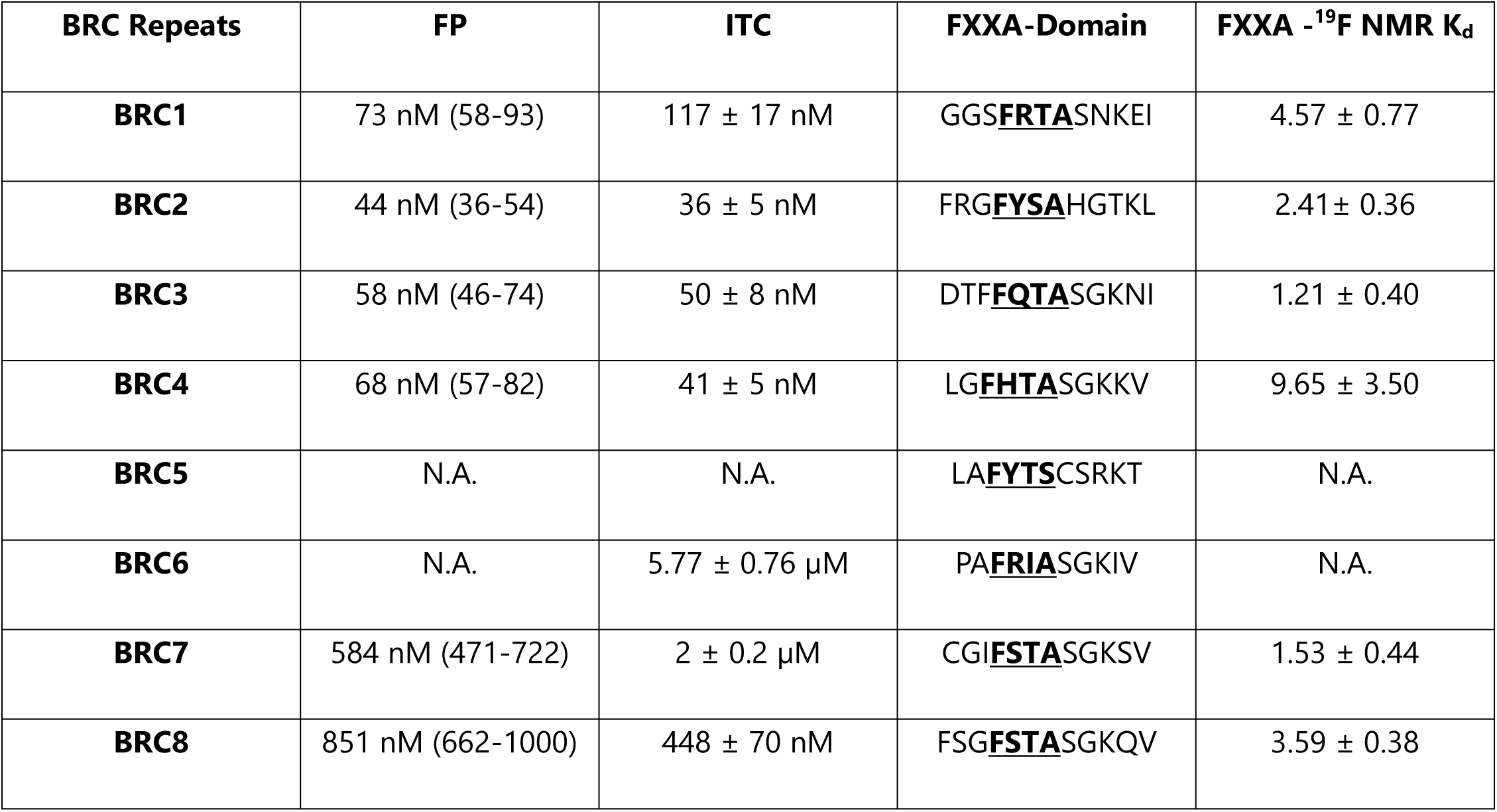
K_d_ values summary of the complete BRC-repeats and of their respective FXXA binding motifs measured by ITC, competitive FP and ^19^F NMR. All experiments were performed using a fully human monomeric RAD51. Affinity data of BRC4 are shown as a reference and can be retrieved in (7). FP represent the mean ± standard deviation from three independent experiments while NMR data represent the average of two independent experiments.

BRC repeats (BRC1 - BRC2 - BRC3) showed high affinity for RAD51 (K_d_ = 50-100 nM), like BRC4(7) (Figures **S39, S40** and Table **1**)(7). By contrast, BRC6-8 displayed lower affinity with a K_d_ ranging from 1 μM to 10 μM (Figures **S39, S41** and Table **1**). Unfortunately, the poor solubility of BRC5 prevented its biophysical characterization. To clarify the role of each conserved motif within the BRC repeats, we also performed NMR competition experiments using fluorinated peptides containing the FXXA domains of peptides displaying the highest affinity for RAD51 (BRC1-3, BRC7-8) in the presence of BRC4 or the small-molecule inhibitor CAM833 (Figures **S42-S48**). NMR analysis revealed both competitors’ displacement of fluorinated peptides, suggesting competition for the same binding site (Figure **S48**) and yielding similar K_d_ values across all peptides analyzed (Figure **S48**, Table **1**). While indirect, these findings provide meaningful evidence that the LFDE domain, rather than the FXXA module, is likely the primary determinant modulating the peptides’ affinities for RAD51 (Table **1**). Subsequently, the interaction of the BRC-repeats with RAD51 WT was characterized to assess the effect of BRC peptides binding on RAD51 WT oligomerization(19). RAD51 WT exists as a heterogeneous mixture of self-assembled oligomeric structures in solution, as observed by Mass Photometry (MP) (Figure **S49**)(7,19). The ability to disassemble RAD51 fibrils was tested for all the BRC repeats by performing in parallel SEC and MP experiments. Interestingly, BRC1-4 disassembled RAD51 fibrils, while BRC6-8 could not affect oligomers’ size. (Figure **3** and Figures **S50, S51**)(19). Following the previous analyses, the effect of BRC1-3 on RAD51 fibril morphology was investigated through Negative Staining Transmission Electron Microscopy (NS-TEM) experiments, as previously done for the BRC4 peptide (Figures **S52-S61**)(19). Without BRC repeats, RAD51 average fibril length was ∼ 50-60 nm. However, upon addition of increasing concentrations of peptides, a progressive reduction in fibril length was observed, with an average minimum length of 30-36 nm at the highest peptide/protein stoichiometric ratio of 2/1. Incubation with increasing concentrations of BRC1-3 thus resulted in a statistically significant reduction in RAD51 WT fibrils’ length (Figure **3**). Finally, for BRC1, BRC2, and BRC3, which induced RAD51 defibrillation, MST analyses were performed, enabling the determination of K_d_ values for the defibrillation processes(19) (∼ 0.5-2 µM) (Figure **3**). Biophysical results obtained using a fully human monomeric RAD51 are consistent with previous studies employing a humanized RadA surrogate(69,70), suggesting that the first four repeats differ in behavior from the last four and likely serve distinct functional roles(26,27). The first four repeats, which display high affinity for RAD51, can depolymerize RAD51 fibrils, suggesting their involvement in RAD51 recruitment in the cytosol.

**Figure 3.**
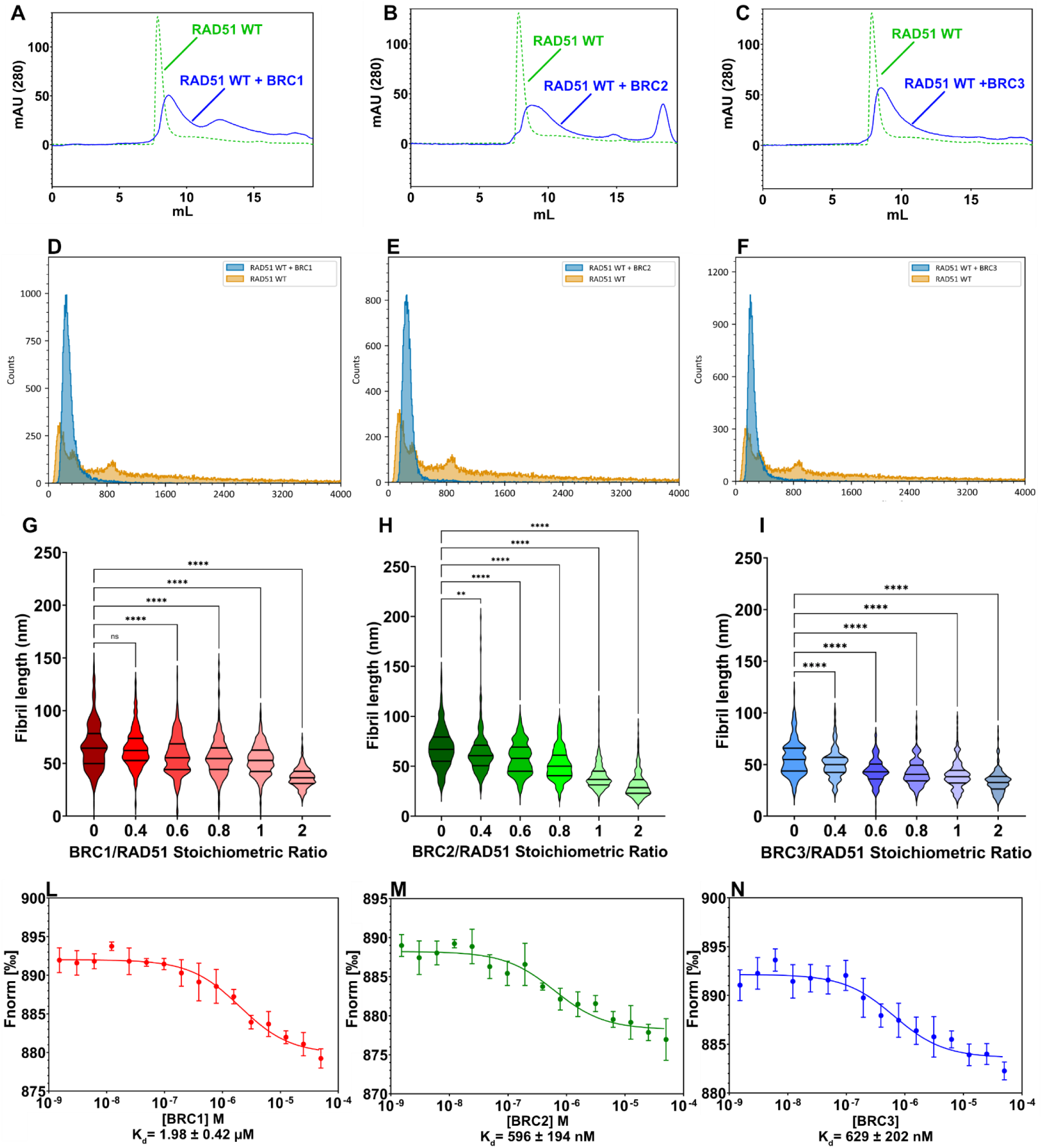
Evaluation of the BRC repeats defibrillation effect on RAD51. Comparison of SEC elution profiles of RAD51 WT in the absence (dashed green line) or in the presence of **A.** BRC1, **B.** BRC2, **C.** BRC3 in 1:4 RAD51/peptide stoichiometric ratio (continuous blue line). Comparison of MP mass distributions of RAD51 WT in the absence (yellow) or in the presence **D.** BRC1, **E.** BRC2, **F.** BRC3 in 1:4 RAD51/peptide stoichiometric ratio (blue). Ordinary one-way ANOVA statistical analysis of RAD51 WT fibrils length (n=300) in NS-TEM experiments in the presence of increasing stoichiometric ratios of **G.** BRC1, **H.** BRC2, **I.** BRC3 MST measurements in the presence of increasing concentration of **L.** BRC1, **M.** BRC2, **N.** BRC3.

In contrast, computational and experimental data indicate that BRC5-BRC8 peptides exhibit weaker binding to RAD51 and are incompetent in disassembling RAD51 fibrils. Notably, NMR data suggest that differences in binding affinities among the peptides likely arise from variations in the aminoacidic composition of their C-terminal regions, which may also influence their ability to promote RAD51 fibril disassembly, consistent with the results from our computational analyses.

### Co-expression, co-purification, and characterization of the complexes of RAD51 with single BRC repeats or BRCA2 truncation harboring multiple repeats

Given that the first four BRC repeats exhibit the strongest affinity for RAD51, they were co-expressed and co-purified with RAD51 to obtain stable complexes for further structural analysis. The following complexes were investigated: Δ97-RAD51/BRC4, RAD51/BRC4, RAD51/BRC3, RAD51/BRC2, and RAD51/BRC1. RAD51 full-length (hereafter referred to as RAD51) was engineered to carry a double mutation in its oligomerization linker to prevent competition between the RAD51 N-terminus and the FXXA motif of the BRC-repeats for the same binding site(7). In the final storage buffer, Na_2_SO_4_(7) was replaced with a mix of NaCl and Li_2_SO_4_, as recent studies highlighted this condition could significantly improve RAD51 stability(71). This protocol enabled the successful purification of all complexes (Figures **4****, S62**), except for RAD51/BRC3, most likely due to the BRC3’s possible degradation or its toxicity in bacteria, a previously reported phenomenon for other peptides(72). Subsequently, also BRCA2 truncations encompassing multiple BRC repeats and their connecting regions (BRC1-2, BRC2-3, BRC3-4, BRC2-4, and BRC1-4) were co-expressed with RAD51, and all corresponding soluble complexes were successfully purified (Figures **4****, S62**). The purified complexes were then further characterized using orthogonal biophysical techniques, enabling accurate estimation of their molecular weights (Figure **4** and Figures **S63, S64**). Samples were essentially homogeneous (Figure **4** and Figures **S63, S64**), and SLS data highlighted an increment of the molecular weights proportional to the number of BRC repeats included in the expressed BRCA2 truncations (Figure **4**). This observation suggests that each BRC repeat binds a single RAD51 monomer with affinities consistent with those previously obtained through biophysical methods (K_d_ ∼ 50-100 nM, Table **1**). To support this hypothesis, MP analyses were performed on RAD51-BRCA2 truncations across various concentrations. Complex disassembly was detected between 50 and 75 nM, a range consistent with the K_d_ values determined by biophysical assays for the first four BRC-repeats (Table **1**, Figures **S65**-**S69**). Given that the K_d_ defines the concentration at which equilibrium between bound and unbound sites is achieved, the MP data provided independent corroboration of the biophysical measurements.

**Figure 4.**
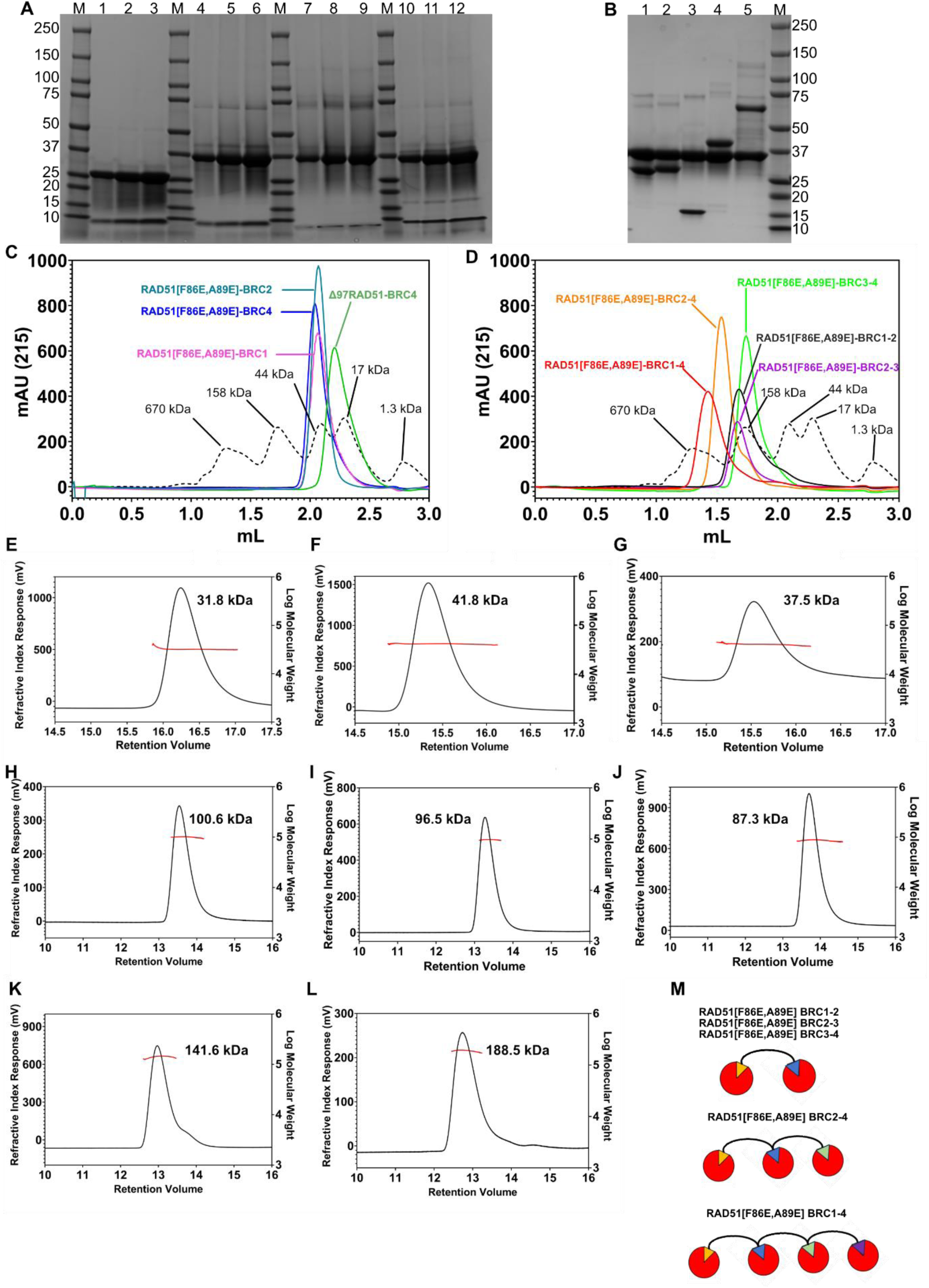
Characterization of the BRC repeats – RAD51 complexes A. SDS Page gel Coomassie Blue Staining. M = Marker, 1 = Δ97-RAD51/BRC4 (10 µg), 2 = Δ97-RAD51/BRC4 (20 µg), 3 = Δ97-RAD51/BRC4 (30 µg), 4 = RAD51/BRC4 (10 µg), 5 = RAD51/BRC4 (20 µg), 6 = RAD51/BRC4 (30 µg), 7 = RAD51/BRC2 (10 µg), 8 = RAD51 /BRC2 (20 µg), 9 = RAD51/BRC2 (30 µg), 10 = RAD51/BRC1 (10 µg), 11 = RAD51/BRC1 (20 µg), 12 = RAD51 /BRC1 (30 µg) **B.** SDS Page gel Coomassie Blue Staining M = Marker 1 = RAD51/BRC1-2 (20 µg) 2 = RAD51/BRC2-3 (20 µg) 3 = RAD51/BRC3-4 (20 µg) 4 = RAD51/BRC2-4 (20 µg) 5 = RAD51/BRC1-4 (14 µg) **C.** Analytical SEC elution profiles of Δ97-RAD51/BRC4 (green), RAD51/BRC4 (blue), RAD51/BRC2 (light blue), RAD51/BRC1 (violet). The profile of globular molecular weight markers is shown as a dashed black line. **D.** Analytical SEC elution profiles of RAD51/BRC1-2 (continuous black line), RAD51/BRC2-3 (violet) RAD51 /BRC3-4 (green) RAD51/BRC2-4 (orange) RAD51/BRC1-4 (red). The elution profile of globular molecular weight markers is shown as a dashed black line. SLS analysis of **E.** Δ97-RAD51/BRC4 **F.** RAD51/BRC4 **G.** RAD51/BRC2 **H.** RAD51/BRC1-2, **I.** RAD51/BRC2-3, **J.** RAD51/BRC3-4, **K.** RAD51/BRC2-4, **L.** RAD51/BRC1-4 **M.** Pictorial representation of the proposed stoichiometry of the reported RAD51/BRCA2 truncates complexes. RAD51 is represented as a sphere while BRC1, BRC2, BRC3 and BRC4 are shown as triangles of different colors.

A protocol for the co-expression and co-purification of RAD51 in complex with BRC2 and with BRCA2 truncations containing multiple repeats was developed. This approach enabled the preparation of complex samples suitable for downstream structural studies. Biophysical characterization of the purified complexes suggests that each BRC repeat interacts with a single RAD51 monomer, with affinities consistent with previously reported values, thereby supporting initial biophysical observations.

### XL-MS on the RAD51-BRC repeats complexes, AlphaFold models generation and validation

To gain additional and more detailed structural insights into the RAD51/BRC-repeats interaction, XL-MS experiments were performed, following a similar approach to previously reported analyses of the interaction between RAD51 and BRC4(25). Initially, Biolayer Interferometry (BLI) measurements were performed to assess whether peptide biotinylation (BRC1-BRC3, BRC7-8) required to detect the efficiency of cross-linking experiments, affected their binding to monomeric RAD51 (Figure **S70**)(25). Once confirmed that this modification did not alter the interaction (Figure **S70**), RAD51 complexes with biotinylated peptides were reconstituted and cross-linked using bis(sulfosuccinimidyl) suberate (BS3) (Figure **S71**)(25).

In addition, we also cross-linked all the other co-expressed and co-purified complexes, namely Δ97-RAD51/BRC4, RAD51/BRC4, RAD51/BRC2, RAD51/BRC1-2, RAD51/BRC2-3, RAD51/BRC3-4, RAD51/BRC2-4, and RAD51/BRC1-4 and we assessed cross-linking efficiency through Coomassie blue stained SDS-Page gels and/or western blots (Figures **S71, S72**). LC-MS/MS analyses led to the identification of high-quality cross-links between BRC1, BRC2, BRC3, BRC4, and the N-terminal domain of RAD51 (Figure **5**, Table **2**). For the BRC4 peptide, we observed that K1536 was cross-linked with RAD51 K58 and K73, while K1541 was cross-linked with RAD51 K40, K58, K64, and K73, located in a cluster of alpha-helices, which supports RAD51 interaction with the DNA(73) (Figure **5**, Table **2**). Similar interactions could also be identified between RAD51 and the first three repeats. Indeed, K58 was also observed to be cross-linked with BRC1 K1015/1025/K1028, BRC2 K1226/K1236 and BRC3 K1440 (Figure **5**, Table **2**).

**Figure 5.**
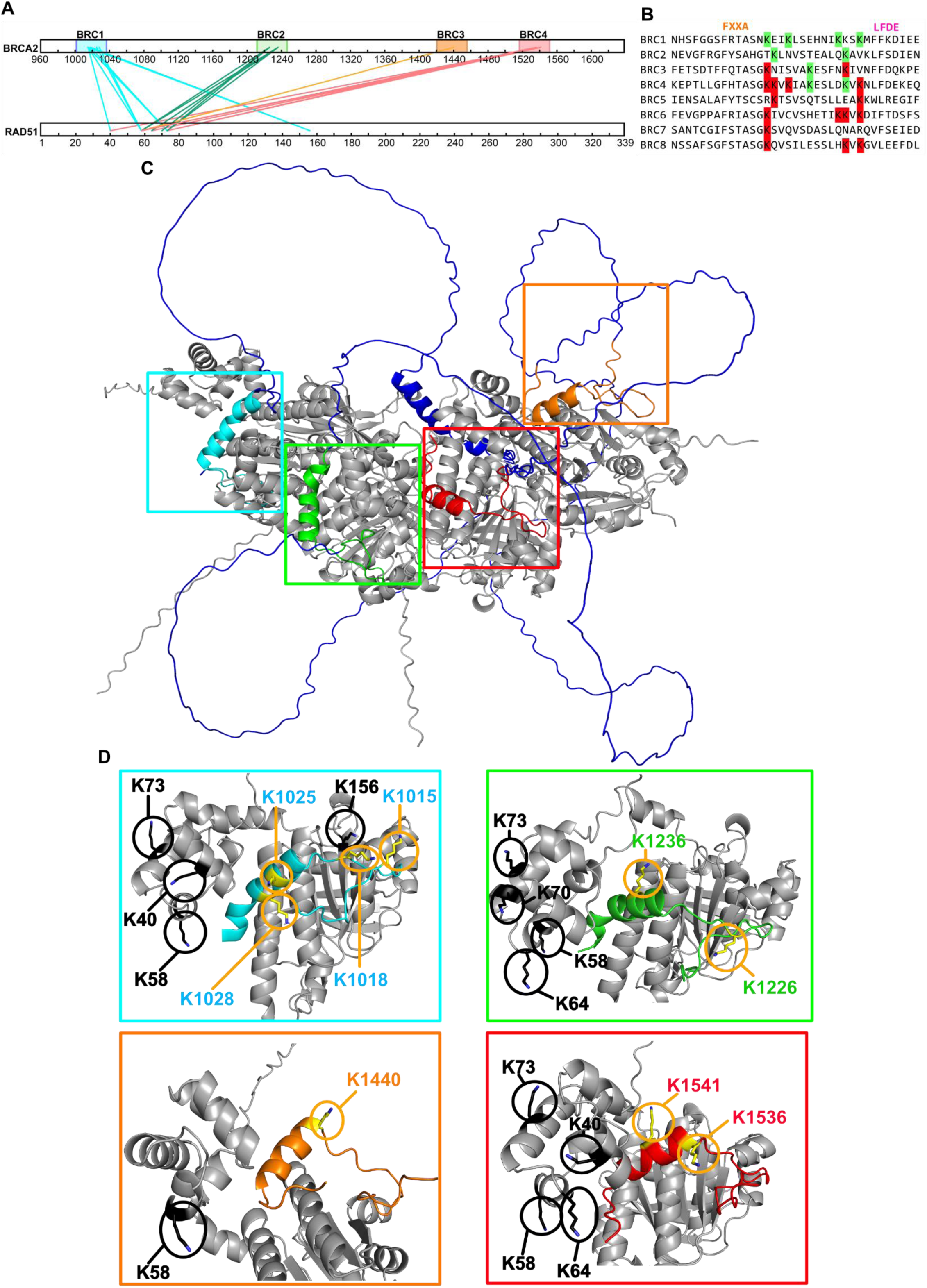
BS3 cross-linking mass spectrometric analysis of the RAD51 interaction with the first four BRC repeats. **A.** Representative map of the crosslinks observed between BRC1, BRC2, BRC3, and BRC4 and RAD51, are displayed in different colors (cyan, green, orange, red respectively). **B.** BRC-repeats multiple sequence alignment, highlighting in green the K residues identified through XL-MS and in red K residues that have a similar position in other peptides, which, however, were not identified in our XL-MS experiments. We speculate that the different folding of the peptides can prevent the interaction of K residues with the RAD51 N-terminus. **C.** AF prediction of the RAD51/BRC1-4 complex. Each peptide has been colored differently: BRC1, cyan; BRC2, green; BRC3, orange, BRC4 red. The BRCA2 regions connecting BRC repeats have been colored in blue, while the RAD51 monomers are depicted in grey. **D.** Close-up view of the single BRC-repeats highlighting the identified cross-linked residues in RAD51 (black) and in BRC1 (cyan), BRC2 (green), BRC3 (orange), and BRC4 (red).

**Table 2.** Detected inter-protein crosslinks. Both the BS3 cross-linked amino acids in BRCA2 and RAD51 are reported as well as their corresponding position in BRC repeats.

| Position BRCA2 | Position RAD51 | Repeat | Identified in samples cross-linked with BS<sub>3</sub> |
| --- | --- | --- | --- |
| 1015 | 40 | **BRC1** | RAD51/BRC1 |
| 1015 | 58 |  | RAD51/BRC1 |
| 1015 | 73 |  | RAD51/BRC1 |
| 1015 | 156 |  | RAD51/BRC1, RAD51/BRC1-2, His-RAD51-Mon/bioBRC1 |
| 1018 | 73 |  | RAD51/BRC1 |
| 1018 | 156 |  | RAD51/BRC1 |
| 1025 | 58 |  | RAD51/BRC1 |
| 1028 | 58 |  | RAD51/BRC1 |
| 1226 | 58 | **BRC2** | RAD51/BRC2 |
| 1226 | 70 |  | RAD51/BRC2 |
| 1226 | 73 |  | RAD51/BRC2 |
| 1236 | 58 |  | RAD51/BRC2, RAD51/BRC1-2, RAD51/BRC2-3, RAD51/BRC2-4 |
| 1236 | 64 |  | RAD51/BRC2 |
| 1236 | 73 |  | RAD51/BRC2, RAD51/BRC2-3, RAD51/BRC2-4 |
| 1440 | 58 | **BRC3** | RAD51/BRC3-4, RAD51/BRC1-4 |
| 1536 | 58 | **BRC4** | RAD51/BRC4 |
| 1536 | 73 |  | RAD51/BRC4 |
| 1541 | 40 |  | RAD51/BRC4, RAD51/BRC3-4 |
| 1541 | 58 |  | RAD51/BRC4, RAD51/BRC3-4, RAD51/BRC2-4, RAD51/BRC1-4 |
| 1541 | 64 |  | RAD51/BRC4, RAD51/BRC3-4 |
| 1541 | 73 |  | RAD51/BRC2-4 |

Moreover, K40 and K73 in the same RAD51 region were also observed to be cross-linked with BRC1 K1015/K1018 and BRC2 K1236/K1226, respectively (Figure **5**, Table **2**). Notably, all the crosslinked residues identified within the first four peptides were localized in the region between the FXXA and the LFDE domains, highlighting the importance of the BRC-repeats in facilitating interaction with the RAD51 N-terminal domain (Figure **5**). At this stage, the consistency of the AF predictions was verified with the obtained XL-MS data. We first considered the RAD51-BRC4 complex by generating AF2 and AF3 models and comparing their outputs (Figures **S73**-**S75**). The five best-ranked AF2 models were quite similar and compact, while the AF3 predictions displayed a higher conformational heterogeneity as the RAD51 N-terminal domain was modeled either more compact or extended (Figures **S73**, **S74**). Consistent with our previous observations(25), the generated AF2 models displayed Cα-Cα distances compatible with the experimental results and within the typical distance constraints for BS3 described in the literature(74–76) (**Table S1**). In contrast, we noticed that the Cα-Cα distance of the identified cross-linked residue pairs varied significantly depending on the conformation of the RAD51 N-terminal domain modeled by AF3, except for K1541-K58 and K1541-K40, which consistently remained within the expected BS_3_ cross-linking distance constraints across all AF simulations (**Table S2**). Similar trends were observed for RAD51-BRC1, RAD51-BRC2, RAD51-BRC3, and their corresponding AF3 predictions (Figures **S76-S81,** Tables **S3-S5**). These results may be explained by the intrinsic flexibility of the RAD51 N-terminal domain in solution, as indicated by the high positional error in the PAE matrix (Figures **S76, S78, S80**).

We then focused on the larger complexes comprising BRCA2 truncations harboring multiple repeats in complex with RAD51. Interestingly, we observed that AF2/AF3 outputs displayed similar features to those observed in the RAD51/BRC3-4 complex (Figures **S82, S83**). Indeed, BRC3 and BRC4 were modeled as binding to RAD51 as the canonical BRC4 binding site. However, both AF2/AF3 predictions were quite compact and characterized by the presence of intrinsically disordered loops connecting the BRC repeats (Figures **S82-S83**). For these reasons, we simplified XL-MS analyses considering only one prediction for each cross-linked BRCA2 truncations/RAD51 complex, selecting models that did not display any significant steric clash (Figures **S84-S88**). In these models, most of the identified cross-link pairs between the peptides and RAD51 N-terminus matched permissive distances in the predicted structures, despite some pairs that systematically displayed Cα-Cα distances > 30 Å (**Tables S6-S10**). These data allowed us to validate the BRC-repeats/RAD51 interaction, providing valuable constraints for subsequent structural studies.

### SAXS and NMR studies on the co-expressed and co-purified RAD51 – BRC repeats complexes

To further characterize the interaction between BRC-repeats and RAD51, we performed a systematic study using SEC and batch SAXS on all purified complexes (Figures **S89-S104**, Tables **S11-S19**). Primary data analysis of the collected SEC-SAXS data revealed significant differences in the structural parameters of the analyzed samples (Figures **S89-S96,** Table **3**). Guinier analysis showed that the radii of gyration (R_g_) varied considerably depending on the analyzed complex (Figures **S89-S96,** Table **3**). Indeed, single BRC repeats/RAD51 complexes showed smaller R_g_ values, whereas BRCA2 truncations encompassing multiple peptides bound to RAD51 monomers exhibited increased R_g_ values, directly correlating with the number of peptides and, consequently, the number of coordinated RAD51 subunits (Figures **S89-S96,** Table **3**). These results were further supported by the calculation of the p(r) function, which revealed similar trends in Dmax and demonstrated good agreement between p(r) and Guinier R_g_ values (Figures **S89-S96,** Table **3**). Interestingly, dimensionless Kratky plots showed varying degrees of structural compactness across the analyzed complexes, consistent with features observed in the previously calculated p(r) functions (Figures **6A** and **S89-S96**). Specifically, the Δ97-RAD51/BRC4 complex displayed a compact conformation, whereas all the other complexes showed characteristics of flexible particles, with the degree of flexibility correlating with the size of the BRCA2 truncations (Figures **6A** and **S89-S96)**. In addition, good agreement was observed between the molecular weights of the complexes estimated from correlation volume (V_c_), SLS/MP data, and theoretical calculations, further supporting the purported 1:1 stoichiometry of BRC-repeats to RAD51 in the BRCA2 truncations-RAD51 complexes (Table **3**, Tables **S11-S19**).

**Figure 6.**
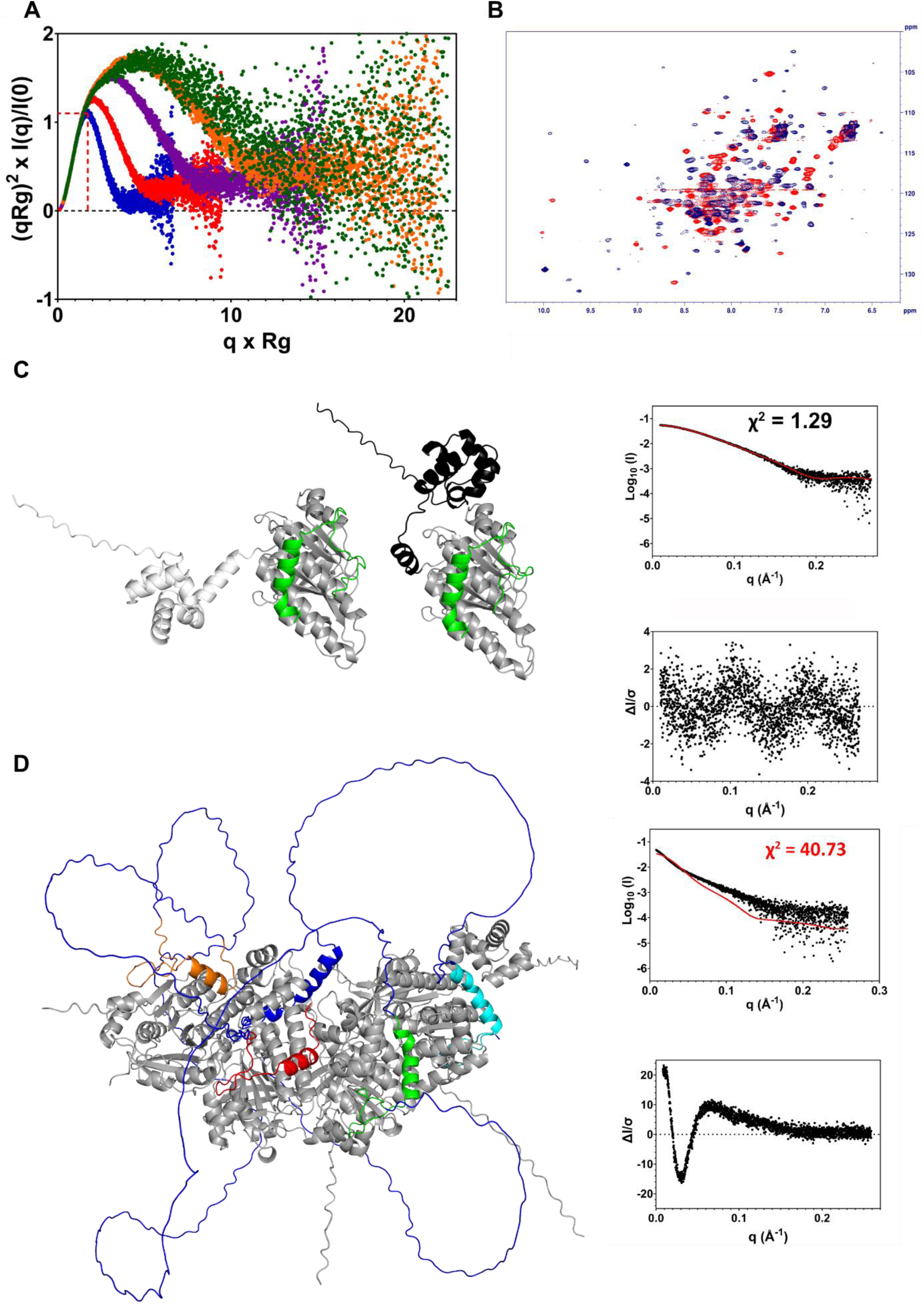
SAXS and NMR studies on BRC repeats – RAD51 complexes A. Dimensionless. Kratky Plot overlay of Δ97-RAD51/BRC4 (Blue), RAD51/BRC4 (Red), RAD51/BRC3-4 (Violet), RAD51/BRC2-4 (Orange), RAD51/BRC1-4 (Dark Green) **B.** Overlay of the ^1^H-^15^N Heteronuclear Single Quantum Coherence (HSQC) spectra of Δ97-RAD51/BRC4 (blue) and RAD51/BRC4 (red) **C.** Multi-FoXS modeling of RAD51/BRC2 utilizing the five AlphaFold3 models as initial ensemble. Left: the two best scoring Multi-FoXS 2-state models for the AF RAD51/BRC2 complex. Right: model fit to experimental SAXS data and the relative residuals. RAD51 C-terminus is colored grey, while the two different conformation of the RAD51 N-terminal domain are highlighted in white and black. The BRC2 peptide is shown in green. **D.** FoXS modelling of BRC1-4/RAD51 complex utilizing the AlphaFold prediction as an input model. RAD51 monomers are colored grey. BRC1 is cyan, BRC2 is green, BRC3 is orange, BRC4 is red and the BRC-repeats connecting loops are colored blue. FoXS modelling for the input structure (red curve) is compared with the experimental SAXS data (black dots), showing the relative residuals.

**Table 3.** SEC-SAXS primary data analysis of different BRC repeats-RAD51 complexes.

| Sample | R<sub>g</sub> - Guinier | R<sub>g</sub> – p(r) | D<sub>max</sub> | V<sub>c</sub> MW | SLS | Expected-MW |
| --- | --- | --- | --- | --- | --- | --- |
| Δ97-RAD51/BRC4 | 20.30 ± 0.03 Å | 20.18 ± 0.02 Å | 66.0 Å | 29.6 kDa | 31.8 kDa | 30.9 kDa |
| RAD51/BRC4 | 27.82 ± 0.03 Å | 28.37 ± 0.04 Å | 105.0 Å | 40.3 kDa | 41.8 kDa | 41.5 kDa |
| RAD51 /BRC2 | 27.29 ± 0.07 Å | 27.51 ± 0.04 Å | 96.0 Å | 37.9 kDa | 37.5 kDa | 41.4 kDa |
| RAD51/BRC1-2 | 52.54 ± 0.20 Å | 55.30 ± 0.20 Å | 210.0 Å | 98.9 kDa | 100.6 kDa | 101.7 kDa |
| RAD51/BRC2-3 | 54.39 ± 0.19 Å | 58.82 ± 0.22 Å | 230.0 Å | 93.9 kDa | 96.5 kDa | 101.8 kDa |
| RAD51/BRC3-4 | 45.61 ± 0.10 Å | 47.51 ± 0.09 Å | 180.0 Å | 85.5 kDa | 87.3 kDa | 89.7 kDa |
| RAD51/BRC2-4 | 66.37 ± 0.30 Å | 69.87 ± 0.26 Å | 280.0 Å | 152.1 kDa | 141.6 kDa | 149.8 kDa |
| RAD51/BRC1-4 | 77.09 ± 0.47 Å | 84.26 ± 0.78 Å | 340.0 Å | 220.9 kDa | 188.5 kDa | 210.2 kDa |

In parallel to SEC-SAXS experiments, batch SAXS measurements (Figures **S97**-**S104,** Tables **S11-S19**) were performed, providing structural parameters in line with SEC-SAXS results, further supporting the robustness of the data presented in this work (Figures **S97**-**S104,** Tables **S11-S19**). The comparison of the dimensionless Kratky plot data for Δ97-RAD51/BRC4 and RAD51/BRC4 complexes suggested that while the former complex behaved as a compact and mostly folded particle, the latter was more flexible (Figure **6A**). To further support this analysis, both these complexes were produced using a ^15^N isotopic enrichment protocol to record ^1^H - ^15^N Heteronuclear Single Quantum Coherence (HSQC) spectroscopy experiments(51)(Figures **6B** and **S105, S106**). The collected ^1^H -^15^N NMR spectra showed peaks coalescence in the 7-8 ppm region for the RAD51/BRC4 complex, indicating underlying conformational dynamics (Figures **6B** and **S106**). This behavior was only partially observed in the spectrum of the N-terminal domain truncated counterpart (Figures **6B** and **S106**). These observations were consistent with SAXS data, further corroborating the hypothesis that RAD51 N-terminal domain behaves as a flexible region upon binding of the BRC4 peptide (Figures **6B** and **S106**)(7,25). SAXS data were subsequently used for atomistic modeling. As Δ97-RAD51/BRC4 AF2/AF3 prediction substantially overlaid (R.M.S.D = 0.166), the latter was used for fitting SAXS data, yielding a final χ^2^ value of 1.15 with FoXS (Figures **S107, S108**). For the RAD51-BRC4 complex, a significant difference was observed between the predictions generated by AF2 and AF3. While AF2 produced compact and structurally consistent models, AF3 generated a more heterogeneous ensemble in which the RAD51 N-terminus was predicted to adopt multiple distinct conformations (Figures **S109**, **S110**). Interestingly, despite the AF2 models satisfying all the identified cross-link pairs exhibiting Cα-Cα distances ≤ 30 Å (Table **S1**), they poorly fitted experimental data with a χ^2^ ∼ 35, suggesting that they were not representative of the experimental data (Figure **S109**). By contrast, the AF3 models exhibited significant variability in their fitting (Figure **S110**). Indeed, while the most extended structures provided the best fitting (χ^2^ ∼ 1.9), the most compact ones led to the worst scores (χ^2^ ∼ 38, Figure **S110**). Notably, the best-fitting predictions showed poor compliance with the XL-MS data, whereas the structures poorly fitting to the SAXS profile exhibited Cα-Cα distances within the limits defined by BS_3_ cross-linkers (Table **S2**, Figure **S110**). Strikingly, by employing five AF3 models for multi-state modeling using Multi-FoXS, a two-state model was obtained that nicely fitted the experimental data, yielding a χ^2^ value of approximately 1.1 (Figure **S111**). Application of the same methodology to the AF3-generated RAD51/BRC2 models yielded comparable results in both FoXS and Multi-FoXS modeling, suggesting similar behavior of the complex in solution (Figures **6C** and **S112**). These results further support our hypothesis that the RAD51 N-terminal domain undergoes structural rearrangements upon BRC-repeats binding(7,25), and demonstrate that AF3 can predict conformational ensembles that closely align with the experimental SAXS data. We then turned our attention to modeling of the BRCA2 truncations – RAD51 complexes. FoXS modeling of the AF-generated structures resulted in high χ^2^ for all the analyzed protein complexes in solution, suggesting that the input models were not representative of the experimental data (Figures **6D** and **S113-116**). Indeed, we did not account for the flexibility of our systems, which was observed in the dimensionless Kratky plots. This behavior could possibly derive from the conformational flexibility of the RAD51 N-terminus and from the BRCA2 regions, bridging together different peptides. Indeed, AF predicted these regions as intrinsically disordered loops. Moreover, we also observed, in all the predicted models, a significant positional error in the PAE matrix, suggesting that these regions may be quite dynamic in solution (Figures **S84-S88**). Recent NMR studies by Julien et al. showed that the BRC1-BRC2 connecting region is essentially disordered, possibly suggesting similar behaviors also for the other connecting regions(77). Therefore, to rationalize the behavior of these regions in solution and to guide our SAXS modeling efforts, we employed CALVADOS(78,79). Specifically, CALVADOS enables the study of IDR conformational ensemble in the absence of experimental data, providing a scaling exponent *v* that describes the structural compactness of the studied chains(78,79). Interestingly, the calculated *v* for BRC1-2, BRC2-3, and BRC3-4 connecting regions were similar, suggesting that these polypeptide chains have comparable behavior in solution (Figure **S117**). We deemed it unlikely that these complexes could adopt such compact structures as in the AF predictions and concluded that they would most likely behave as an ensemble of conformations. Therefore, we performed multi-state modeling through MultiFoXS on all these complexes by specifically assigning flexibility to the interconnecting BRCA2 regions. To improve the computational efficiency of our modeling, we did not account for the flexibility of the RAD51 N-termini, and we utilized XL-MS data as a constraint for the BRC repeats/RAD51 interfaces. While we recognized that this approach would significantly simplify the actual behavior of these complexes in solution, it also enables MultiFoXS to explore a broader range of possible conformations for the interconnecting regions (Tables **S11-S19**). Given the ambiguity of the collected datasets, as indicated by the Ambimeter(58) algorithm (Tables **S11-S19**), we performed two replicates of multi-state modeling for each complex (Tables **S11-S19**, Figures **7** and **S118**-**S145**). Interestingly, four and five-state models provided the best fits, leading to a χ^2^ value of approximately 1/2, thus remarkably improving the agreement with the experimental data compared to the initial models (Tables **S11-S19**, Figures **S118**-**S145**). For example, a four-state model of the BRC1-4/RAD51 complex, which encompasses all the BRC repeats, provided the best fit to the experimental data (**Figure 7**). Upon closer inspection of the four most representative models, we observed both extended and compact states, suggesting significant conformational rearrangements of the structures in solution (**Figure 7**). Structural SAXS and NMR data on BRC-RAD51 complexes support the hypothesis that binding of BRC-repeats induces a re-arrangement of the RAD51 N-terminal domain, which becomes a flexible region capable of exploring multiple conformations in solution. Moreover, structural modelling of SAXS data of the BRCA2 truncation in complex with RAD51 clearly indicates the presence of intrinsically disordered regions connecting different BRC-repeats.

**Figure 7.**
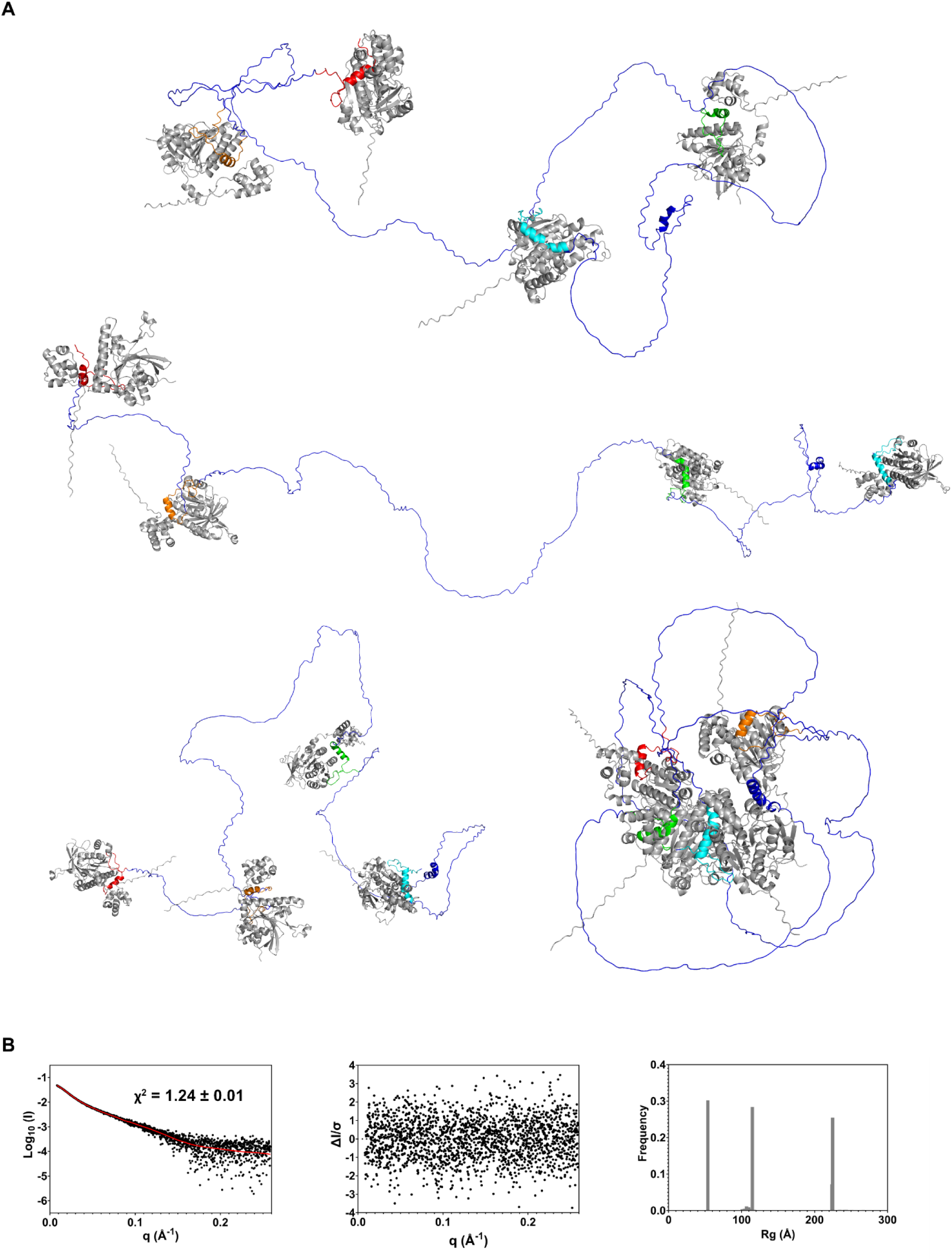
Multi-FoXS modeling of RAD51/BRC1-4. **A.** The Multi-FoXS 4-state models for the BRC1-4/RAD51 complex. RAD51 monomers are colored in grey, BRC1 in cyan, BRC2 in green, BRC3 in orange, and BRC4 in red, and the region connecting the different peptides in blue **B**. Left: model fit to experimental SAXS data. Center: model residuals. Right: distribution of radii of gyration in the 5-state models.

## DISCUSSION

HR requires a series of coordinated steps, involving different sensors and effectors, to ensure the high-fidelity fixing of deleterious DSBs(1,2). Recent findings have highlighted the central role of BRCA2 in orchestrating several key factors (e.g., RAD51 paralogs, PALB2, etc.) to facilitate the recruitment and regulation of RAD51 activity, which lies at the heart of a sophisticated “recombinosome” machinery(31,80). A detailed understanding of the behavior of this multifaceted and multifunctional protein in its primary interaction with RAD51 is crucial, as this protein-protein interaction underpins the coordinated activity of the other components within the HR machinery. Such insight will be fundamental for establishing an integrated and mechanistic view of the entire HR process. To the best of our knowledge, most existing data on the human RAD51–BRCA2 interaction focus on the DNA repair phase, i.e., in the presence of DNA(26,27). However, it is well established that these two proteins first interact in the cytosol, and any disruption at this early stage can severely impair the DNA repair process, potentially leading to irreversible genomic damage(8). It has already been suggested that upon DNA damage, BRC4, the fourth repeat of BRCA2, disassembles RAD51 fibrils gradually from their termini, detaching one RAD51 monomer at a time, rather than acting randomly at different fibril positions(19). Despite this, it is still unclear how, during cytosolic RAD51 nuclear recruitment, each of the other seven BRC repeats interacts with RAD51, how they coordinate their interactions, and ultimately what their role is(8,25). In this work, we address this long-lasting question through a multidisciplinary approach, gathering information through biochemical, biophysical, structural, and computational studies. Orthogonal approaches clearly show that BRC1-4 repeats bind RAD51 with high affinity, compared to BRC5-8 ones. Notably, the peptides exhibiting higher affinities are also those capable of disassembling RAD51 WT fibrils. Importantly, we also correlate peptides’ amino acid composition with interaction effects. Consistent with previous studies focused on BRC4(81), our analyses highlight that the LFDE motif is critical for mediating high-affinity interactions across all BRC repeats, guiding the peptides’ ability to initiate interactions with RAD51 fibrils. Moreover, XL-MS studies suggest that residues located between the FXXA and the LFDE domains are essential for detaching each RAD51 monomer from its neighbor in the fibrils, destabilizing the interaction between each RAD51 N-terminus and the adjacent protomer, and promoting its re-arrangement into a flexible region. This behavior is essential for proper filament disassembly, possibly enabling the sequential engagement of individual BRC-repeats to RAD51 monomers, as also corroborated by further studies carried out on larger BRCA2 truncations in complex with RAD51, where each BRC-repeat coordinates a single RAD51 monomer. Moreover, structural studies also highlight that BRC-repeats, like other peptides(82), are embedded into flexible and disordered regions, thus allowing these complexes to adopt multiple conformations. This structural adaptability is likely essential for maintaining RAD51 monomers in a separate state, thus preventing premature reassembly and enabling their proper nuclear import. Moreover, once in the nucleus, the dynamic behavior of BRCA2 flexible interconnecting regions is also likely designed to prime each RAD51 monomer to a specific position onto the resected ssDNA, which is itself highly dynamic and capable of adopting multiple conformations(83). This would agree with recent findings assessing that BRCA2 modulates the recombinase’s binding selectivity for ssDNA(8). Overall, these findings suggest that the first four BRC-repeats function like flexible fingers, each designed to detach individual monomers from cytosolic RAD51 fibrils and guide them to specific positions along the ssDNA (Figure **8**). Nevertheless, the role of BRC5–BRC8 remains unclear. Our evidence suggests that they are unlikely to significantly contribute to RAD51 recruitment in the cytosol. However, as highlighted by other studies(26,27), the presence of cofactors in the nuclear environment may substantially alter protein–protein affinities, potentially allowing BRC5–BRC8 to play important roles in supporting RAD51 activity during HR, thus preventing premature disassembly. Furthermore, emerging structural and functional data suggest that BRC-repeats may serve as a hub for interactions with other essential components of the HR pathway. Indeed, BRC1 and BRC2 have already been reported to interact with the BCDX2 complex, composed of the RAD51 paralogs RAD51B, RAD51C, RAD51D, and XRCC2, which play a key role in RAD51 filament assembly and stabilization(31). Notably, despite the poor sequence identity of RAD51 and the RAD51 paralogs, they all contain conserved FXXA/FXXS motifs, similar to those found in BRC-repeats and in other HR factors, suggesting a shared mode of interaction(31). Therefore, it is plausible that sequence variations, particularly within the LFDE motifs of the BRC repeats, could be crucial in determining the specificity for different interaction partners. This raises the possibility that the last four repeats, while showing weaker affinity for RAD51, may preferentially engage other HR factors in the nuclear environment.

**Figure 8.**
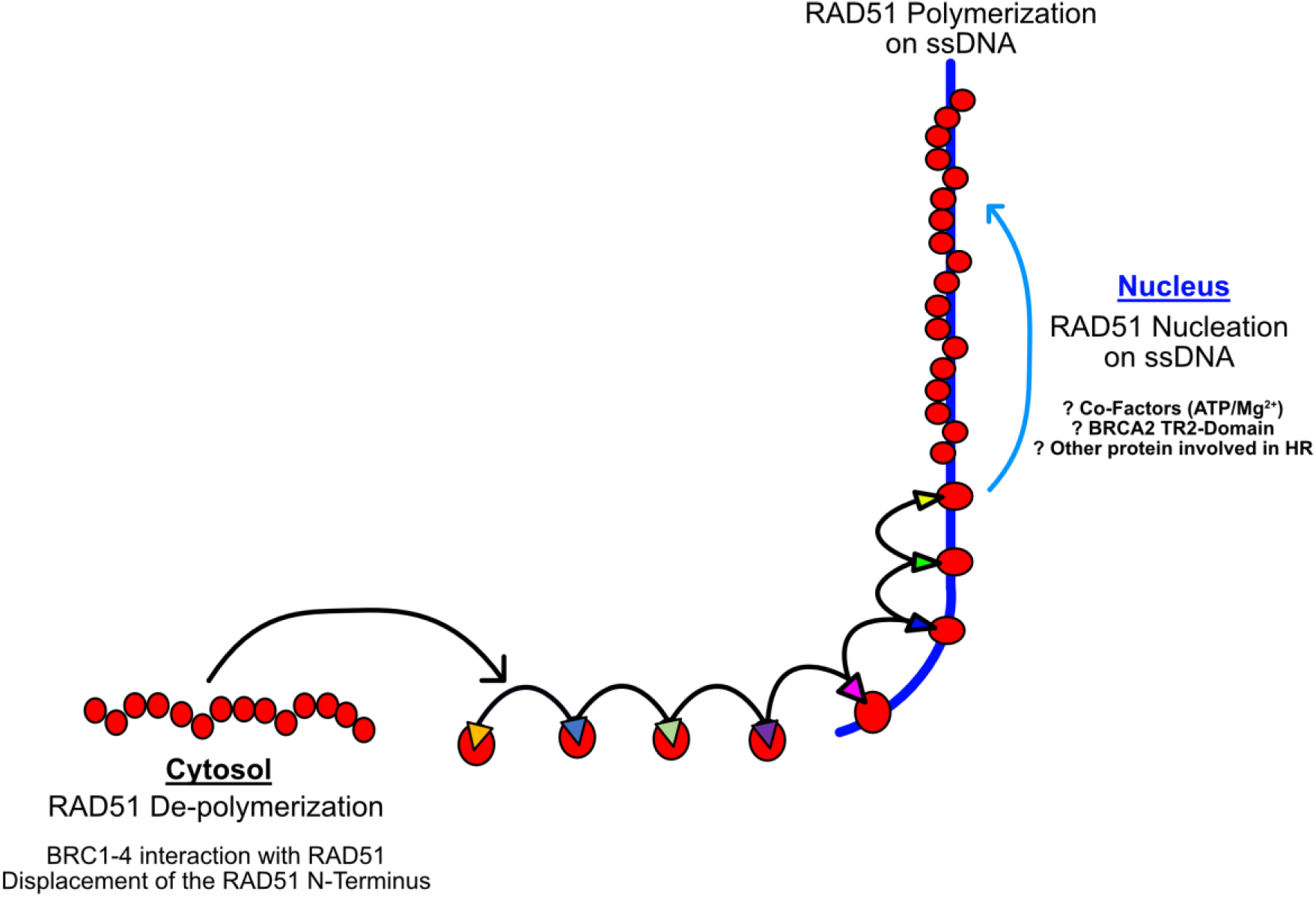
Pictorial view of the BRC repeats role in the HR pathway. The first four BRC repeats present an aminoacidic composition that allows the depolymerization of RAD51 by triggering the displacement of the RAD51 N-terminus. Once RAD51 is recruited, it is translocated into the nucleus where it is delivered onto the resected ssDNA. Here, BRC5-8 could be crucial to allow the optimal priming and spatial organization for the RAD51 loading onto the DNA. Nevertheless, other factors are also involved in this process, and the mechanistic details are still unclear.

Altogether, our results show that BRC repeats are not merely ancestral remnants; rather, each repeat is finely tuned through its amino acid composition to play a specific role within the HR. Moreover, the highly flexible interconnecting regions between the repeats are specifically designed to functionally contribute to the recombination machinery, pinpointing the range of action of the connected repeats and contributing to cytosolic RAD51 recruitment. The complex and finely tuned mechanism involving BRCA2 and its interaction with RAD51, and the indispensability of each of its elements, helps explain why approximately 33% of BRCA2 mutations associated with increased cancer risk^1,2,3^ occur within the BRC repeats. Several of these pathogenic variants have already been associated with tumor onset and progression(84,85). The data presented here further justify why mutations affecting a single BRC repeat cannot be compensated for by the others(8), as, despite their high sequence homology, each repeat performs a distinct and essential role within the RAD51-BRCA2 machinery. However, many questions concerning the structural organization of the BRC-repeats in the nuclear compartment remain. Highly performing folding predictors (AF2/3) only partially help to shed light on this structural complexity. Future studies involving multiprotein complexes that include the presence of additional factors such as DNA, ATP, and multiple key HR factors will be essential to fully understand the complex and tightly regulated mechanism that governs the HR machinery and the specific role of BRC-repeats within this context. These studies will be crucial to unravel the detailed mechanisms underlying this intricate and sophisticated DNA-repair system, but also to provide a characterization of its different layers that could provide critical targets for innovative cancer therapies. Moreover, they could provide mechanistic explanations for the functional failures induced by specific mutations, thereby opening the way toward personalized treatment strategies.

## ACKNOWLEDGEMENTS

The authors gratefully thank the European Institute of Oncology (IEO) Biochemistry and Structural Biology Unit, Federico Donà, Sara Martin, Edoardo Gelardi and Cristina Cecchetti for useful discussions. Moreover, the authors acknowledge the Data Science and Computation Facility and its Support Team at Fondazione Istituto Italiano di Tecnologia for computing time and support on the Franklin HPC system, and Imke Wüllenweber for excellent technical assistance for proteomics sample preparation. The authors would like also to express their gratitude to the High-throughput SAXS B21 at Diamond Light Source (DLS) for providing access to the synchrotron radiation facility for SAXS measurements (rapid access proposal SM37027, awarded to Dr. Stefania Girotto and Dr. Francesco Rinaldi) and the beamline scientists Dr. Nathan Cowieson, Mr. Nikul Khunti and Dr. Katsuaki Inoue for fruitful discussions and continuous support.

## AUTHOR CONTRIBUTIONS

Conceptualization and methodology: F.R., J.L., S.G. and A.C. Investigation: Biophysical Assays, Protein characterization, SAXS data collection and modelling, AlphaFold predictions: F.R.; Protein expression and purification: F.R. and G.V.; Cloning: E.R.; Cross-linking mass spectrometry P.F. and J.L.; NMR studies: M.V.; Alanine scanning, Molecular dynamics simulations and ligand interaction diagrams V. B. and M.B.; Transmission Electron Microscopy: F.C. Formal analysis: F.R., F.C. M.B., V.B., M.V, P.F., J.L. Writing - Original Draft: F.R., S.G., M.B., E.R., P.F., J.L., M.V. Writing- Review & Editing: All authors. Project administration: F.R., S.G. and A.C. Supervision: F.R., S.G. and A.C. Funding acquisition: A.C.

## SUPPLEMENTARY DATA

Supplementary Data are available at NAR online.

## DATA AVAILABILITY

The data and models that support the findings of this study are openly available in SASBDB at https://www.sasbdb.org/(64,65) with the following reference numbers: SASDWT5, SASDWU5, SASDWV5, SASDWW5, SASDWX5, SASDWY5, SASDWZ5, SASDW26, SASDW36, SASDW46, SASDW56, SASDW66, SASDW76, SASDW86, SASDW96, SASDWA6, SASDWB6, SASDWC6, SASDWD6, SASDWE6, SASDWF6, SASDWG6, SASDWH6, SASDWJ6, SASDWK6, SASDWL6, SASDWM6, SASDWN6, SASDWP6, SASDWQ6, SASDWR6, SASDX95, SASDXA5, SASDXB5, SASDXC5.

The mass spectrometry proteomics data have been deposited to the ProteomeXchange Consortium via the PRIDE partner repository with the identifier PXD065998(86). We freely provide AlphaFold models not deposited in SASBDB, as well as Jupyter notebook to reproduce our analyses, results, and the plots reported in this work in Zenodo with accession code 10.5281/zenodo.15553042.

