## Supplementary Information for "Breaking new ground into RAD51–BRC repeats interplay in Homologous Recombination"

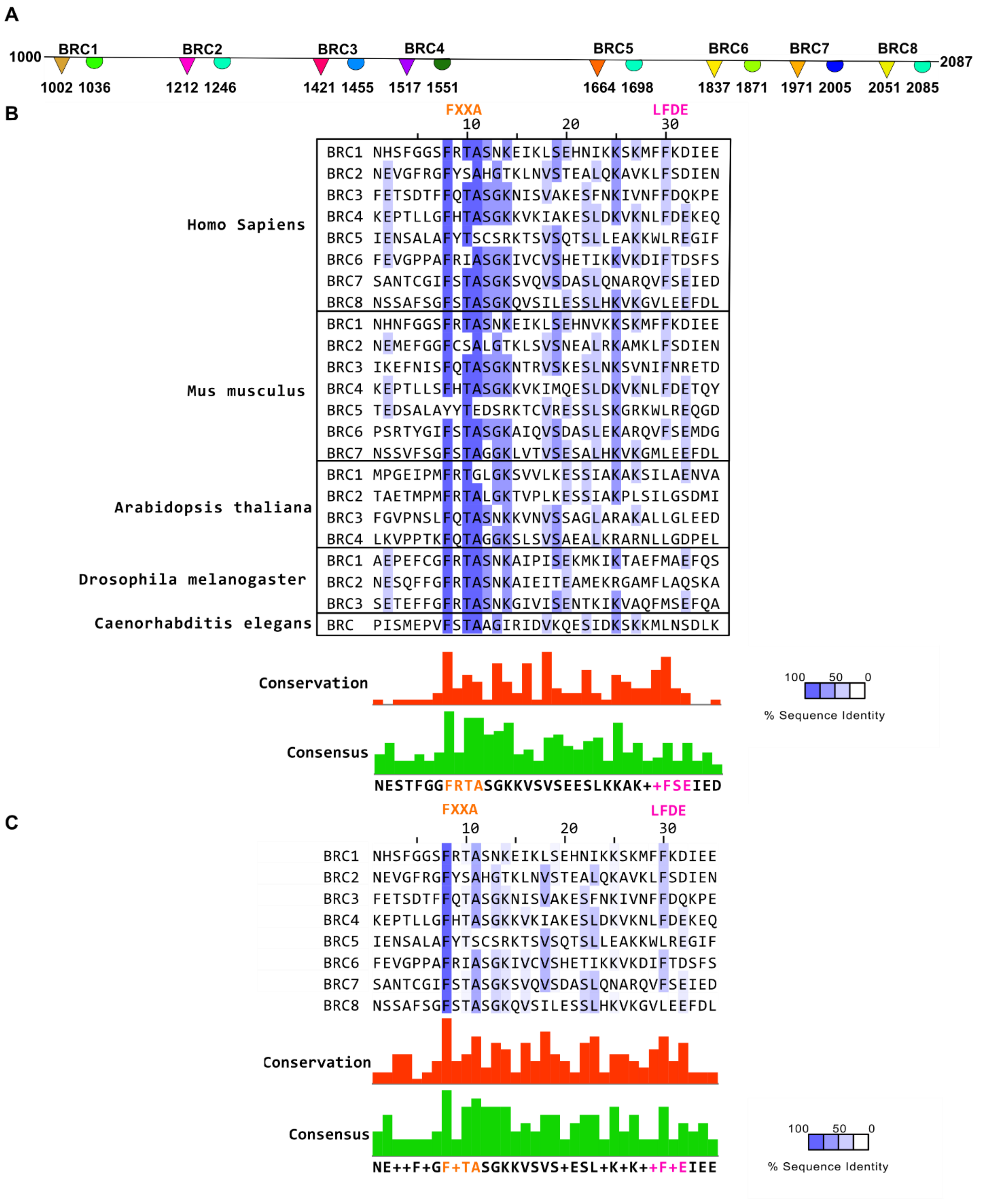


**Supplementary Figure 1 BRC repeats aminoacidic conservation. A.** Schematic representation of human BRC repeats region. Triangles and spheres represents the FXXA and LFDE domains of each BRC repeats. **B.** Multiple sequence alignment of the BRC repeats of different species **C.** Sequence alignment of the human BRC repeats.

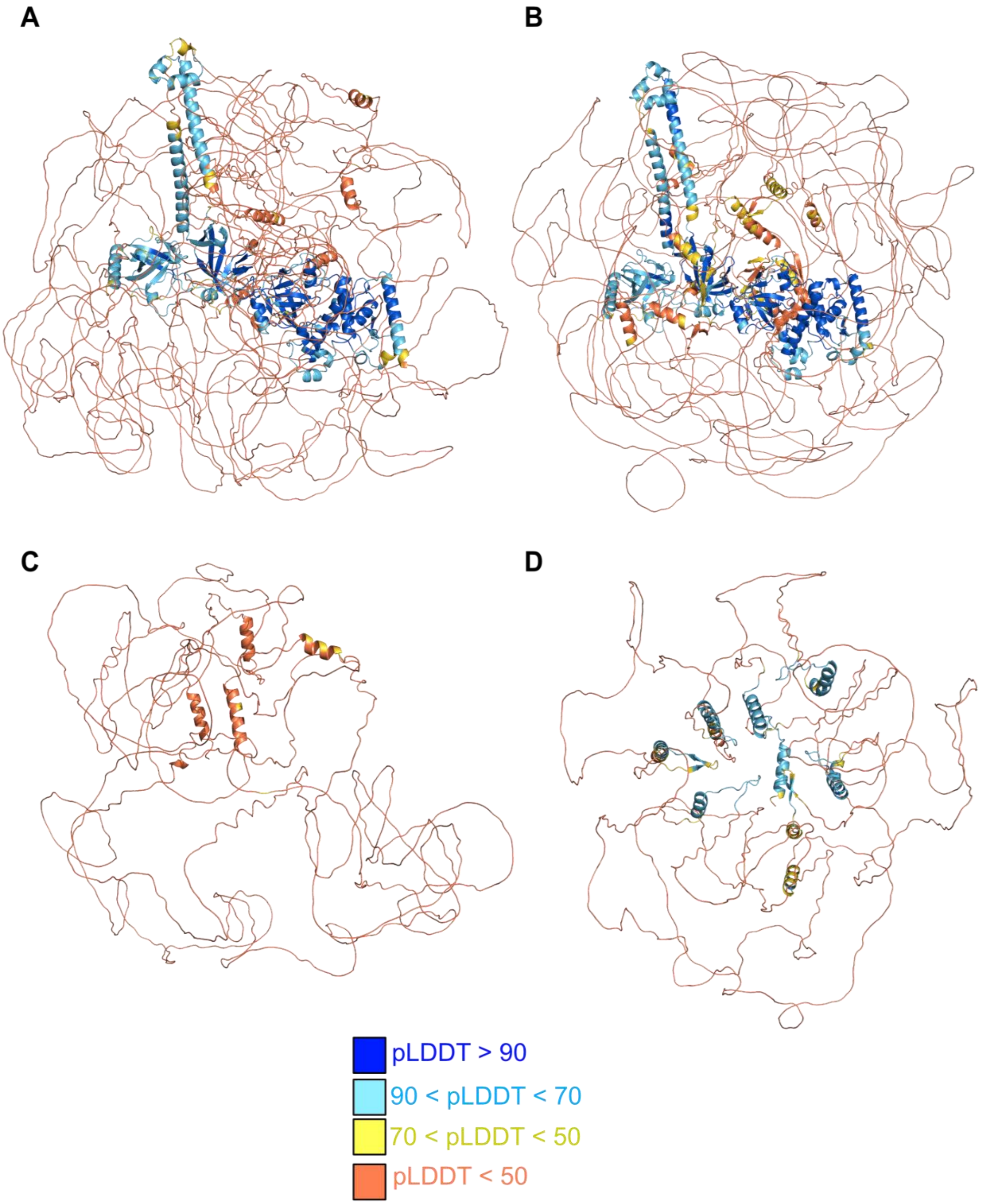


**Supplementary Figure 2 AlphaFold models of BRCA2 and the BRC repeats. A.** AlphaFold2.3 and **B.** AlphaFold3 predictions of BRCA2-FL **C.** BRC-repeats region extracted from the prediction of FL-BRCA2 **D.** AlphaFold prediction of the isolated BRC-repeats region. Residues have been colored based on predicted Local Distance Difference Test (pLDDT).

**A**

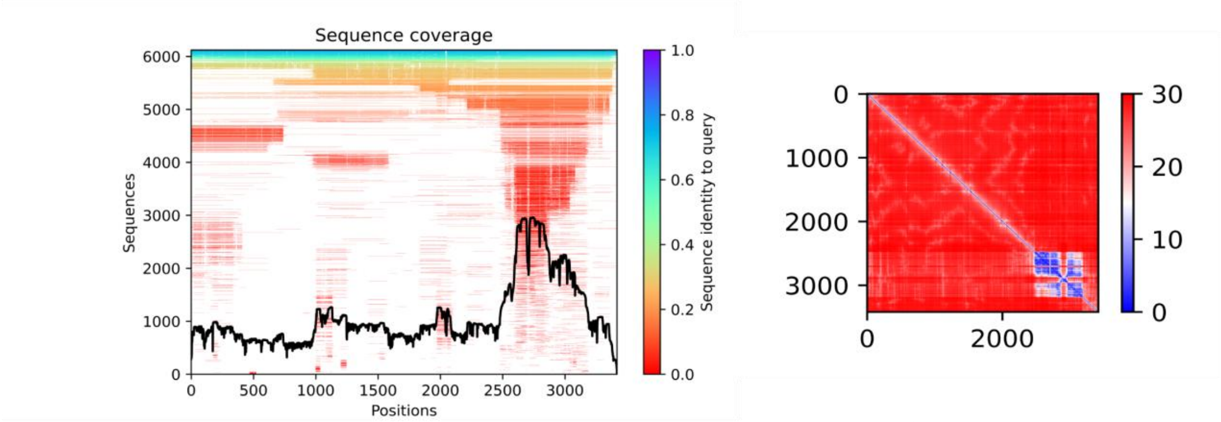


**B**

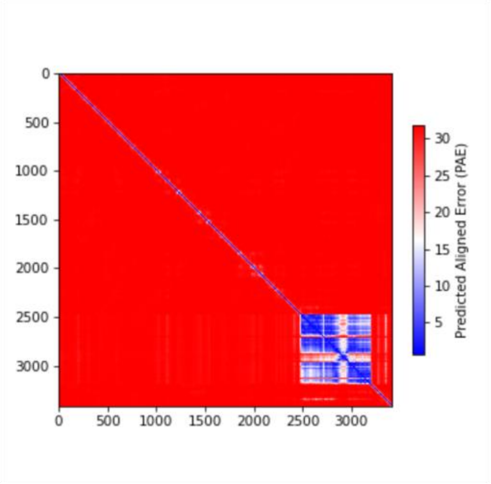


**Supplementary Figure 3 Multiple Sequence Alignment (MSA) analysis and Predicted Align Error matrices of BRCA2 predictions A.** Left, Alphafold 2 Multiple Sequence Alignment (MSA). In this representation, the black line defines the coverage of the input sequence with respect to the total number of aligned sequences. The heat-map representation provides a metric of the identity score of each aligned sequence: sequences are ordered from largest identity (top) to lowest identity (bottom) and colored by their identity score according to the color scale reported on the right (red lower identity, blue higher identity). White regions are not covered by the sequence alignment. Right, Predicted Align Error (PAE) plot of the Alphafold2 BRCA2 prediction, providing the expected positional error at residue x if the predicted structure is aligned on residue y. **B.** Predicted Align Error (PAE) plot of the Alphafold3 BRCA2 prediction.

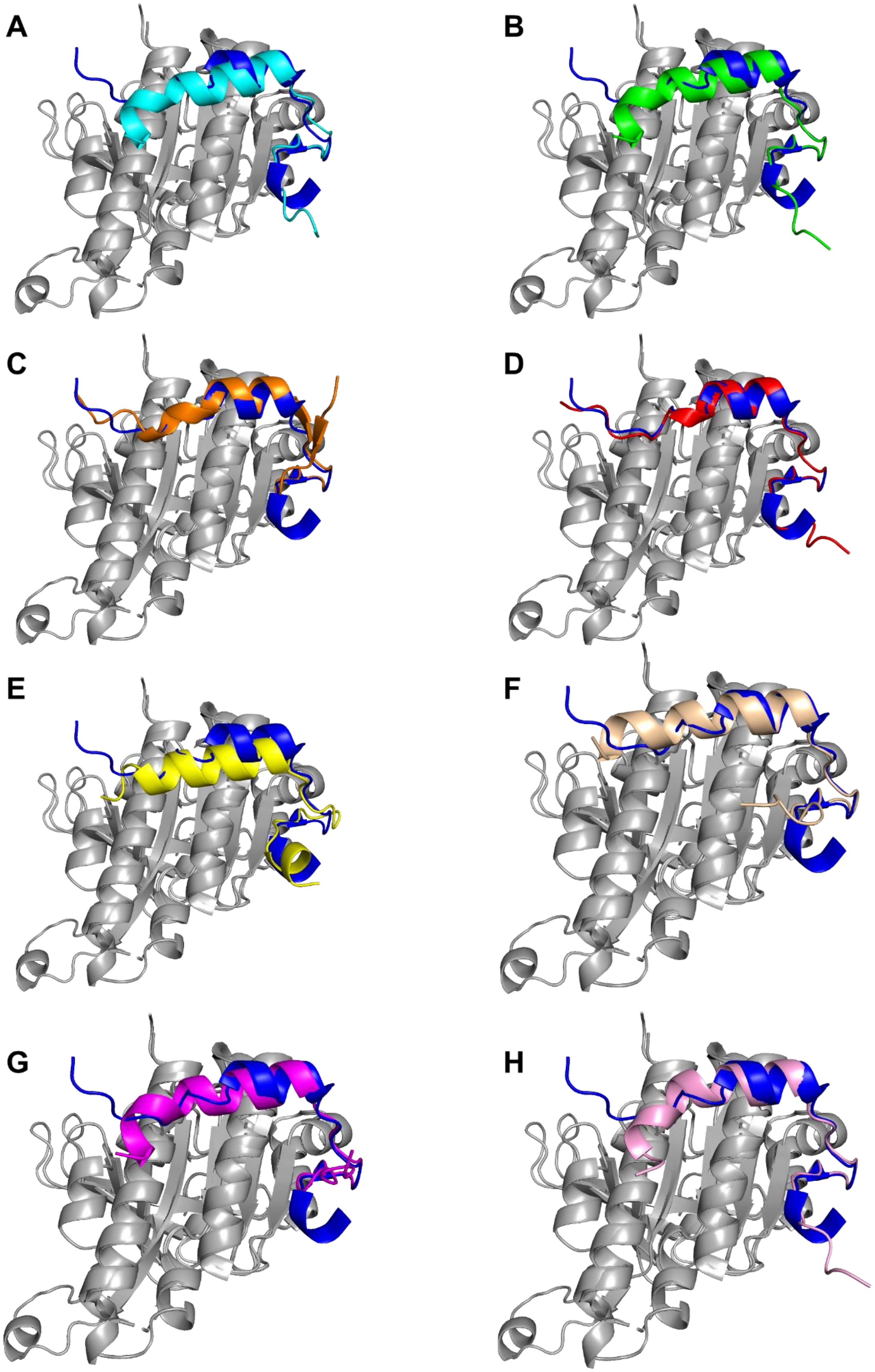


**Supplementary Figure 4 AlphaFold prediction of the Δ97-RAD51/BRC repeats complex.** Peptides have been presented with different colors: **A. BRC1** (cyan) **B. BRC2** (green) **C. BRC3** (orange) **D. BRC4** (red) **E. BRC5** (yellow) **F. BRC6** (wheat) **G. BRC7** (magenta) **H. BRC8** (pink). Predictions have been aligned to the RAD51-BRC4 complex structure (PDB entry: 1n0w) and the experimentally determined structure of the BRC4 peptide is displayed in dark blue.

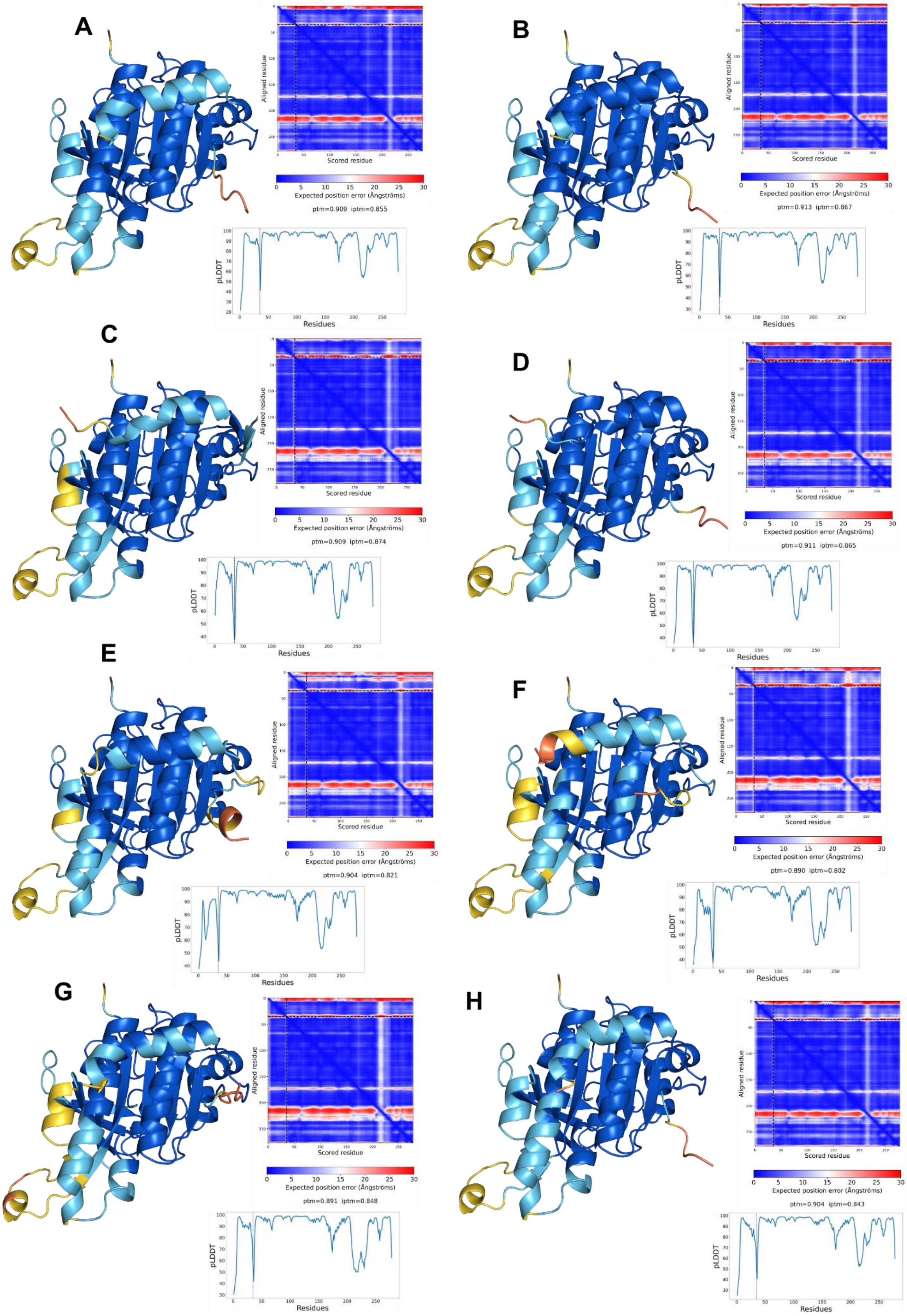


**Supplementary Figure 5 AlphaFold prediction of the Δ97-RAD51/BRC repeats complex, colored by pLDDT A. BRC1 B. BRC2 C. BRC3 D. BRC4 E. BRC5 F. BRC6 G. BRC7 H. BRC8**. On the right of each model the PAE matrices are reported along with ptm and iptm scores.

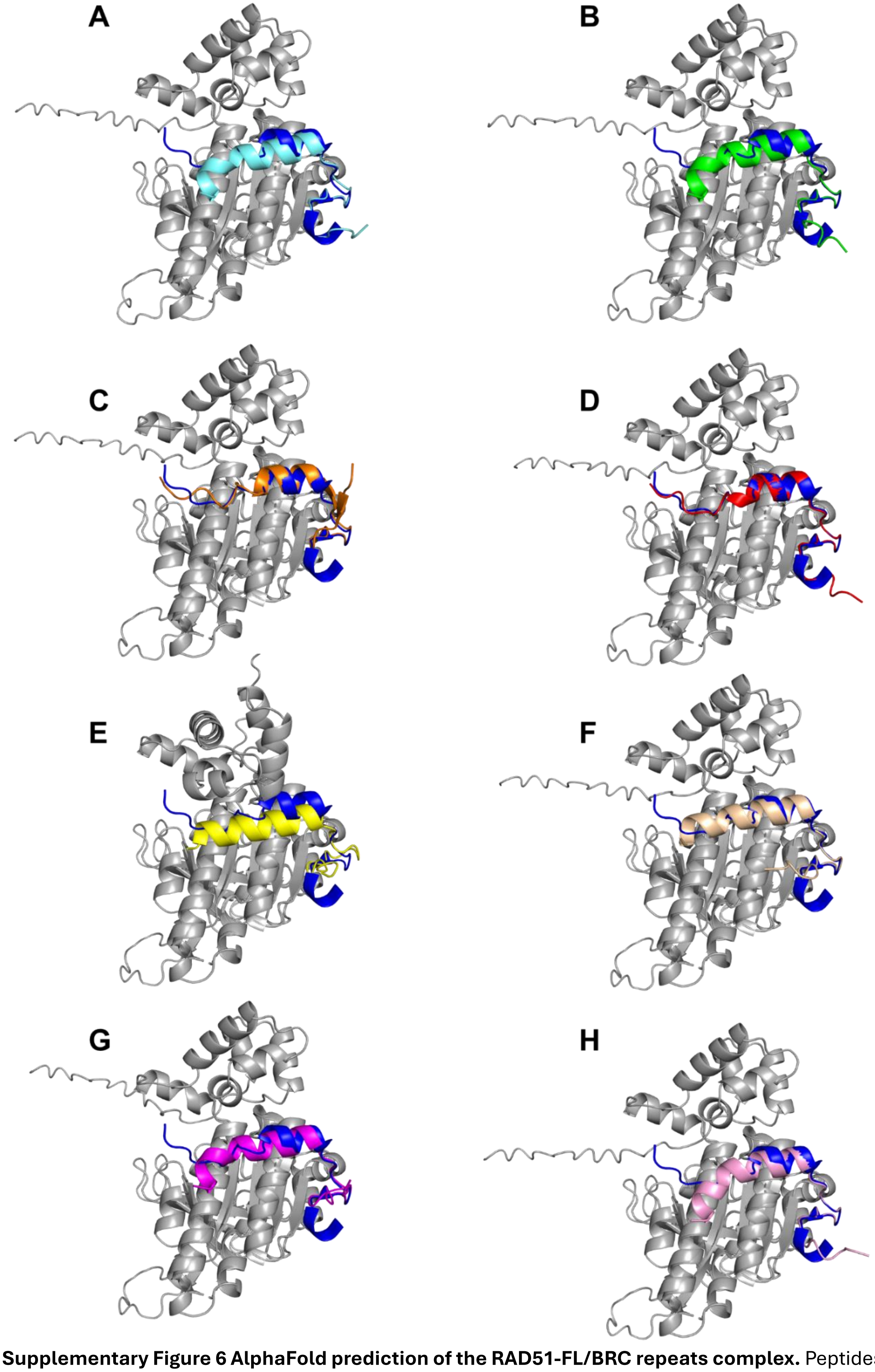


**Supplementary Figure 6 AlphaFold prediction of the RAD51-FL/BRC repeats complex.** Peptides have been presented with different colors: **A. BRC1** (cyan) **B. BRC2** (green) **C. BRC3** (orange) **D. BRC4** (red) **E. BRC5** (yellow) **F. BRC6** (wheat) **G. BRC7** (magenta) **H. BRC8** (pink). Predictions have been aligned to the available RAD51-BRC4 complex structure (PDB entry: 1n0w). The experimentally determined structure of the BRC4 peptide is displayed in dark blue.

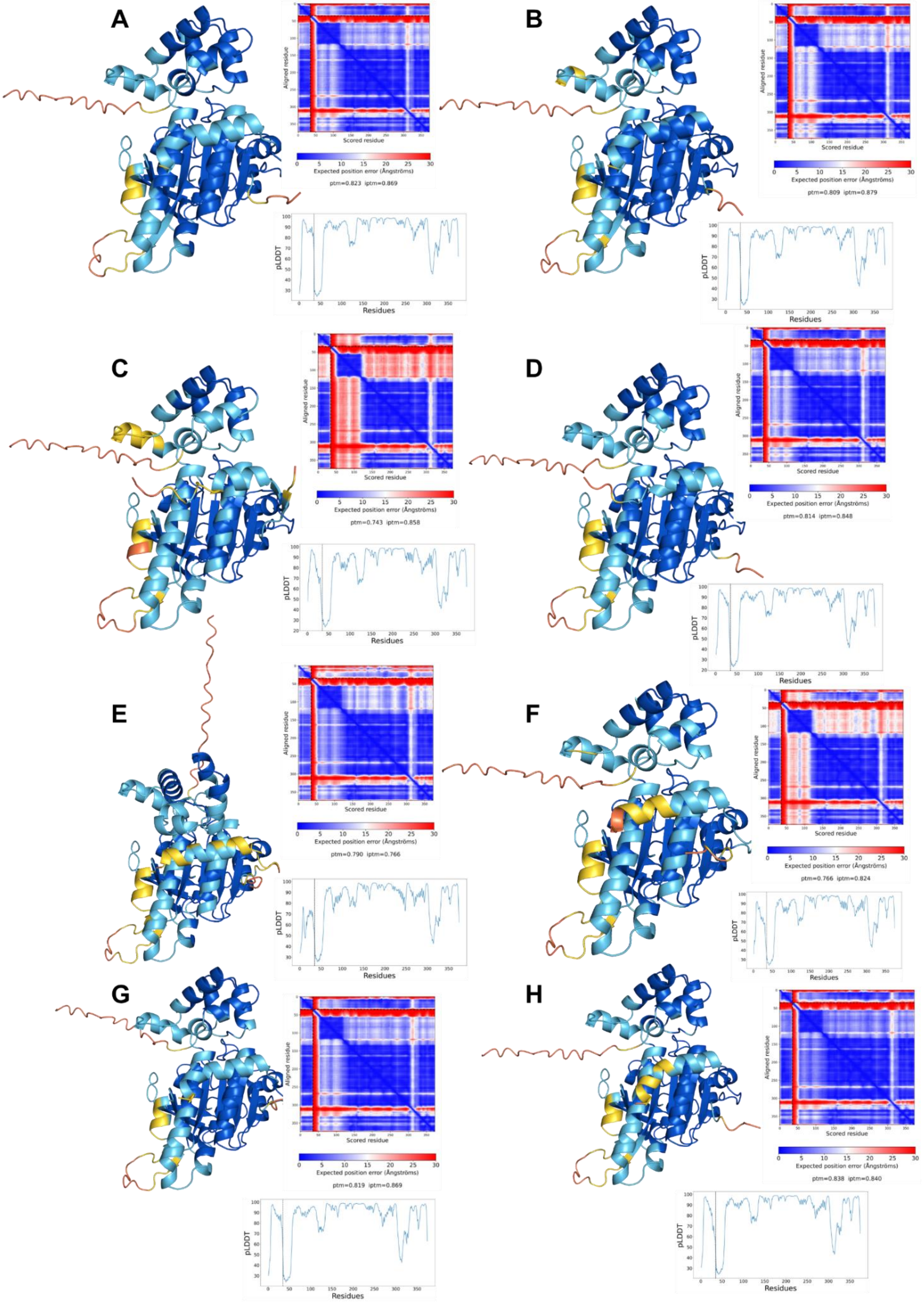


**Supplementary Figure 7 AlphaFold prediction of the RAD51-FL/BRC repeats complex, colored by pLDDT A. BRC1 B. BRC2 C. BRC3 D. BRC4 E. BRC5 F. BRC6 G. BRC7 H. BRC8**. On the right of each model the PAE matrices are reported along with ptm and iptm scores.

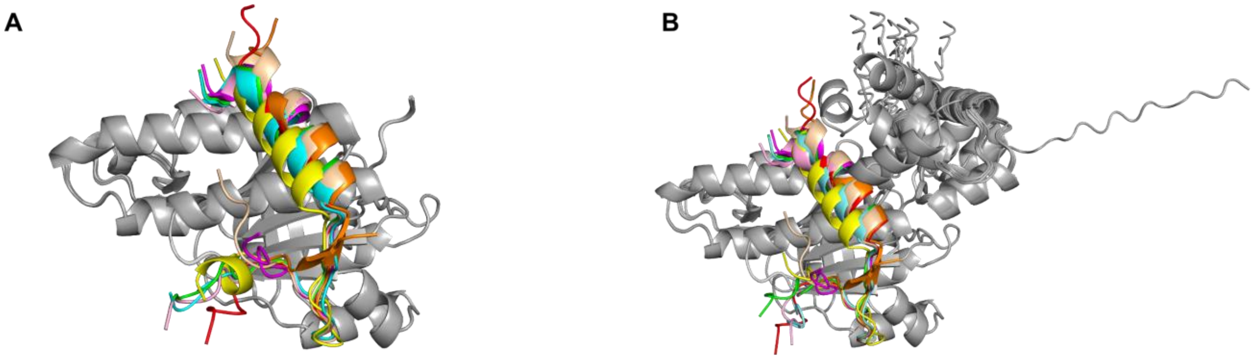


**Supplementary Figure 8 AlphaFold predictions of the BRC repeats-RAD51 binding. A.** Overlay of the predictions generated with BRC1-8 and Δ97-RAD51 **B.** Overlay of the predictions generated with BRC1-8 and RAD51

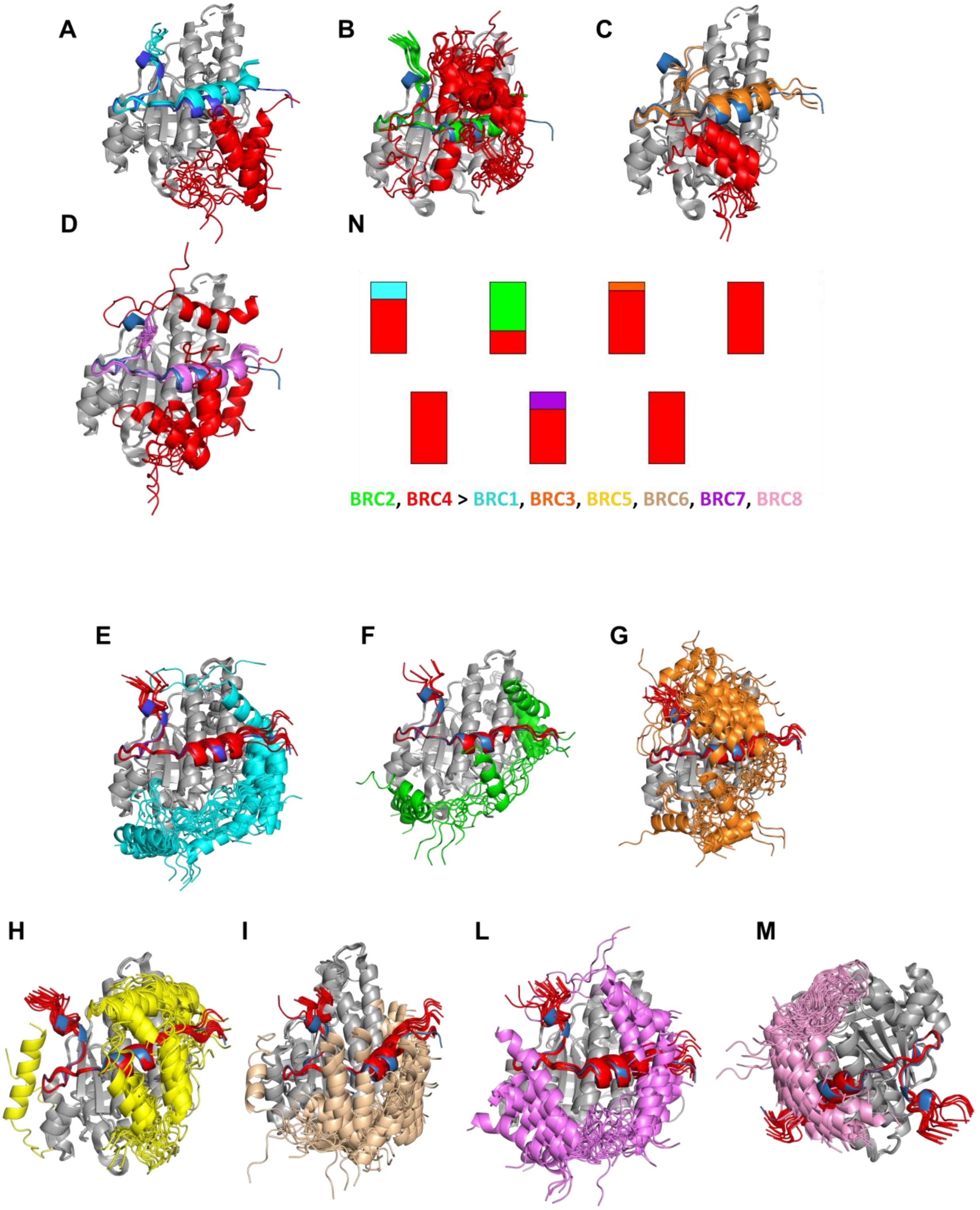


**Supplementary Figure 9** Competitive AlphaFold predictions of the BRC-repeats vs BRC4, using a N-terminal truncated RAD51 (Δ97-RAD51) as receptor. Simulations' outputs were aligned to the available RAD51-BRC4 complex structure (PDB: 1n0w) in which the experimentally determined BRC4 peptide is shown in blue. Overlay of simulations using RAD51-FL where BRC4 (red) was misplaced while **A.** BRC1 (cyan) **B.** BRC2 (green) **C.** BRC3 (orange) **D.** BRC7 (violet) were positioned in the BRC4 binding site. Overlay of simulations where the BRC4 (red) peptide is correctly modelled in its binding site and the other BRC-repeats: **E.** BRC1 (cyan) **F.** BRC2 (green) **G.** BRC3 (orange) **H.** BRC5 (yellow) **I.** BRC6 (wheat) **L.** BRC7 (violet) **M.** BRC8 (pink) display a different binding mode. In panel **N.** a summary graph reports the results of competitive simulations between BRC4 and the other BRC-repeats (25 models for each competitive simulation).

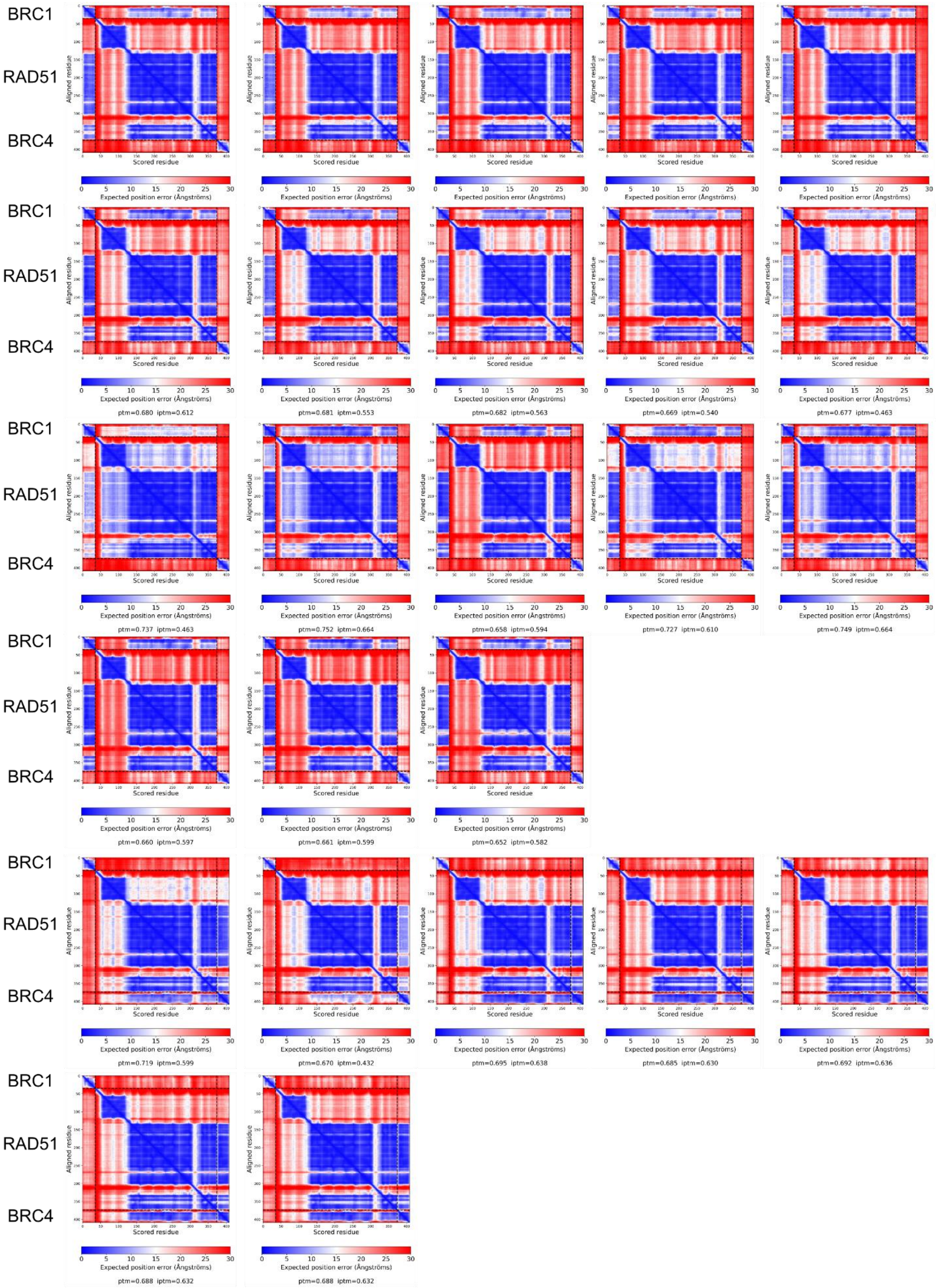


**Supplementary Figure 10 Predicted Align Error plots of predictions carried out with BRC1 and BRC4 sequences utilizing RAD51-FL sequence as receptor.** Predicted Align Error (PAE) plots provide the expected positional error at residue x if the predicted structure is aligned on residue y.

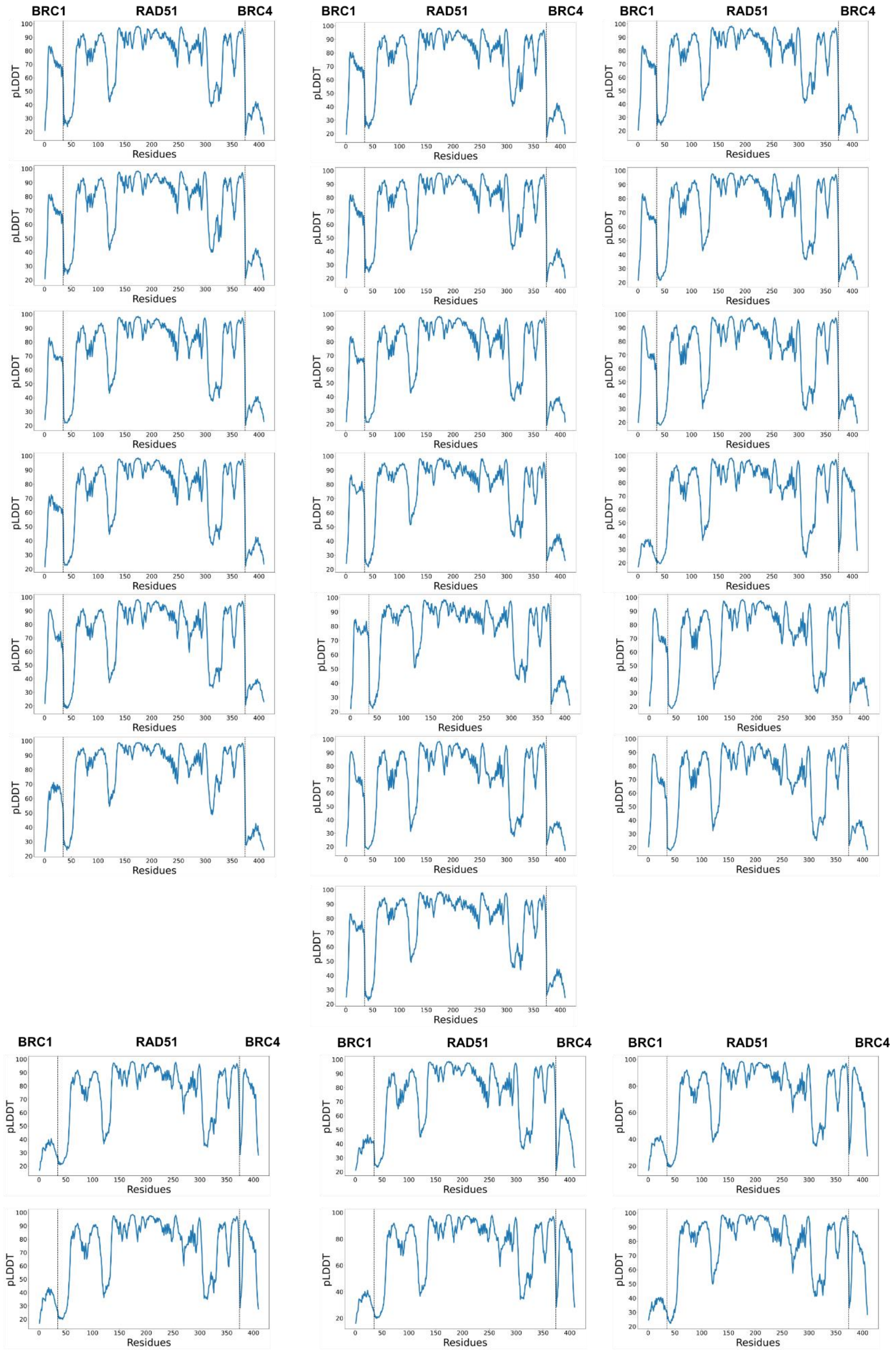


**Supplementary Figure 11 predicted Local Distance Difference Test (pLDDT) predictions carried out with BRC1 and BRC4 sequences utilizing RAD51-FL sequence as receptor.** pLDDT representation provides a per-residue accuracy metric.

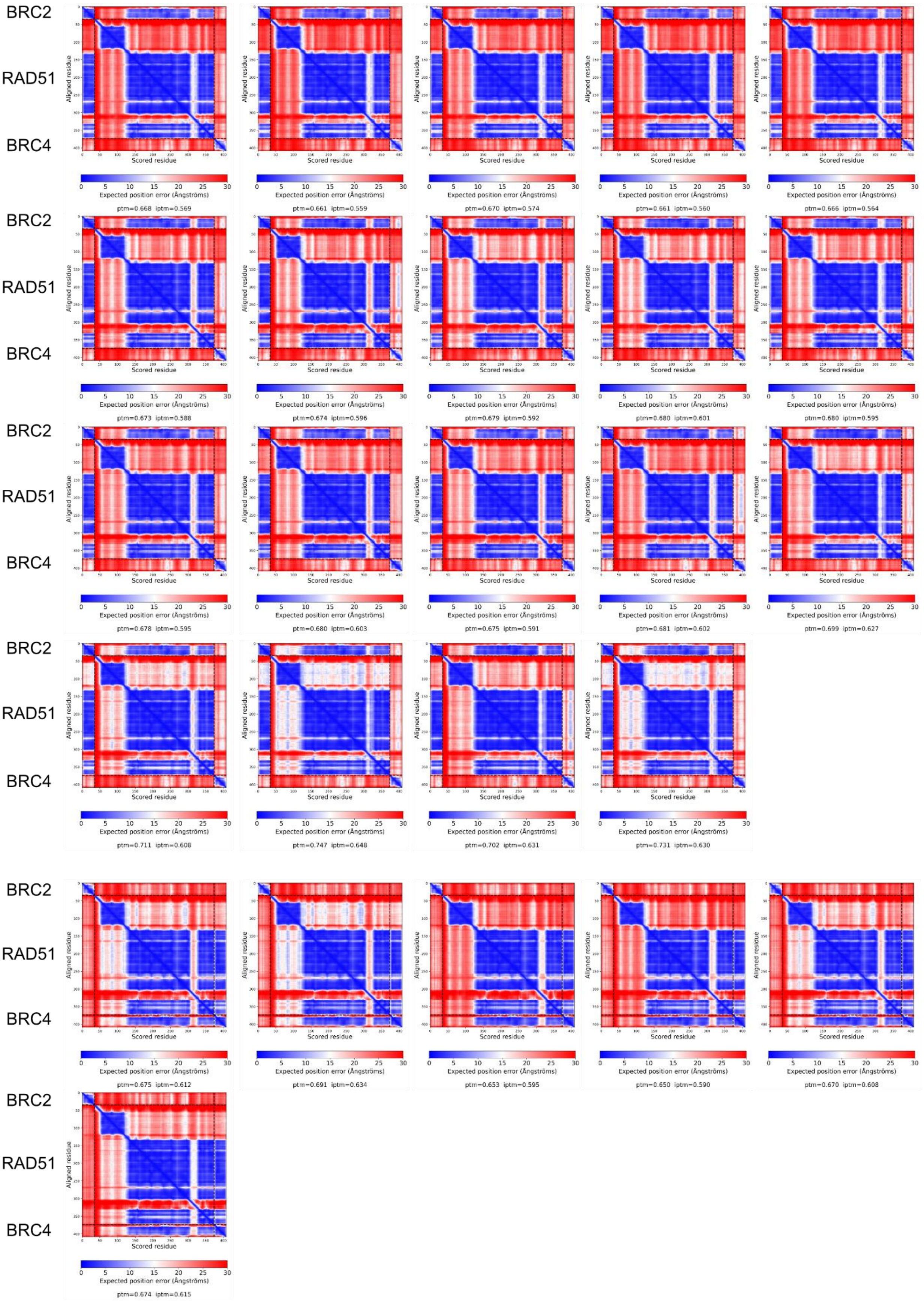


**Supplementary Figure 12 Predicted Align Error plots of predictions carried out with BRC2 and BRC4 sequences utilizing RAD51-FL sequence as receptor.** Predicted Align Error (PAE) plots provide the expected positional error at residue x if the predicted structure is aligned on residue y.

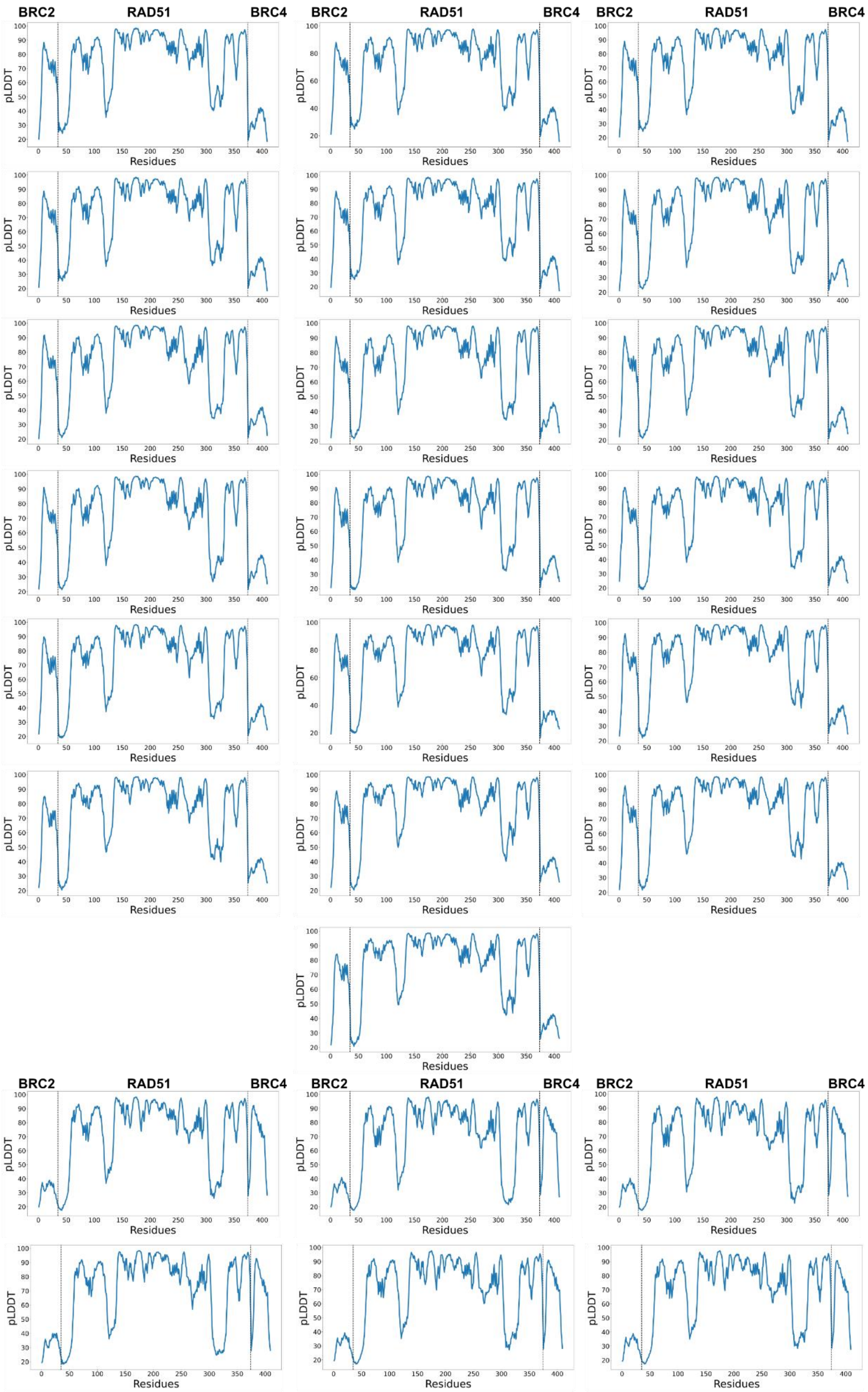


**Supplementary Figure 13 predicted Local Distance Difference Test (pLDDT) predictions carried out with BRC2 and BRC4 sequences utilizing RAD51-FL sequence as receptor.** pLDDT representation provides a per-residue accuracy metric.

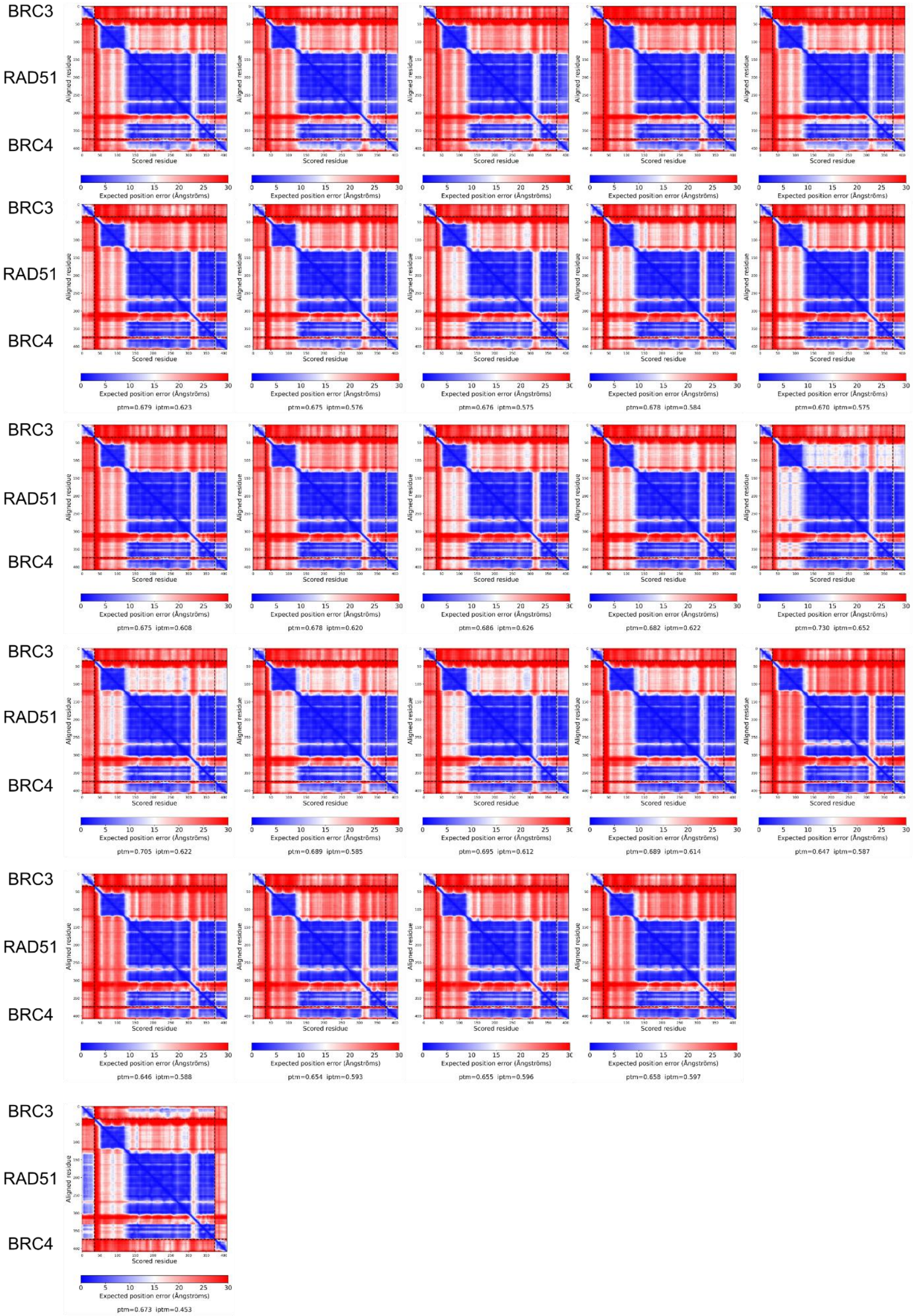


**Supplementary Figure 14 Predicted Align Error plots of predictions carried out with BRC3 and BRC4 sequences utilizing RAD51-FL sequence as receptor.** Predicted Align Error (PAE) plots provide the expected positional error at residue x if the predicted structure is aligned on residue y.

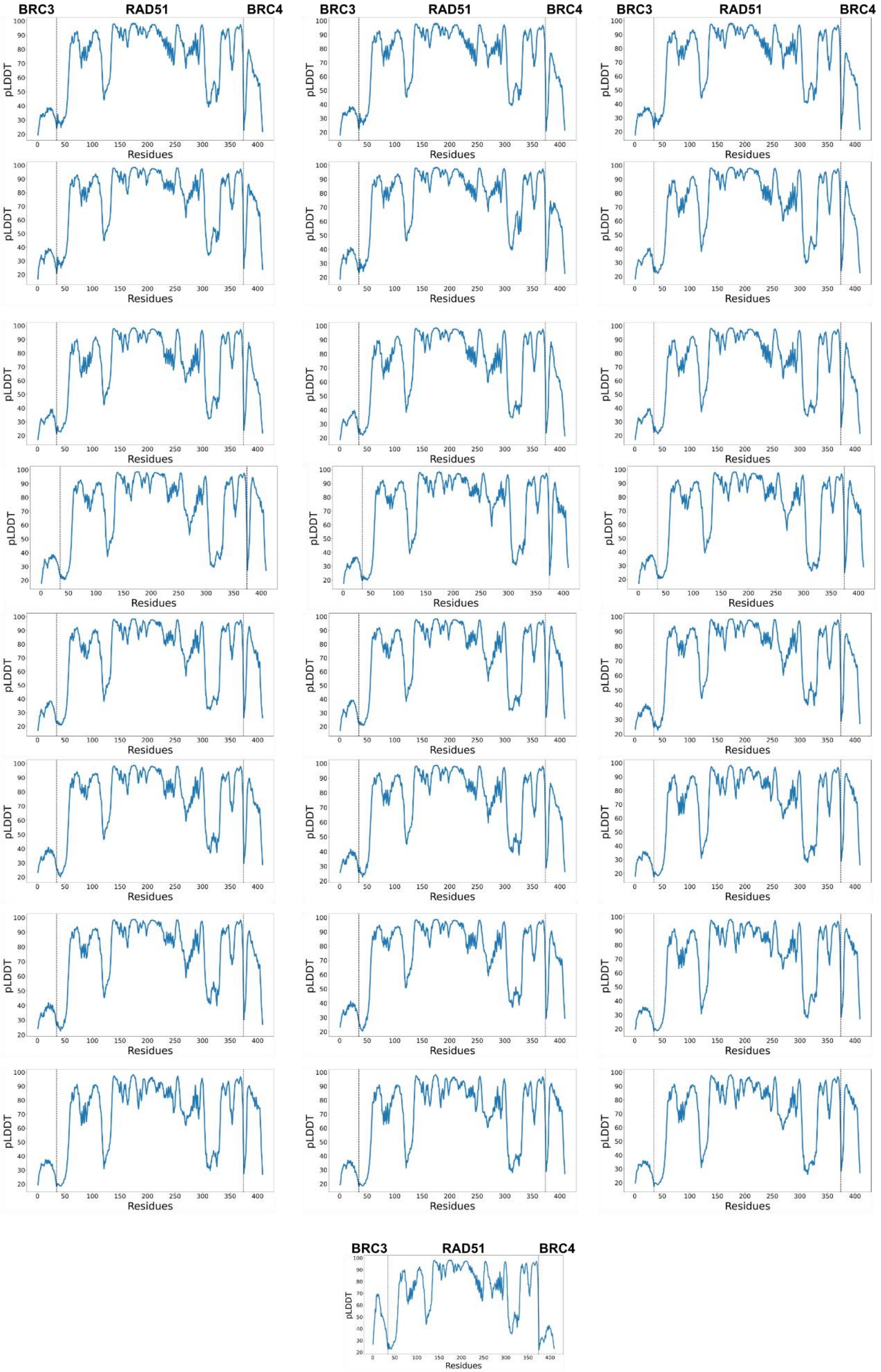


**Supplementary Figure 15 predicted Local Distance Difference Test (pLDDT) predictions carried out with BRC3 and BRC4 sequences utilizing RAD51-FL sequence as receptor.** pLDDT representation provides a per-residue accuracy metric.

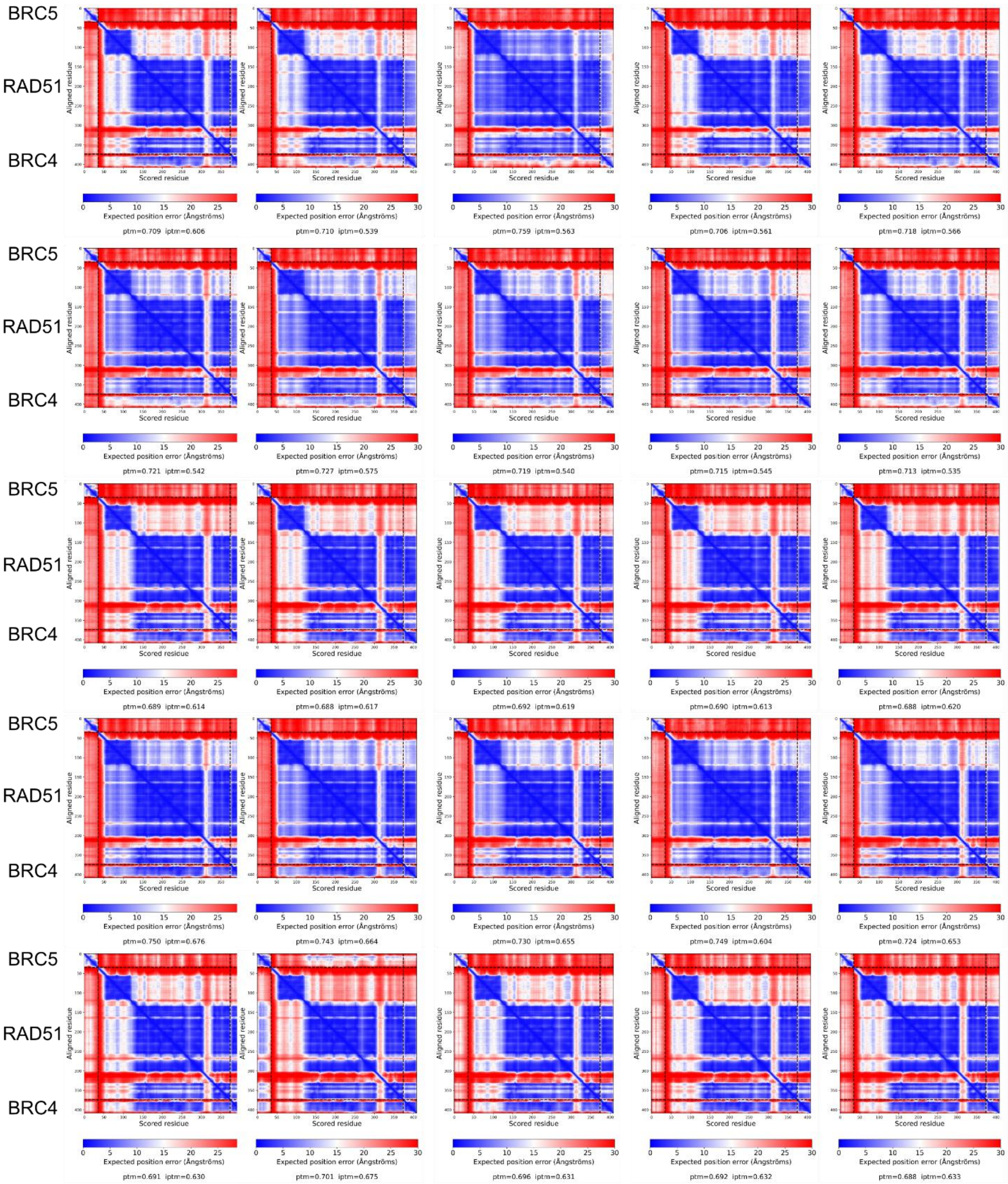


**Supplementary Figure 16 Predicted Align Error plots of predictions carried out with BRC5 and BRC4 sequences utilizing RAD51-FL sequence as receptor.** Predicted Align Error (PAE) plots provide the expected positional error at residue x if the predicted structure is aligned on residue y.

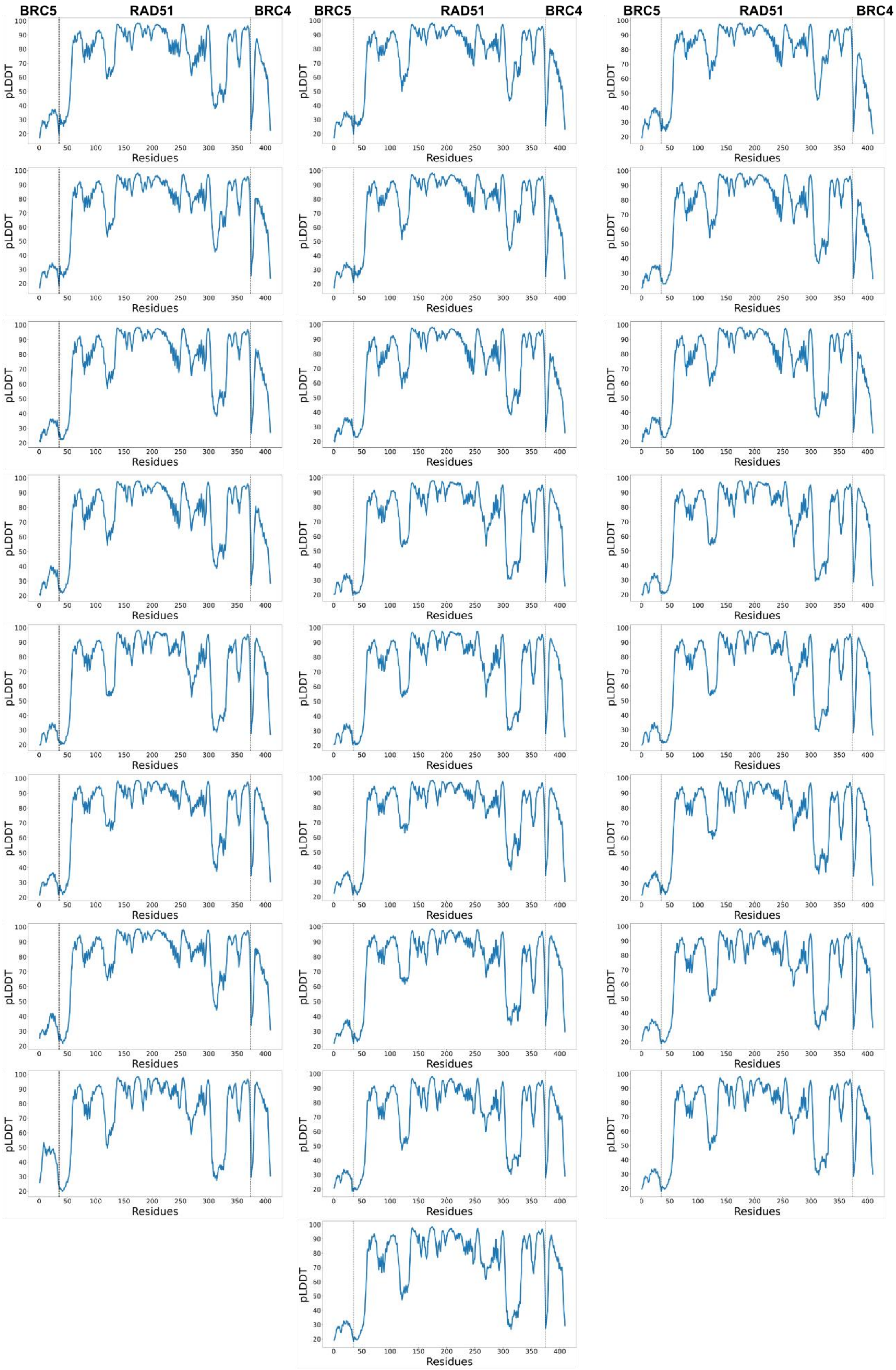


**Supplementary Figure 17 predicted Local Distance Difference Test (pLDDT) predictions carried out with BRC5 and BRC4 sequences utilizing RAD51-FL sequence as receptor.** pLDDT representation provides a per-residue accuracy metric.

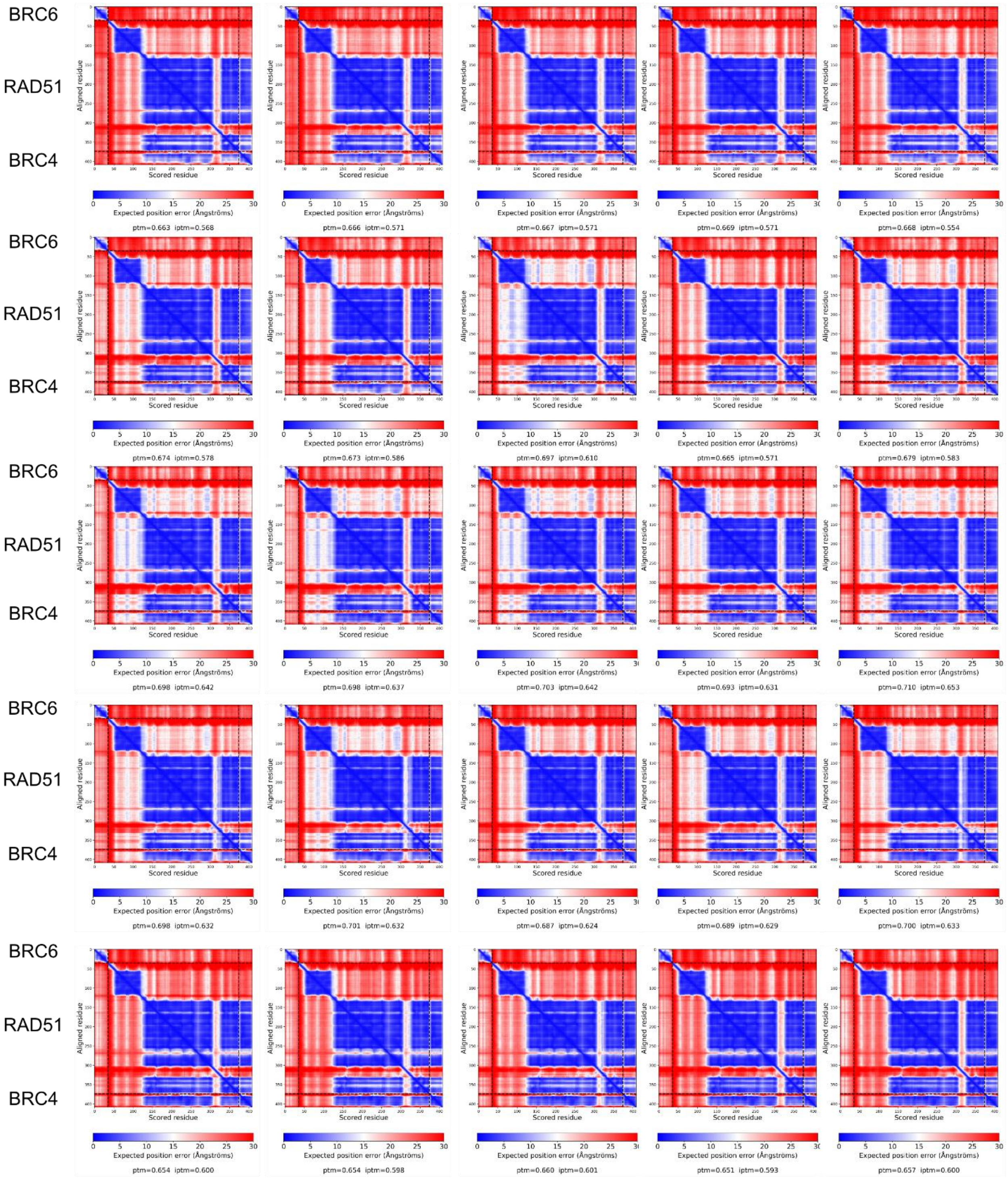


**Supplementary Figure 18 Predicted Align Error plots of predictions carried out with BRC6 and BRC4 sequences utilizing RAD51-FL sequence as receptor.** Predicted Align Error (PAE) plots provide the expected positional error at residue x if the predicted structure is aligned on residue y.

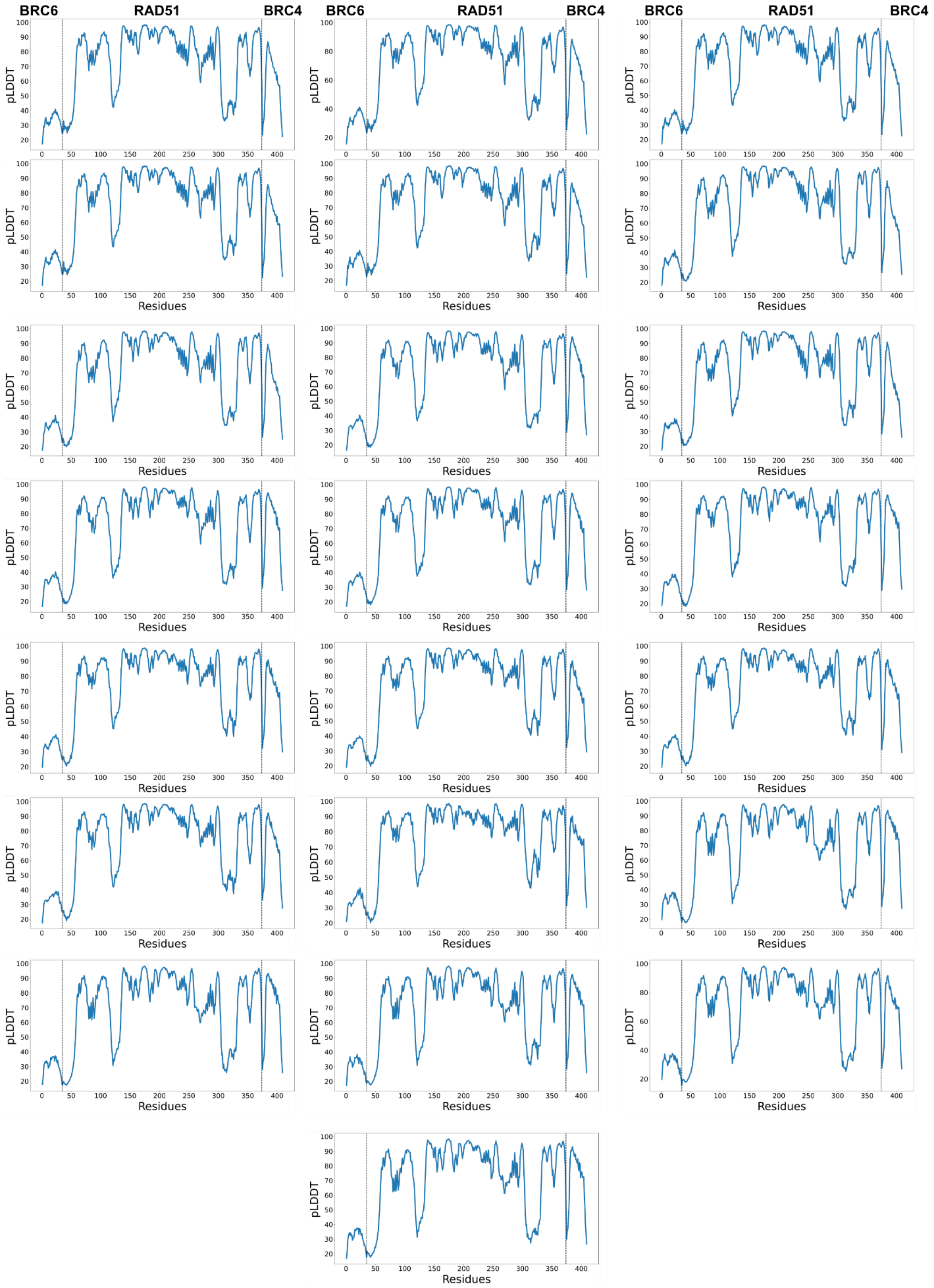


**Supplementary Figure 19 predicted Local Distance Difference Test (pLDDT) predictions carried out with BRC6 and BRC4 sequences utilizing RAD51-FL sequence as receptor.** pLDDT representation provides a per-residue accuracy metric.

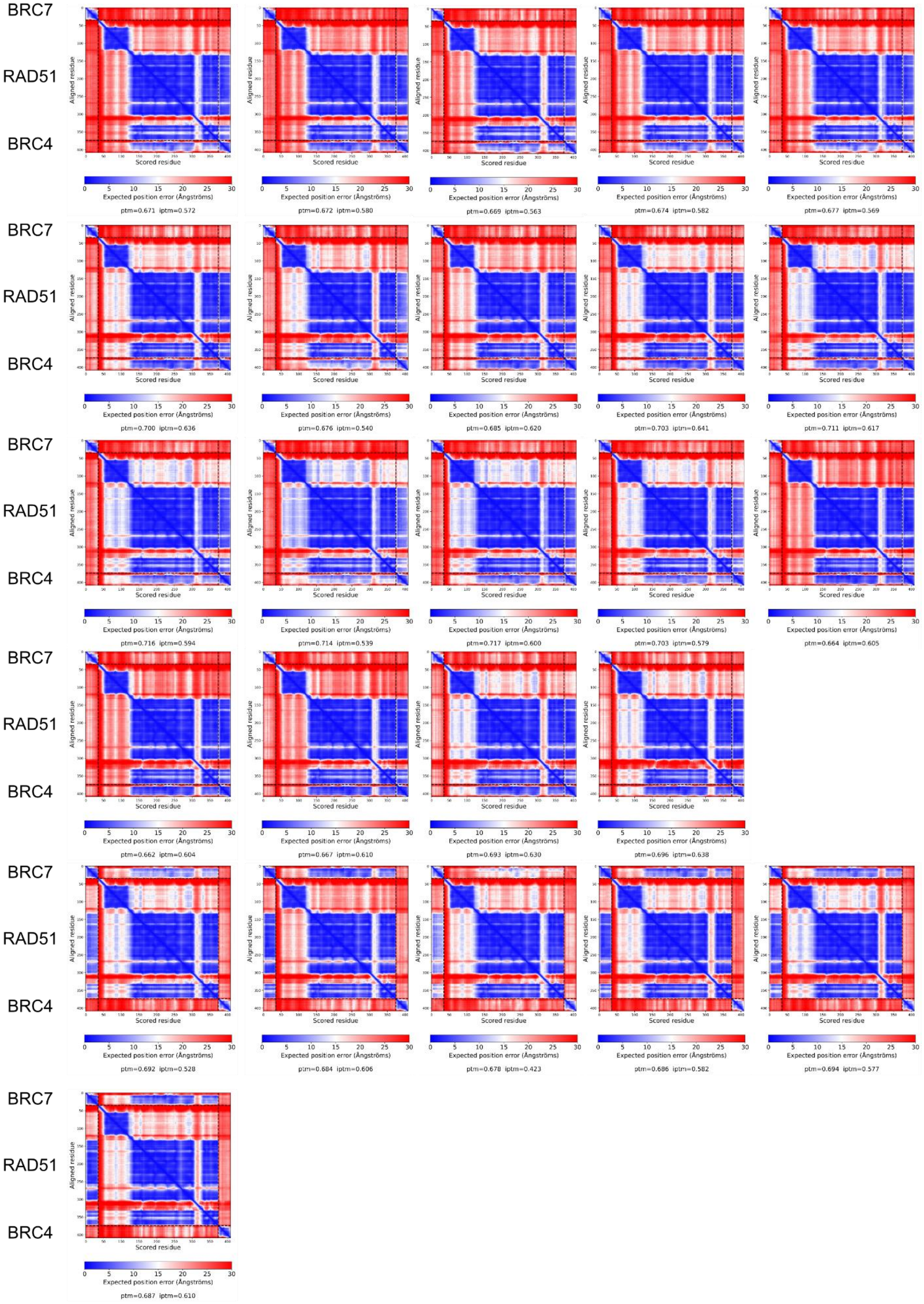


**Supplementary Figure 20 Predicted Align Error plots of predictions carried out with BRC7 and BRC4 sequences utilizing RAD51-FL sequence as receptor.** Predicted Align Error (PAE) plots provide the expected positional error at residue x if the predicted structure is aligned on residue y.

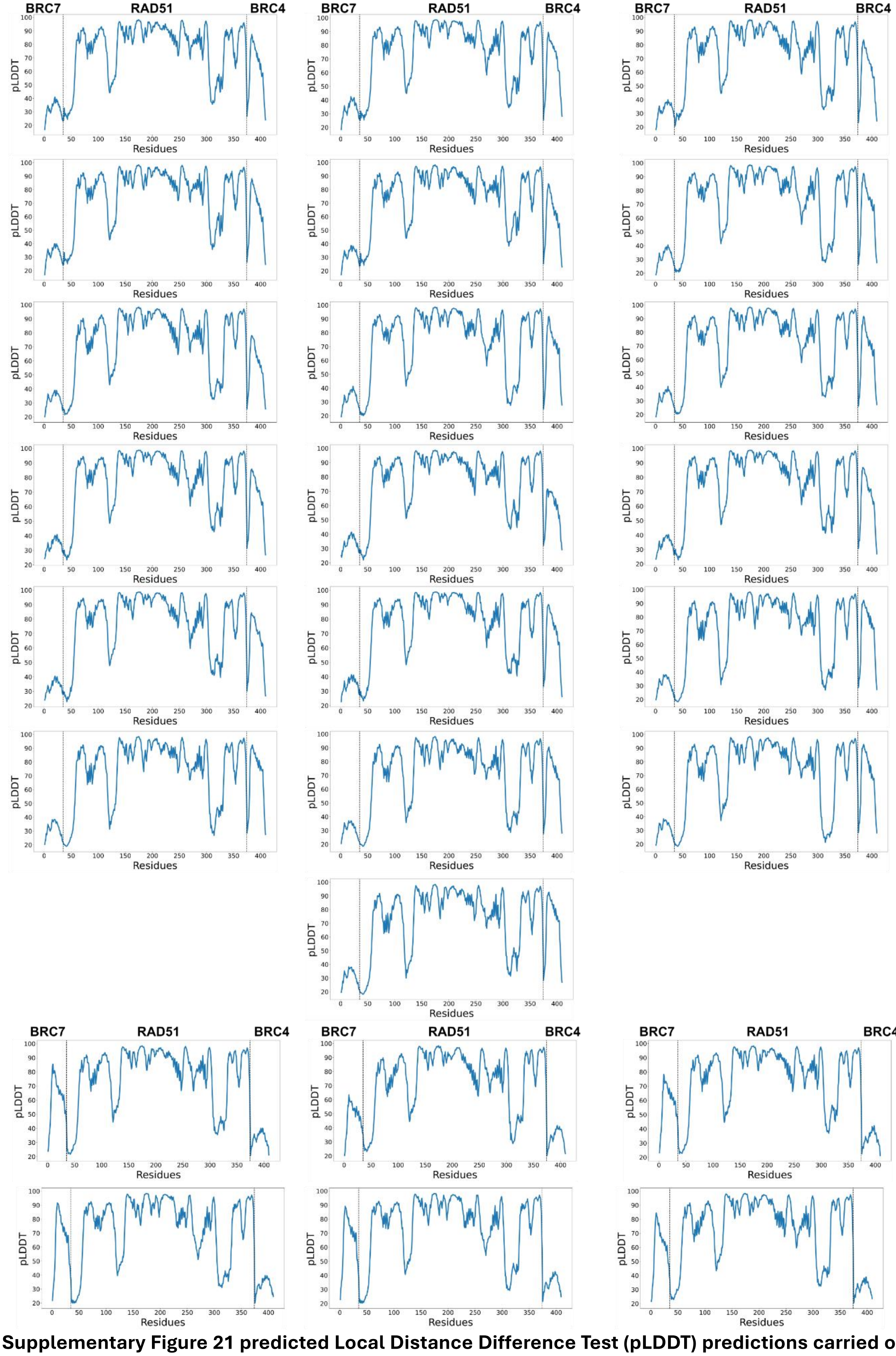


**Supplementary Figure 21 predicted Local Distance Difference Test (pLDDT) predictions carried out with BRC7 and BRC4 sequences utilizing RAD51-FL sequence as receptor.** pLDDT representation provides a per-residue accuracy metric.

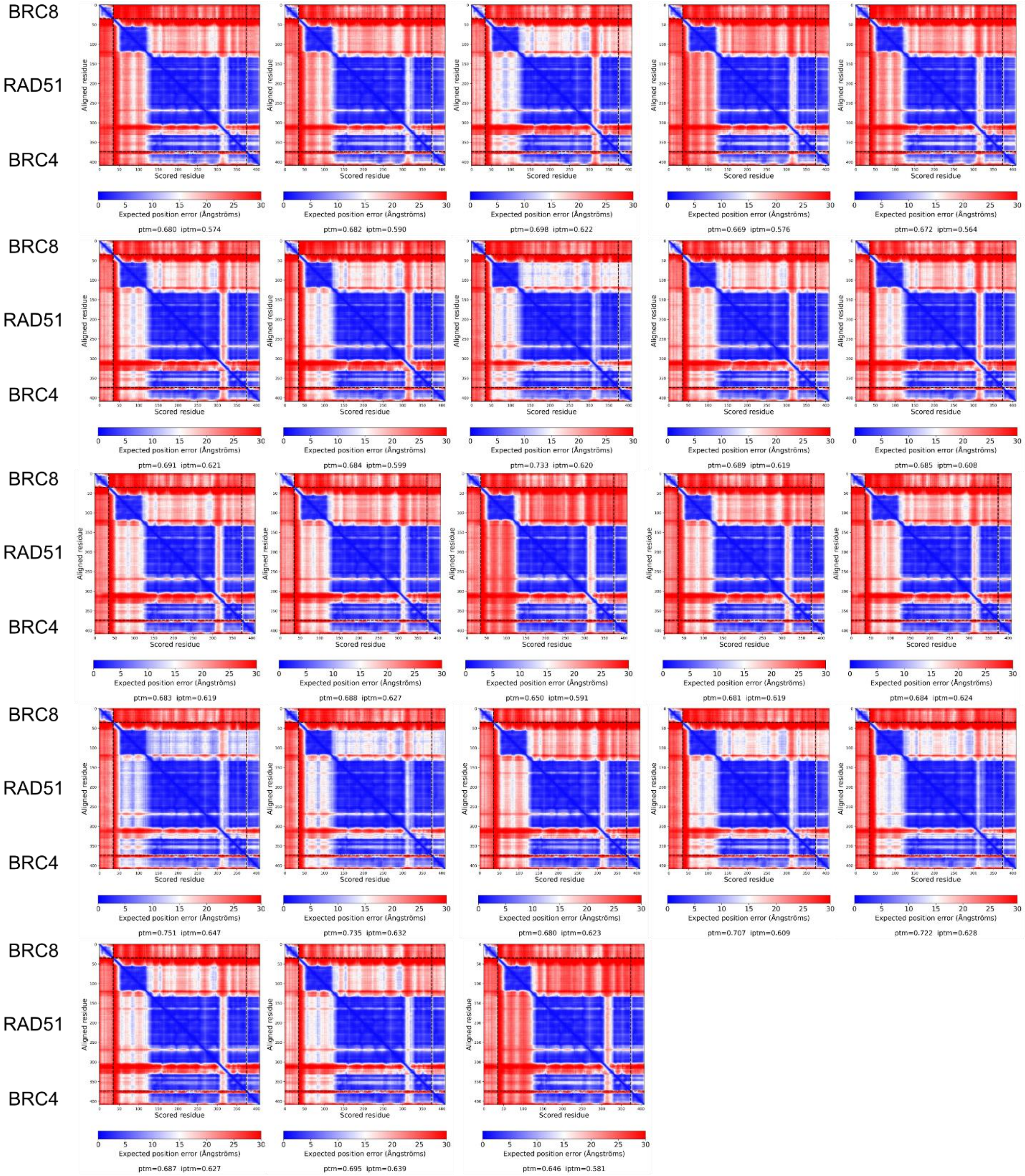


**Supplementary Figure 22 Predicted Align Error plots of predictions carried out with BRC8 and BRC4 sequences utilizing RAD51-FL sequence as receptor.** Predicted Align Error (PAE) plots provide the expected positional error at residue x if the predicted structure is aligned on residue y.

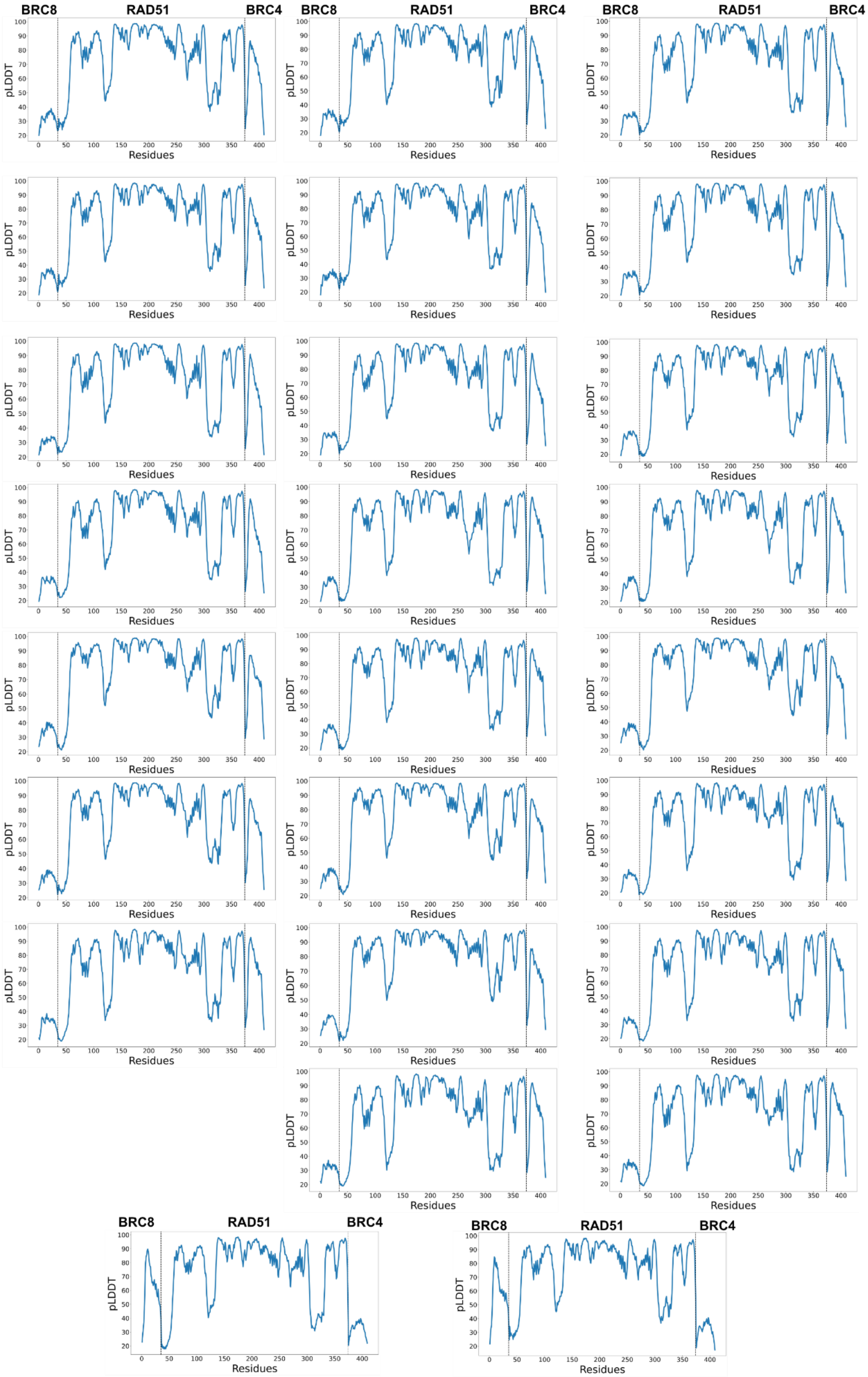


**Supplementary Figure 23 predicted Local Distance Difference Test (pLDDT) predictions carried out with BRC8 and BRC4 sequences utilizing RAD51-FL sequence as receptor.** pLDDT representation provides a per-residue accuracy metric.

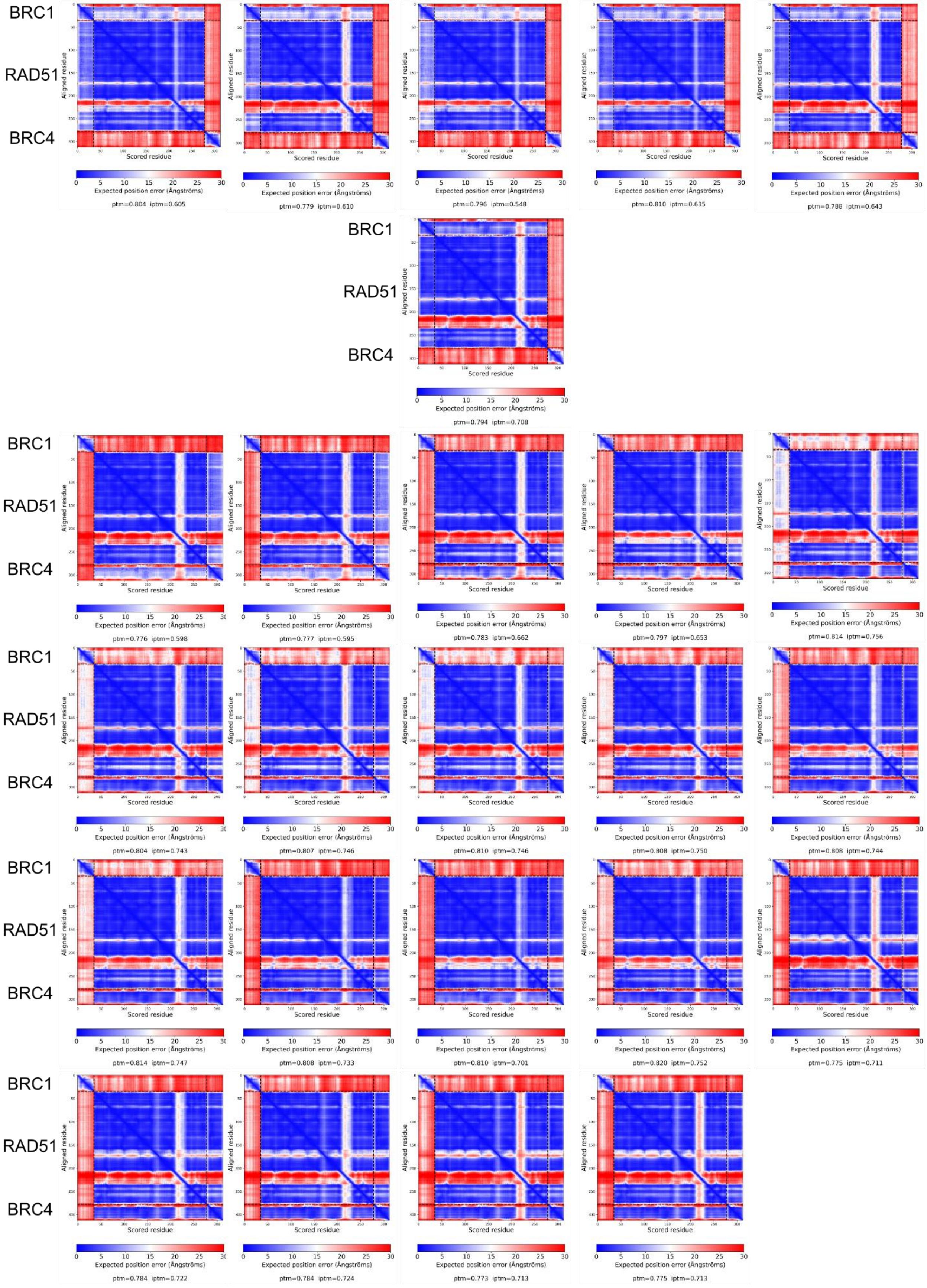


**Supplementary Figure 24 Predicted Align Error plots of predictions carried out with BRC1 and BRC4 sequences utilizing an N-terminal truncated RAD51 (Δ97-RAD51) sequence as receptor.** Predicted Align Error (PAE) plots provide the expected positional error at residue x if the predicted structure is aligned on residue y.

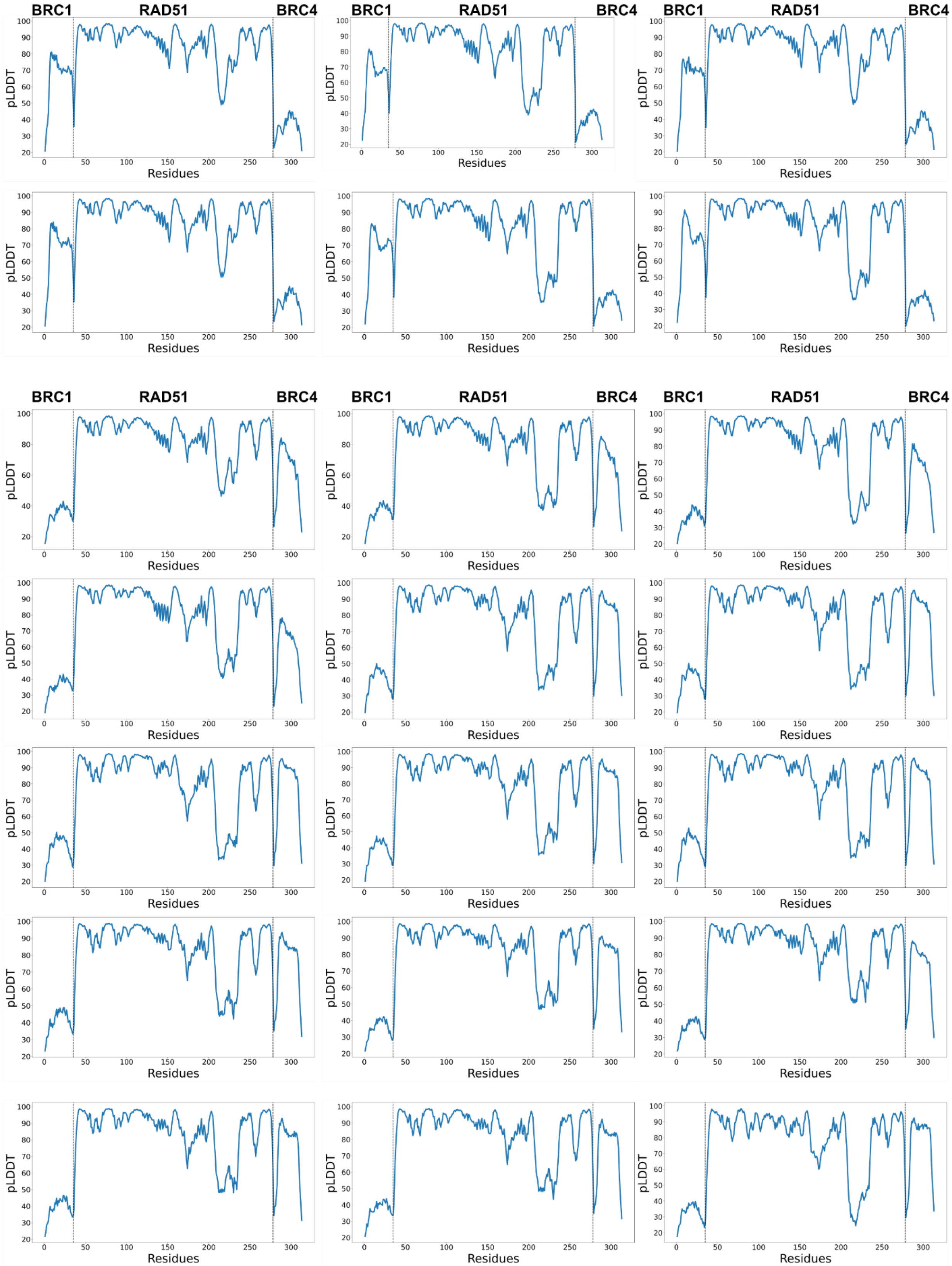


**Supplementary Figure 25 predicted Local Distance Difference Test (pLDDT) predictions carried out with BRC1 and BRC4 sequences utilizing a N-terminal truncated RAD51 (Δ97-RAD51) sequence as receptor.** pLDDT representation provides a per-residue accuracy metric.

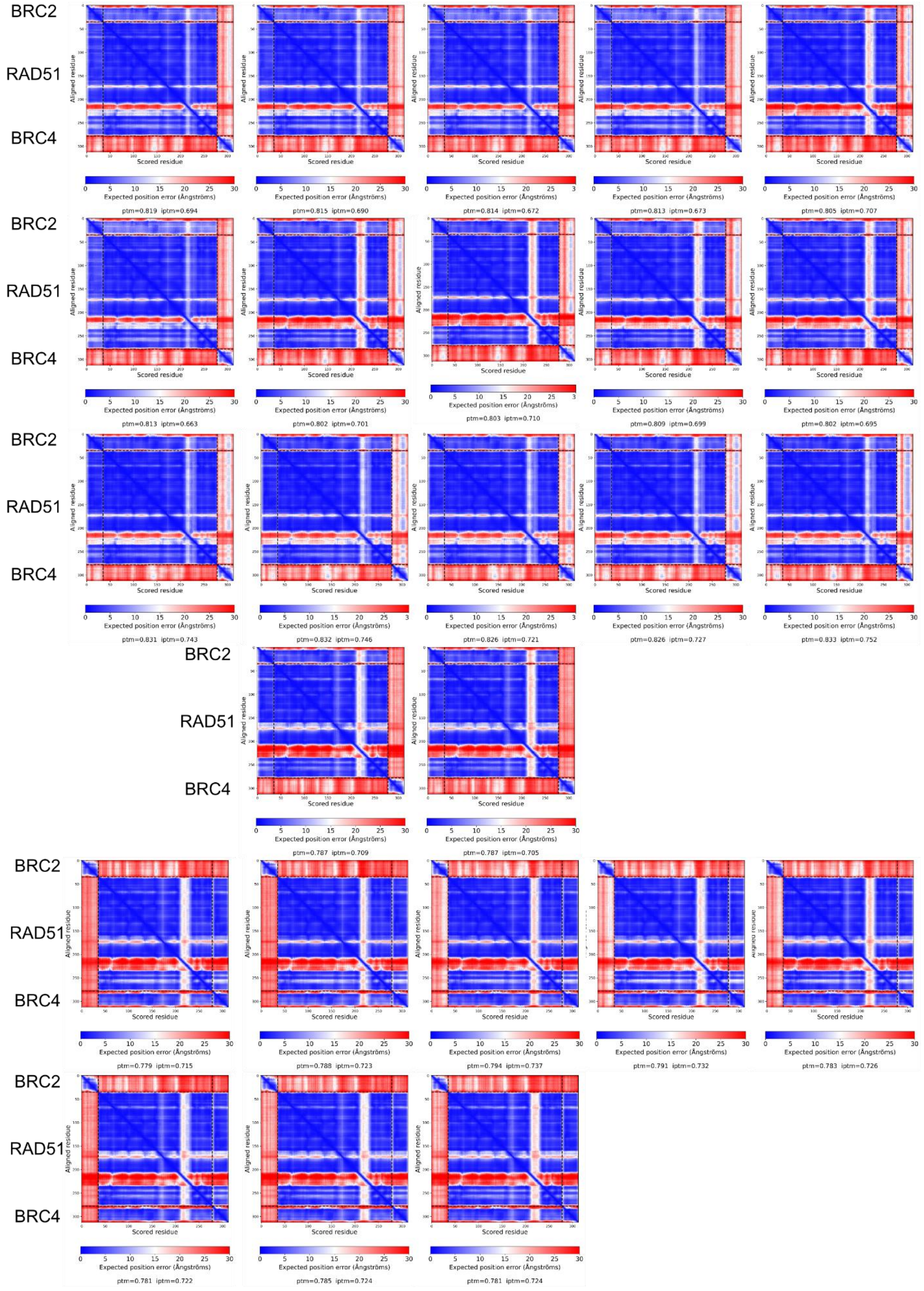


**Supplementary Figure 26 Predicted Align Error plots of predictions carried out with BRC2 and BRC4 sequences utilizing an N-terminal truncated RAD51 (Δ97-RAD51) sequence as receptor.** Predicted Align Error (PAE) plots provide the expected positional error at residue x if the predicted structure is aligned on residue y.

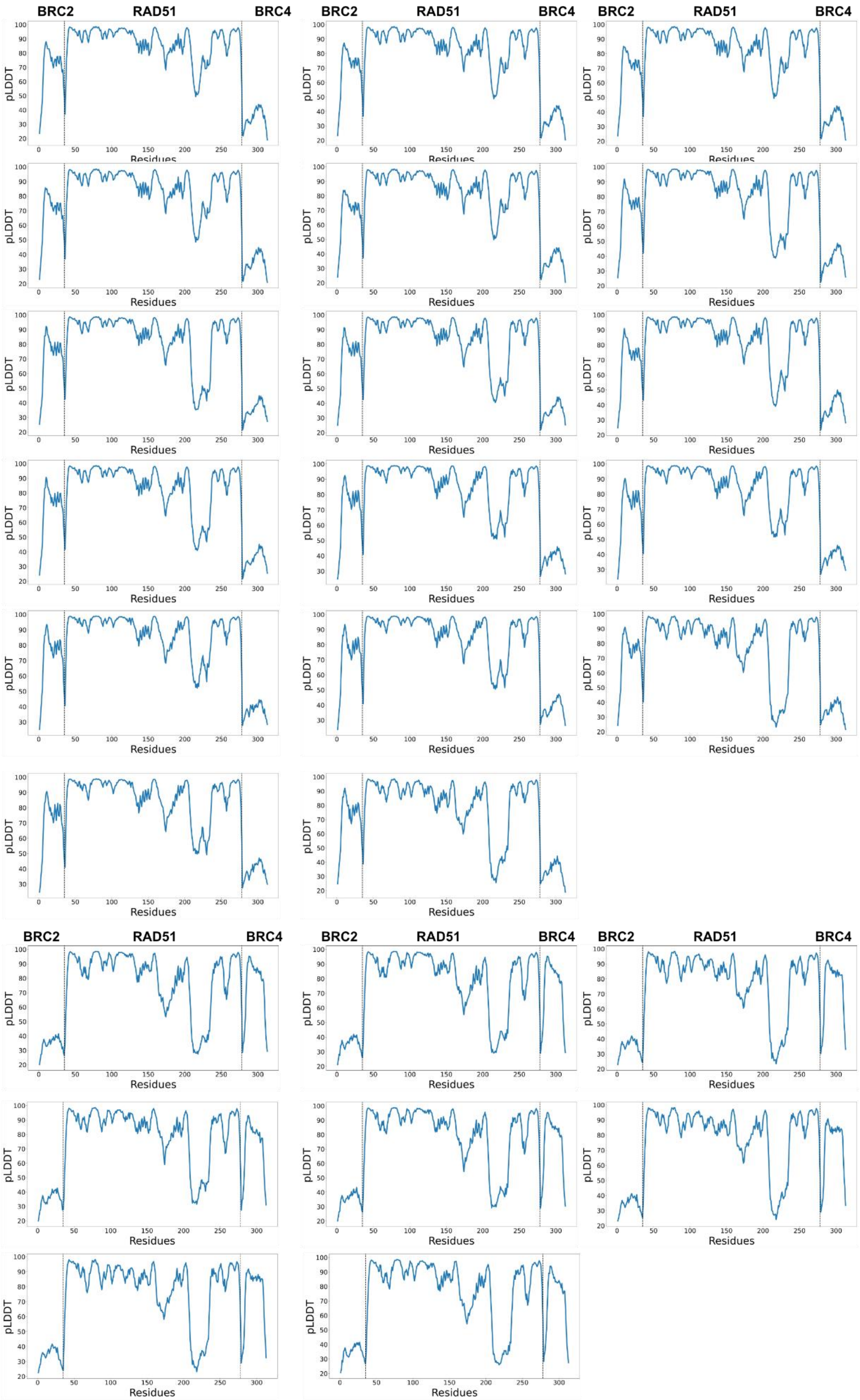


**Supplementary Figure 27 predicted Local Distance Difference Test (pLDDT) predictions carried out with BRC2 and BRC4 sequences utilizing a N-terminal truncated RAD51 (Δ97-RAD51) sequence as receptor.** pLDDT representation provides a per-residue accuracy metric.

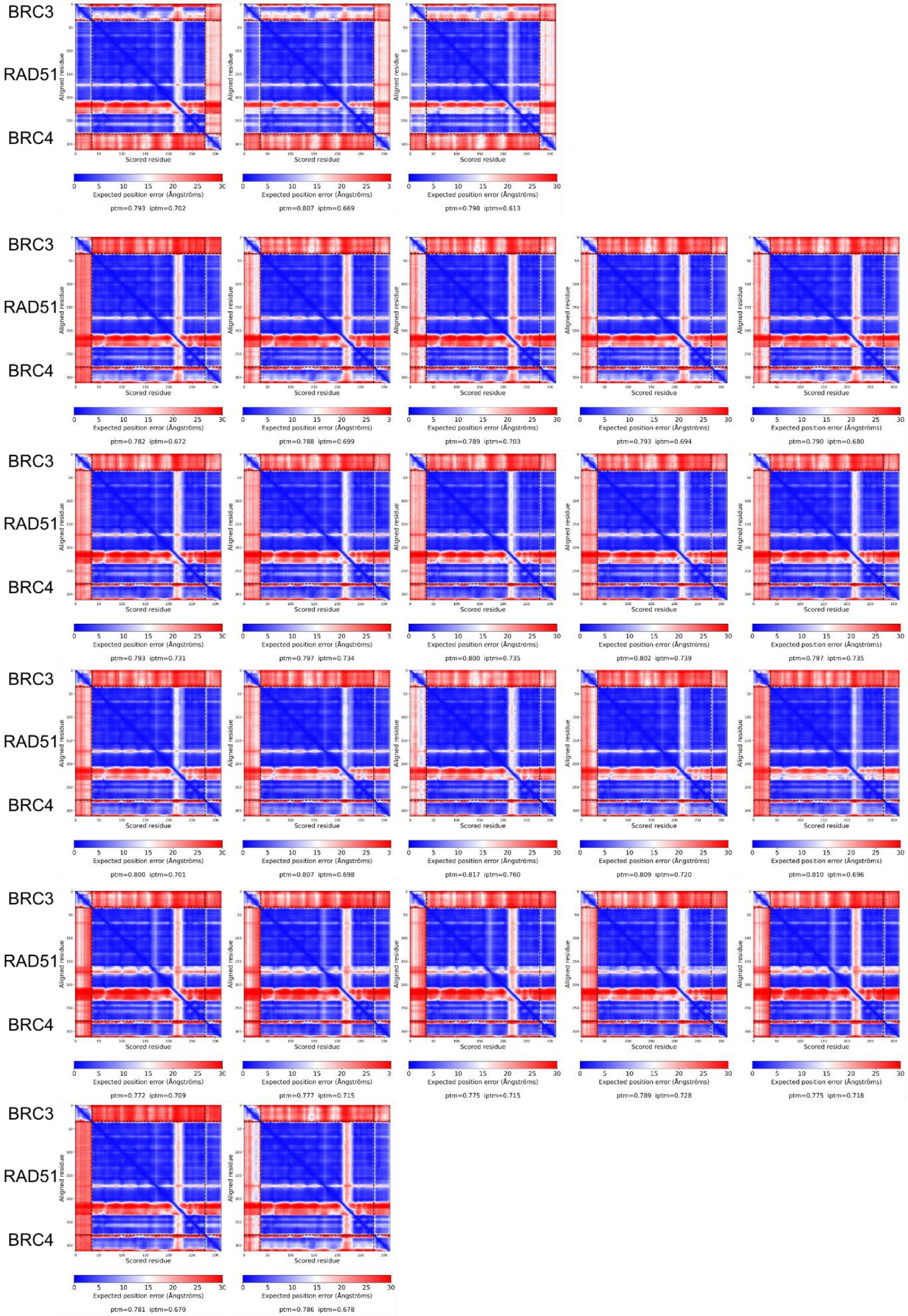


**Supplementary Figure 28 Predicted Align Error plots of predictions carried out with BRC3 and BRC4 sequences utilizing an N-terminal truncated RAD51 (Δ97-RAD51) sequence as receptor.** Predicted Align Error (PAE) plots provide the expected positional error at residue x if the predicted structure is aligned on residue y.

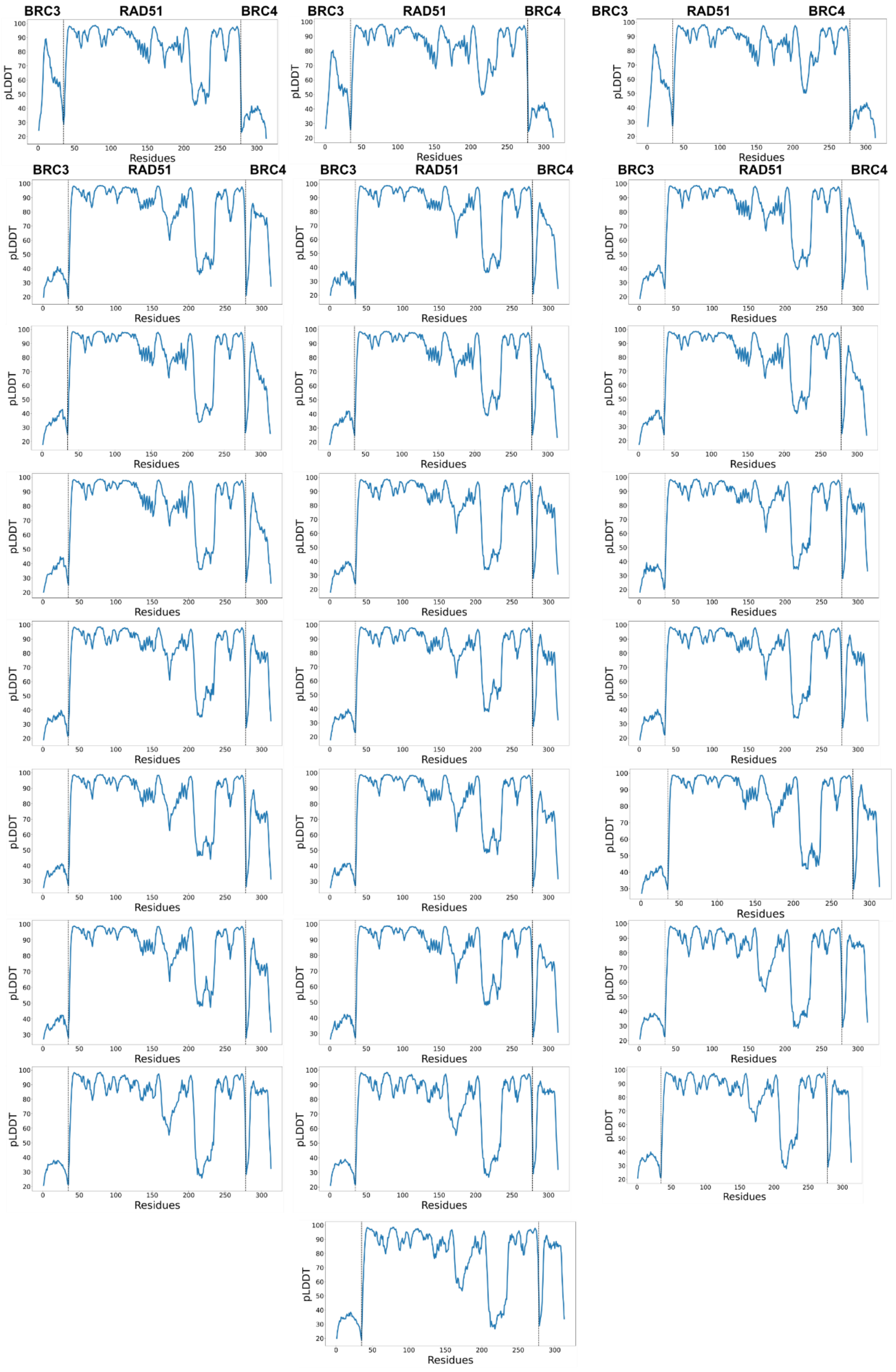


**Supplementary Figure 29 predicted Local Distance Difference Test (pLDDT) predictions carried out with BRC3 and BRC4 sequences utilizing a N-terminal truncated RAD51 (Δ97-RAD51) sequence as receptor.** pLDDT representation provides a per-residue accuracy metric.

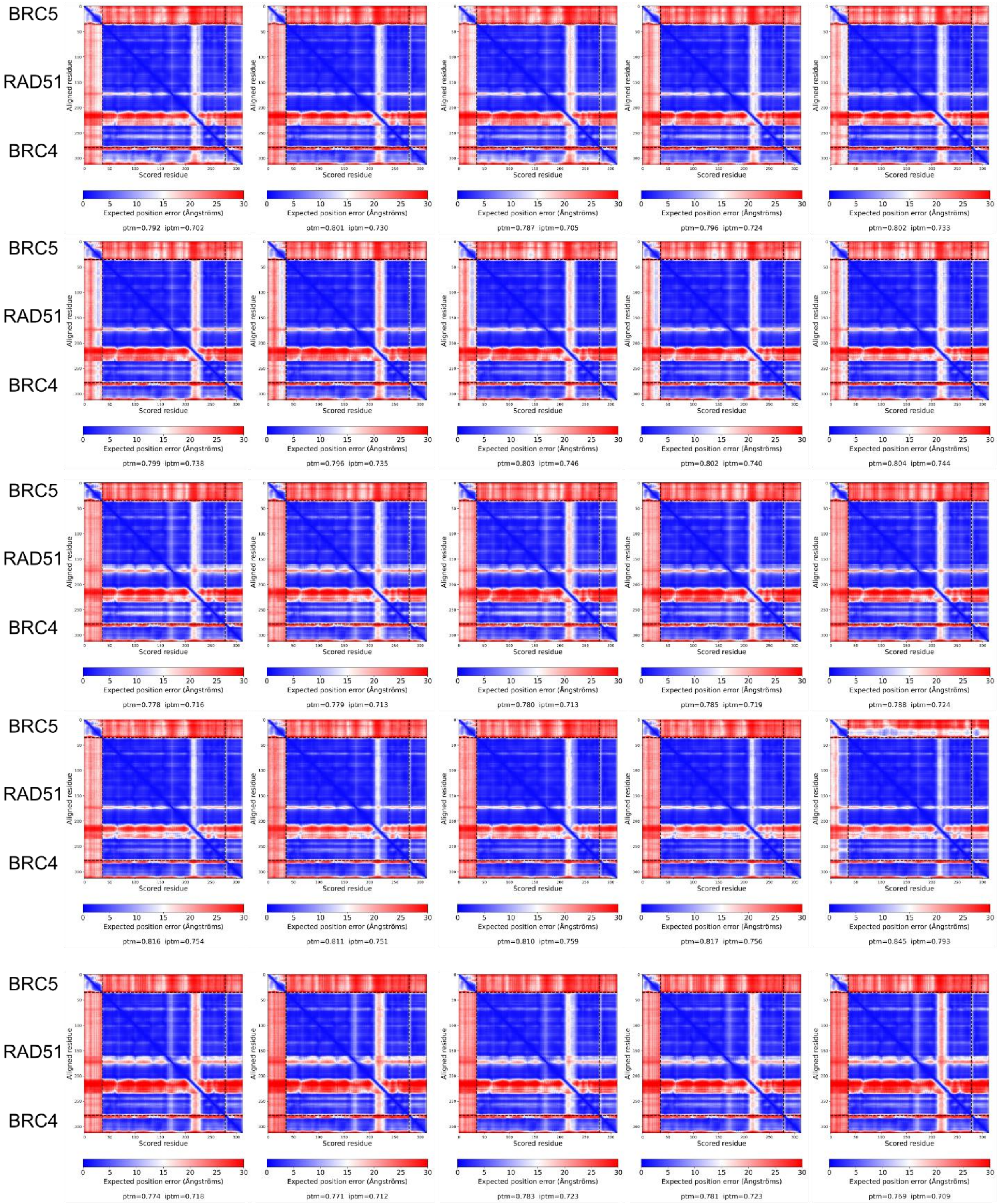


**Supplementary Figure 30 Predicted Align Error plots of predictions carried out with BRC5 and BRC4 sequences utilizing an N-terminal truncated RAD51 (Δ97-RAD51) sequence as receptor.** Predicted Align Error (PAE) plots provide the expected positional error at residue x if the predicted structure is aligned on residue y.

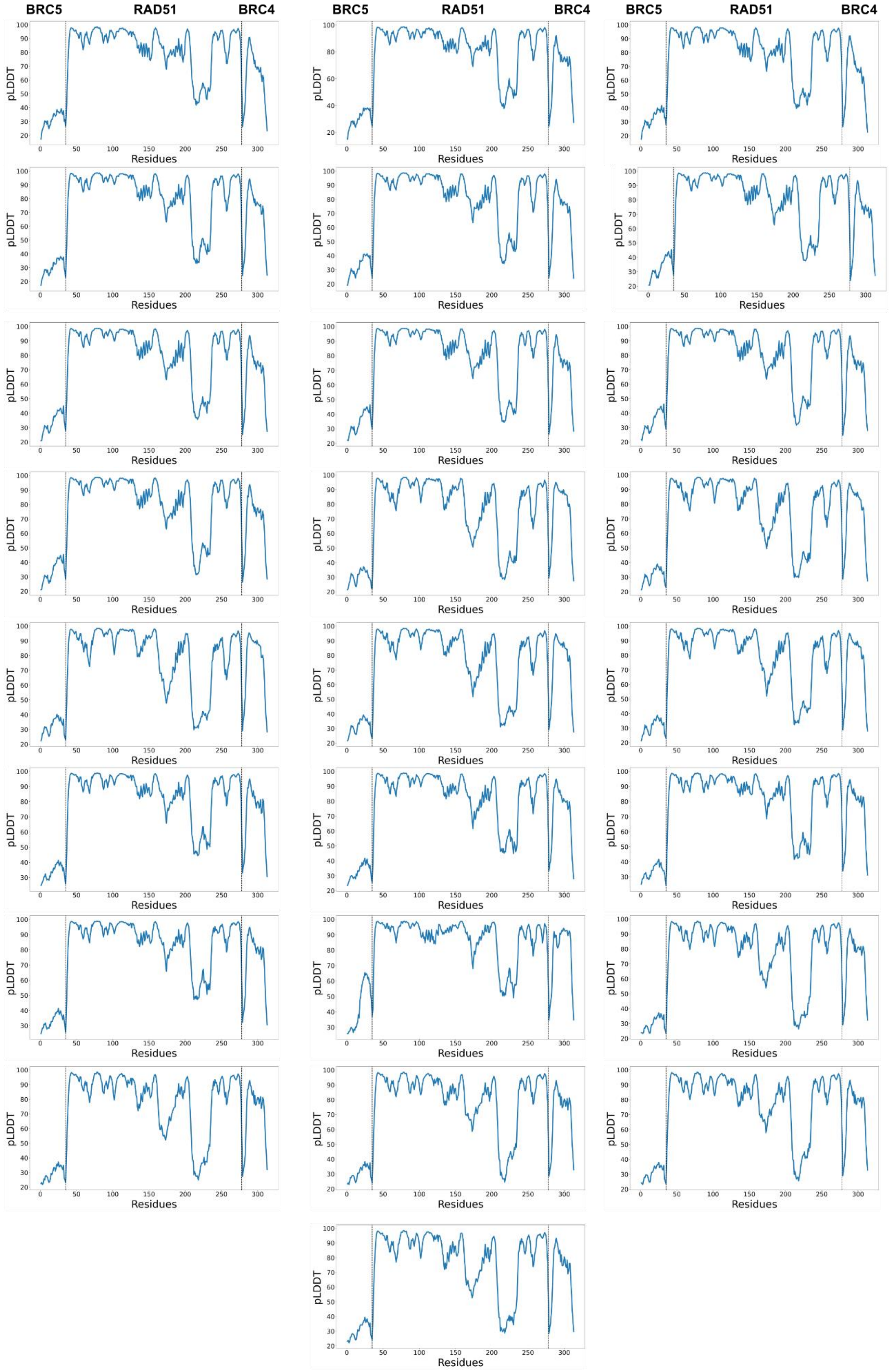


**Supplementary Figure 31 predicted Local Distance Difference Test (pLDDT) predictions carried out with BRC5 and BRC4 sequences utilizing a N-terminal truncated RAD51 (Δ97-RAD51) sequence as receptor.** pLDDT representation provides a per-residue accuracy metric.

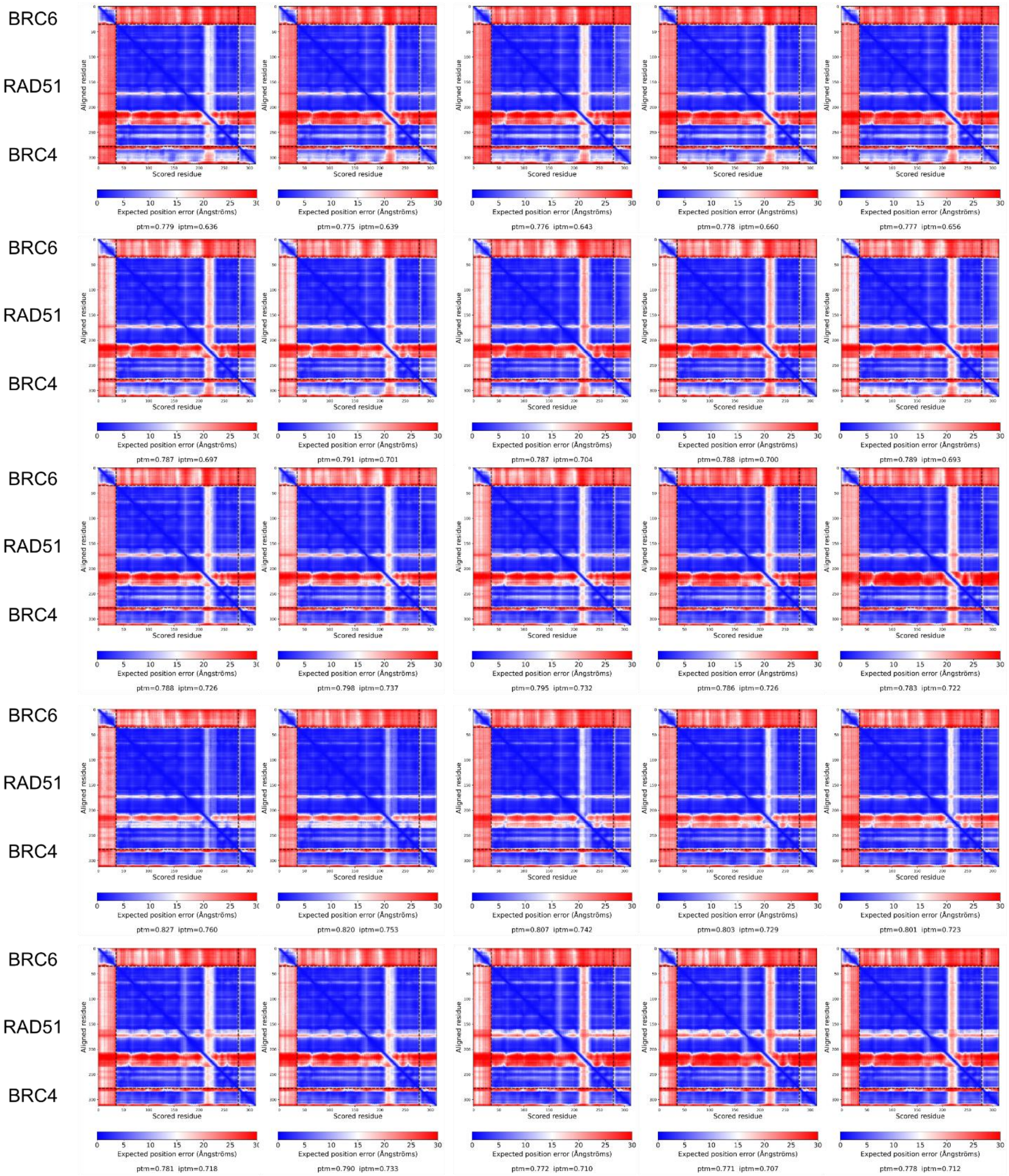


**Supplementary Figure 32 Predicted Align Error plots of predictions carried out with BRC6 and BRC4 sequences utilizing an N-terminal truncated RAD51 (Δ97-RAD51) sequence as receptor.** Predicted Align Error (PAE) plots provide the expected positional error at residue x if the predicted structure is aligned on residue y.

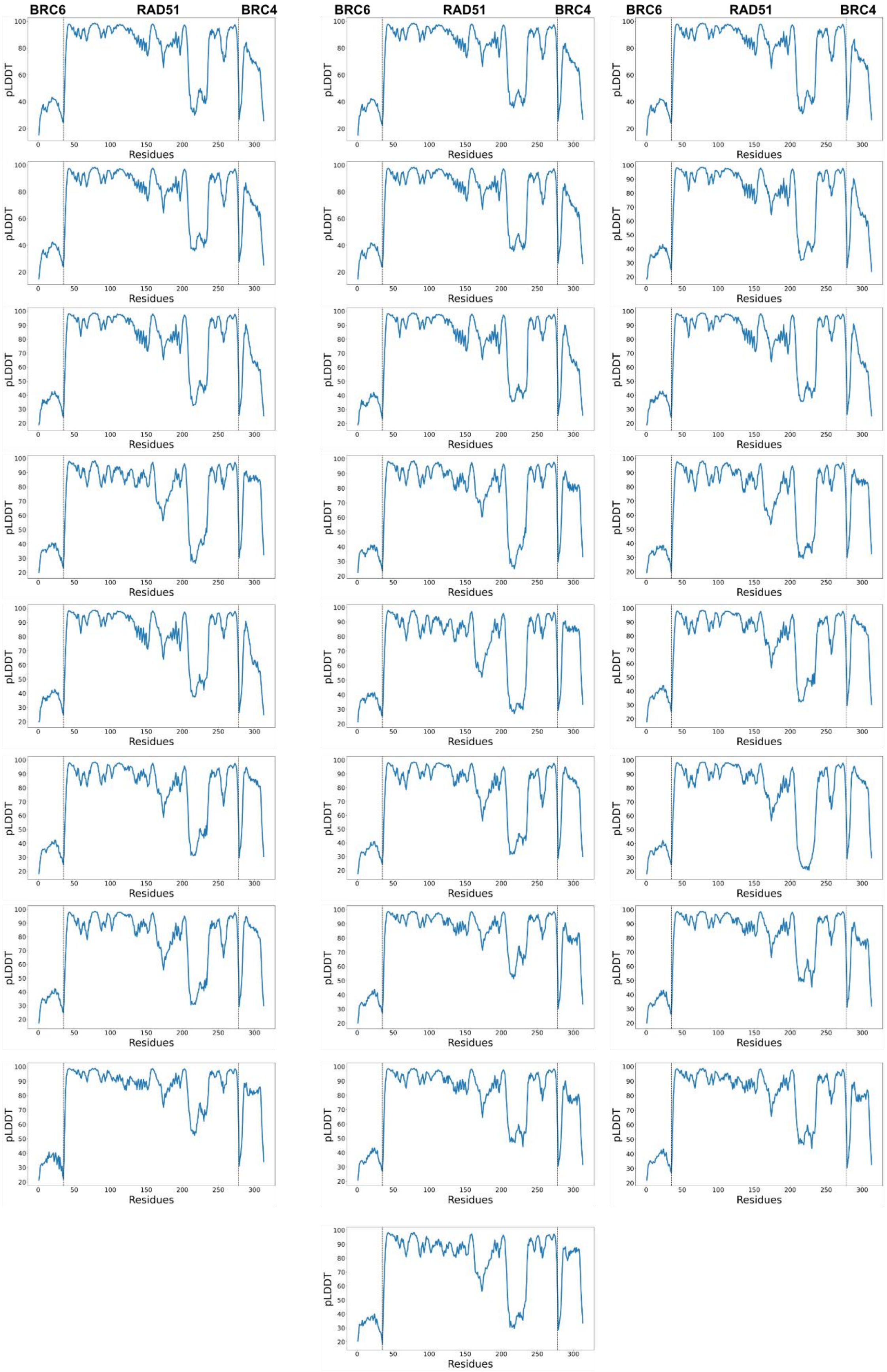


**Supplementary Figure 33 predicted Local Distance Difference Test (pLDDT) predictions carried out with BRC6 and BRC4 sequences utilizing a N-terminal truncated RAD51 (Δ97-RAD51) sequence as receptor.** pLDDT representation provides a per-residue accuracy metric.

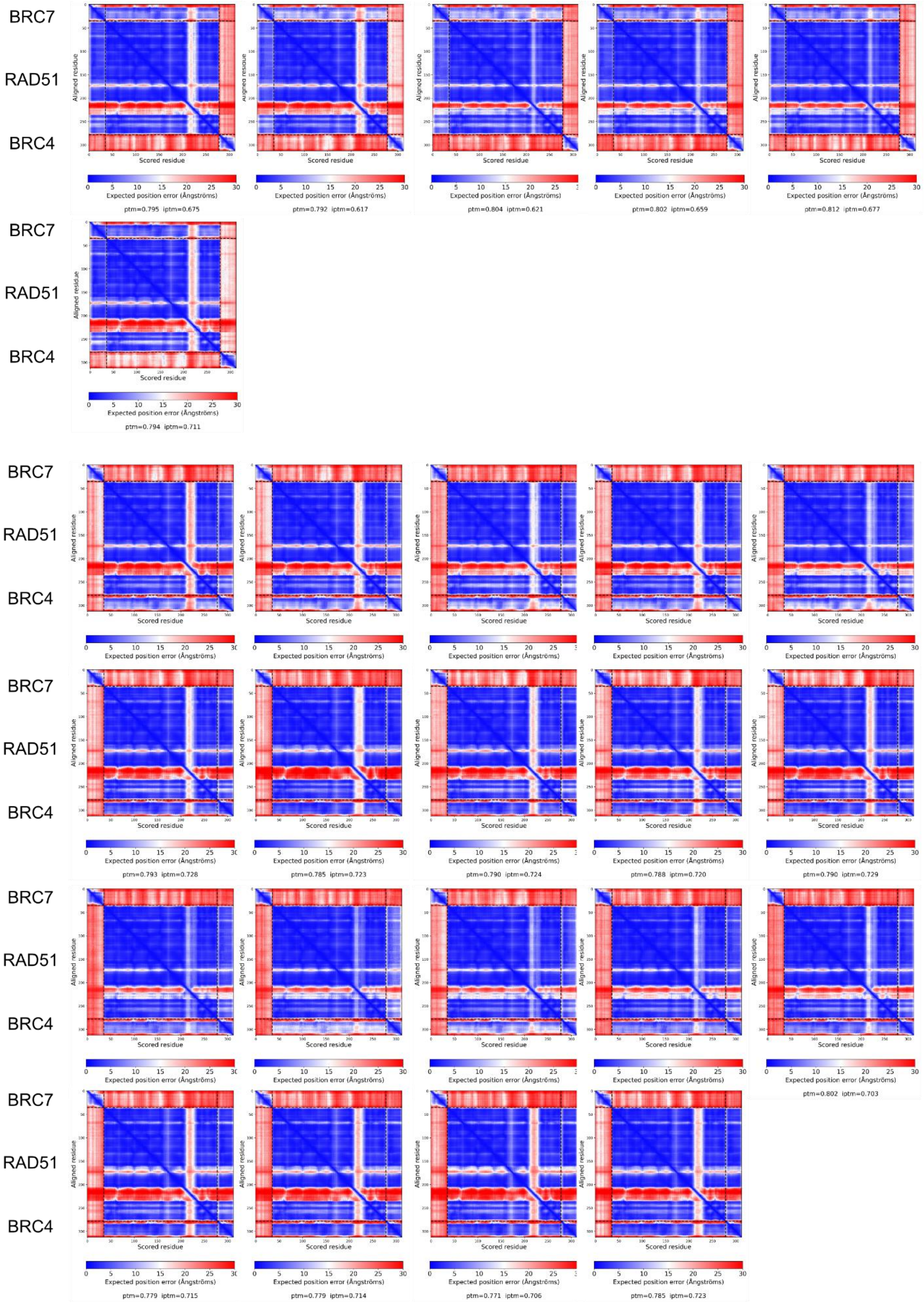


**Supplementary Figure 34 Predicted Align Error plots of predictions carried out with BRC7 and BRC4 sequences utilizing an N-terminal truncated RAD51 (Δ97-RAD51) sequence as receptor.** Predicted Align Error (PAE) plots provide the expected positional error at residue x if the predicted structure is aligned on residue y.

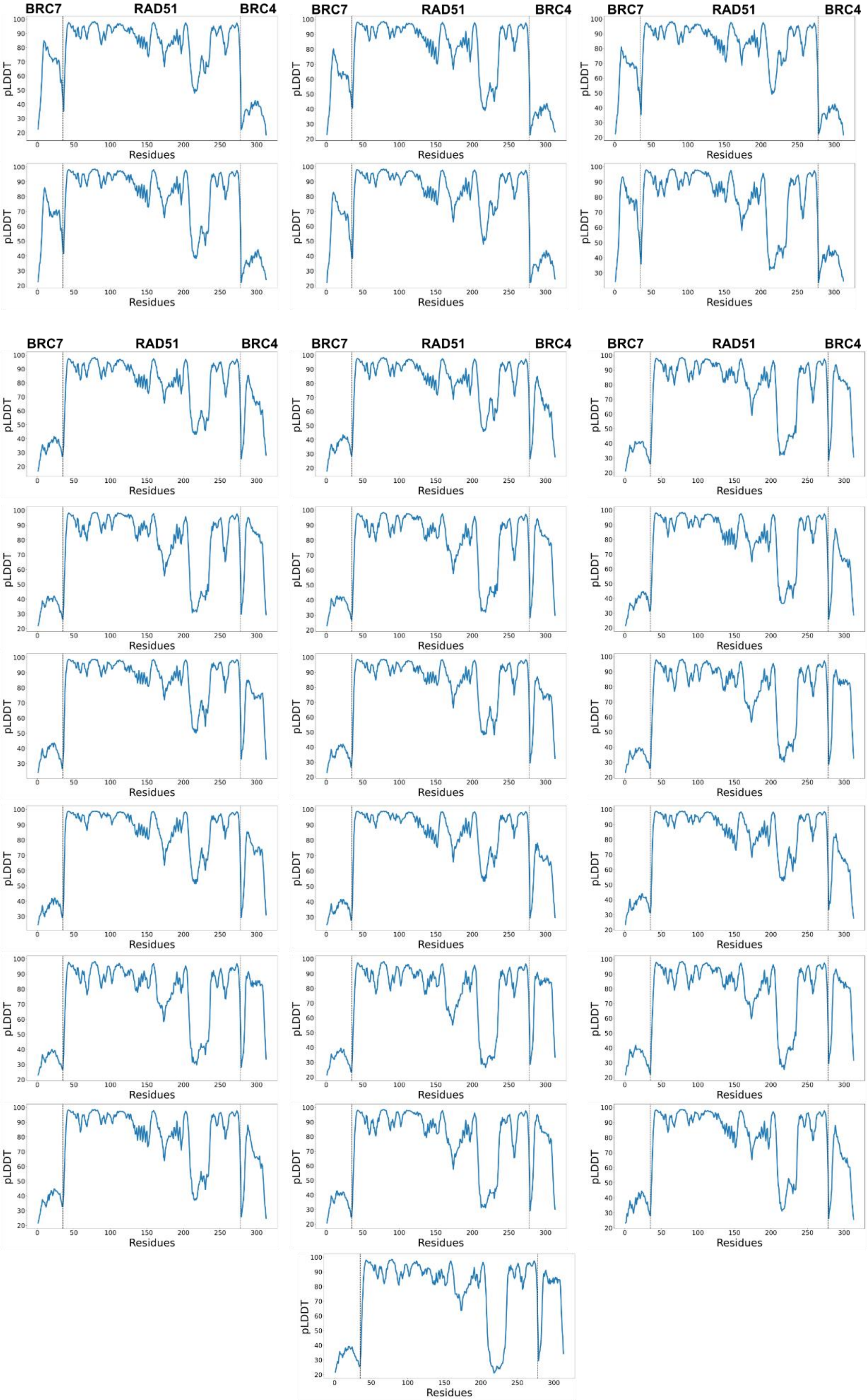


**Supplementary Figure 35 predicted Local Distance Difference Test (pLDDT) predictions carried out with BRC7 and BRC4 sequences utilizing a N-terminal truncated RAD51 (Δ97-RAD51) sequence as receptor.** pLDDT representation provides a per-residue accuracy metric.

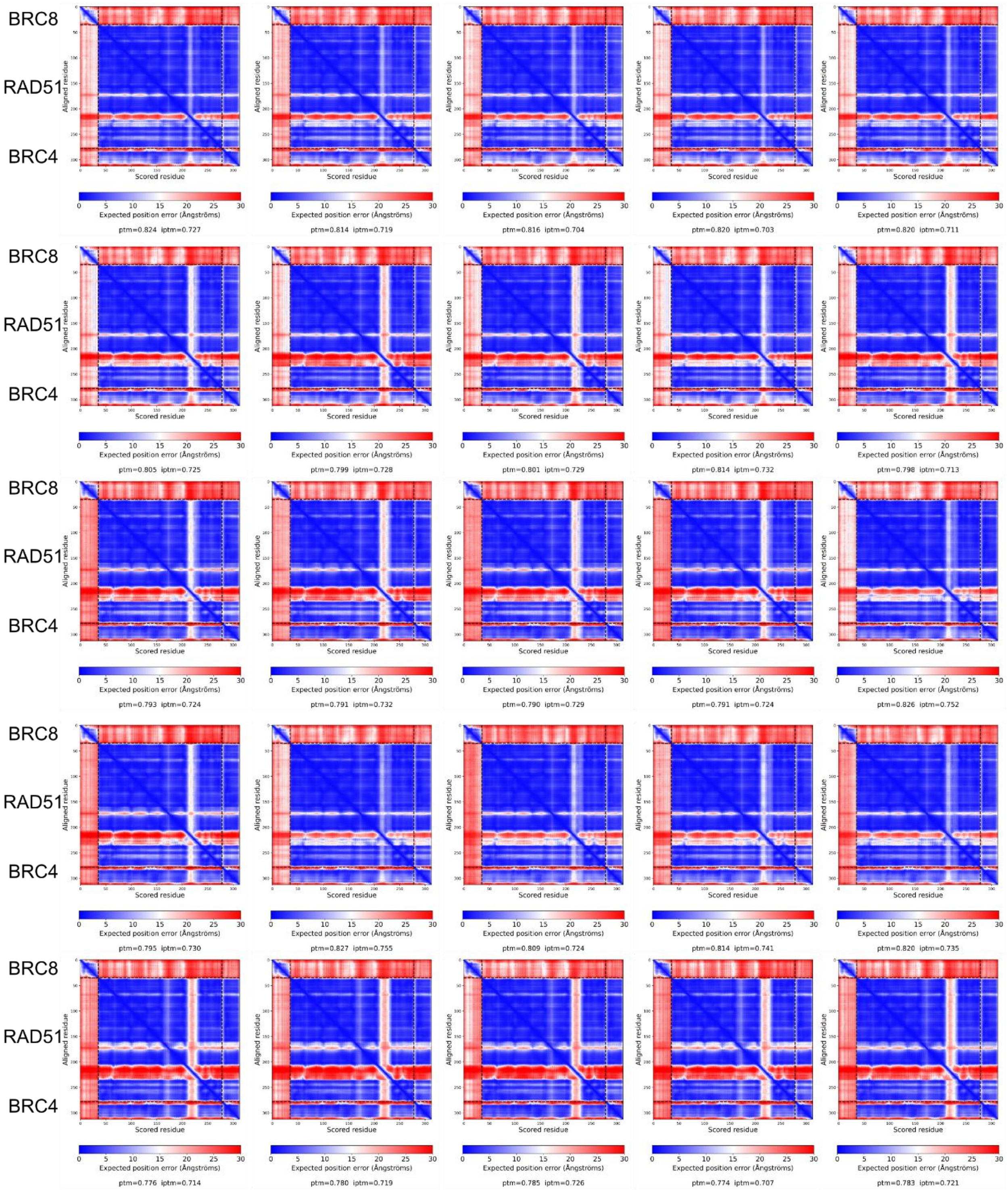


**Supplementary Figure 36 Predicted Align Error plots of predictions carried out with BRC8 and BRC4 sequences utilizing an N-terminal truncated RAD51 (Δ97-RAD51) sequence as receptor.** Predicted Align Error (PAE) plots provide the expected positional error at residue x if the predicted structure is aligned on residue y.

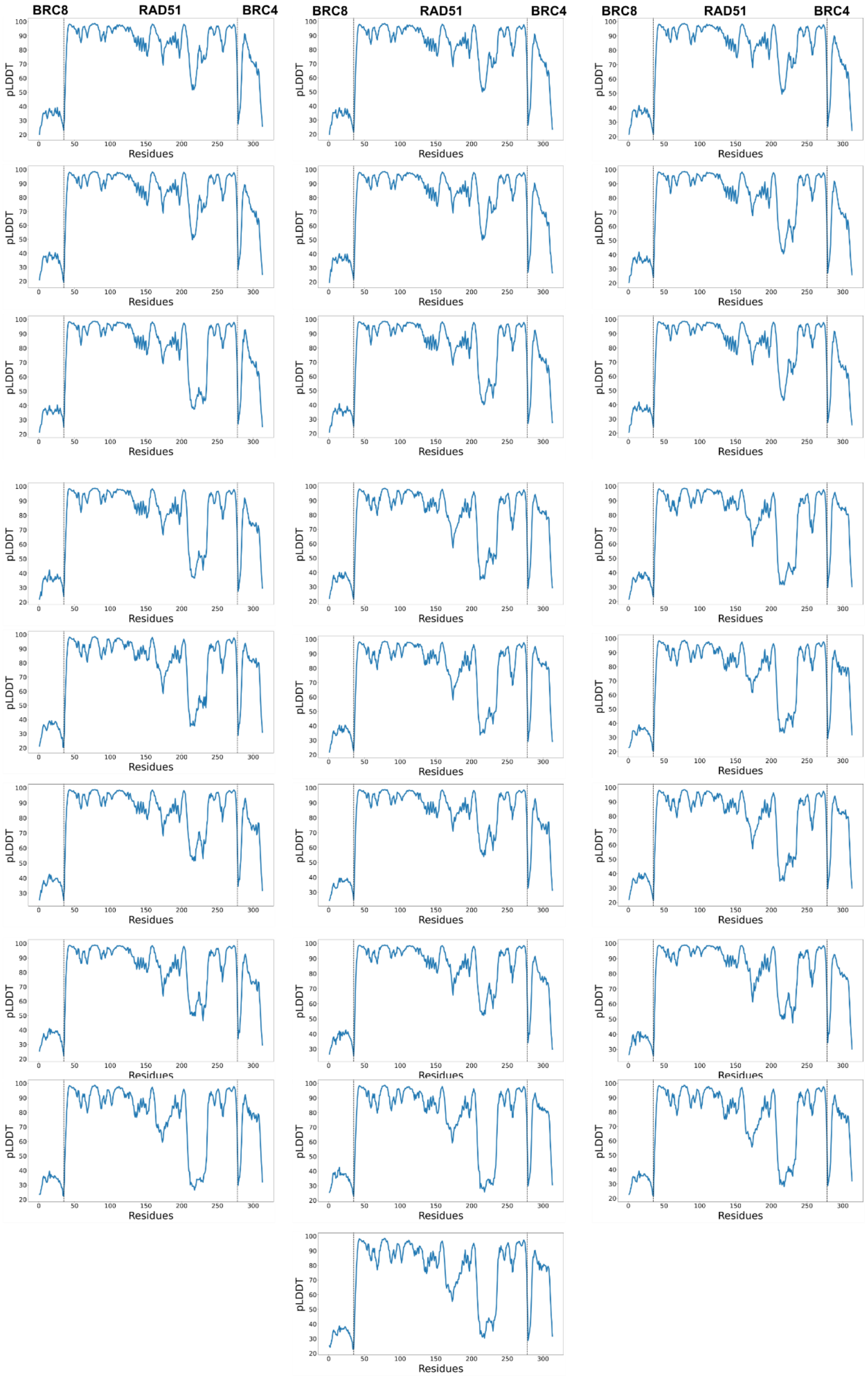


**Supplementary Figure 37 predicted Local Distance Difference Test (pLDDT) predictions carried out with BRC8 and BRC4 sequences utilizing a N-terminal truncated RAD51 (Δ97-RAD51) sequence as receptor.** pLDDT representation provides a per-residue accuracy metric.

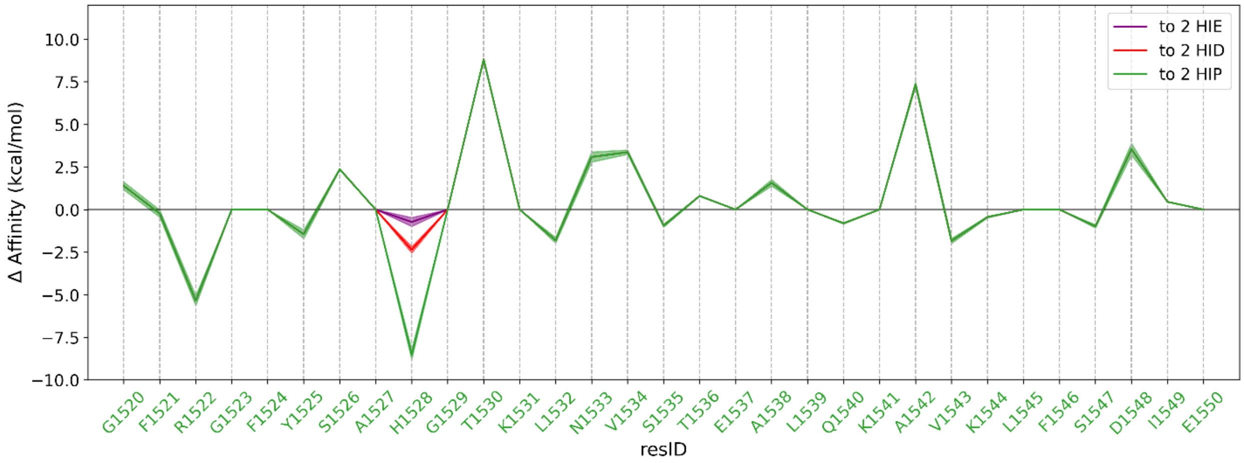


**Supplementary Figure 38** Residue scanning analysis for mutating the BRC4 into the BRC2 repeat. The results considering the possible tautomeric (epsilon and delta, indicated as HIE and HID, respectively) and protonated (HIP) forms for histidine 1528 are reported.

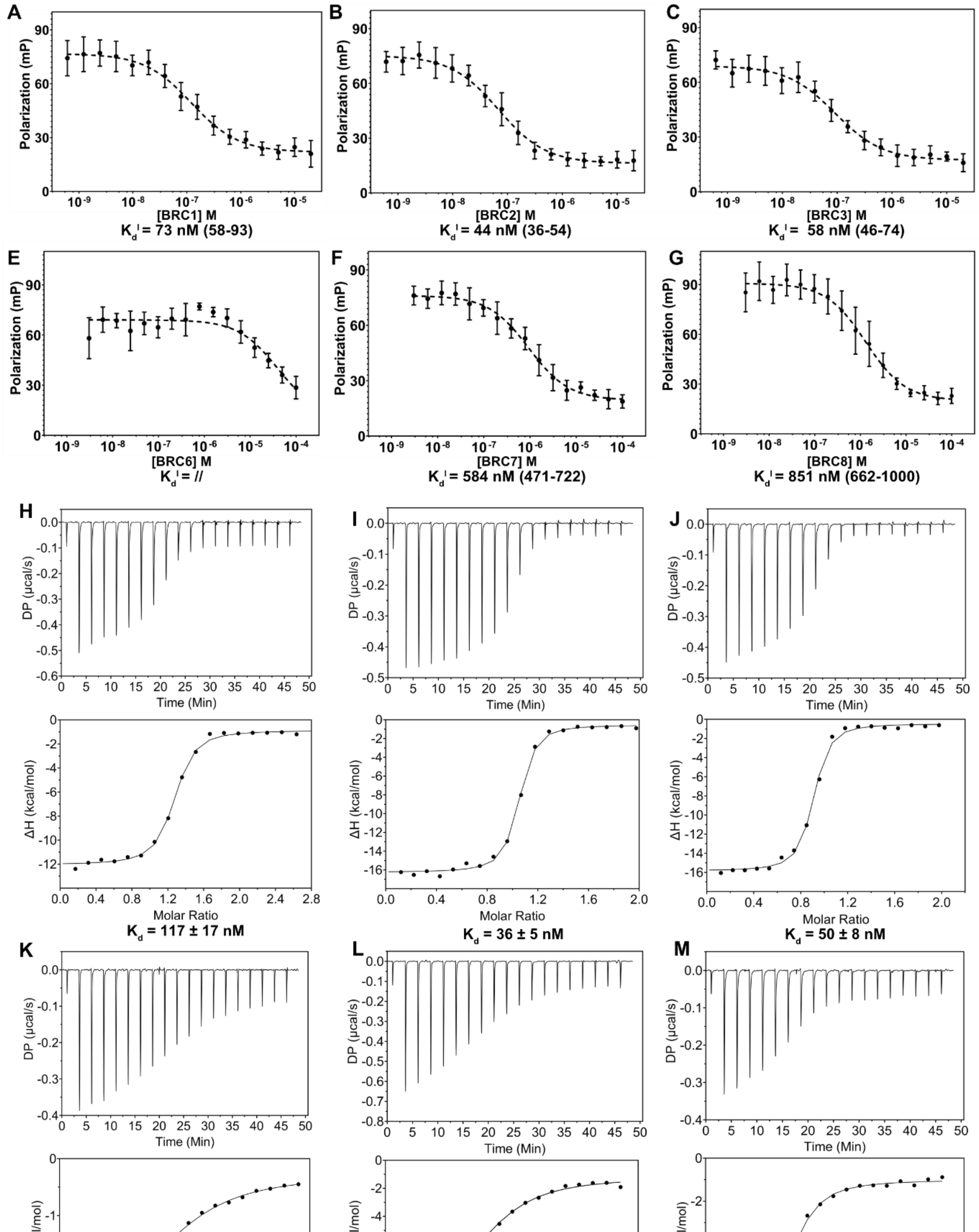


**Supplementary Figure 39 Biophysical analysis of the BRC repeats RAD51 interaction through monomeric RAD51** Competitive FP assay in which a pre-formed labelled BRC4 (BRC4*)-monomeric RAD51 complex is titrated with increasing concentrations of unlabeled **A.** BRC1, **B.** BRC2, **C.** BRC3, **D.** BRC6, **E.** BRC7, **F.** BRC8 (n=3). ITC Analyses of the RAD51 monomer interaction with **G.** BRC1, **H.** BRC2, **I.** BRC3, **L.** BRC6, **M.** BRC7, **N.** BRC8.

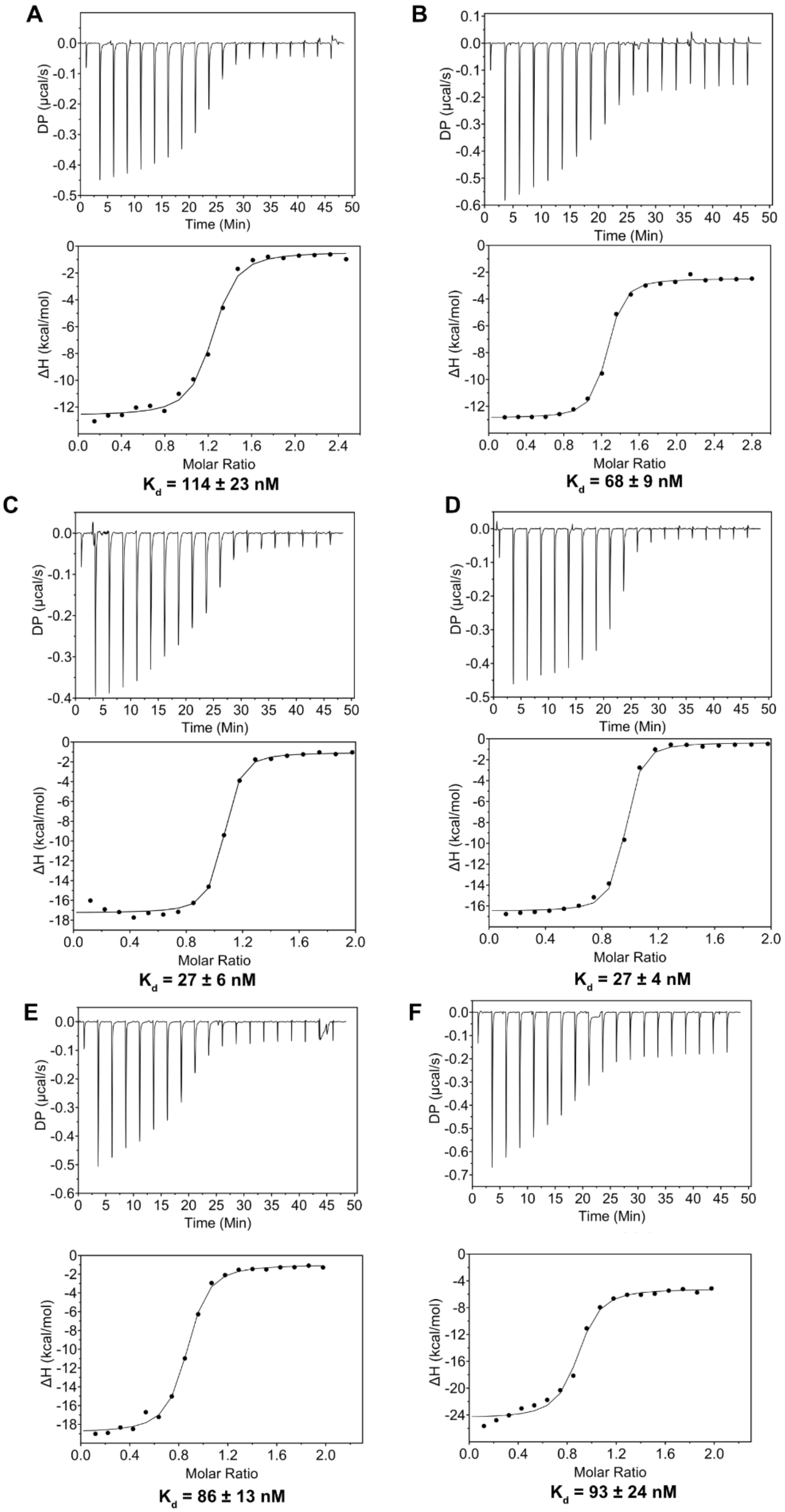


**Supplementary Figure 40** ITC Analysis of **A., B.** BRC1 – His-RAD51 [F86E, A89E] interaction. **C., D.** BRC2 - His-RAD51 [F86E, A89E] interaction **E., F.** BRC3 - His-RAD51 [F86E, A89E] interaction. The reaction are exothermic.

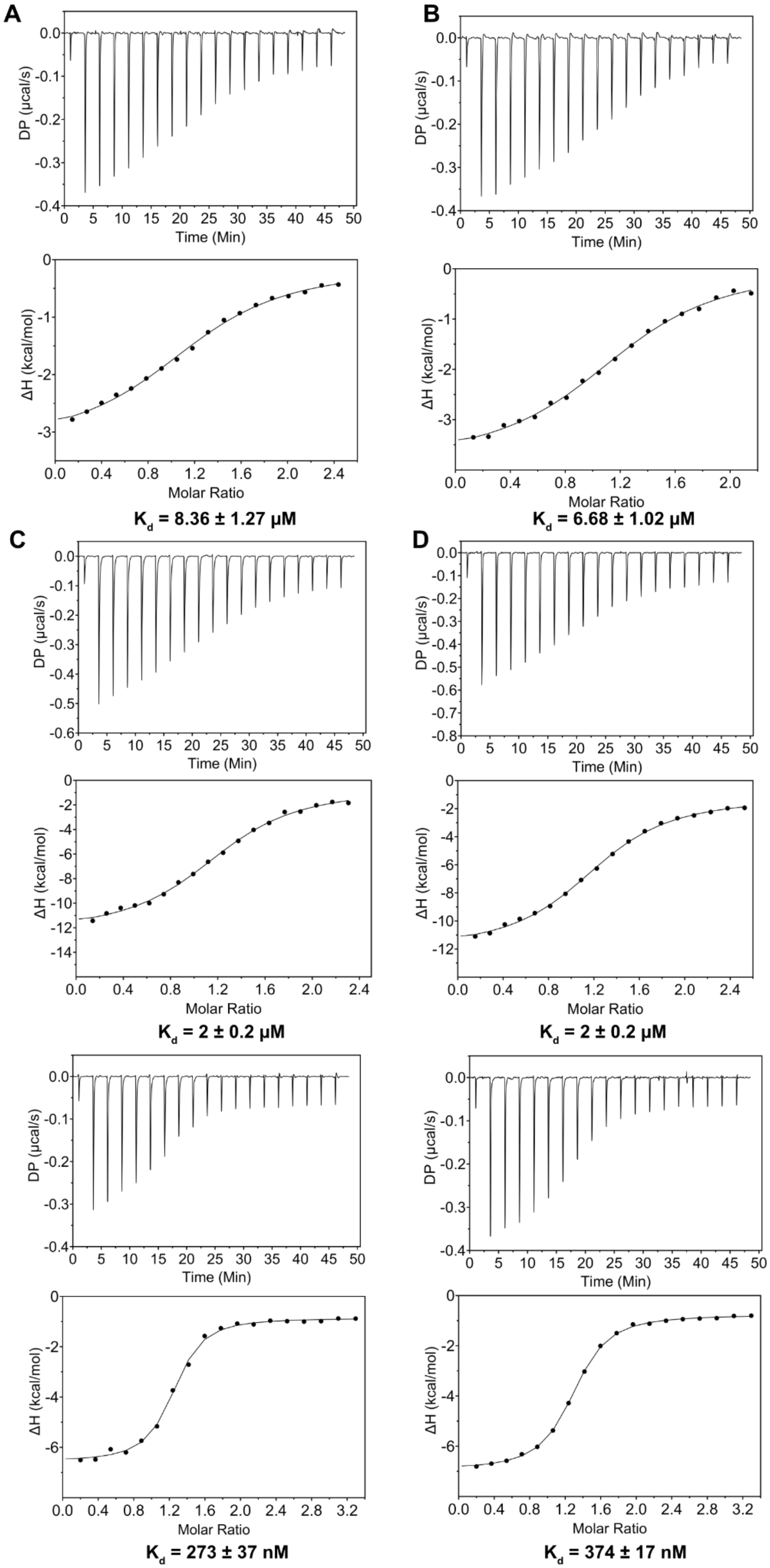


**Supplementary Figure 41** ITC Analysis of **A., B.** BRC6 – His-RAD51 [F86E, A89E] interaction. **C., D.** BRC7 - His-RAD51 [F86E, A89E] interaction **E., F.** BRC8 - His-hRAD51 [F86E, A89E] interaction. The reaction are exothermic.

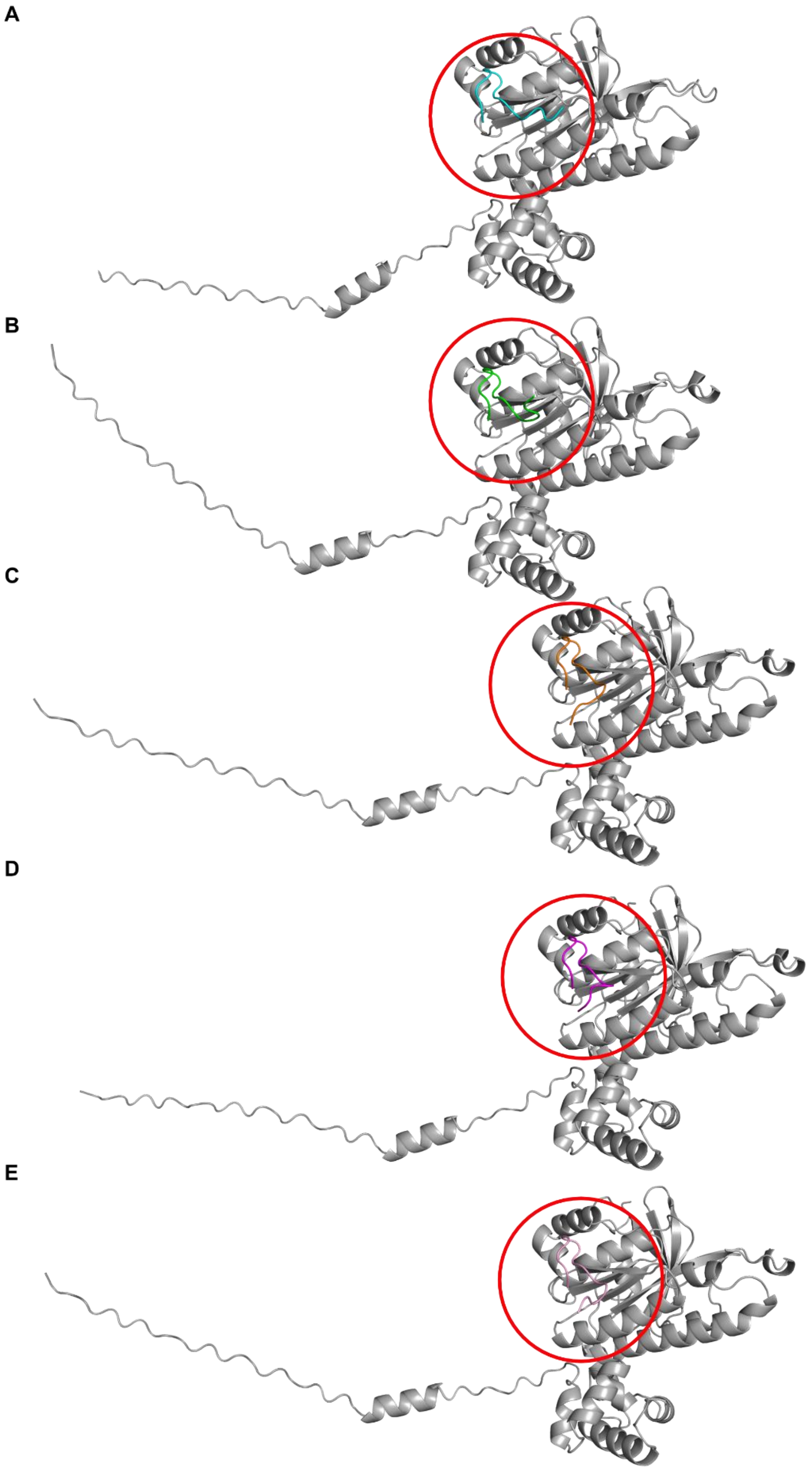


**Supplementary Figure 42 AlphaFold prediction of the His-RAD51[F86E, A89E] in complex with FXXA domains of BRC-repeats utilized for NMR competition experiments.** Peptides have been presented with different colors: **A. BRC1** (cyan) **B. BRC2** (green) **C. BRC3** (orange) **D. BRC7** (magenta) **E. BRC8** (pink).

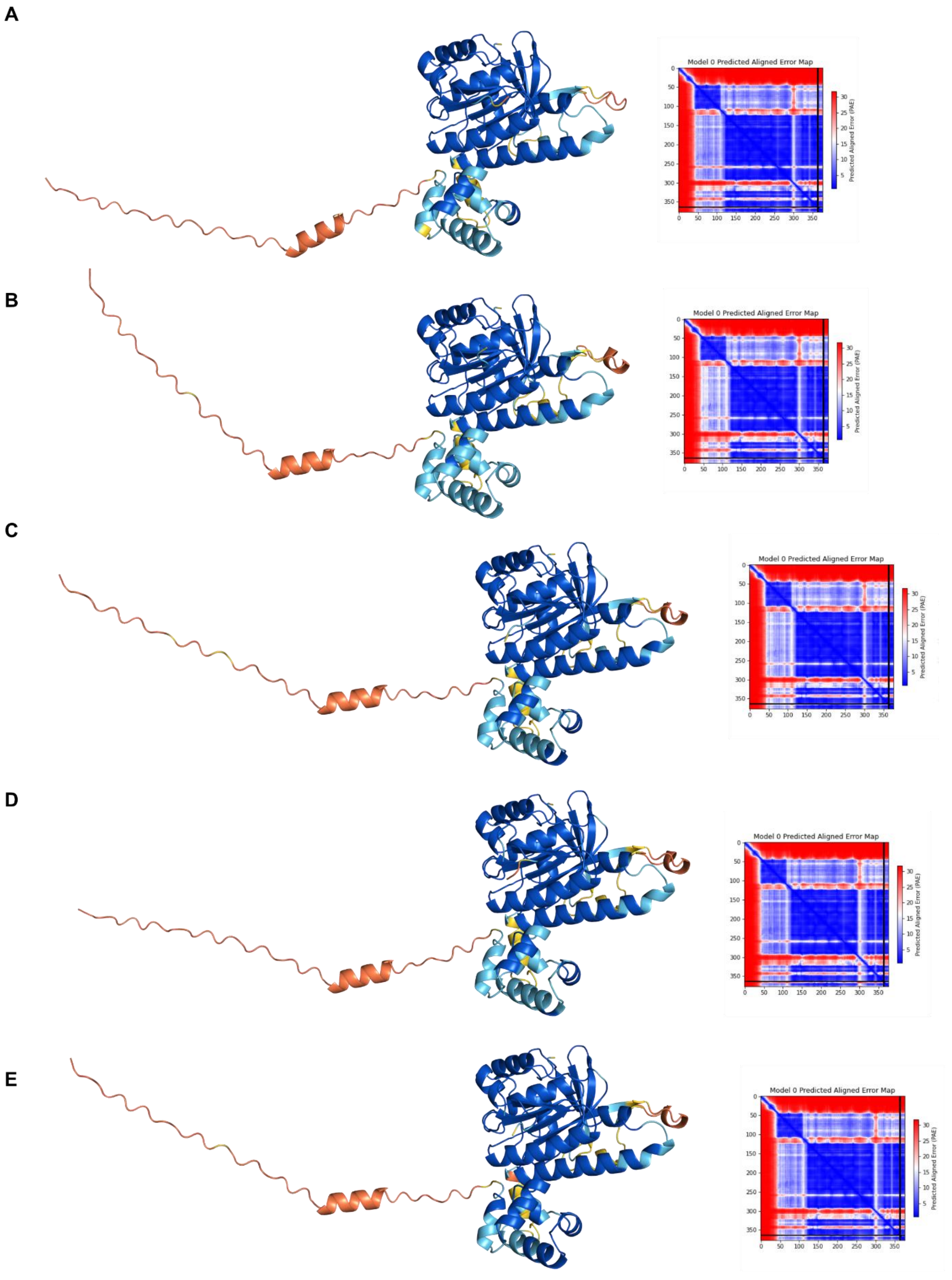


**Supplementary Figure 43 AlphaFold prediction of the His-RAD51[F86E, A89E] in complex with FXXA domains of BRC-repeats utilized for NMR competition experiments, colored by pLDDT. A. BRC1 B. BRC2 C. BRC3 D. BRC7 E. BRC8**. On the right of each model the PAE matrices are reported.

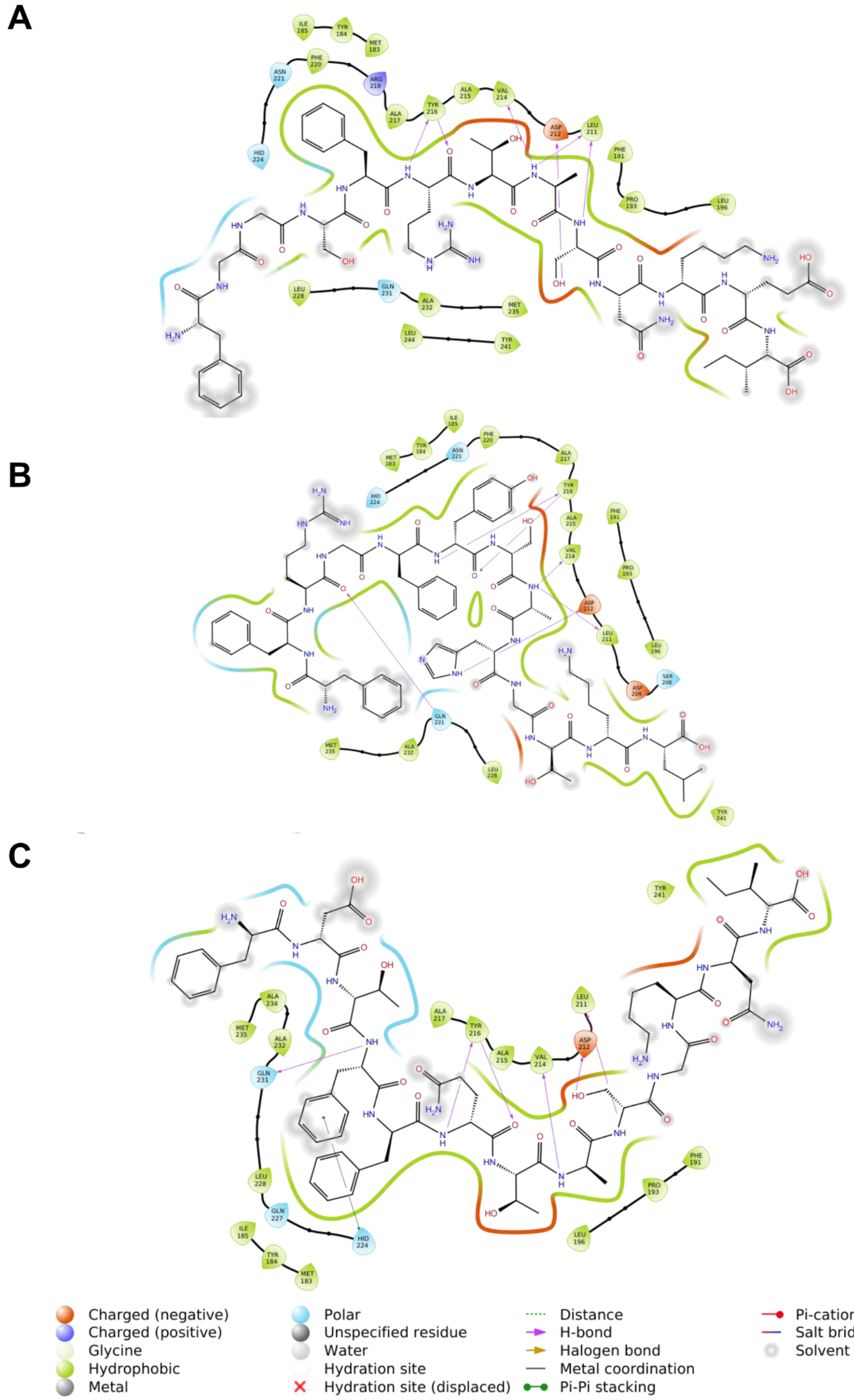


**Supplementary Figure 44 Ligand interaction diagrams of peptides' FXXA domains with RAD51 predicted by AlphaFold. A.** BRC1, **B.** BRC2, **C.** BRC3. On the bottom of the figure the legend provides information concerning the charge of residues, and type of interactions.

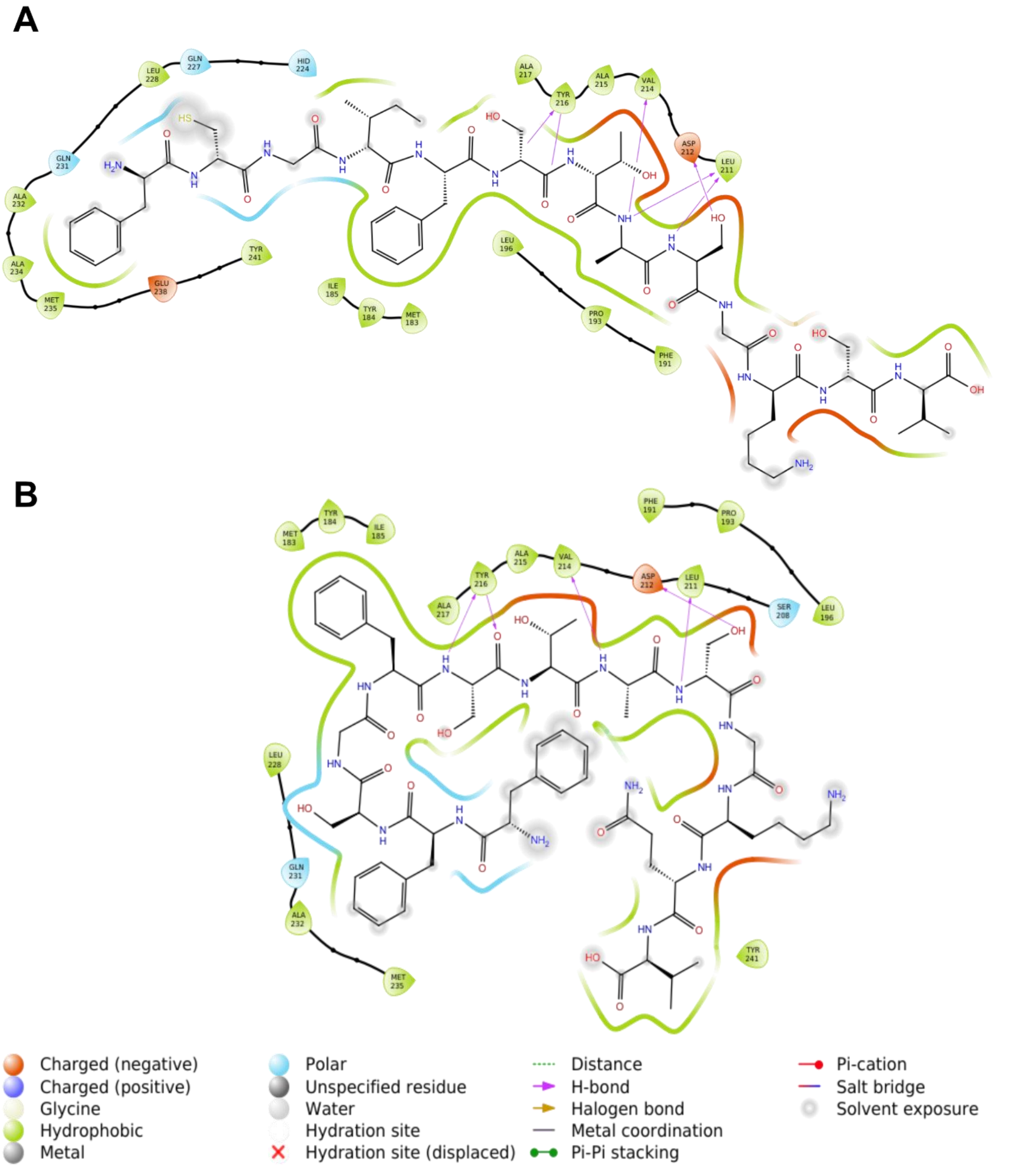


**Supplementary Figure 45 Ligand interaction diagrams of peptides' FXXA domains with RAD51 predicted by AlphaFold. A.** BRC7, **B.** BRC8. On the bottom of the figure the legend provides information concerning the charge of residues, and type of interactions.

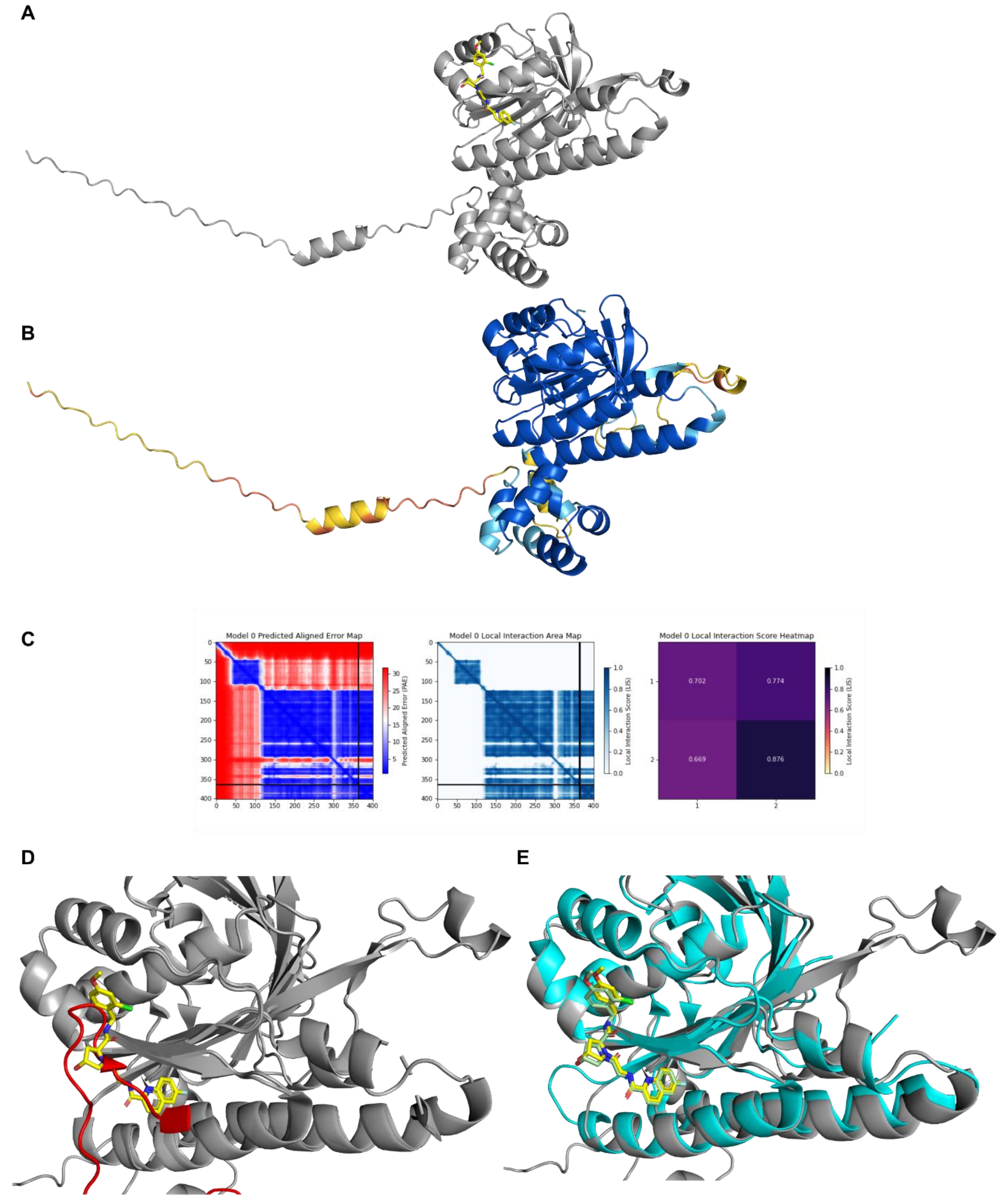


**Supplementary Figure 46 RAD51-CAM833 interaction A.** AlphaFold prediction of the His-RAD51[F86E, A89E] (grey) in complex with CAM833 (yellow). **B.** AlphaFold prediction of the His-RAD51[F86E, A89E] in complex with CAM833 colored by pLDDT. **C.** PAE matrix for the generated complex and Ligand Interaction Score (LIS) which provides an estimation of the interaction strength and quality within protein complexes. **D.** Overlay of the generated AlphaFold model with the available PDB entry of the RAD51-BRC4 complex (PDB entry: 1n0w), highlighting that CAM833 and the FXXA binding motif overlaps **E.** Overlay of the generated AlphaFold model with the available PDB entry of HumRadA22F in complex with CAM833 (PDB entry: 6TW9)

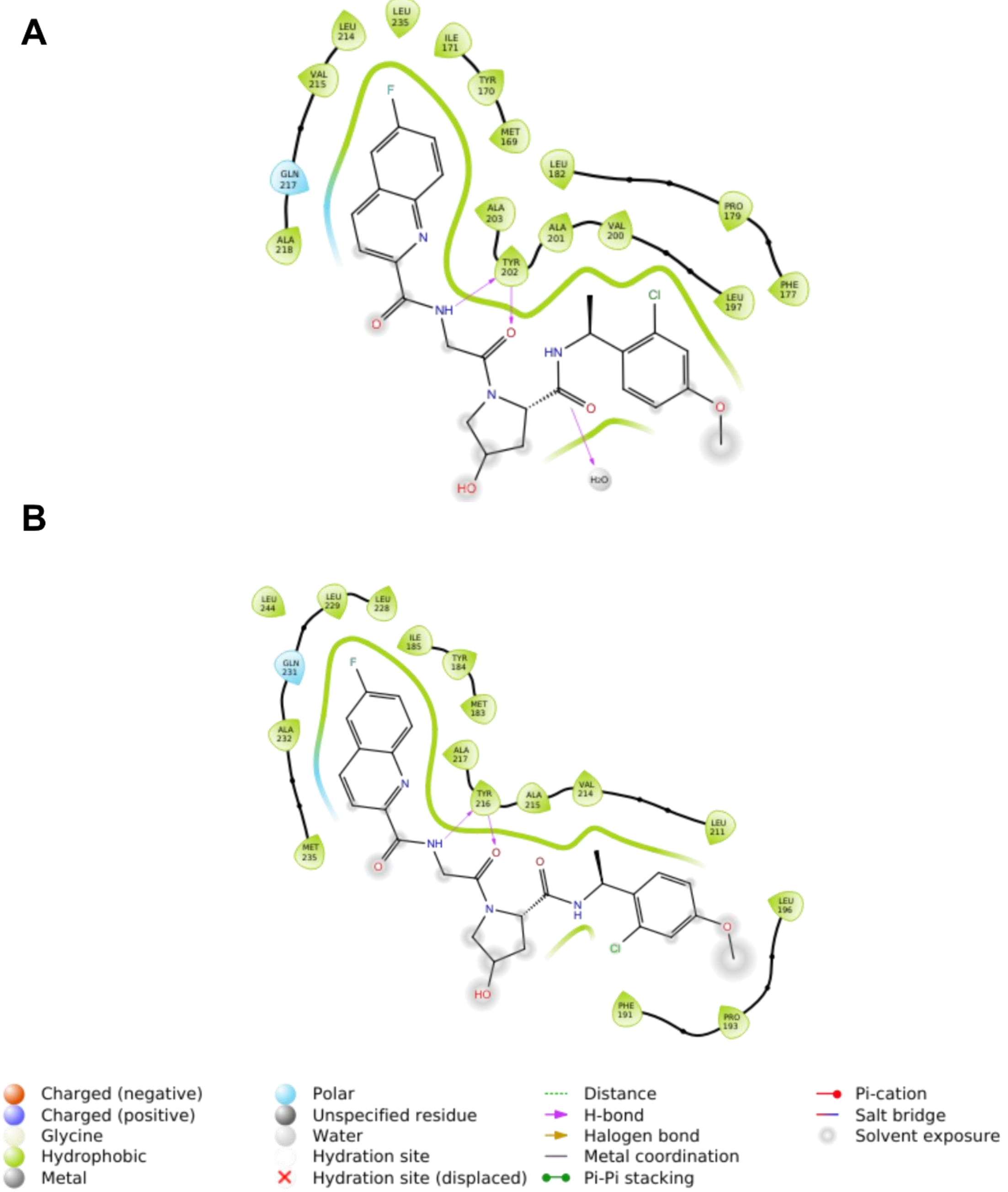


**Supplementary Figure 47 Ligand interaction diagrams of CAM833A interaction with A.** HumRadA22F (PDB entry: 6tw9) **B.** human RAD51 predicted by AlphaFold

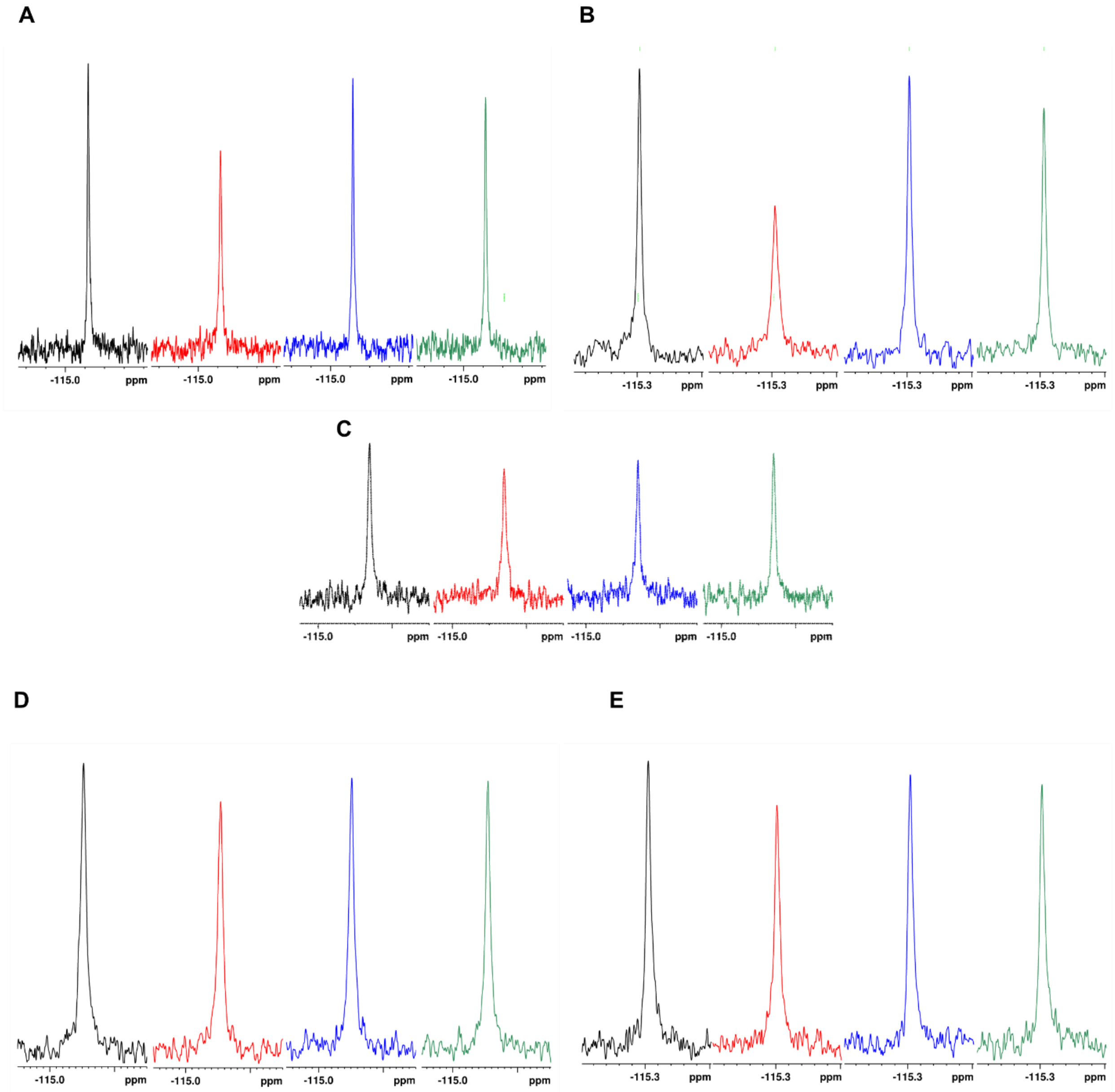


**Supplementary Figure 48 Characterization of the fluorinated FXXA BRC-repeats peptides' interaction with RAD51 [F86E, A89E] Monomer by competitive NMR experiments.** $^{19}F$ $T_2$ filter experiments of $^{19}F$-peptide in the absence (black) and presence (red) of His-RAD51 [F86E, A89E] monomer in presence of 1.92 µM BRC4 (blue) or 10 µM CAM833A (green). In all experiments we can observe the line broadening of the peptides' $^{19}F$ NMR signal in the presence of protein, due to its binding to RAD51 monomer. Additionally, in the presence of competitors we can observe the sharpening of the peptide $^{19}F$ signal due to its displacement from the monomer by BRC4 or CAM833A, thus suggesting direct competition for the same binding site.

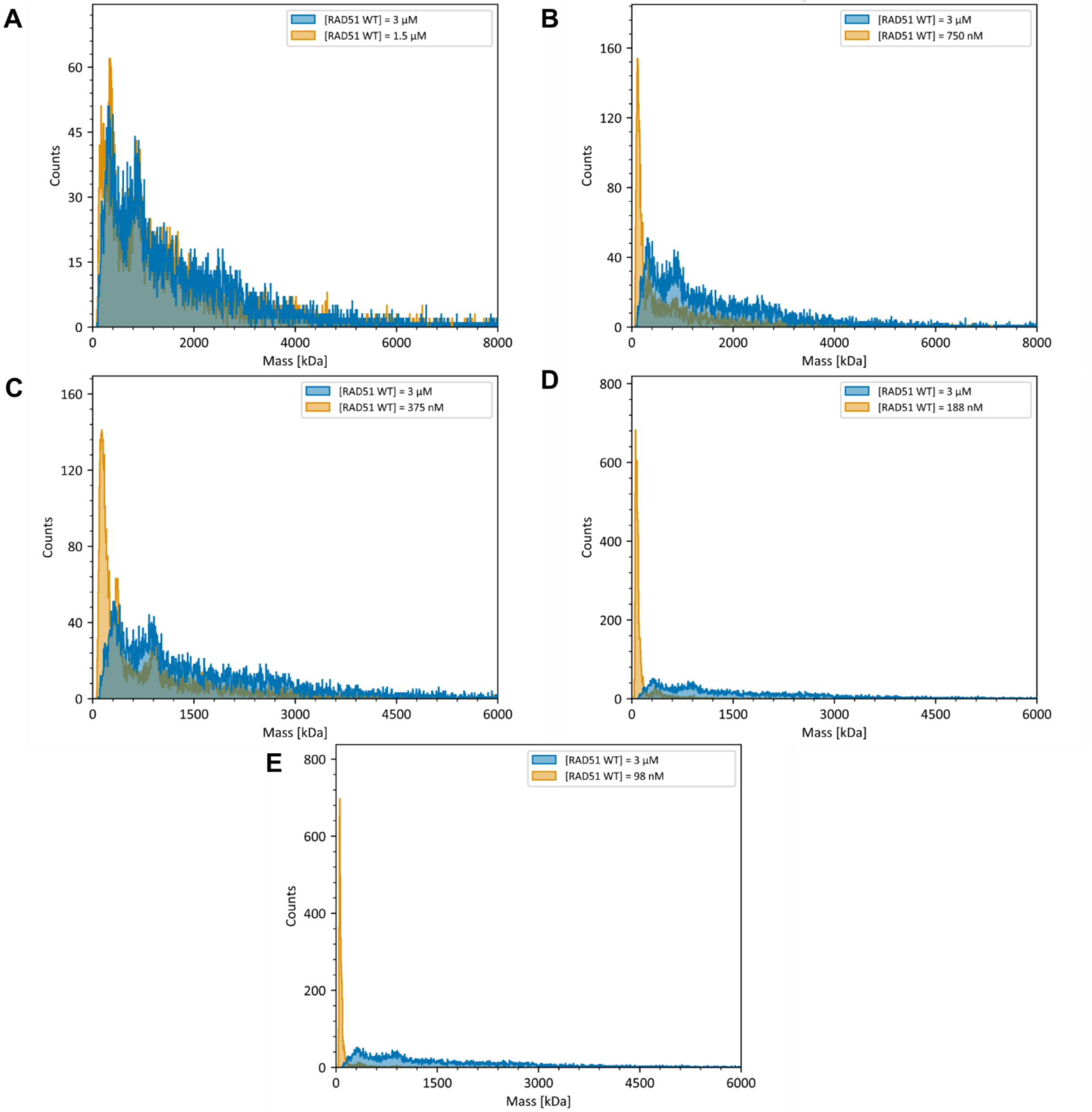


**Supplementary Figure 49 Mass Photometry analysis of the RAD51 WT mass distribution variation at different concentrations A.** 1.5 µM, **B.** 0.75 µM, **C.** 0.375 µM, **D** 0.188 µM, **E** 0.98 µM. As a comparison, in all graphs it is reported the mass distribution at 3 µM concentration (blue).

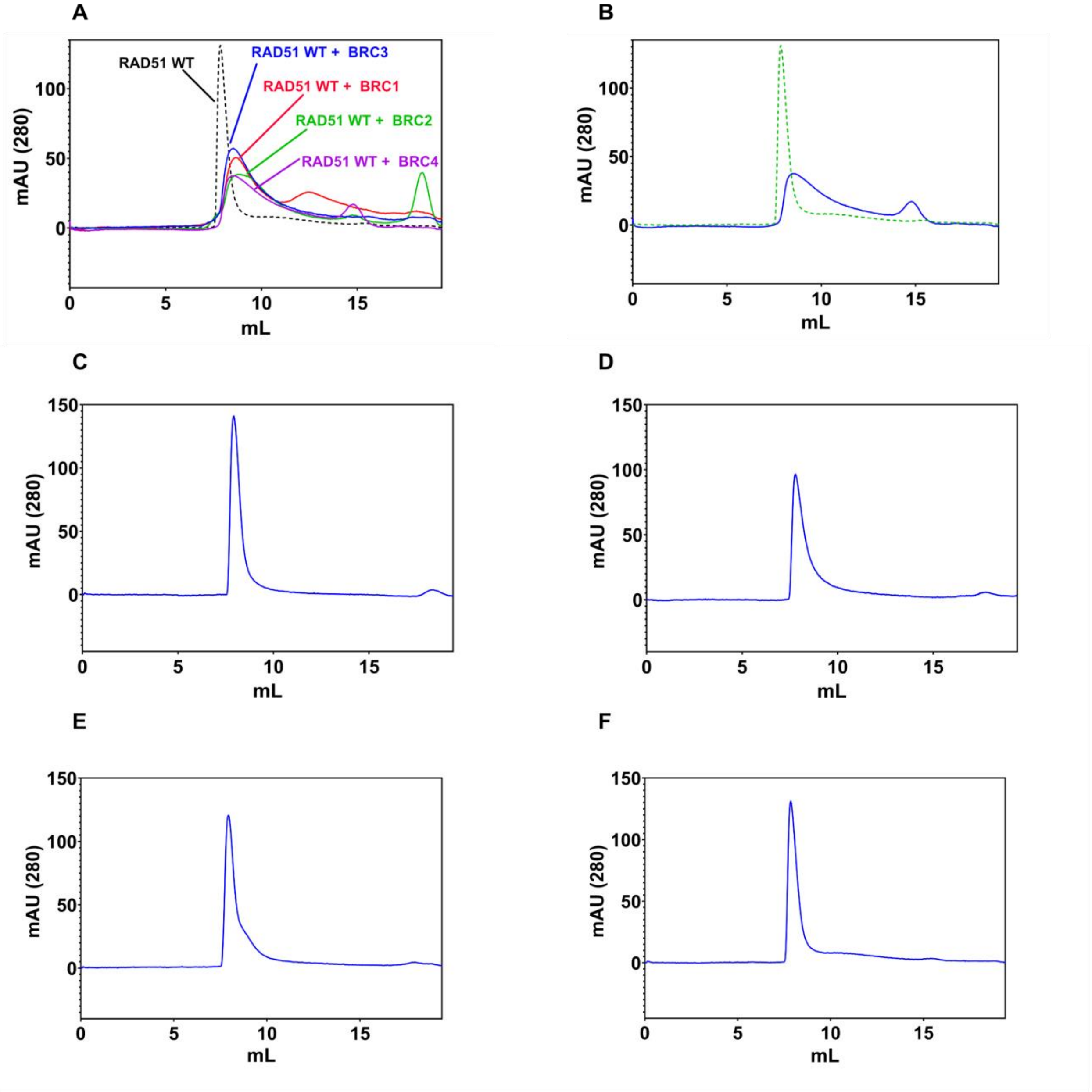


**Supplementary Figure 50 A. Elution profiles of RAD51 WT (25 µM) in presence of a four-fold higher stoichiometric excess of BRC-repeat peptides (100 µM) loaded onto a Superdex 200 Increase 10/300 GL.** The 280 nm absorbance of samples where the peptides have been added is recorded and displayed as a continuous curve while the 280 nm absorbance of RAD51 WT in absence of peptides is displayed as a dashed line. **A.** Overlay of the elution profiles of RAD51 in presence of BRC1 (red), BRC2 (green), BRC3 (blue) and BRC4 (violet) **B.** Elution profile in presence of BRC4 **C.** Elution profile in presence of BRC6 **D.** Elution profile in presence of BRC7 **E.** Elution profile in presence of BRC8 **F.** RAD51 elution profile in absence of peptides.

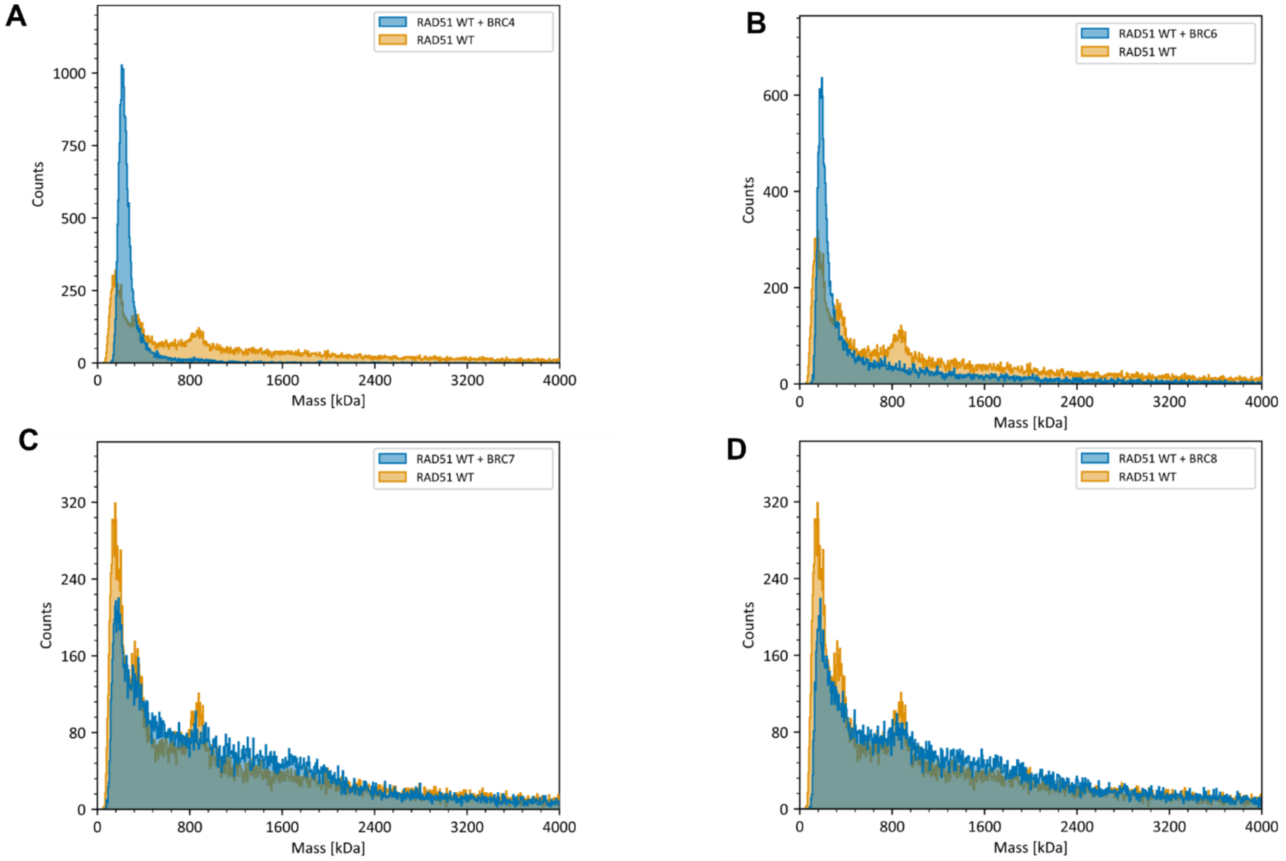


**Supplementary Figure 51 Mass Photometry analysis of the RAD51 WT mass distribution in presence (yellow) or absence (blue) of different peptides at final peptide/protein stoichiometric ratio 4:1. A.** In presence of BRC4, **B.** In presence of BRC6 **C.** In presence of BRC7, **D** In presence of BRC8.

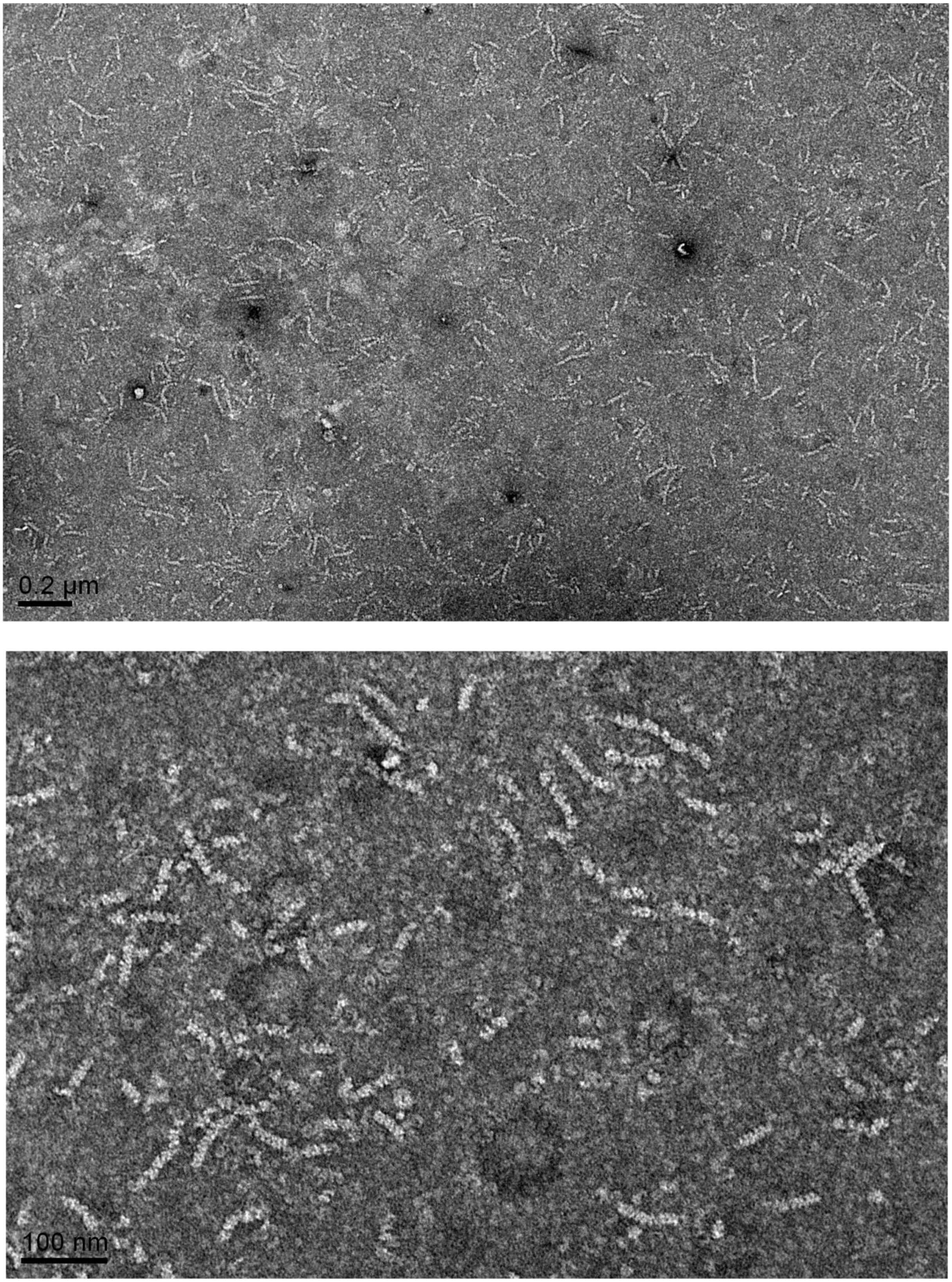


**Supplementary Figure 52** Two representative micrographs of RAD51 WT in the absence of BRC repeat peptides. It is clearly possible to observe the self-assembled RAD51 fibrils as worm-like structures. In the following micrographs it is possible to qualitatively appreciate a reduction of fibril length as highlighted by the statistical analysis reported in the main figures.

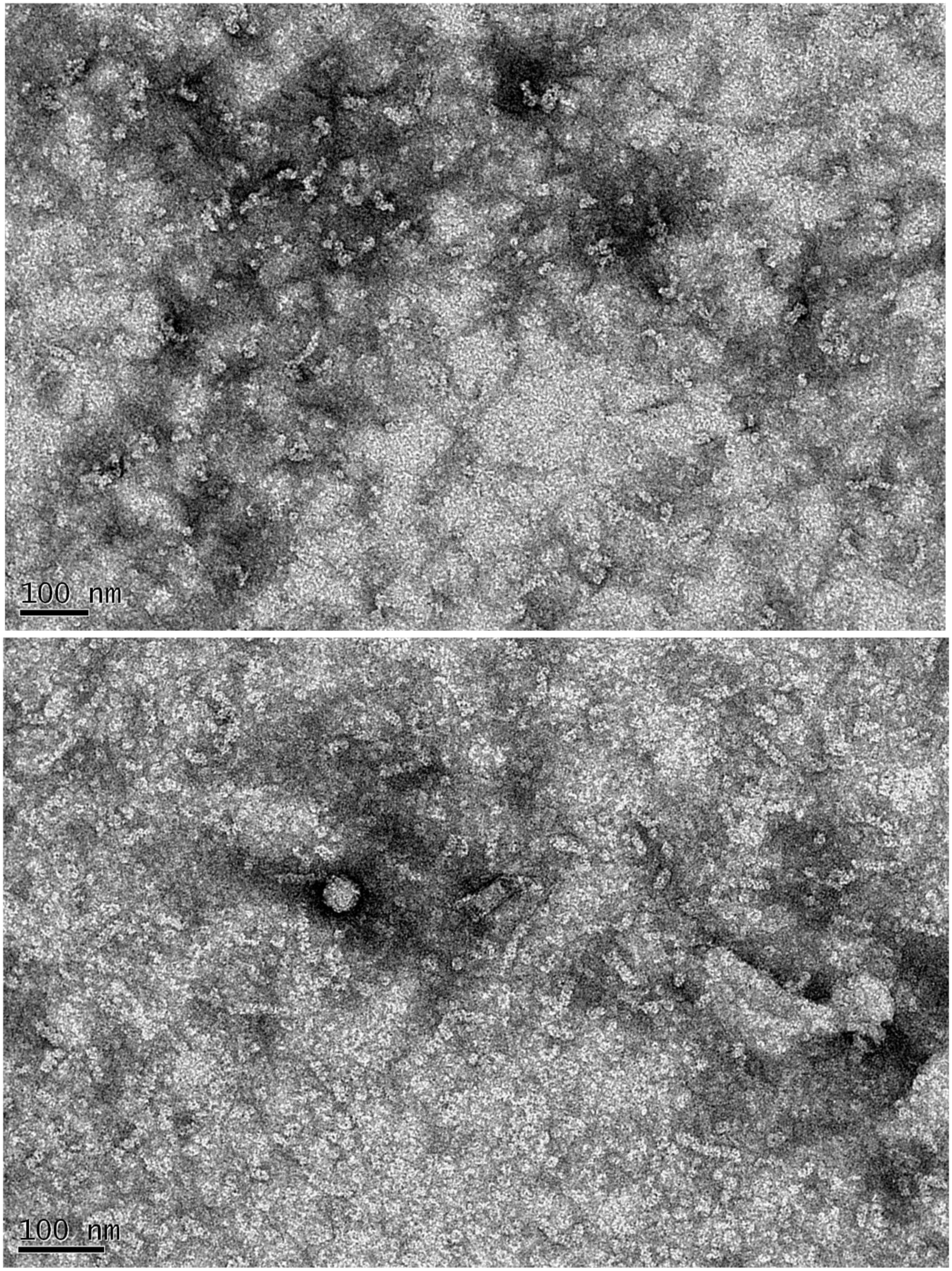


**Supplementary Figure 53** Two representative micrographs of RAD51 WT in the presence of BRC1 repeat in protein/peptide stoichiometric ratio Top. 1/0.4, Bottom. 1/0.6

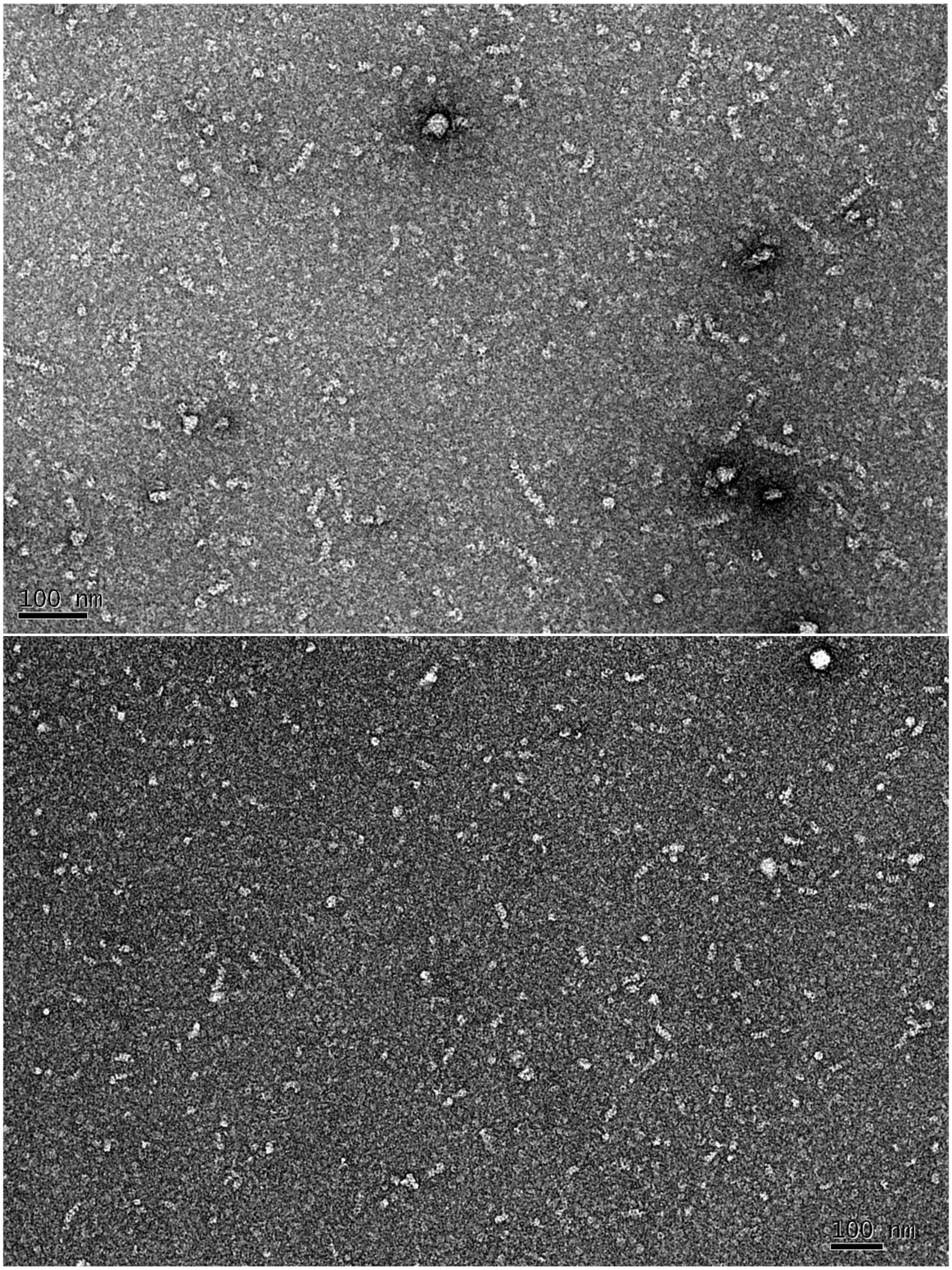


**Supplementary Figure 54** Two representative micrographs of RAD51 WT in the presence of BRC1 repeat in protein/peptide stoichiometric ratio Top. 1/0.8 Bottom. 1/1

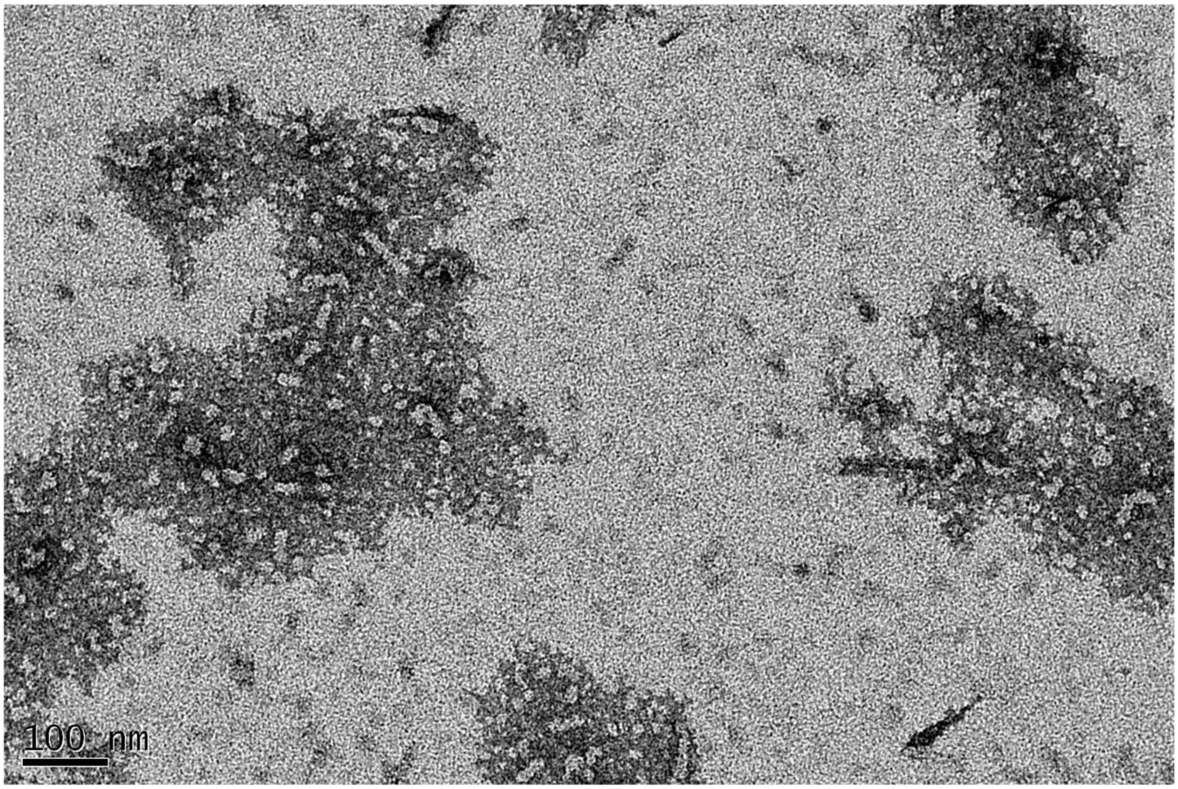


**Supplementary Figure 55** A representative micrographs of RAD51 WT in the presence of BRC1 repeat in protein/peptide stoichiometric ratio 1/2

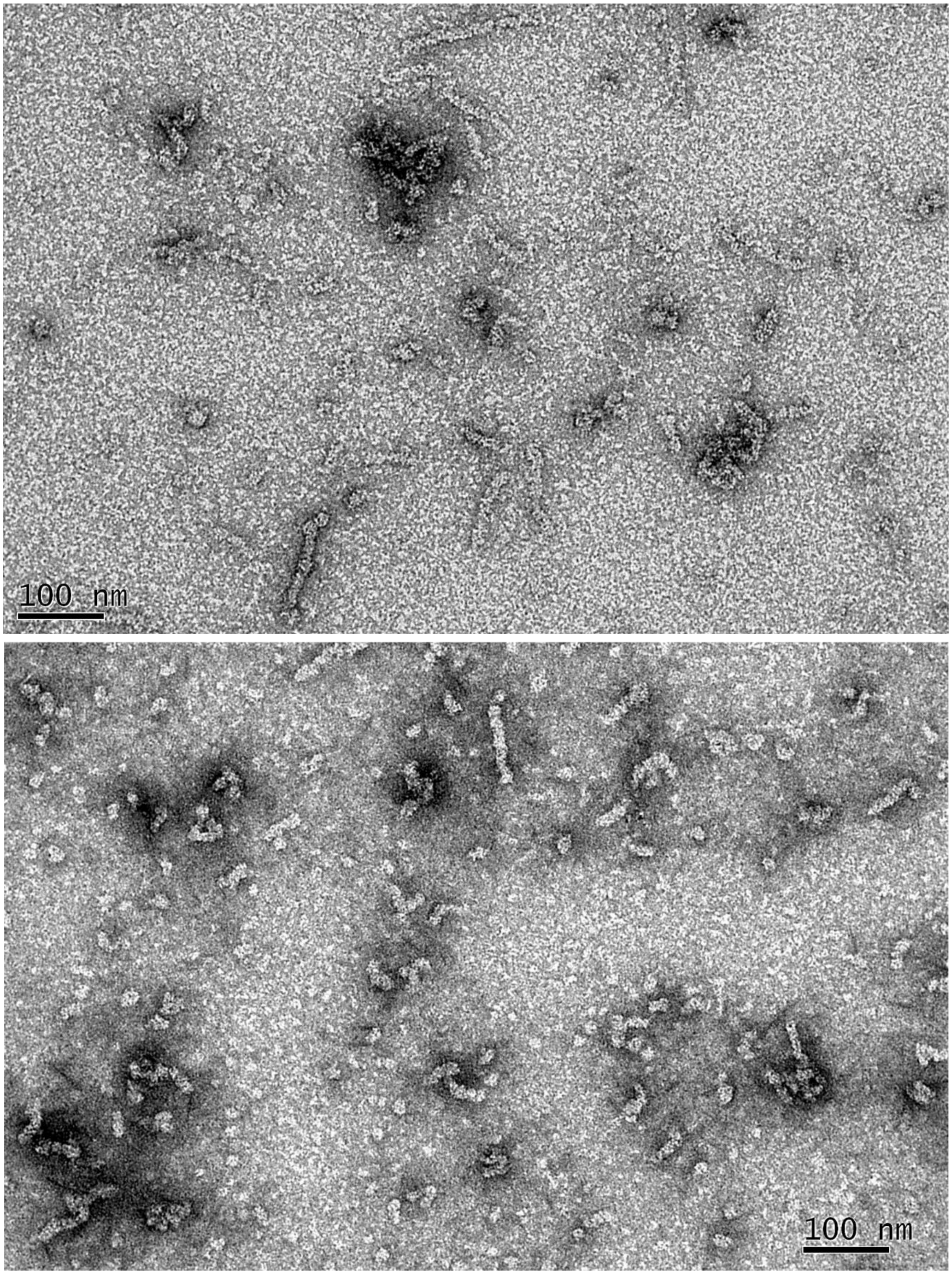


**Supplementary Figure 56** Two representative micrographs of RAD51 WT in the presence of BRC2 repeat in protein/peptide stoichiometric ratio Top. 1/0.4 Bottom. 1/0.6

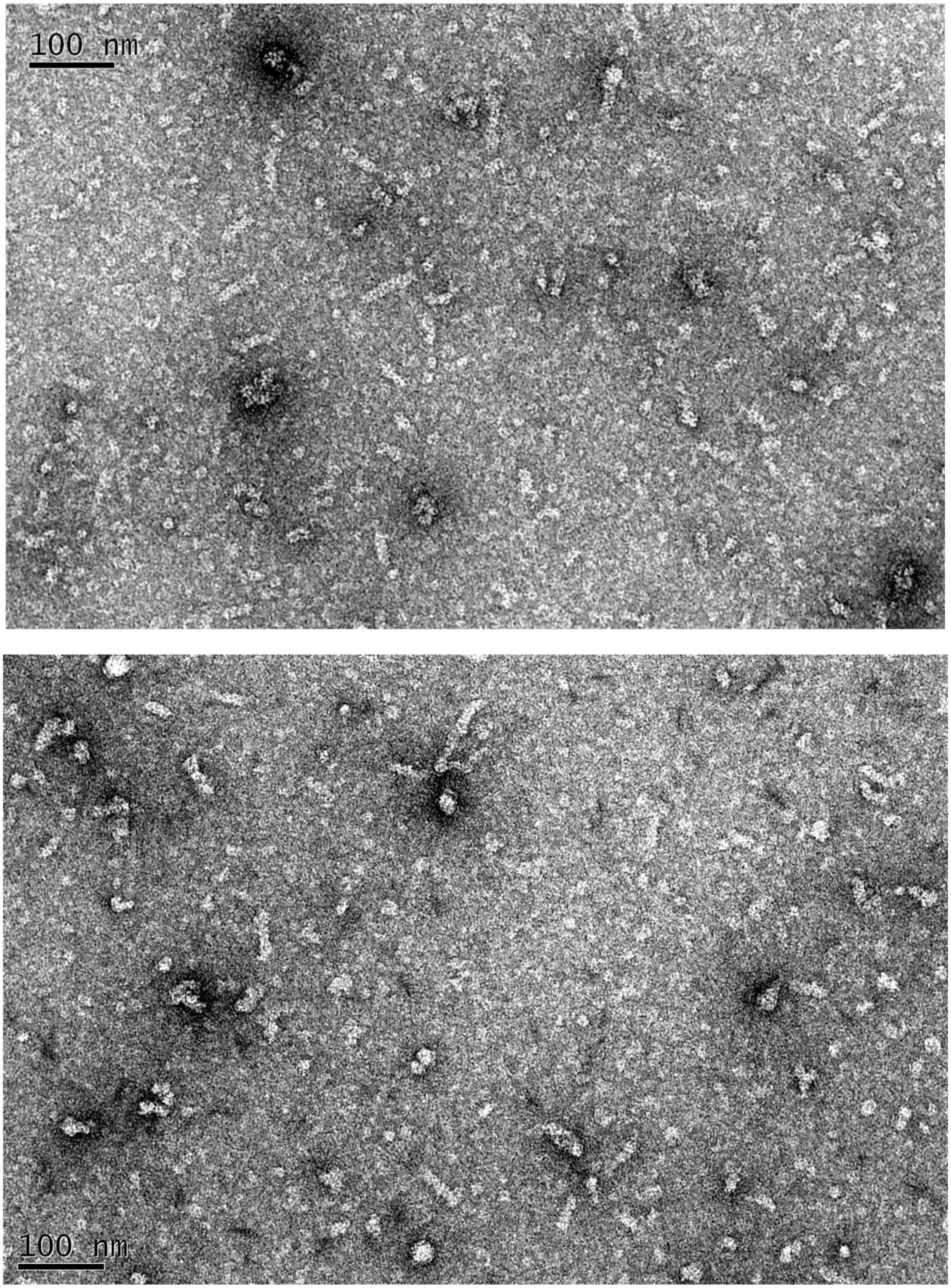


**Supplementary Figure 57** Two representative micrographs of RAD51 WT in the presence of BRC2 repeat in protein/peptide stoichiometric ratio Top. 1/0.8 Bottom. 1/1

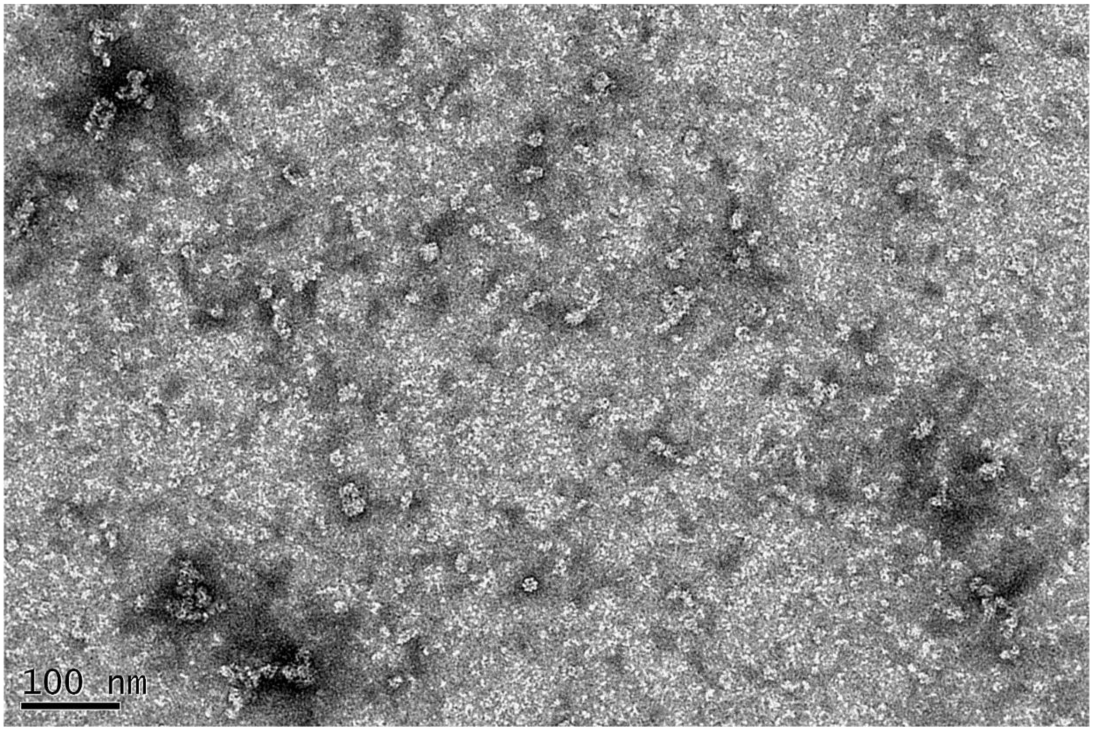


**Supplementary Figure 58** A representative micrograph of RAD51 WT in the presence of BRC2 repeat in protein/peptide stoichiometric ratio 1/2

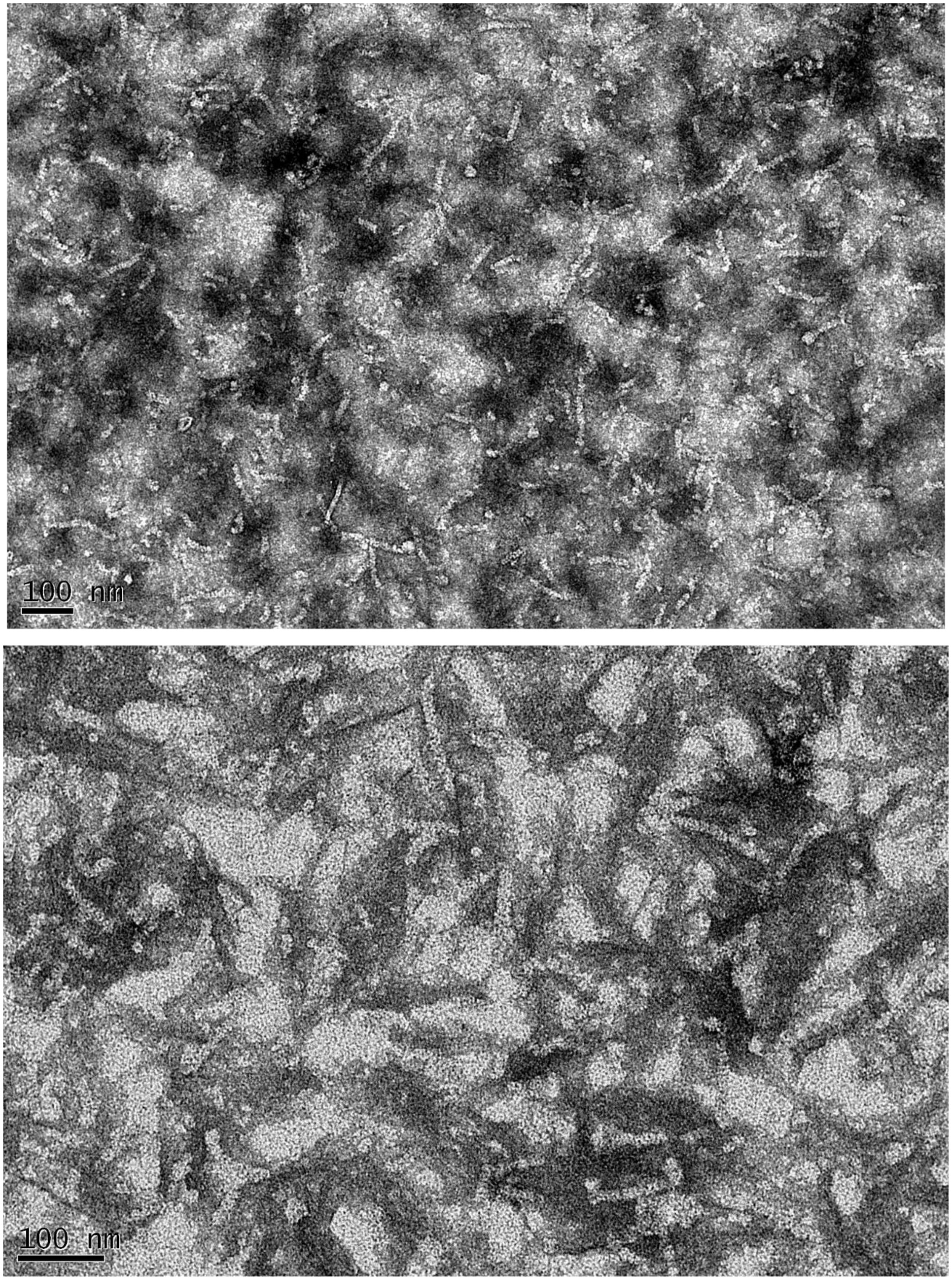


**Supplementary Figure 59** Two representative micrographs of RAD51 WT in the presence of BRC3 repeat in protein/peptide stoichiometric ratio Top. 1/0.4 Bottom 1/0.6

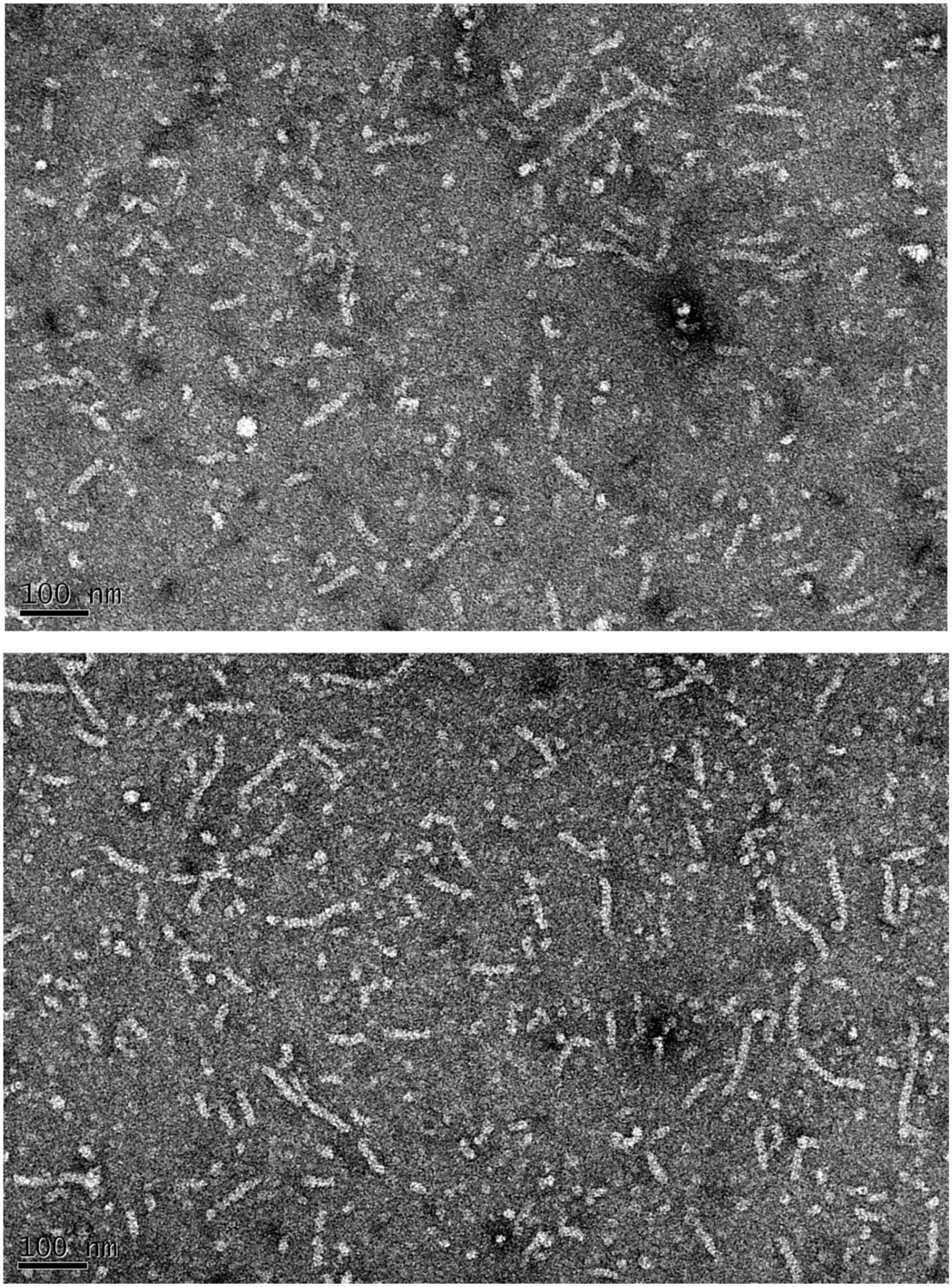


**Supplementary Figure 60** Two representative micrographs of RAD51 WT in the presence of BRC3 repeat in protein/peptide stoichiometric ratio Top 1/0.8 Bottom 1/1

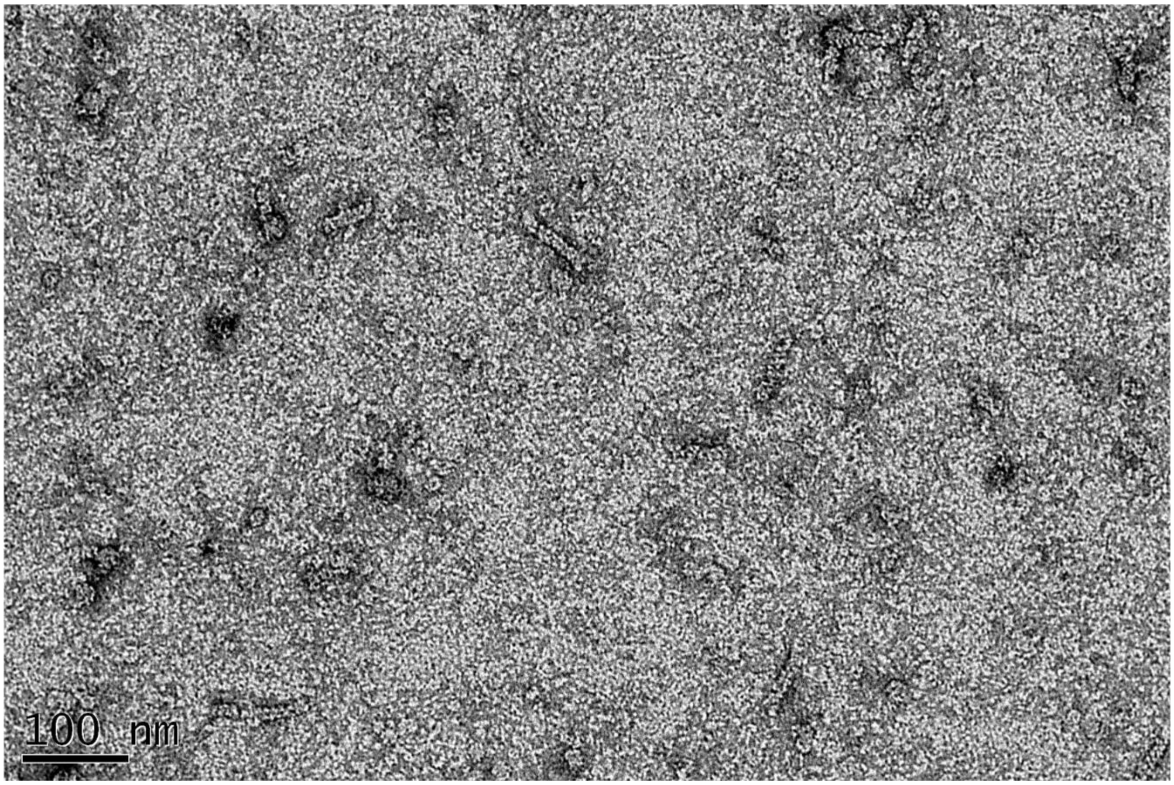


**Supplementary Figure 61** Two representative micrographs of RAD51 WT in the presence of BRC3 repeat in protein/peptide stoichiometric ratio 1/2

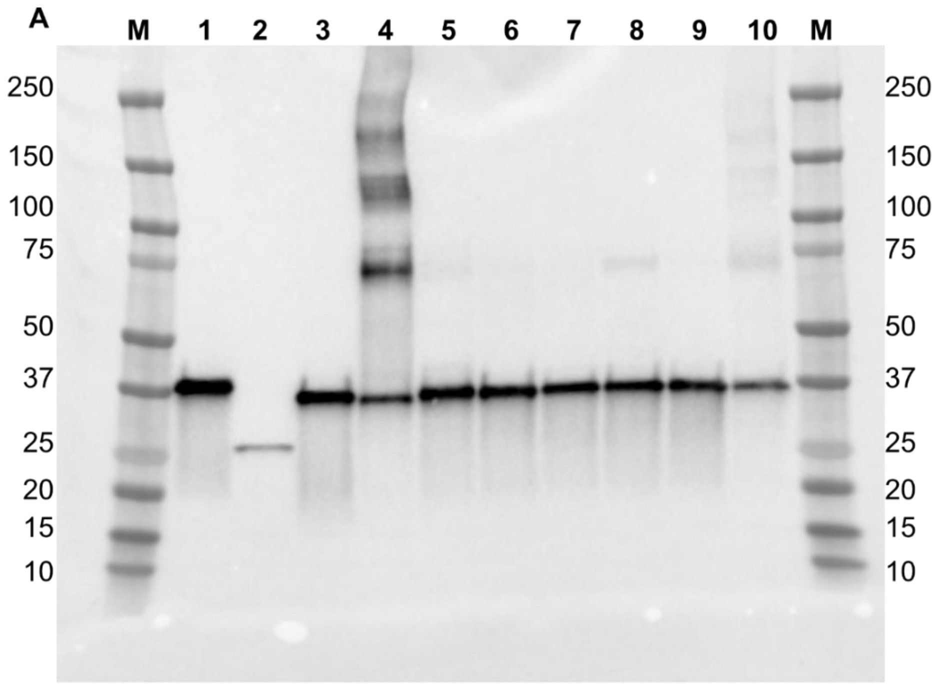


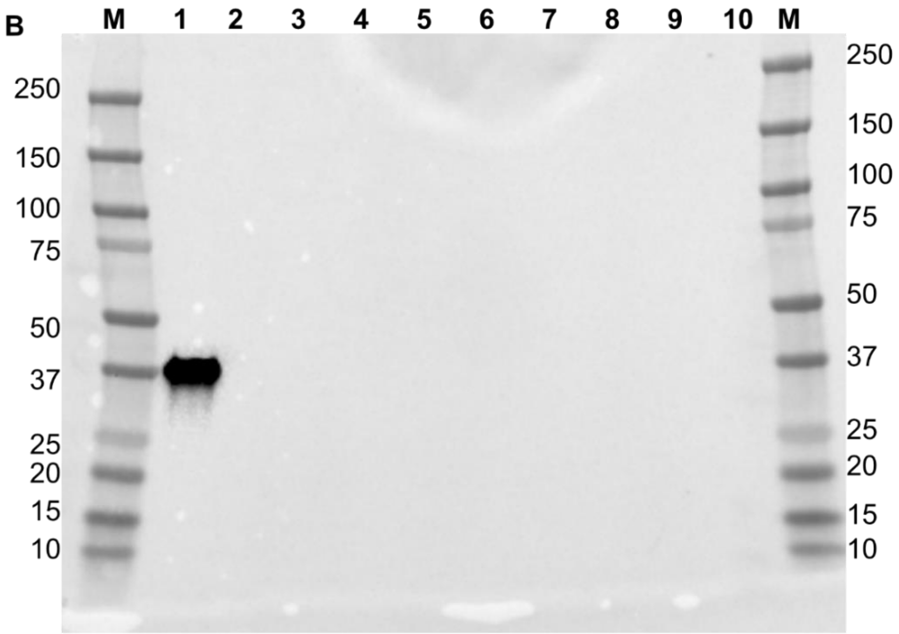


**Supplementary Figure 62 Evaluation of TEV-Cleavage efficiency by Western Blot analyses. A. Western blot using an anti-RAD51 antibody. M** = Marker **1** = His-RAD51-FL/His-BRC4 **2** = Cleaved Δ97-RAD51/BRC4 complex **3** = Cleaved RAD51-FL/BRC4 complex **4** = Cleaved RAD51-FL/BRC2 **5** = Cleaved RAD51-FL/BRC1 complex **6** = Cleaved RAD51-FL/BRC1-2 complex **7** = Cleaved RAD51-FL/BRC2-3 complex **8** = Cleaved RAD51-FL/BRC3-4 complex **9** = Cleaved RAD51-FL/BRC2-4 **10** = Cleaved RAD51-FL/BRC1-4 complex **B. Western blot using an anti-His antibody M** = Marker **1** = His-RAD51-FL/His-BRC4 **2** = Cleaved Δ97-RAD51/BRC4 complex **3** = Cleaved RAD51-FL/BRC4 complex **4** = Cleaved RAD51-FL/BRC2 **5** = Cleaved RAD51-FL/BRC1 complex **6** = Cleaved RAD51-FL/BRC1-2 complex **7** = Cleaved RAD51-FL/BRC2-3 complex **8** = Cleaved RAD51-FL/BRC3-4 complex **9** = Cleaved RAD51-FL/BRC2-4 **10** = Cleaved RAD51-FL/BRC1-4 complex

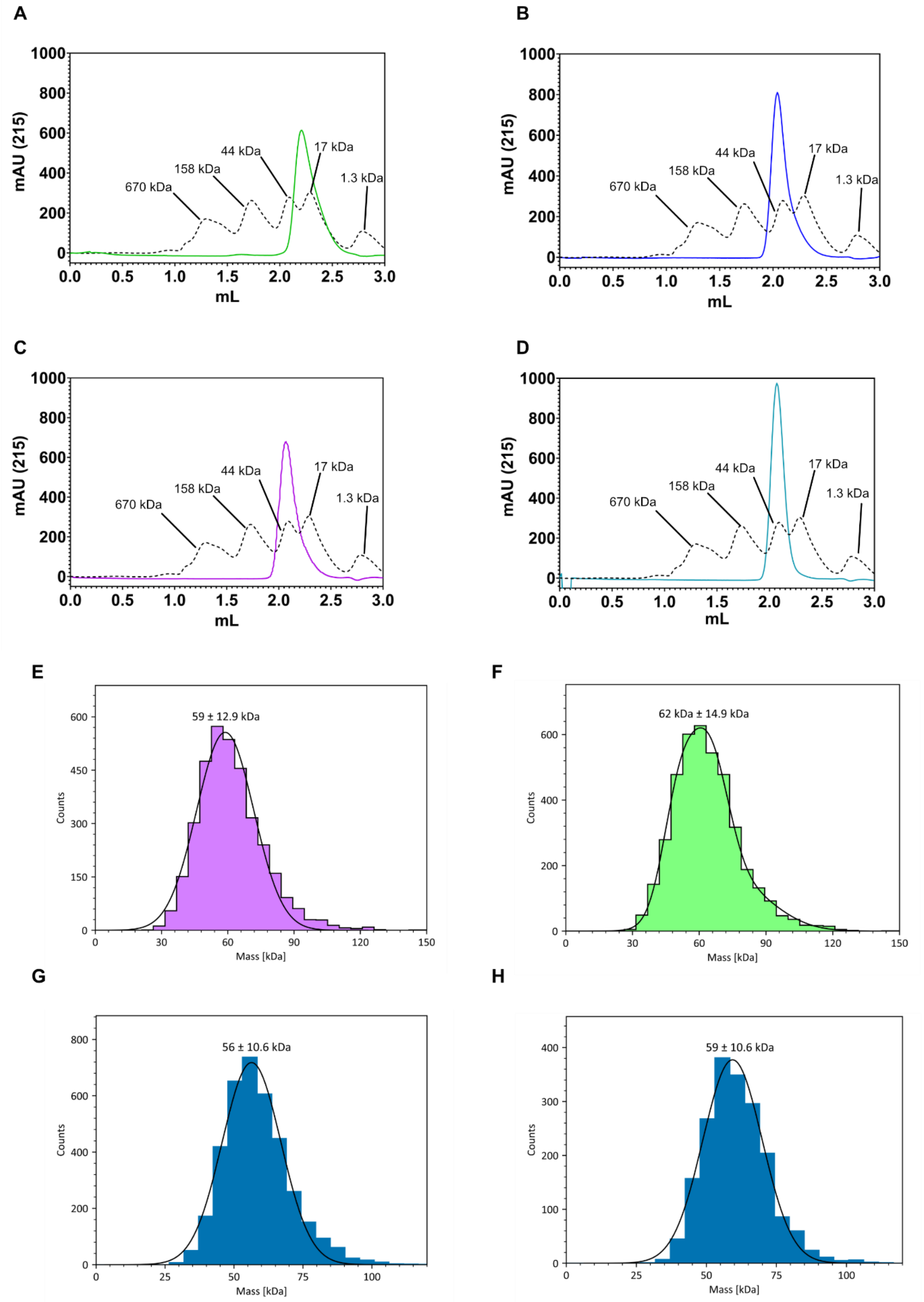


**Supplementary Figure 63 Characterization of the endogenously expressed single BRC-repeats RAD51complexes. Analytical SEC elution profiles** of **A.** Δ97-RAD51/BRC4 **B.** RAD51-FL/BRC4 **C.** RAD51-FL/BRC1 **D.** RAD51-FL/BRC2. Elution profile of globular molecular weight markers are shown as a dashed black line. **Mass Photometry analysis of E.** RAD51-FL/BRC1 **B.** RAD51-FL/BRC2 **C.** RAD51-FL/BRC4 **D.** Δ97-RAD51/BRC4

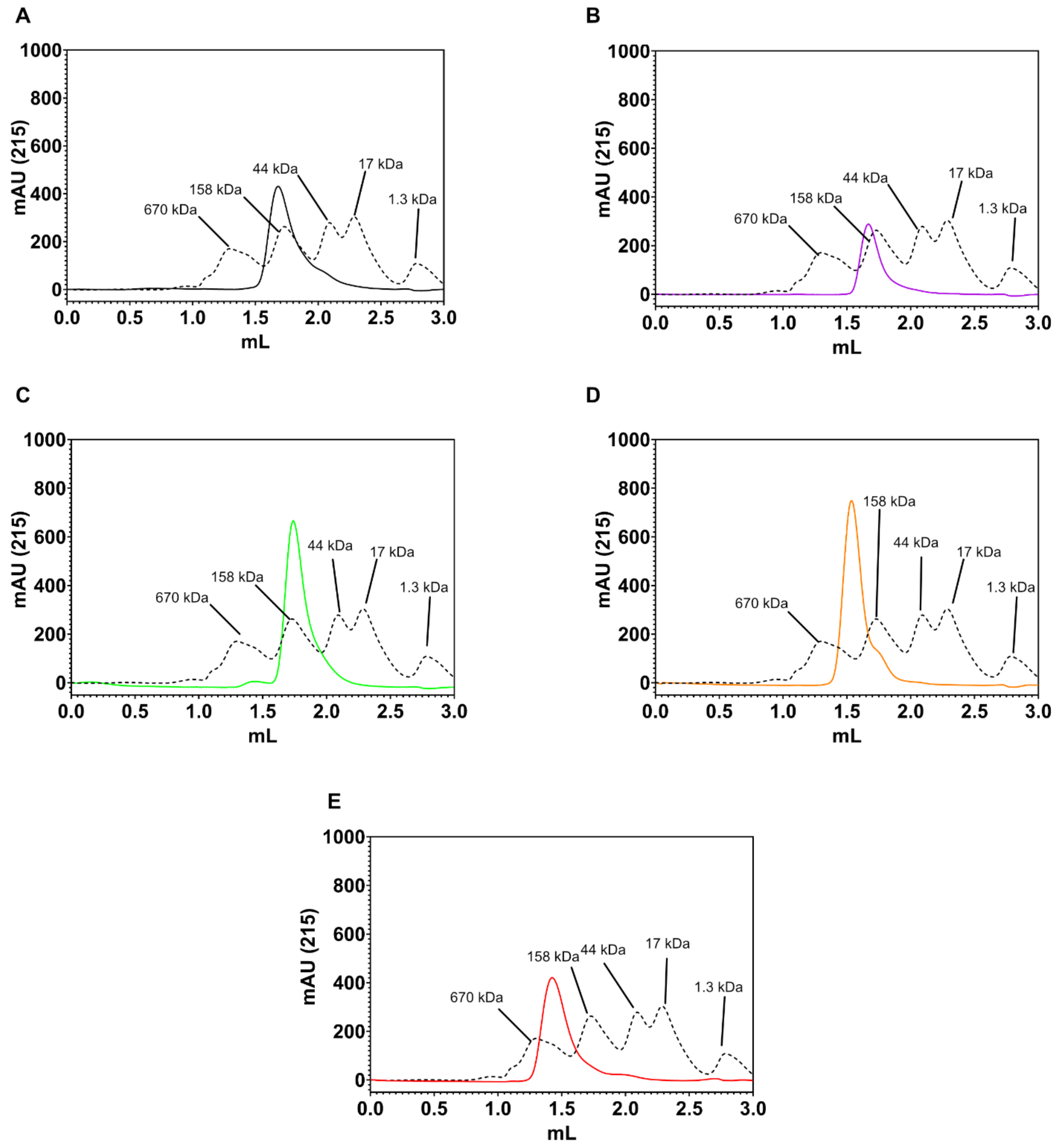


**Supplementary Figure 64 Characterization of the endogenously expressed BRCA2 Truncates-RAD51 complexes. Analytical SEC elution profiles** of **A.** RAD51-FL/BRC1-2 **B.** RAD51-FL/BRC2-3 **C.** RAD51-FL/BRC3-4 **D.** RAD51-FL/BRC2-4. **E.** RAD51-FL/BRC1-4. Elution profile of globular molecular weight markers are shown as a dashed black line.

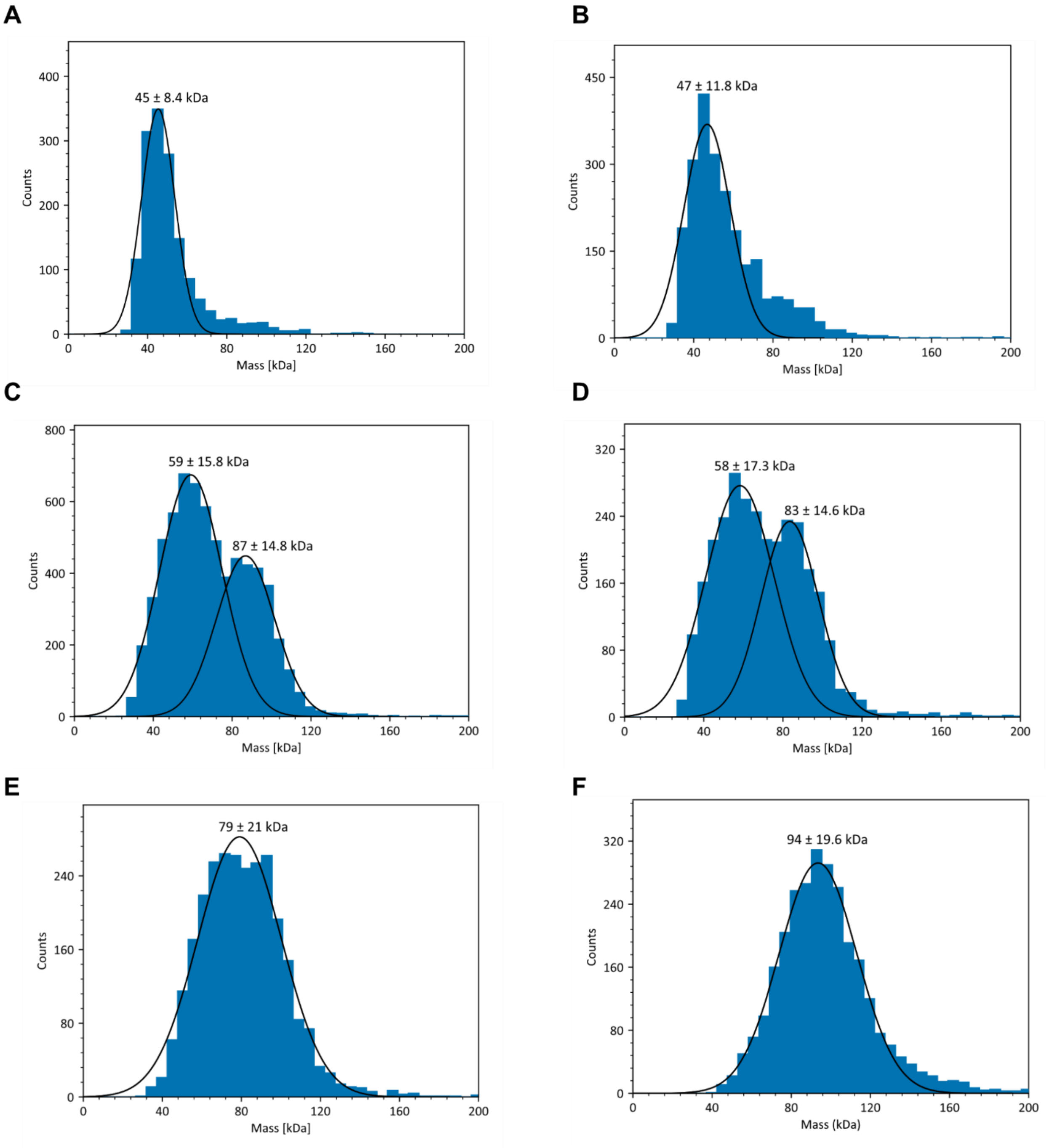


**Supplementary Figure 65 Mass Photometry evaluation of RAD51-FL/BRC1-2 complex stability at different concentrations. A.** 5 nM, **B.** 10 nM, **C.** 25 nM**, D.** 50 nM, **E.** 75 nM **F.** 100 nM

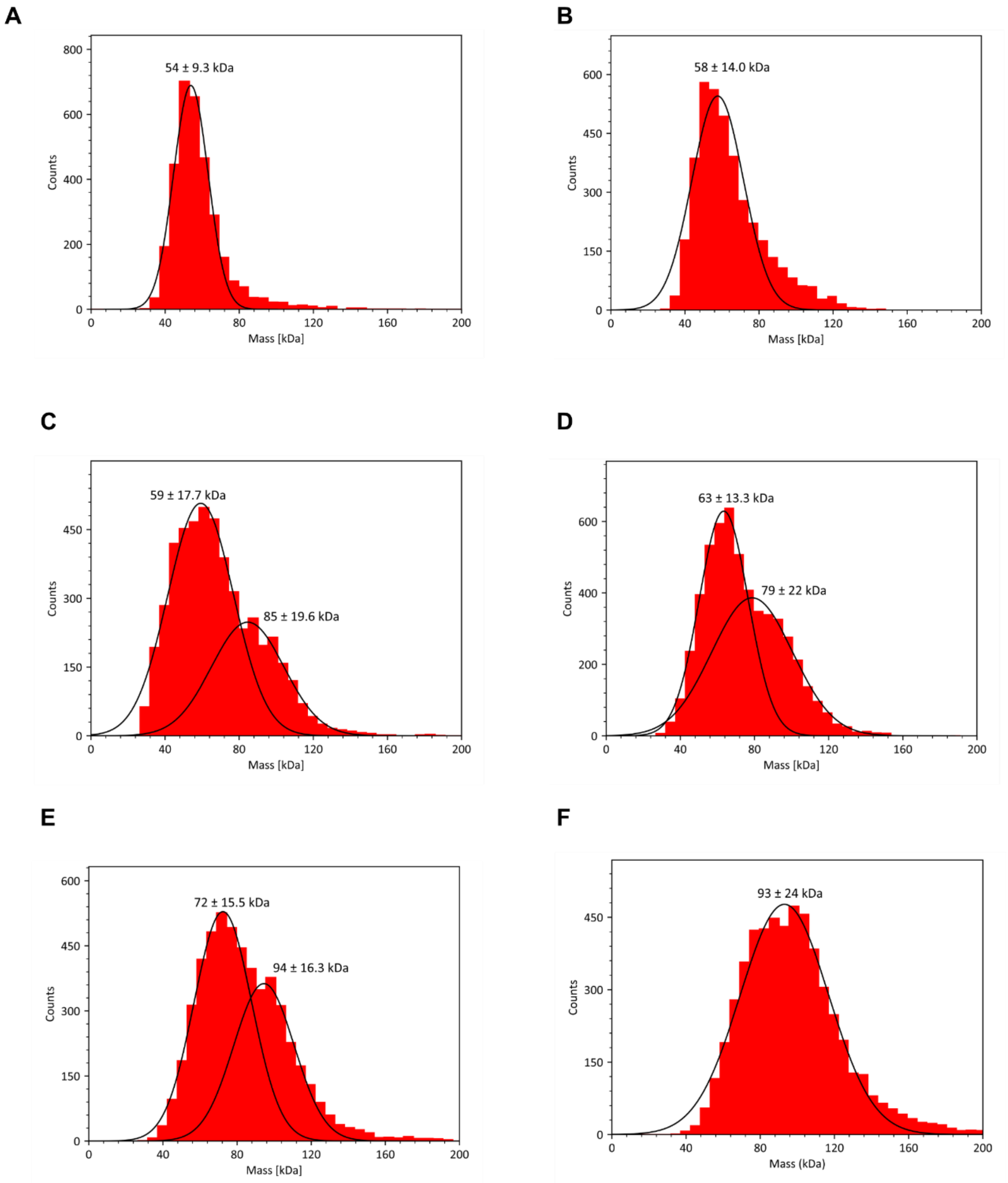


**Supplementary Figure 66 Mass Photometry evaluation of RAD51-FL/BRC2-3 complex stability at different concentrations. A.** 5 nM, **B.** 10 nM, **C.** 25 nM**, D.** 50 nM, **E.** 75 nM **F.** 100 nM

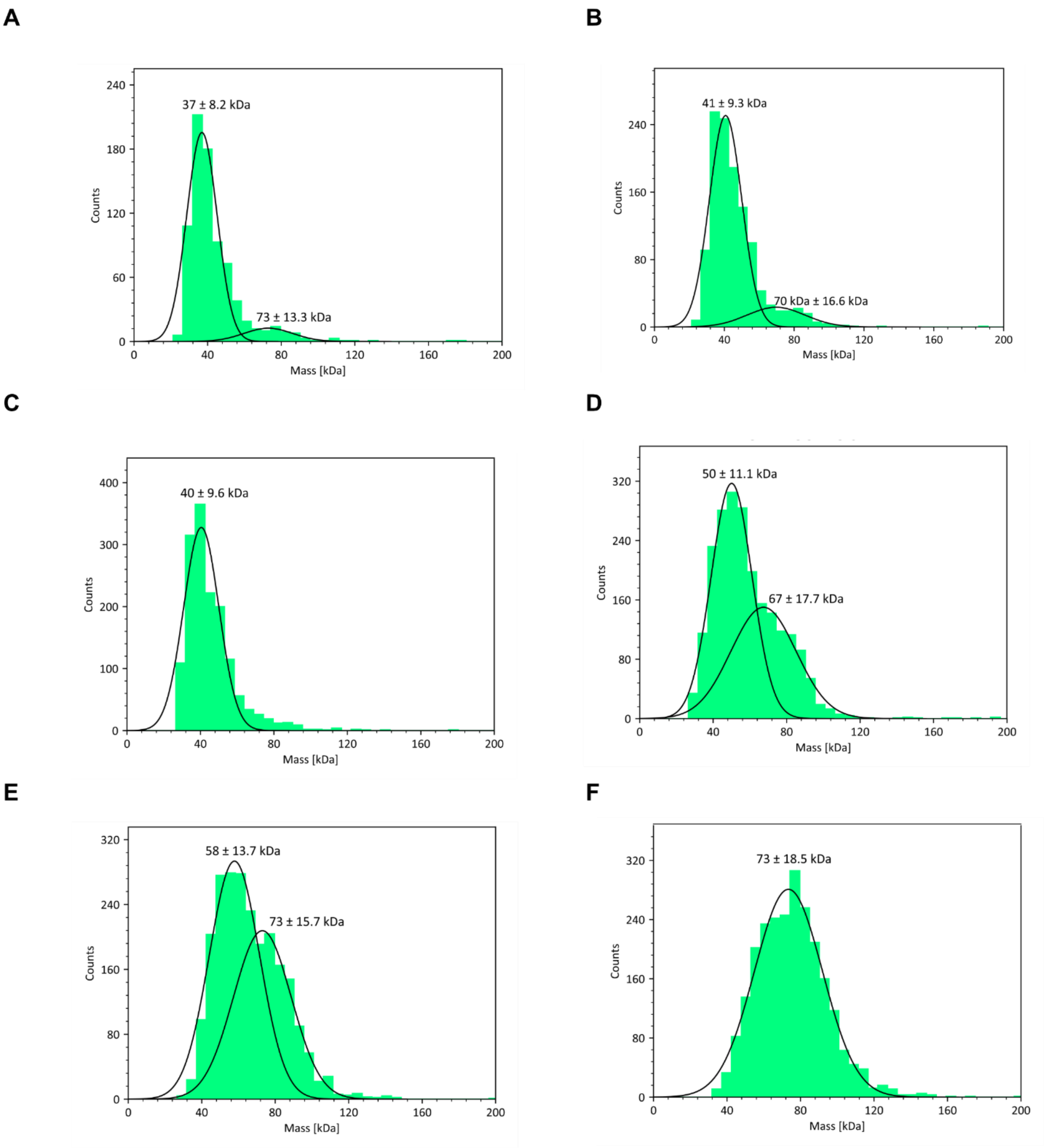


**Supplementary Figure 67 Mass Photometry evaluation of RAD51-FL/BRC3-4 complex stability at different concentrations. A.** 5 nM, **B.** 10 nM, **C.** 25 nM**, D.** 50 nM, **E.** 75 nM **F.** 100 nM

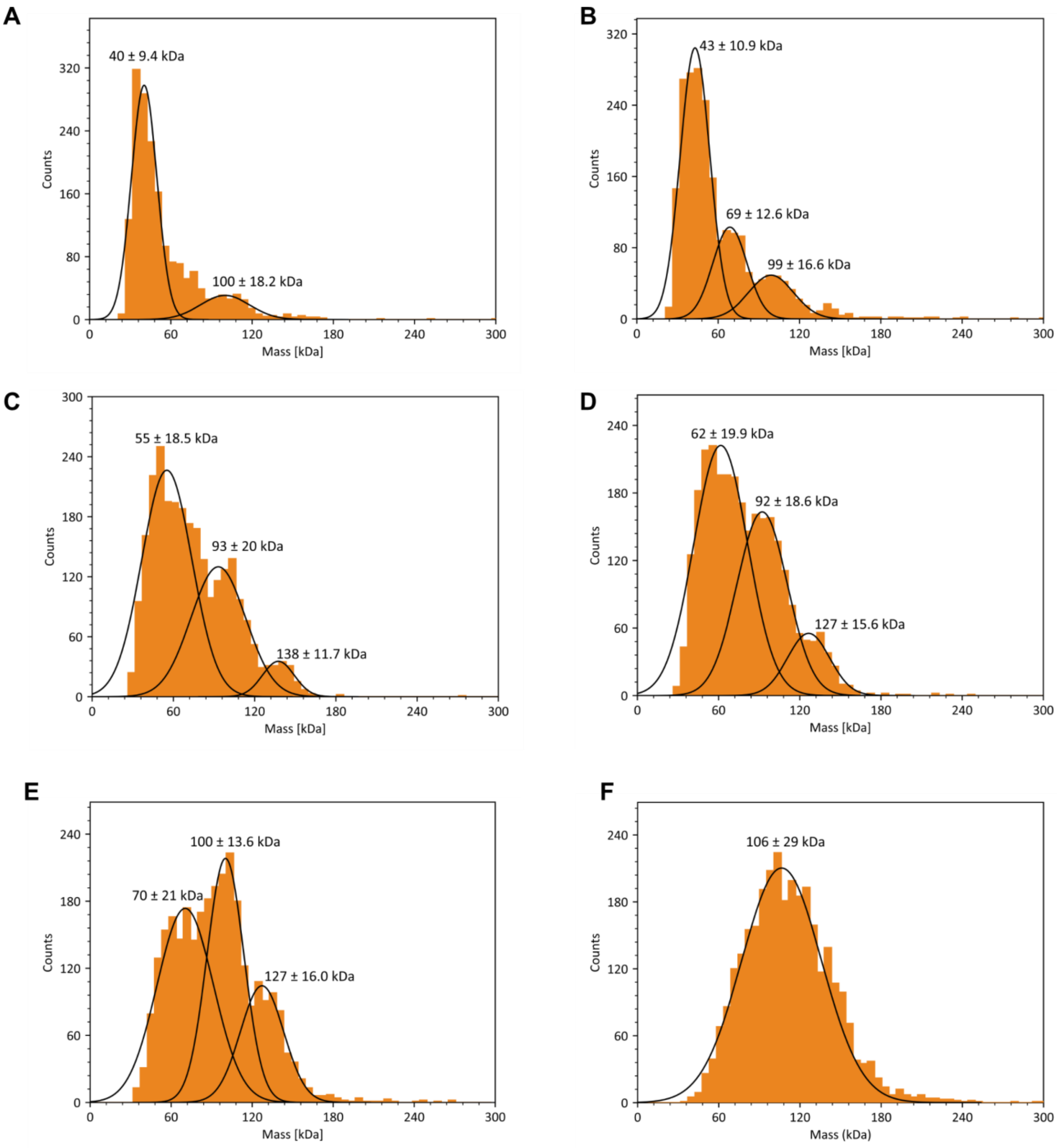


**Supplementary Figure 68 Mass Photometry evaluation of RAD51-FL/BRC3-4 complex stability at different concentrations. A.** 5 nM, **B.** 10 nM, **C.** 25 nM**, D.** 50 nM, **E.** 75 nM **F.** 100 nM

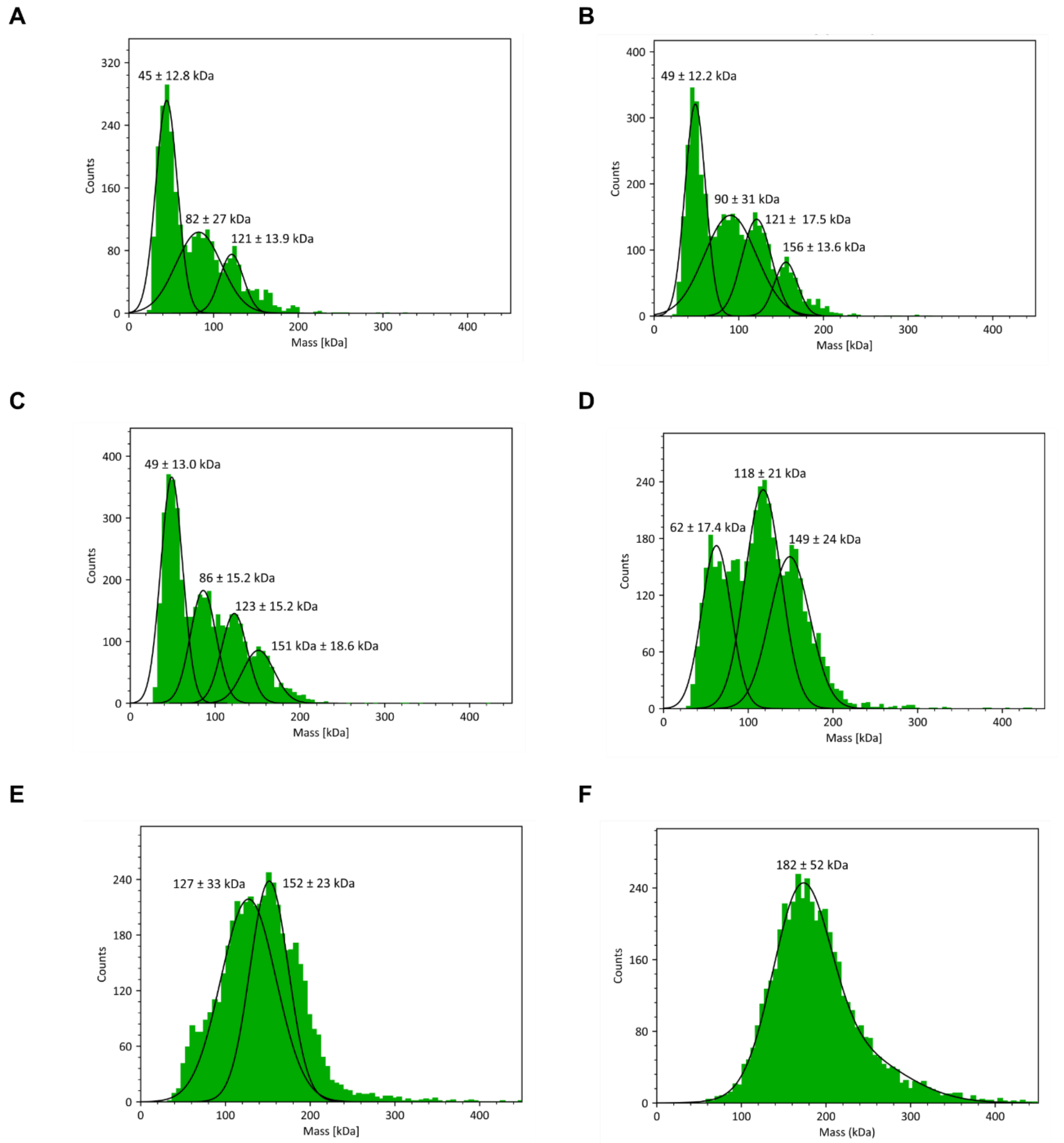


**Supplementary Figure 69 Mass Photometry evaluation of RAD51-FL/BRC1-4 complex stability at different concentrations. A.** 5 nM, **B.** 10 nM, **C.** 25 nM**, D.** 50 nM, **E.** 75 nM **F.** 100 nM

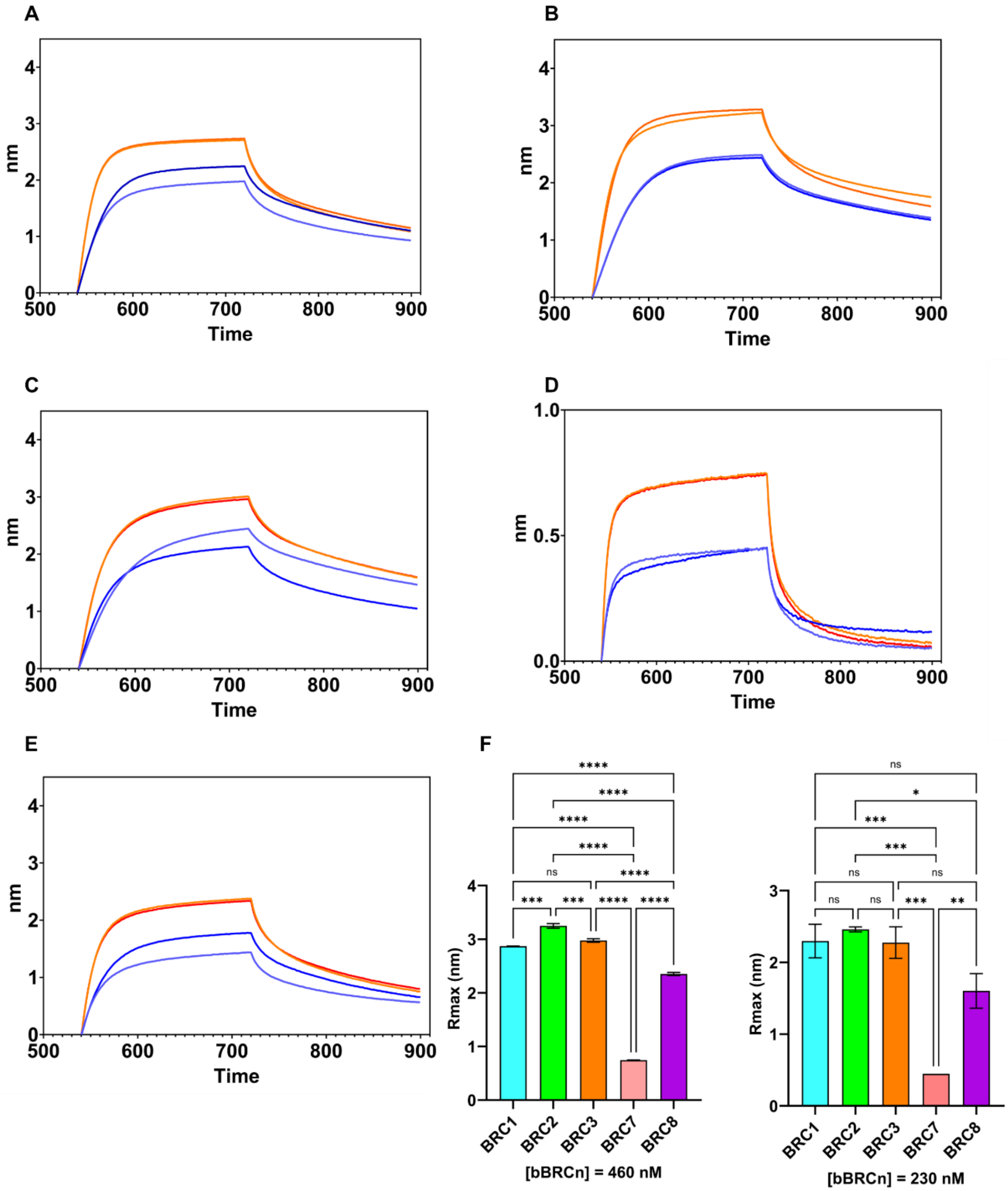


**Supplementary Figure 70 Biolayer interferometry binding tests of the biotinylated BRC-repeats (bioBRCn) peptides to the monomeric His-RAD51 [F86E, A89E].** In each plot are reported the overlay of biolayer interferometry (BLI) sensorgrams showing the binding of monomeric His-RAD51[F86E, A89E] to bioBRCn obtained with two different protein concentrations: 460 nM (red/orange), 230 nM (blue/light blue). For each concentration, two replicates of the measurements were performed **A.** bioBRC1 **B.** bioBRC2 **C.** bioBRC3 **D.** bioBRC7 **E.** bioBRC8 **F.** Mean ± standard deviation of maximum response (Rmax) obtained for BLI experiments carried out at the same protein concentration. In each plot are reported ordinary one-way ANOVA statistical analysis of the plotted Rmax.

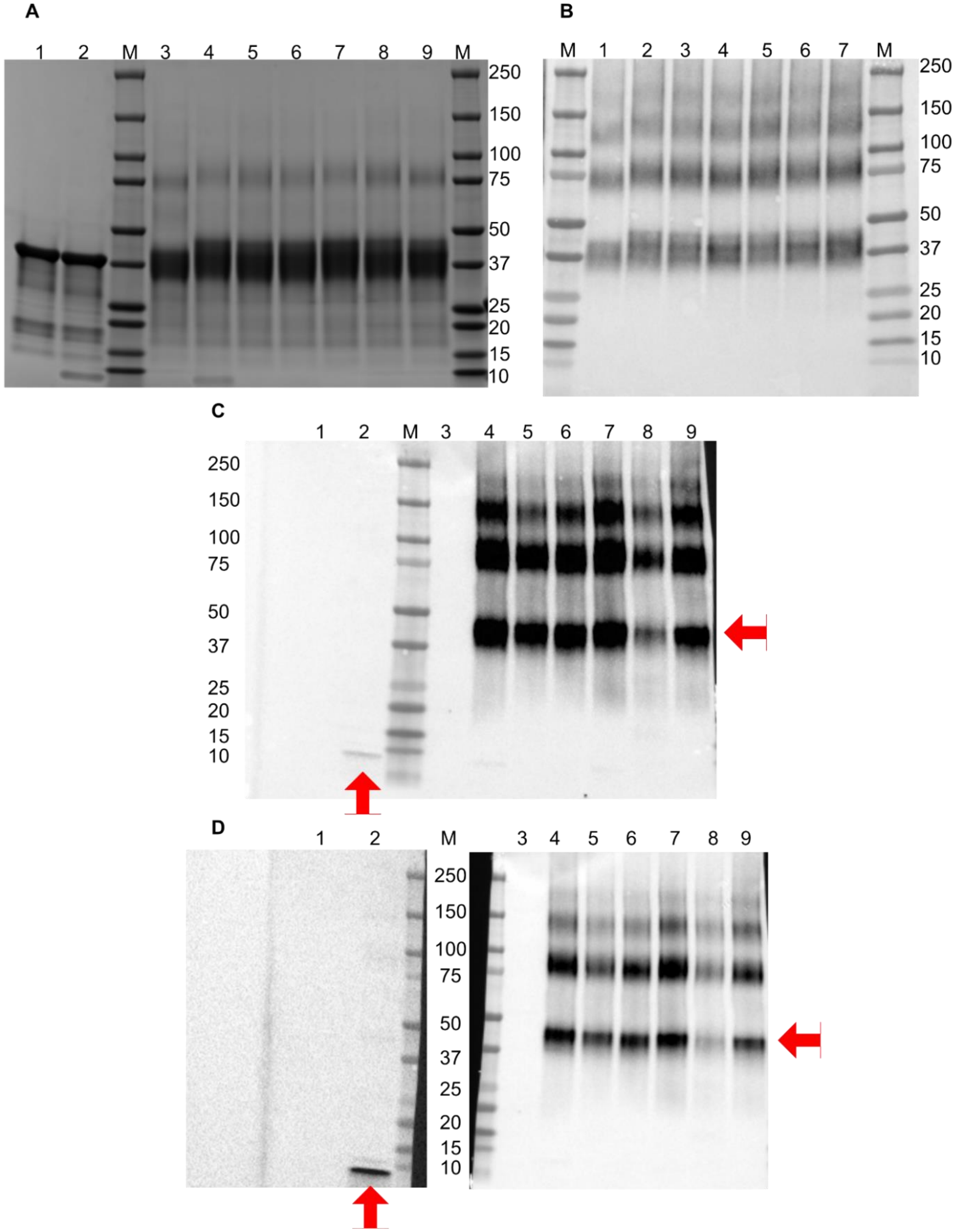


**Supplementary Figure 71 Evaluation of the cross-linking efficiency of the RAD51** **A.** Coomassie Blue SDS-Page analysis of the cross-linked bio-BRC repeats His-RAD51 [F86E, A89E] complexes. M = Marker, 1 = His-RAD51 [F86E, A89E] 2 = His-RAD51 [F86E,A89E] + bioBRC4 (1:1) 3 = His-RAD51 [F86E, A89E] + 1 mM BS3 4 = His-RAD51 [F86E,A89E] + bioBRC1(1:1) 1 mM BS3 5 = His-RAD51 [F86E,A89E] + bioBRC2 (1:1) 1 mM BS3 6 = His-RAD51 [F86E,A89E] + bioBRC3 (1:1) 1 mM BS3 7 = His-RAD51 [F86E,A89E] + bioBRC4 (1:1) 1 mM BS3 8 = His-RAD51 [F86E,A89E] + bioBRC7 (1:1) 1 mM BS3 9 = His-RAD51 [F86E,A89E] + bioBRC8 (1:1) 1 mM BS3 **B.** Western blot using an anti-RAD51 antibody. M = Marker, 1 = His-RAD51 [F86E, A89E] + 1 mM BS3 2 = His-RAD51 [F86E,A89E] + bioBRC1(1:1) 1 mM BS3 3 = His-RAD51 [F86E,A89E] + bioBRC2 (1:1) 1 mM BS3 4 = His-RAD51 [F86E,A89E] + bioBRC3 (1:1) 1 mM BS3 5 = His-RAD51 [F86E,A89E] + bioBRC4 (1:1) 1 mM BS3 6 = His-RAD51 [F86E,A89E] + bioBRC7 (1:1) 1 mM BS3 7 = His-RAD51 [F86E,A89E] + bioBRC8 (1:1) 1 mM BS3 **C.** Western blot using Streptavidin-HRP **D.** The same Western blot reported in **C.** where the membrane was cut and re-developed with higher exposure times to increase the signal of the bioBRC4. Red arrows highlight the shift of the peptides' signals M = Marker, 1 = His-RAD51 [F86E, A89E] 2 = His-RAD51 [F86E,A89E] + bioBRC4 (1:1) 3 = His-RAD51 [F86E, A89E] + 1 mM BS3 4 = His-RAD51 [F86E,A89E] + bioBRC1(1:1) 1 mM BS3 5 = His-RAD51 [F86E,A89E] + bioBRC2 (1:1) 1 mM BS3 6 = His-RAD51 [F86E,A89E] + bioBRC3 (1:1) 1 mM BS3 7 = His-RAD51 [F86E,A89E] + bioBRC4 (1:1) 1 mM BS3 8 = His-RAD51 [F86E,A89E] + bioBRC7 (1:1) 1 mM BS3 9 = His-RAD51 [F86E,A89E] + bioBRC8 (1:1) 1 mM BS3

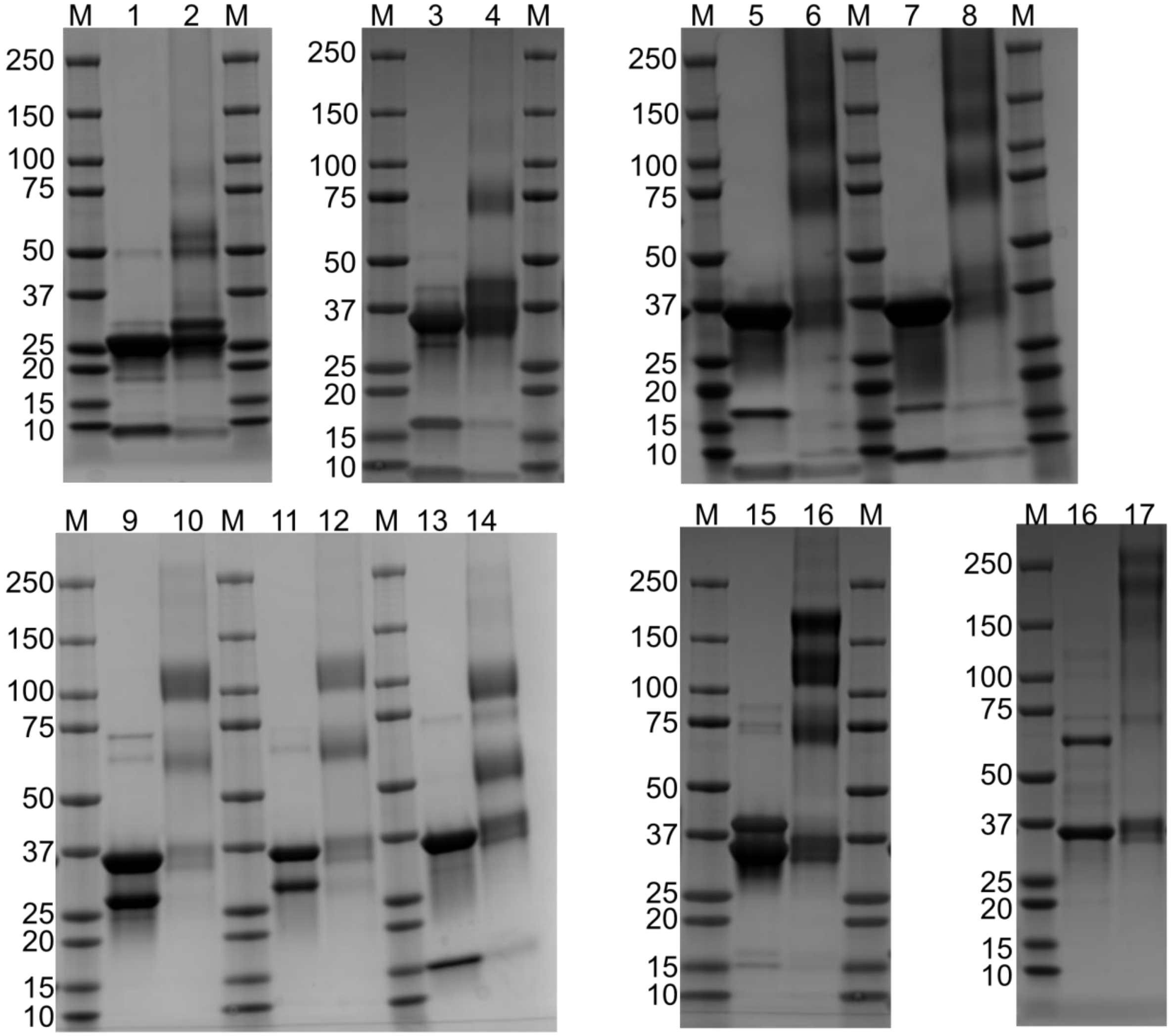


**Supplementary Figure 72 Evaluation of the cross-linking efficiency through Coomassie Blue SDS-Page analysis of the endogenously expressed complexes.** M = Marker, 1 = Δ97-RAD51/BRC4, 2 = Δ97-RAD51/BRC4 BS3 (1 mM), 3 = RAD51-FL/BRC1, 4 = RAD51-FL/BRC1 BS3 (1 mM), 5 = RAD51-FL/BRC2, 6 = RAD51-FL/BRC2 BS3 (1 mM), 7 = RAD51-FL/BRC4, 8 = RAD51-FL/BRC4 BS3 (1 mM), 9 = RAD51-FL/BRC1-2, 10 = RAD51-FL/BRC1-2 BS3 (1 mM), 11 = RAD51-FL/BRC2-3, 12 = RAD51-FL/BRC2-3 BS3 (1 mM), 13 = RAD51-FL/BRC3-4, 14 = RAD51-FL/BRC3-4 BS3(1 mM), 15 = RAD51-FL/BRC2-4, 16 = RAD51 [F86E, A89E]-BRC2-4 BS3 (1 mM), 17 = RAD51 [F86E, A89E]-BRC1-4, 18 = RAD51 [F86E, A89E]-BRC1-4 BS3 (1 mM)

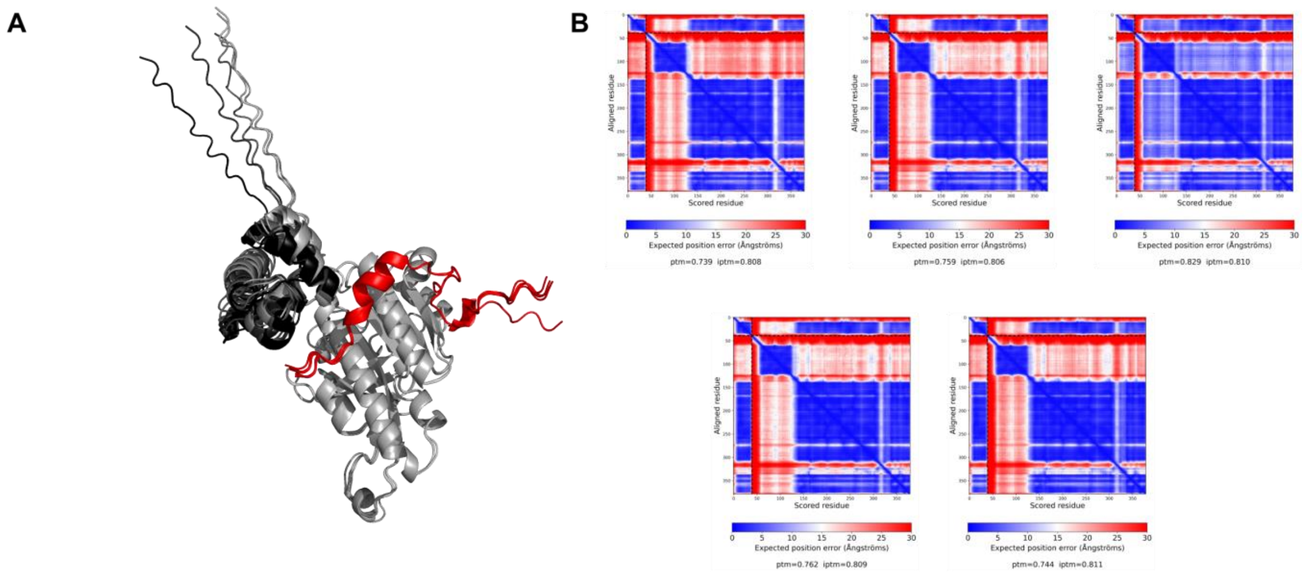


**Supplementary Figure 73 Alphafold2 model generation of full length RAD51-FL/BRC4 complex A.** Overlay of the five best ranked models **B.** Predicted Align Error (PAE) plots of the models displayed in **A.**, providing the expected positional error at residue x if the predicted structure is aligned on residue y

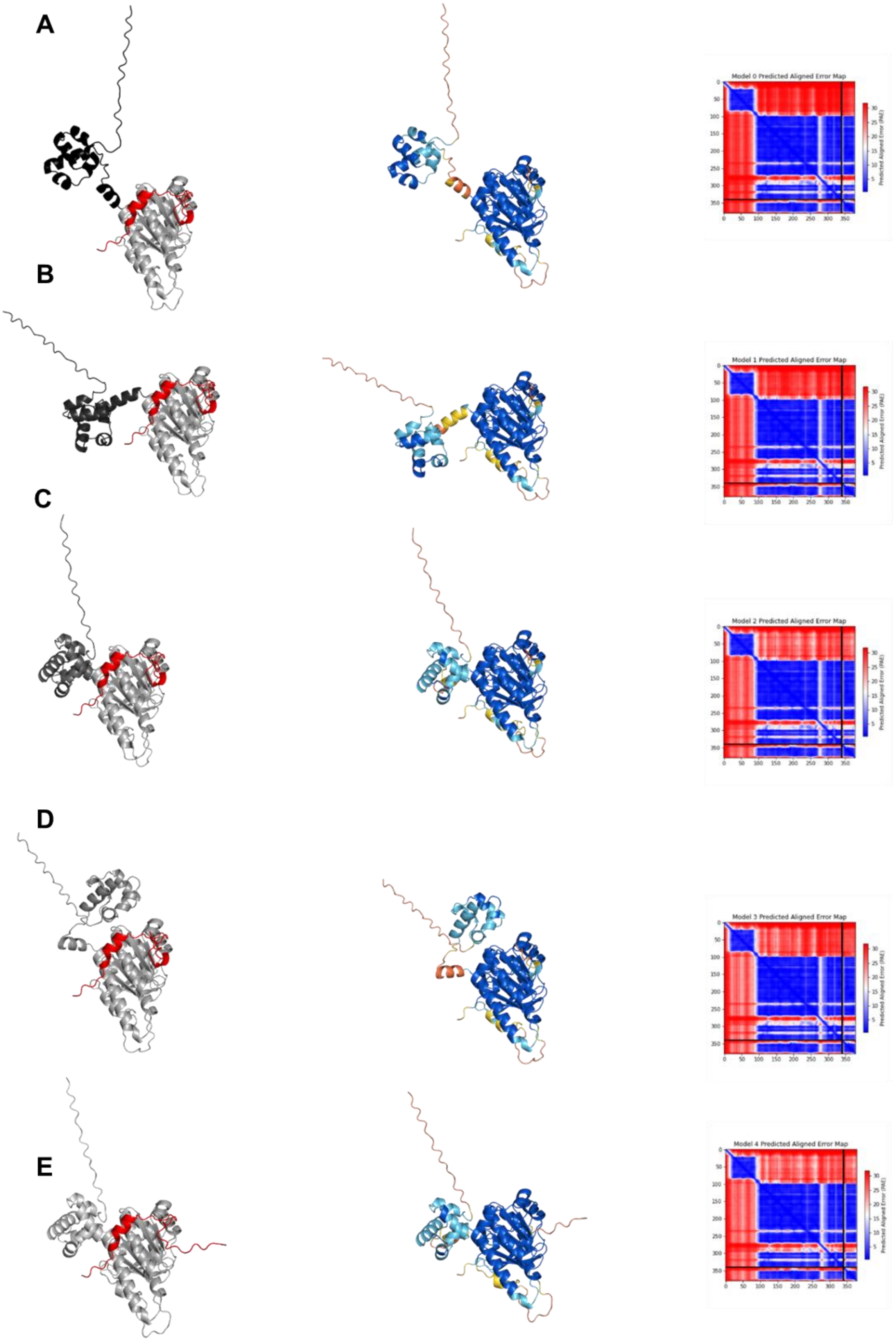


**Supplementary Figure 74 Alphafold3 model generation of full length RAD51-FL/BRC4 complex.** Left: the BRC4 peptide is colored red while the N-Terminal domain is displayed in different shades of grey depending on the model. Center: Residues have been colored for local model confidence estimated with pLDDT. Right: PAE of the corresponding model. **A.** Model 0 **B.** Model 1 **C.** Model 2 **D.** Model 3 **E.** Model 4

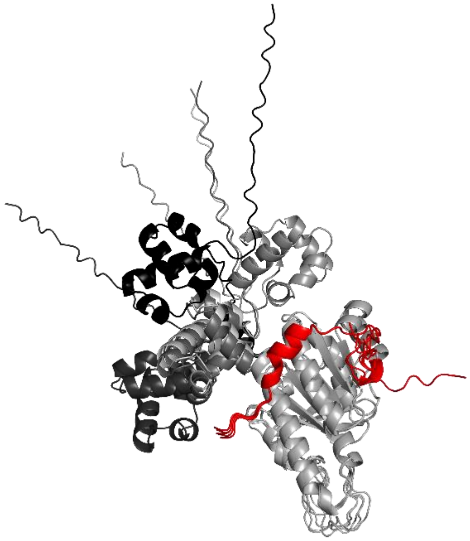

**Supplementary Figure 75 Overlay of the five best ranked AlphaFold3 models of the RAD51-FL/BRC4.**

| XL-Pairs | Distance | | | | |
|---|---|---|---|---|---|
| 58-1536 | 30.38 | 30.09 | 28.59 | 29.67 | 29.06 |
| 73-1536 | 30.61 | 30.73 | 31.52 | 31.07 | 31.19 |
| 40-1541 | 10.19 | 10.27 | 10.95 | 10.31 | 10.78 |
| 58-1541 | 21.76 | 21.49 | 20.42 | 21.12 | 20.67 |
| 64-1541 | 16.04 | 16.1 | 15.83 | 15.76 | 16.02 |
| 73-1541 | 22.76 | 22.69 | 23.15 | 22.93 | 22.92 |
| AF_Model | Model_0 | Model_1 | Model_2 | Model_3 | Model_4 |

**Supplementary Table 1** Table of detected inter-protein crosslinks and their measured Cα-Cα distances in the AlphaFold2 predictions of the RAD51-FL/BRC4 complex.

| XL-Pairs | Distance | | | | |
|---|---|---|---|---|---|
| 58-1536 | 33.34 | 36.61 | 29.81 | 13.95 | 29.67 |
| 73-1536 | 38.7 | 44.86 | 30.77 | 24.77 | 30.63 |
| 40-1541 | 31.76 | 26.03 | 10.18 | 30.05 | 10.02 |
| 58-1541 | 26.08 | 28.49 | 21.32 | 17.16 | 21.1 |
| 64-1541 | 33.77 | 28.24 | 16.05 | 28.97 | 15.7 |
| 73-1541 | 32.9 | 36.15 | 22.59 | 24.01 | 22.65 |
| AF_Model | Model_0 | Model_1 | Model_2 | Model_3 | Model_4 |

**Supplementary Table 2** Table of detected inter-protein crosslinks and their measured Cα-Cα distances in the AlphaFold3 predictions of the RAD51-FL/BRC4 complex.

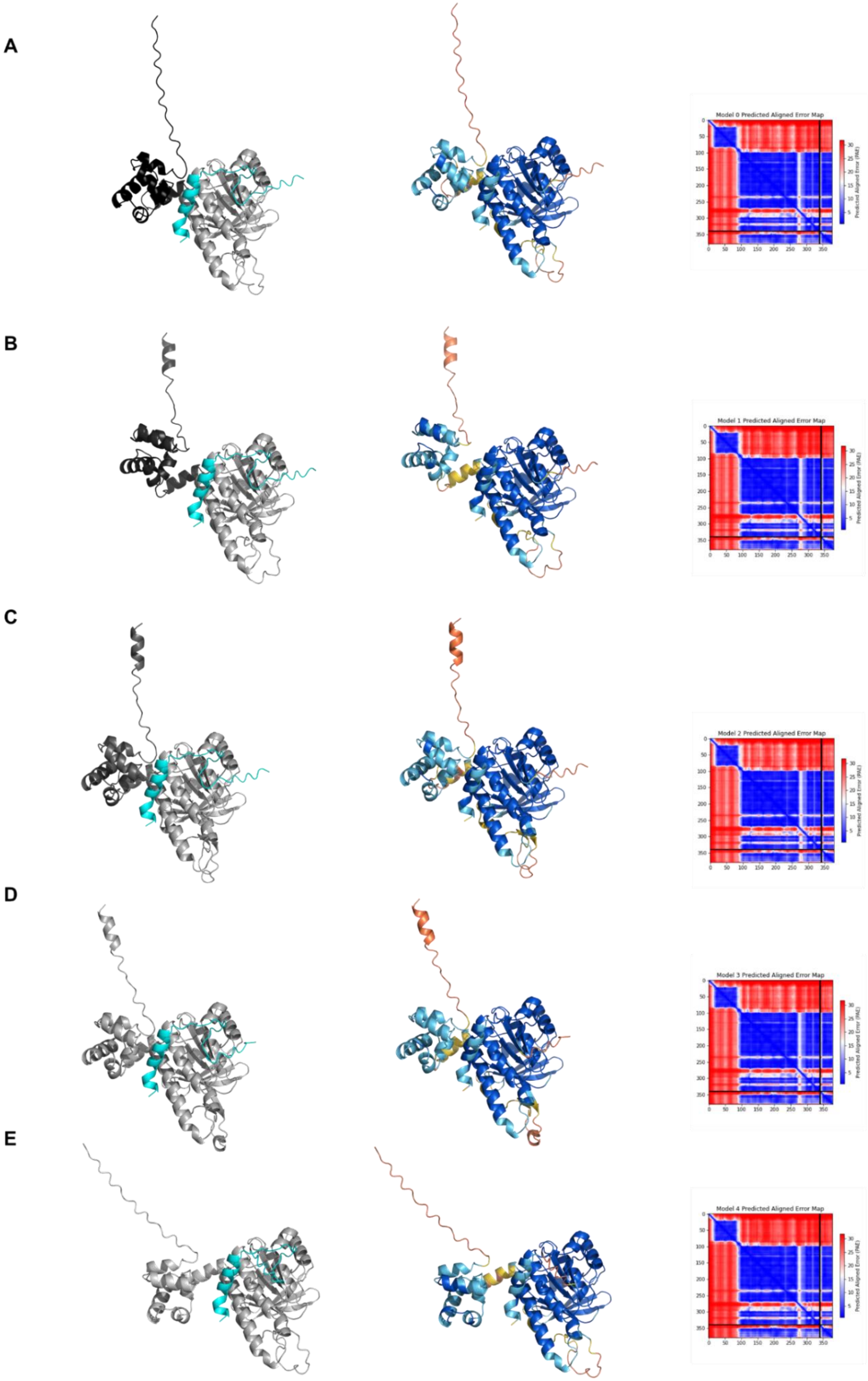


**Supplementary Figure 76** Left: the BRC1 peptide is colored cyan while the N-Terminal domain is displayed in different shades of grey depending on the model. Center: Residues have been colored for local model confidence estimated with pLDDT. Right: PAE of the corresponding model. **A.** Model 0 **B.** Model 1 **C.** Model 2 **D.** Model 3 **E.** Model 4

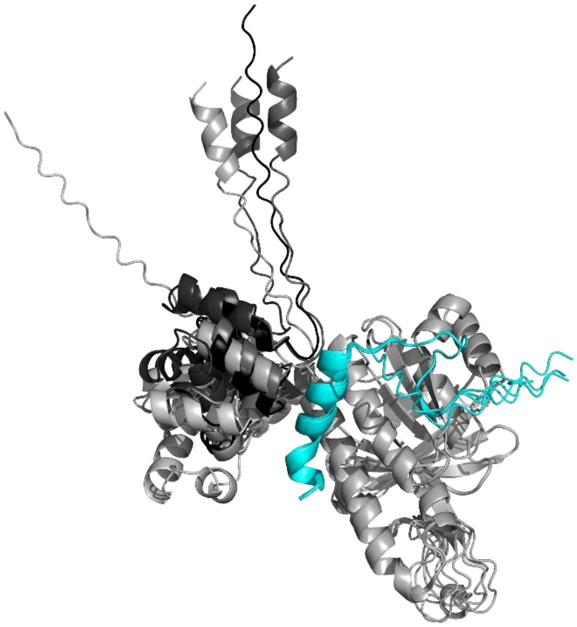

**Supplementary Figure 77 Overlay of the five best ranked AlphaFold3 models of the RAD51-FL/BRC1.**

| XL-Pairs | Distance | | | | |
|---|---|---|---|---|---|
| 40-1015 | 33.94 | 36.81 | 32.81 | 34.38 | 42.34 |
| 58-1015 | 42.07 | 39.53 | 40.46 | 40.71 | 44.45 |
| 73-1015 | 42.92 | 45.68 | 42.59 | 43.93 | 50.18 |
| 156-1015 | 14.43 | 14.38 | 14.47 | 14.49 | 14.5 |
| 73-1018 | 35.27 | 37.8 | 34.88 | 36.19 | 42.38 |
| 156-1018 | 14.99 | 15.04 | 15.16 | 15.26 | 15.29 |
| 58-1025 | 23.74 | 18.91 | 21 | 20.08 | 24.35 |
| 58-1028 | 20.98 | 18.96 | 18.13 | 17.31 | 20.36 |
| AF_Model | Model_0 | Model_1 | Model_2 | Model_3 | Model_4 |

**Supplementary Table 3** Table of detected inter-protein crosslinks and their measured Cα-Cα distances in the AlphaFold3 predictions of the RAD51-FL/BRC1 complex.

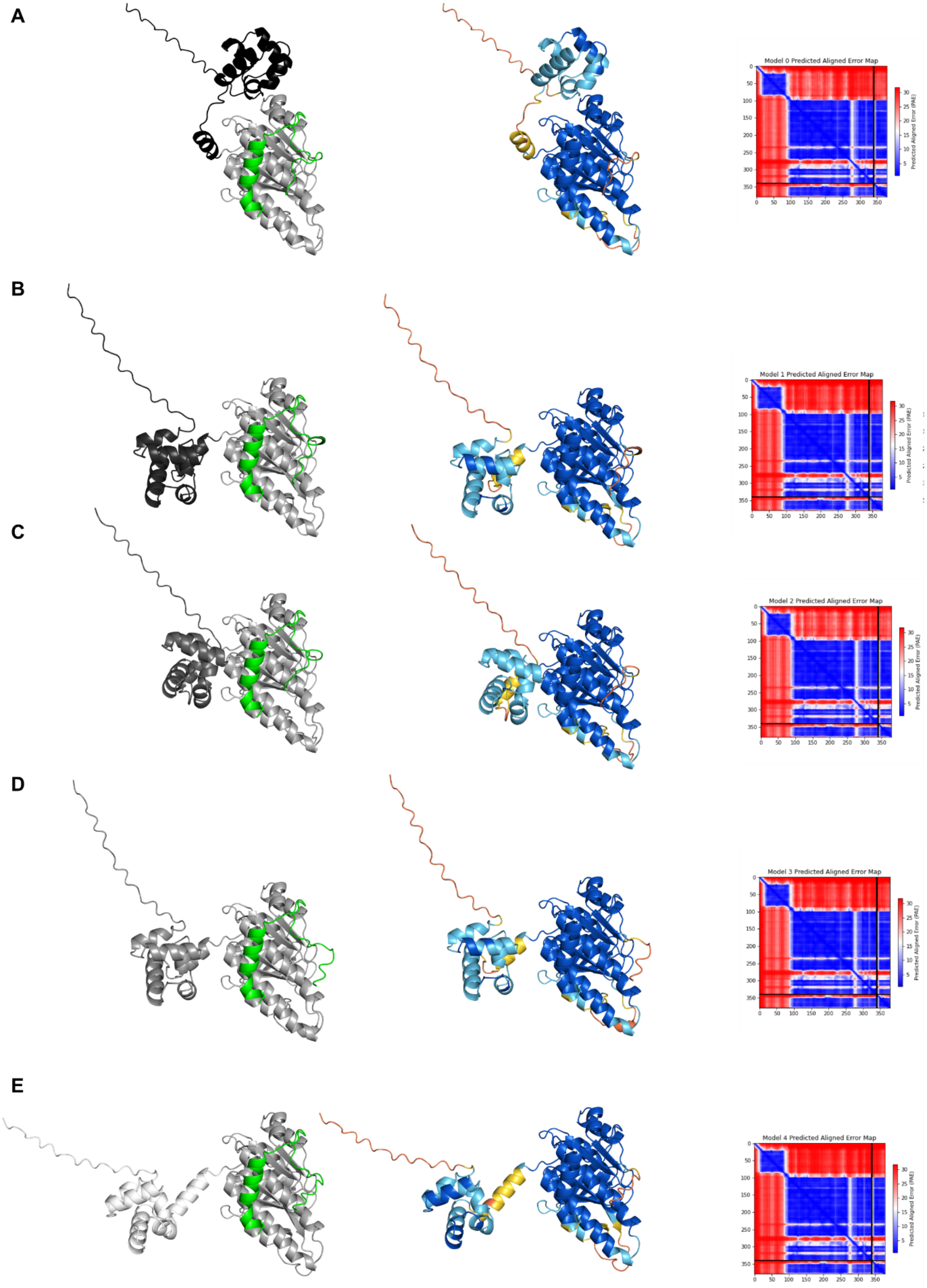


**Supplementary Figure 78** Left: the BRC2 peptide is colored green while the N-Terminal domain is displayed in different shades of grey depending on the model. Center: Residues have been colored for local model confidence estimated with pLDDT. Right: PAE of the corresponding model. **A.** Model 0 **B.** Model 1 **C.** Model 2 **D.** Model 3 **E.** Model 4

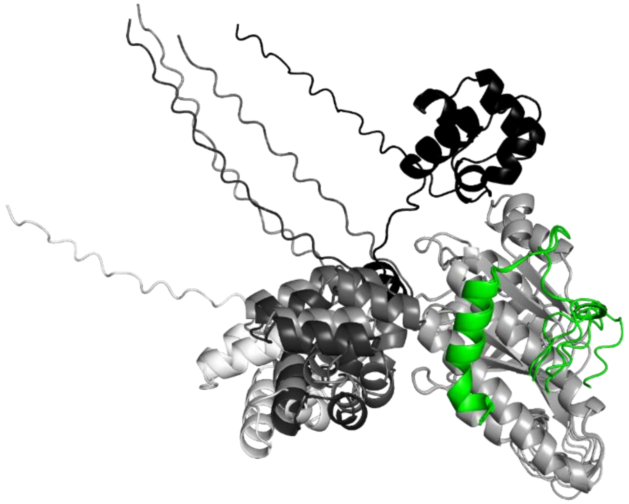

**Supplementary Figure 79 Overlay of the five best ranked AlphaFold3 models of the RAD51-FL/BRC2.**

| XL-Pairs | Distance | | | | |
|---|---|---|---|---|---|
| 58-1226 | 14.83 | 43.97 | 38.67 | 41.47 | 46.88 |
| 70-1226 | 25.71 | 50.27 | 41.92 | 48.88 | 56.66 |
| 73-1226 | 26.59 | 49.54 | 42.07 | 49.22 | 56.67 |
| 58-1236 | 23.93 | 23.27 | 18.01 | 21.08 | 26.44 |
| 64-1236 | 35.67 | 26.92 | 17.76 | 22.79 | 31.59 |
| 73-1236 | 31 | 29.48 | 23.14 | 29.68 | 37.24 |
| AF_Model | Model_0 | Model_1 | Model_2 | Model_3 | Model_4 |

**Supplementary Table 4** Table of detected inter-protein crosslinks and their measured Cα-Cα distances in the AlphaFold3 predictions of the RAD51-FL/BRC2 complex.

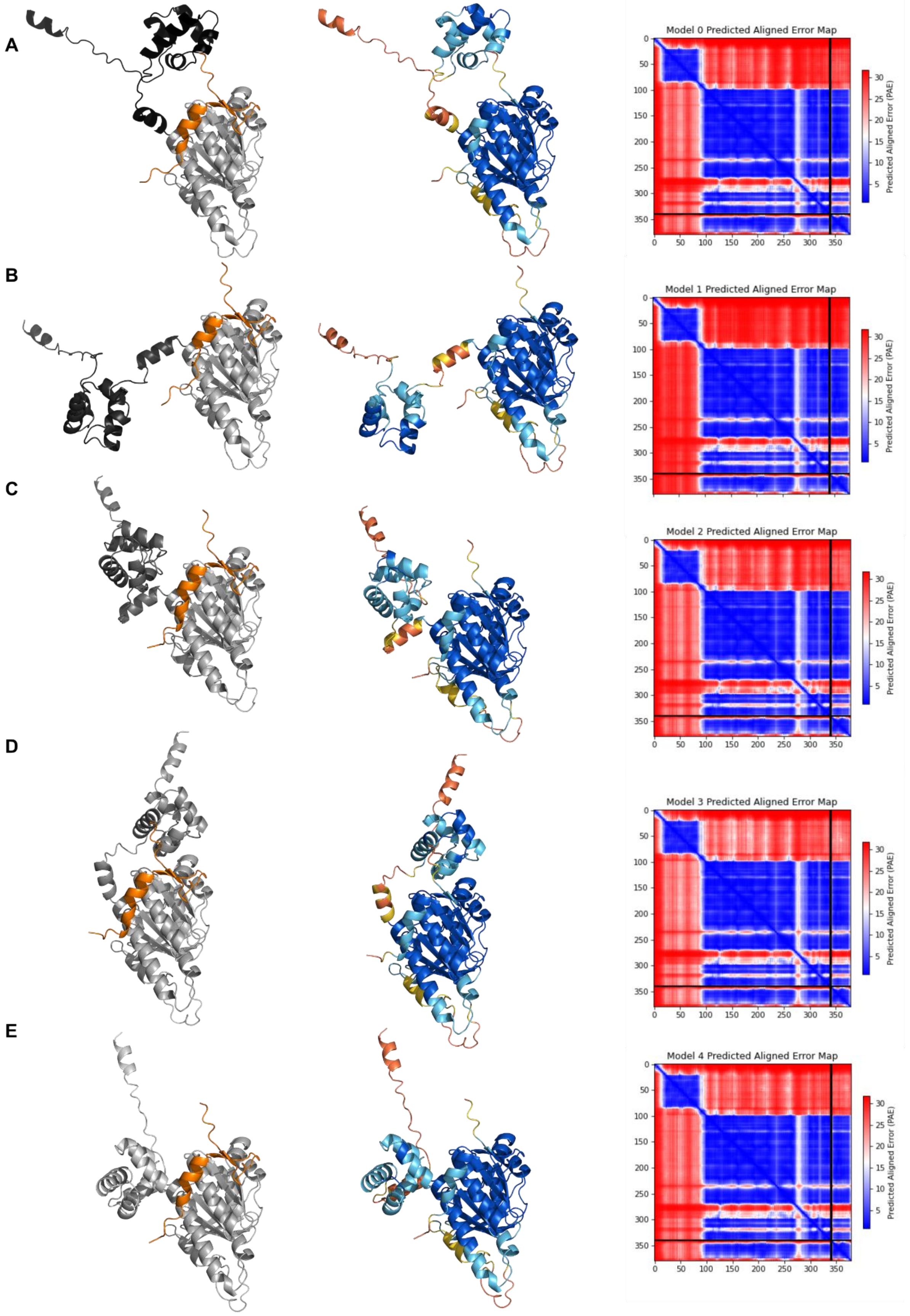


**Supplementary Figure 80** Left: the BRC3 peptide is colored orange while the N-Terminal domain is displayed in different shades of grey depending on the model. Center: Residues have been colored for local model confidence estimated with pLDDT. Right: PAE of the corresponding model. **A.** Model 0 **B.** Model 1 **C.** Model 2 **D.** Model 3 **E.** Model 4

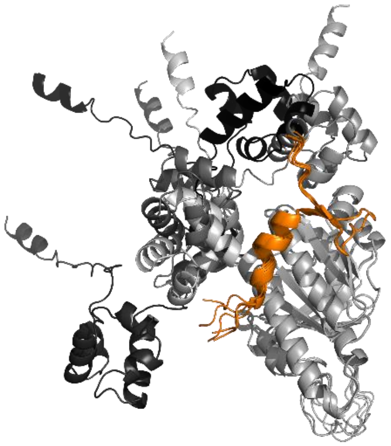

**Supplementary Figure 81 Overlay of the five best ranked AlphaFold3 models of the RAD51-FL/BRC3.**

| XL-Pairs | Distance | | | | |
|---|---|---|---|---|---|
| 58-1440 | 21.07 | 39.73 | 20.22 | 12.3 | 29.16 |
| AF_Model | Model_0 | Model_1 | Model_2 | Model_3 | Model_4 |

**Supplementary Table 5** Table of detected inter-protein crosslinks and their measured Cα-Cα distances in the AlphaFold3 predictions of the RAD51-FL/BRC3 complex.

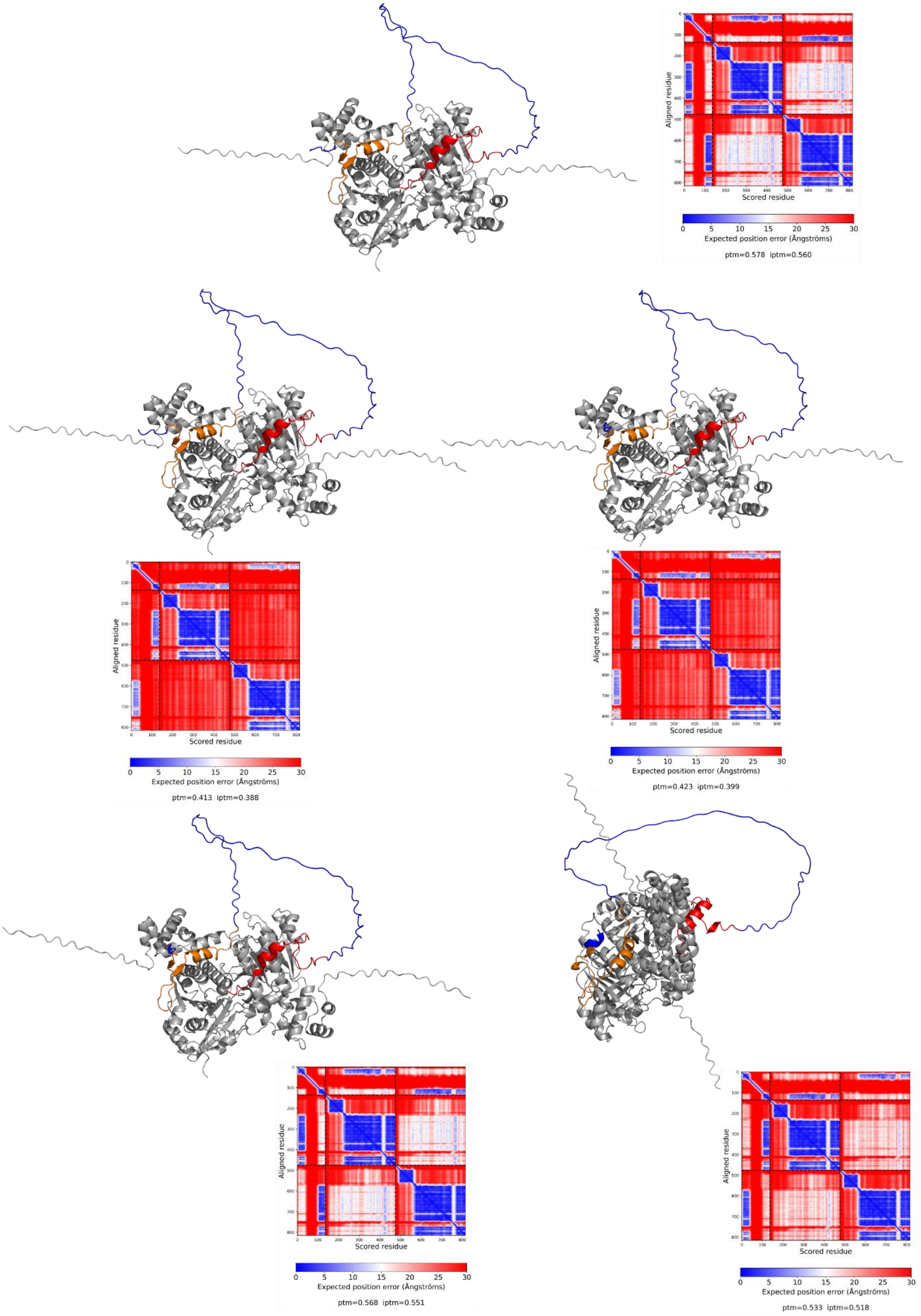


**Supplementary Figure 82 The five best ranked AlphaFold2 models of the RAD51-FL/BRC3-4 complex. Near each model is reported the corresponding PAE matrix.**

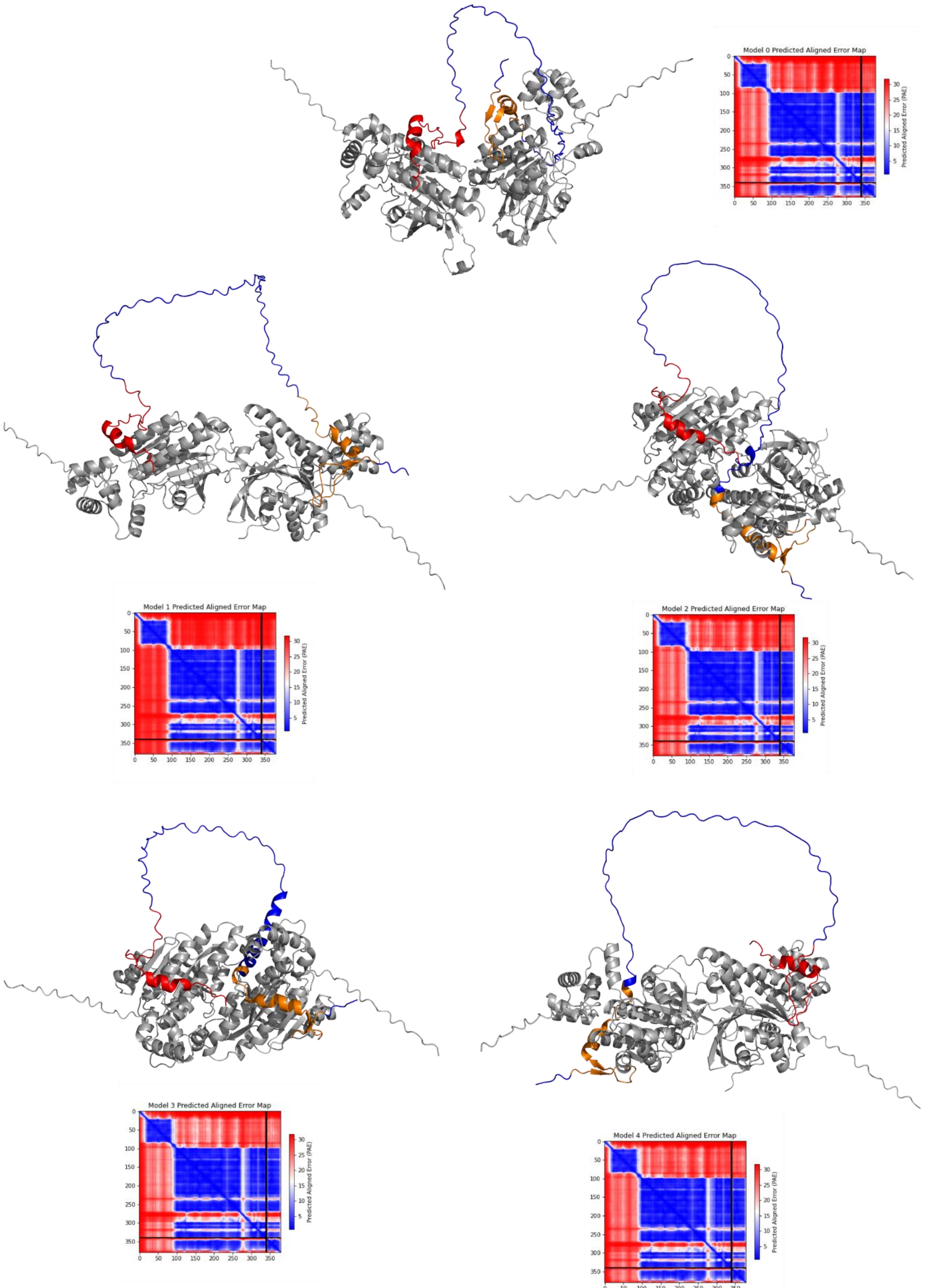


**Supplementary Figure 83 The five best ranked AlphaFold3 models of the RAD51-FL/BRC3-4 complex. Near each model is reported the corresponding PAE matrix.**

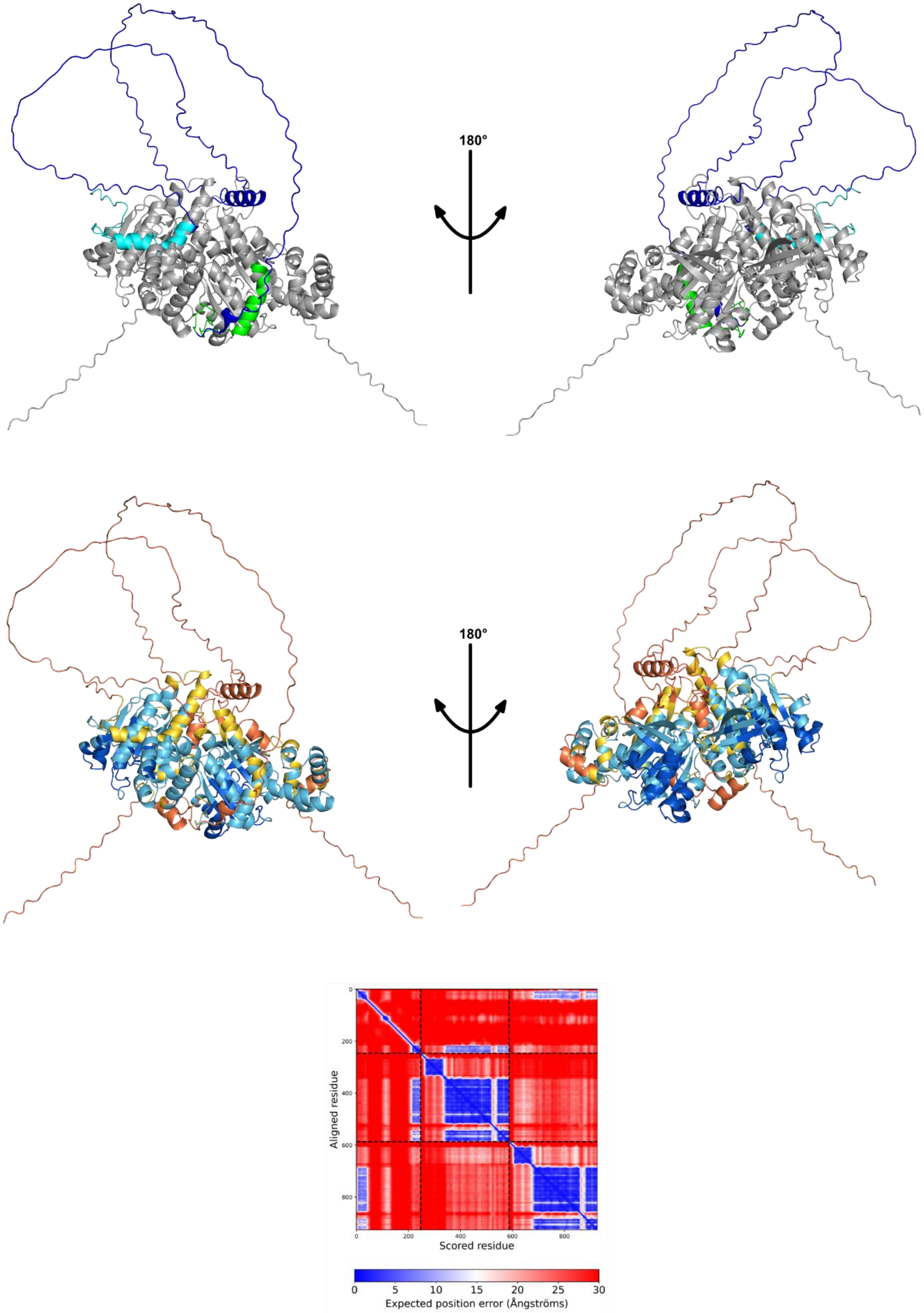


**Supplementary Figure 84 The AlphaFold model of the RAD51-FL/BRC1-2 complex.** Top: The predicted structure of BRC1-2 in complex with two RAD51-FL monomers. BRC1 is colored cyan while BRC2 is green. The two RAD51-FL monomers are highlighted in grey Center: The predicted structure of BRC1-2 in complex with two RAD51-FL monomers. Residues have been colored for local model confidence estimated with pLDDT. Bottom: PAE matrix.

| XL-Pairs | Distance |
|---|---|
| 40-1015 | 33.58 |
| 58-1015 | 40.41 |
| 73-1015 | 43.43 |
| 156-1015 | 14.61 |
| 73-1018 | 35.62 |
| 156-1018 | 15.61 |
| 58-1025 | 27.12 |
| 58-1028 | 17.39 |
| 58-1226 | 27.31 |
| 70-1226 | 27.3 |
| 73-1226 | 22.77 |
| 58-1236 | 23.45 |
| 64-1236 | 19.86 |
| 73-1236 | 23.86 |

**Supplementary Table 6** Table of detected inter-protein crosslinks and their measured Cα-Cα distances in the AlphaFold predictions of the RAD51-FL/BRC1-2 complex.

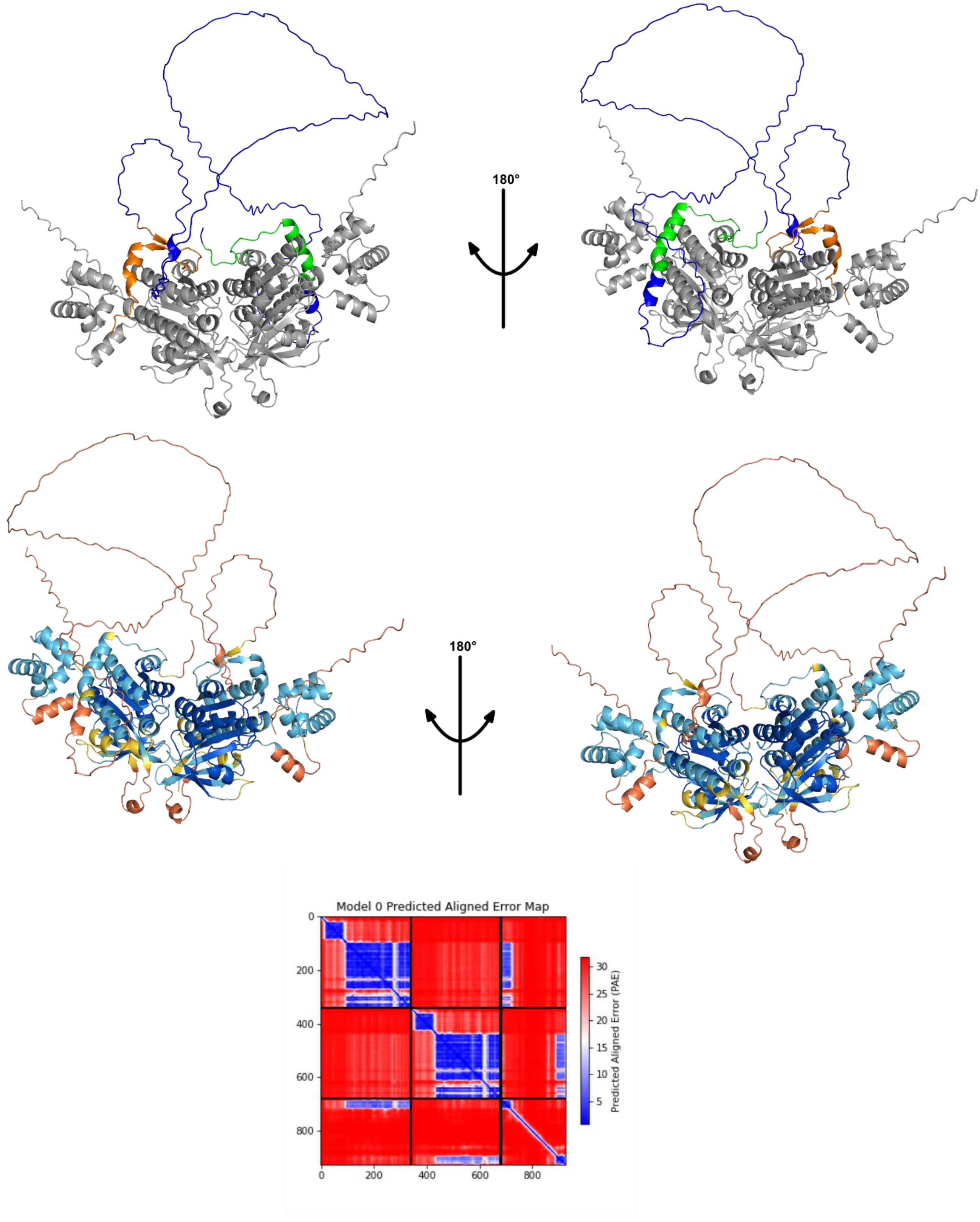


**Supplementary Figure 85 The AlphaFold model of the RAD51-FL/BRC2-3 complex.** Top: The predicted structure of BRC2-3 in complex with two RAD51-FL monomers. BRC2 is colored green while BRC3 is orange. The two RAD51-FL monomers are highlighted in grey. Center: The predicted structure of BRC2-3 in complex with two RAD51-FL monomers. Residues have been colored for local model confidence estimated with pLDDT. Bottom: PAE matrix.

| XL-Pairs | Distance |
| --- | --- |
| 58-1226 | 40.57 |
| 70-1226 | 41.96 |
| 73-1226 | 42.05 |
| 58-1236 | 21.49 |
| 64-1236 | 17.23 |
| 73-1236 | 23.67 |
| 58-1440 | 29.69 |

**Supplementary Table 7** Table of detected inter-protein crosslinks and their measured Cα-Cα distances in the AlphaFold predictions of the RAD51-FL/BRC2-3 complex.

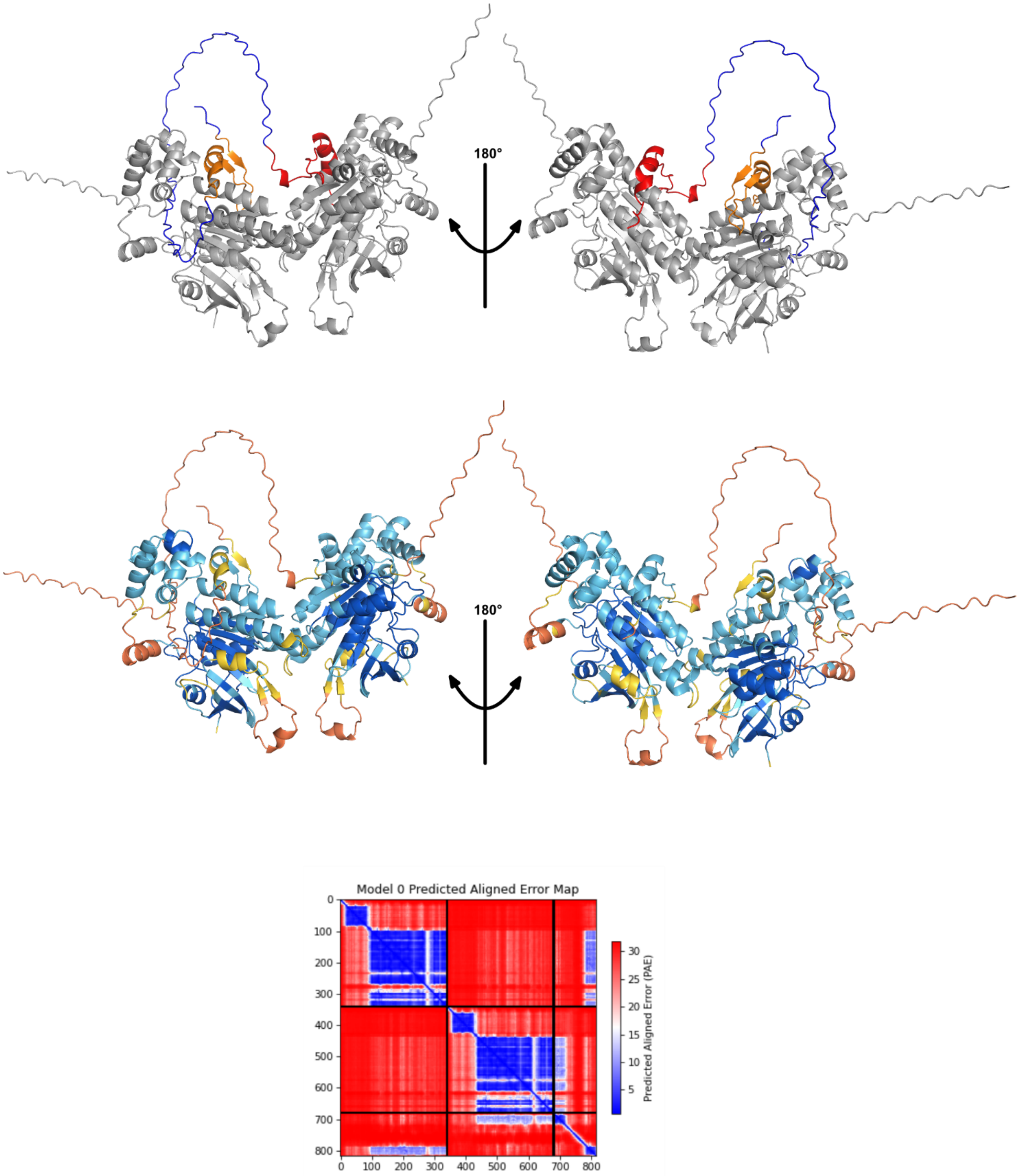


**Supplementary Figure 86 The AlphaFold model of the RAD51-FL/BRC3-4 complex.** Top: The predicted structure of BRC3-4 in complex with two RAD51-FL monomers. BRC3 is colored orange while BRC4 is red. The two RAD51-FL monomers are highlighted in grey. Center: The predicted structure of BRC3-4 in complex with two RAD51-FL monomers. Residues have been colored for local model confidence estimated with pLDDT. Bottom: PAE matrix.

| XL-Pairs | Distance |
|---|---|
| 58-1440 | 28.63 |
| 58-1536 | 29.04 |
| 73-1536 | 29.8 |
| 40-1541 | 9.3 |
| 58-1541 | 20.46 |
| 64-1541 | 14.92 |
| 73-1541 | 22.12 |

**Supplementary Table 8** Table of detected inter-protein crosslinks and their measured Cα-Cα distances in the AlphaFold predictions of the RAD51-FL/BRC3-4 complex.

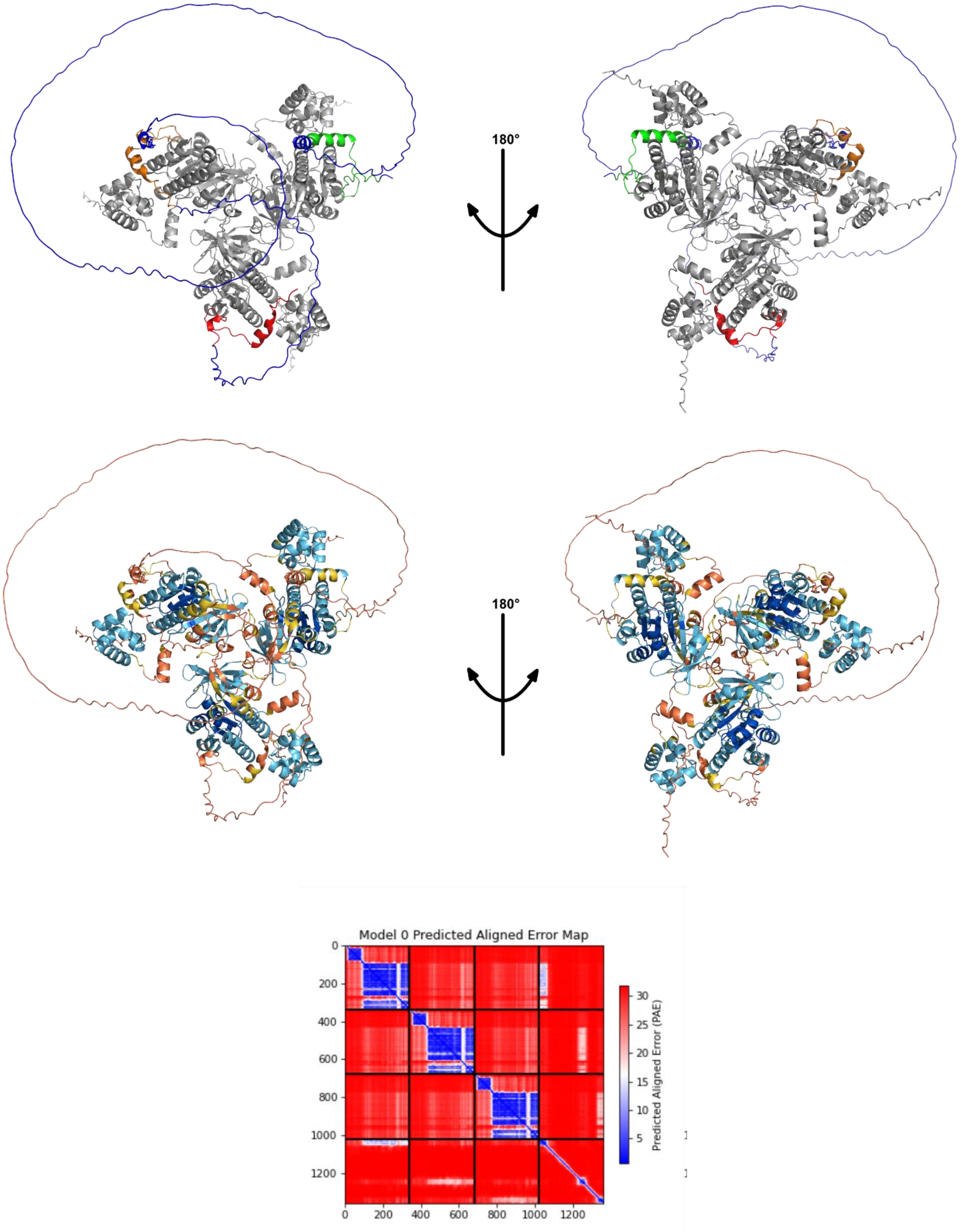


**Supplementary Figure 87 The AlphaFold model of the RAD51-FL/BRC2-4 complex.** Top: The predicted structure of BRC2-4 in complex with three RAD51-FL monomers. BRC2 is highlighted in green, BRC3 is colored orange and BRC4 is red. The three RAD51-FL monomers are highlighted in grey. Center: The predicted structure of BRC2-4 in complex with three RAD51-FL monomers. Residues have been colored for local model confidence estimated with pLDDT. Bottom: PAE matrix.

| XL-Pairs | Distance |
|---|---|
| 58-1226 | 37.09 |
| 70-1226 | 39.92 |
| 73-1226 | 41.28 |
| 58-1236 | 17.55 |
| 64-1236 | 13.31 |
| 73-1236 | 21.33 |
| 58-1440 | 26.03 |
| 58-1536 | 26.08 |
| 73-1536 | 28.89 |
| 40-1541 | 8.35 |
| 58-1541 | 17.92 |
| 64-1541 | 13.02 |
| 73-1541 | 20.86 |

**Supplementary Table 9** Table of detected inter-protein crosslinks and their measured Cα-Cα distances in the AlphaFold predictions of the RAD51-FL/BRC2-4 complex.

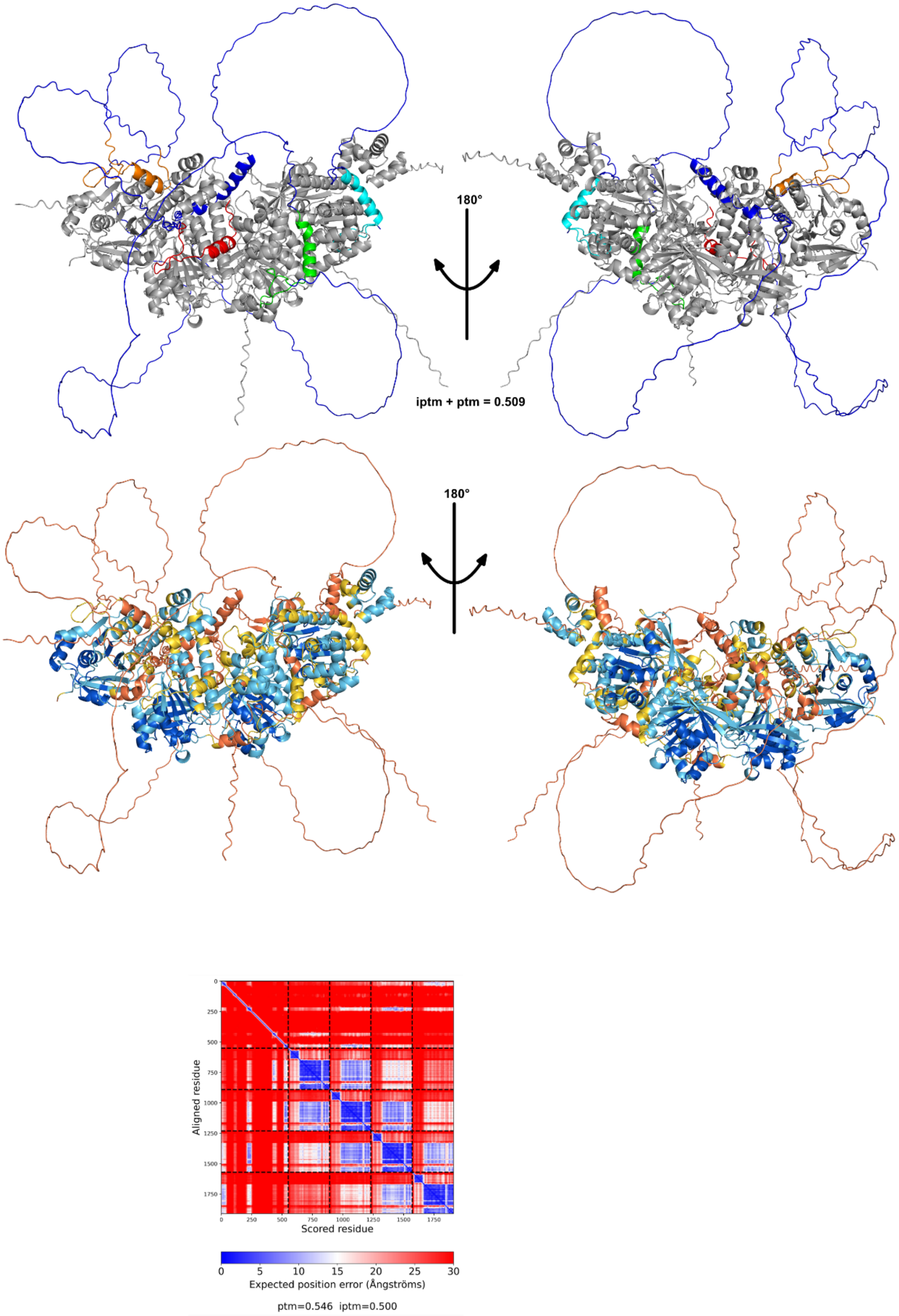


**Supplementary Figure 88 The AlphaFold model of the RAD51-FL/BRC1-4 complex.** Top: The predicted structure of BRC1-4 in complex with four RAD51-FL monomers. BRC1 is cyan, BRC2 is highlighted in green, BRC3 is colored orange and BRC4 is red. The four RAD51-FL monomers are highlighted in grey. Center: The predicted structure of BRC2-4 in complex with four RAD51-FL monomers. Residues have been colored for local model confidence estimated with pLDDT. Bottom: PAE matrix.

| XL-Pairs | Distance |
|---|---|
| 40-1015 | 34.83 |
| 58-1015 | 32.86 |
| 73-1015 | 29.42 |
| 156-1015 | 14.16 |
| 73-1018 | 23.96 |
| 156-1018 | 15.63 |
| 58-1025 | 23.24 |
| 58-1028 | 19.48 |
| 58-1226 | 29.49 |
| 70-1226 | 30.07 |
| 73-1226 | 25.6 |
| 58-1236 | 21.12 |
| 64-1236 | 22.3 |
| 73-1236 | 21.53 |
| 58-1440 | 30.86 |
| 58-1536 | 27.15 |
| 73-1536 | 20.26 |
| 40-1541 | 14.87 |
| 58-1541 | 20.91 |
| 64-1541 | 20.72 |
| 73-1541 | 20.89 |

**Supplementary Table 10** Table of detected inter-protein crosslinks and their measured Cα-Cα distances in the AlphaFold predictions of the RAD51-FL/BRC1-4 complex.

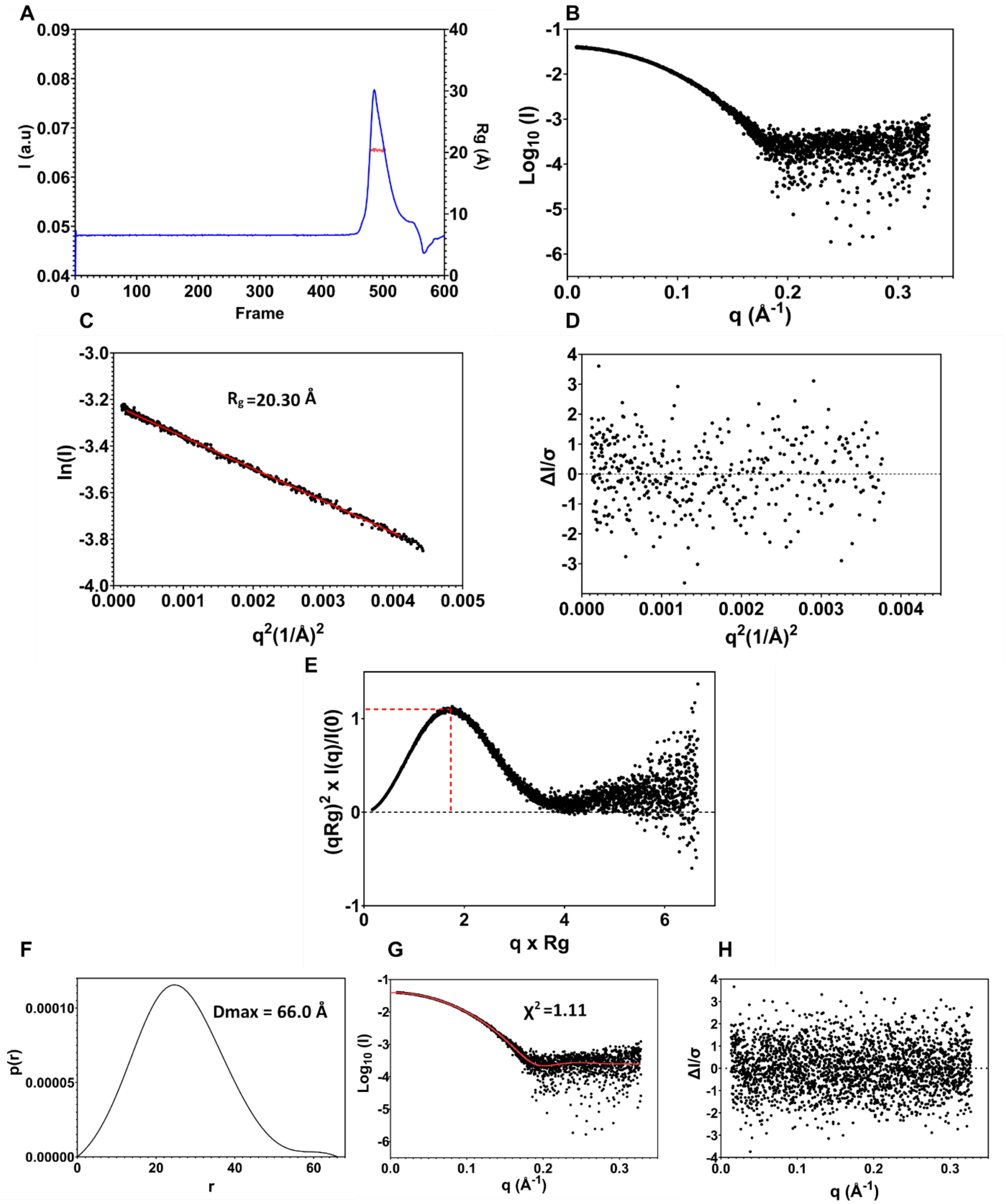


**Supplementary Figure 89 SEC-SAXS analysis of Δ97-RAD51/BRC4 complex A.** SEC - SAXS profile of RAD51 - BRC4 complex. The red line estimates the radius of gyration calculated by the selected sample frames through CHROMIXS. **B.** Scatter plot (logarithmic scale) **C.** Guinier analysis of SAXS Profile. **D.** Residuals of Guinier fitting **E.** Dimensionless Kratky plot. Red crosshairs designate the Guinier-Kratky point. The Guinier-Kratky plot curve peak coincides with this specific point when the protein is behaving as a compact and folded system **F.** SAXS p(r) analysis. Left: p(r) plot and calculated $D_{max}$. Middle: p(r) fitting to experimental data. Right: residuals of p(r) fitting.

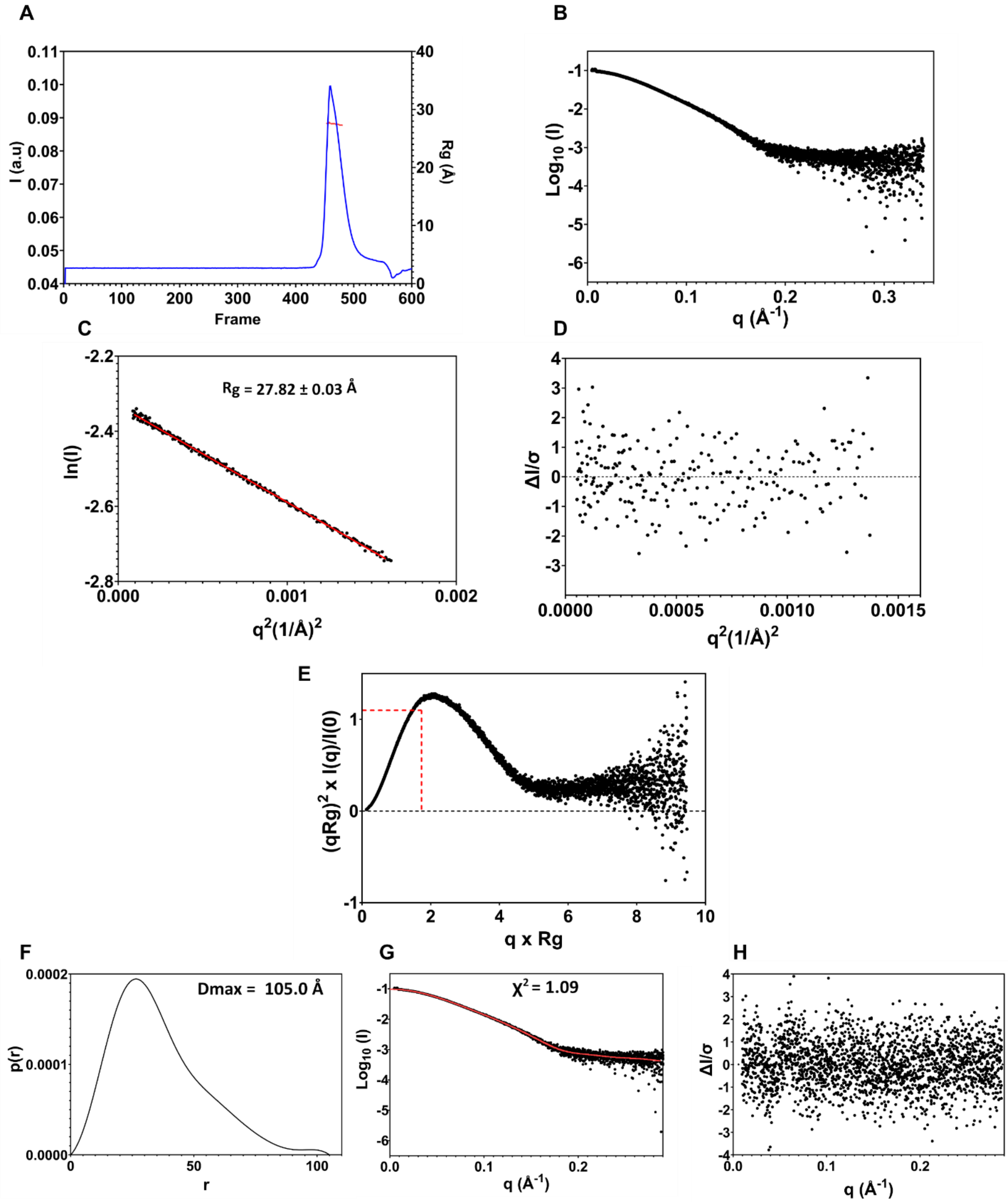


**Supplementary Figure 90 SEC-SAXS analysis of RAD51/BRC4 complex A.** SEC - SAXS profile of RAD51 - BRC4 complex. The red line estimates the radius of gyration calculated by the selected sample frames through CHROMIXS. **B.** Scatter plot (logarithmic scale) **C.** Guinier analysis of SAXS Profile. **D.** Residuals of Guinier fitting **E.** Dimensionless Kratky plot. Red crosshairs designate the Guinier-Kratky point. The Guinier-Kratky plot curve peak coincides with this specific point when the protein is behaving as a compact and folded system **F.** SAXS p(r) analysis. Left: p(r) plot and calculated $D_{max}$. Middle: p(r) fitting to experimental data. Right: residuals of p(r) fitting.

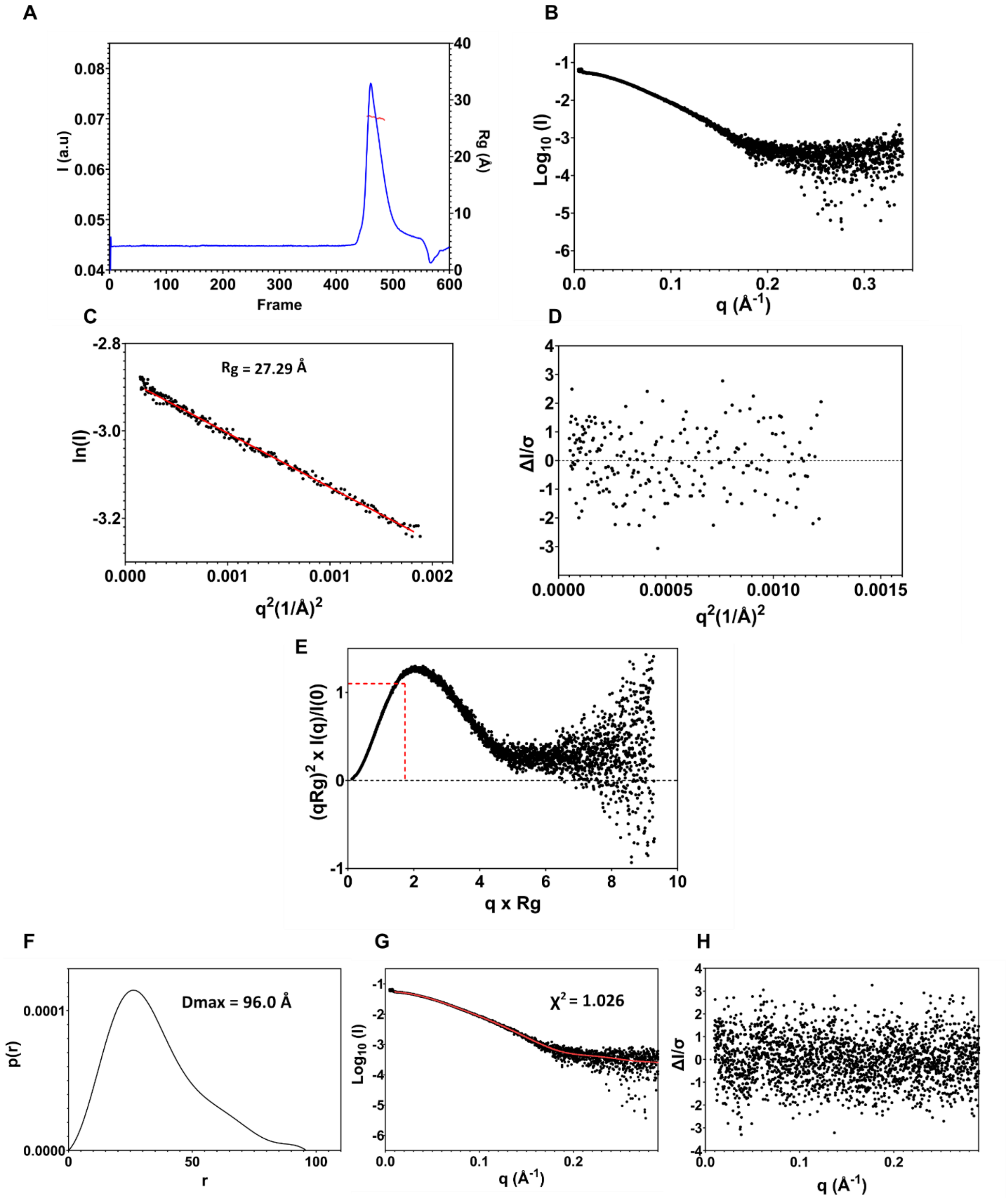


**Supplementary Figure 91 SEC-SAXS analysis of RAD51/BRC2 complex A.** SEC - SAXS profile of RAD51 – BRC2 complex. The red line estimates the radius of gyration calculated by the selected sample frames through CHROMIXS. **B.** Scatter plot (logarithmic scale) **C.** Guinier analysis of SAXS Profile. **D.** Residuals of Guinier fitting **E.** Dimensionless Kratky plot. Red crosshairs designate the Guinier-Kratky point. The Guinier-Kratky plot curve peak coincides with this specific point when the protein is behaving as a compact and folded system **F.** SAXS p(r) analysis. Left: p(r) plot and calculated $D_{max}$. Middle: p(r) fitting to experimental data. Right: residuals of p(r) fitting.

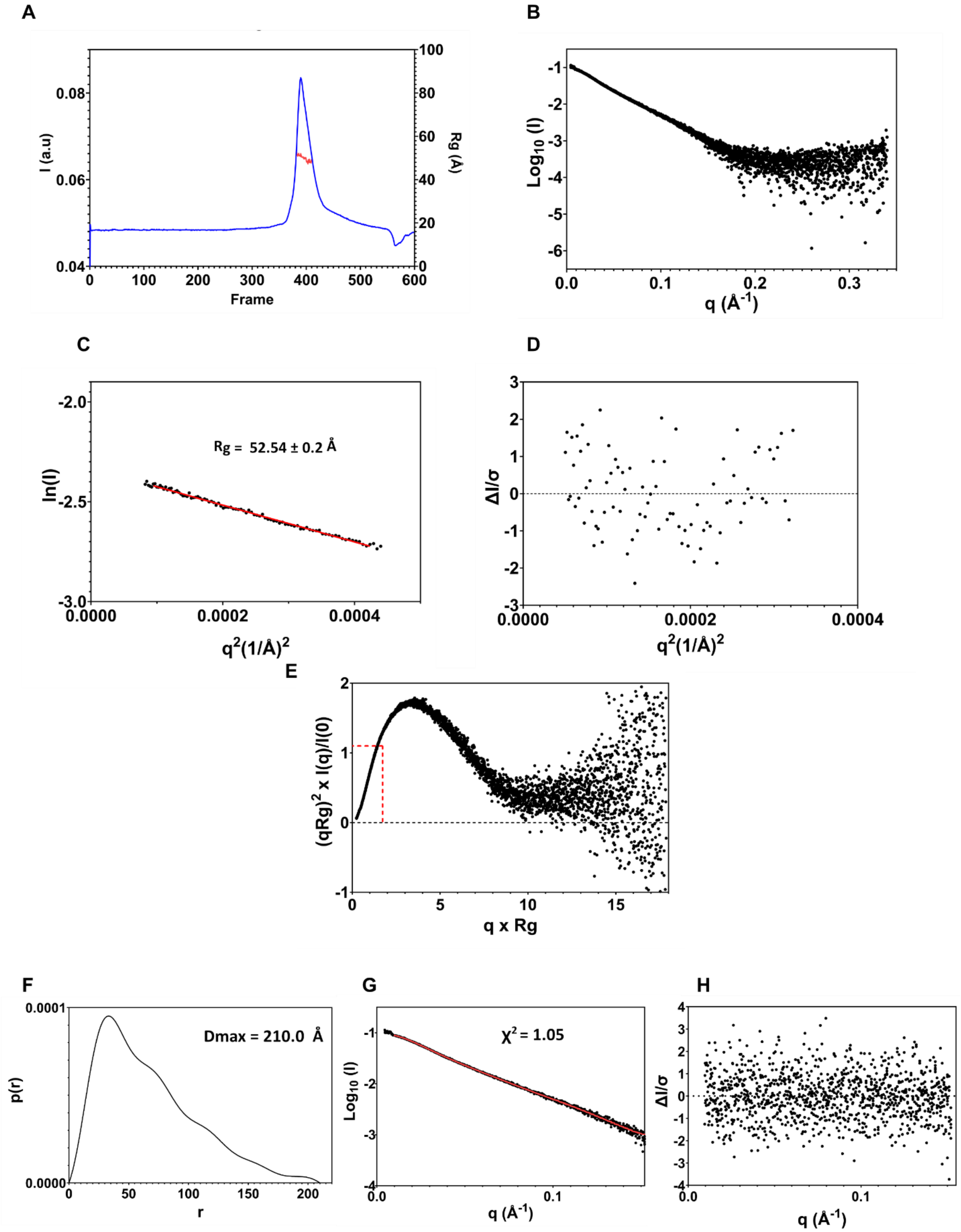


**Supplementary Figure 92 SEC-SAXS analysis of RAD51/BRC1-2 complex A.** SEC - SAXS profile of RAD51 – BRC1-2 complex. The red line estimates the radius of gyration calculated by the selected sample frames through CHROMIXS. **B.** Scatter plot (logarithmic scale) **C.** Guinier analysis of SAXS Profile. **D.** Residuals of Guinier fitting **E.** Dimensionless Kratky plot. Red crosshairs designate the Guinier-Kratky point. The Guinier-Kratky plot curve peak coincides with this specific point when the protein is behaving as a compact and folded system **F.** SAXS p(r) analysis. Left: p(r) plot and calculated $D_{max}$. Middle: p(r) fitting to experimental data. Right: residuals of p(r) fitting.

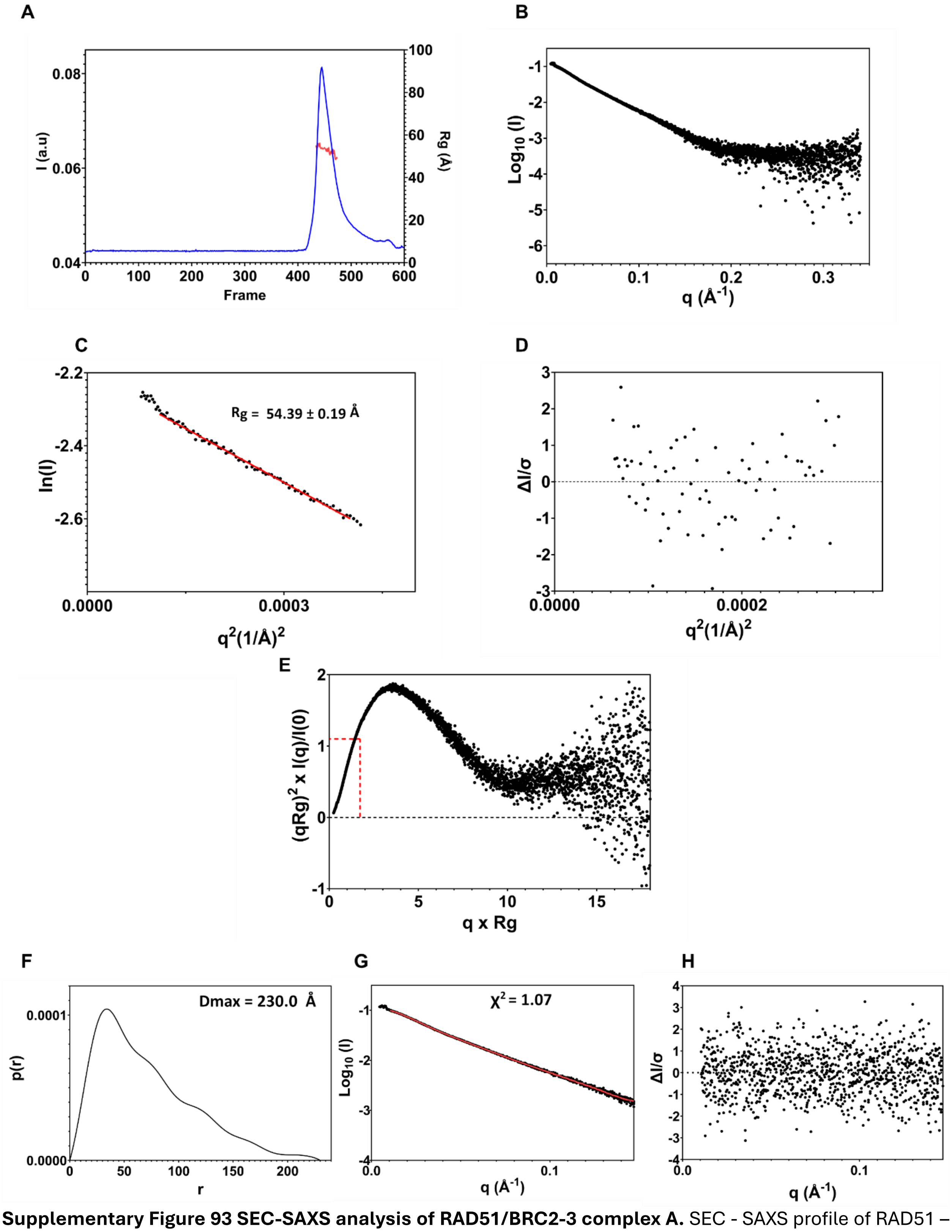


**Supplementary Figure 93 SEC-SAXS analysis of RAD51/BRC2-3 complex A.** SEC - SAXS profile of RAD51 – BRC2-3 complex. The red line estimates the radius of gyration calculated by the selected sample frames through CHROMIXS. **B.** Scatter plot (logarithmic scale) **C.** Guinier analysis of SAXS Profile. **D.** Residuals of Guinier fitting **E.** Dimensionless Kratky plot. Red crosshairs designate the Guinier-Kratky point. The Guinier-Kratky plot curve peak coincides with this specific point when the protein is behaving as a compact and folded system **F.** SAXS p(r) analysis. Left: p(r) plot and calculated $D_{max}$. Middle: p(r) fitting to experimental data. Right: residuals of p(r) fitting.

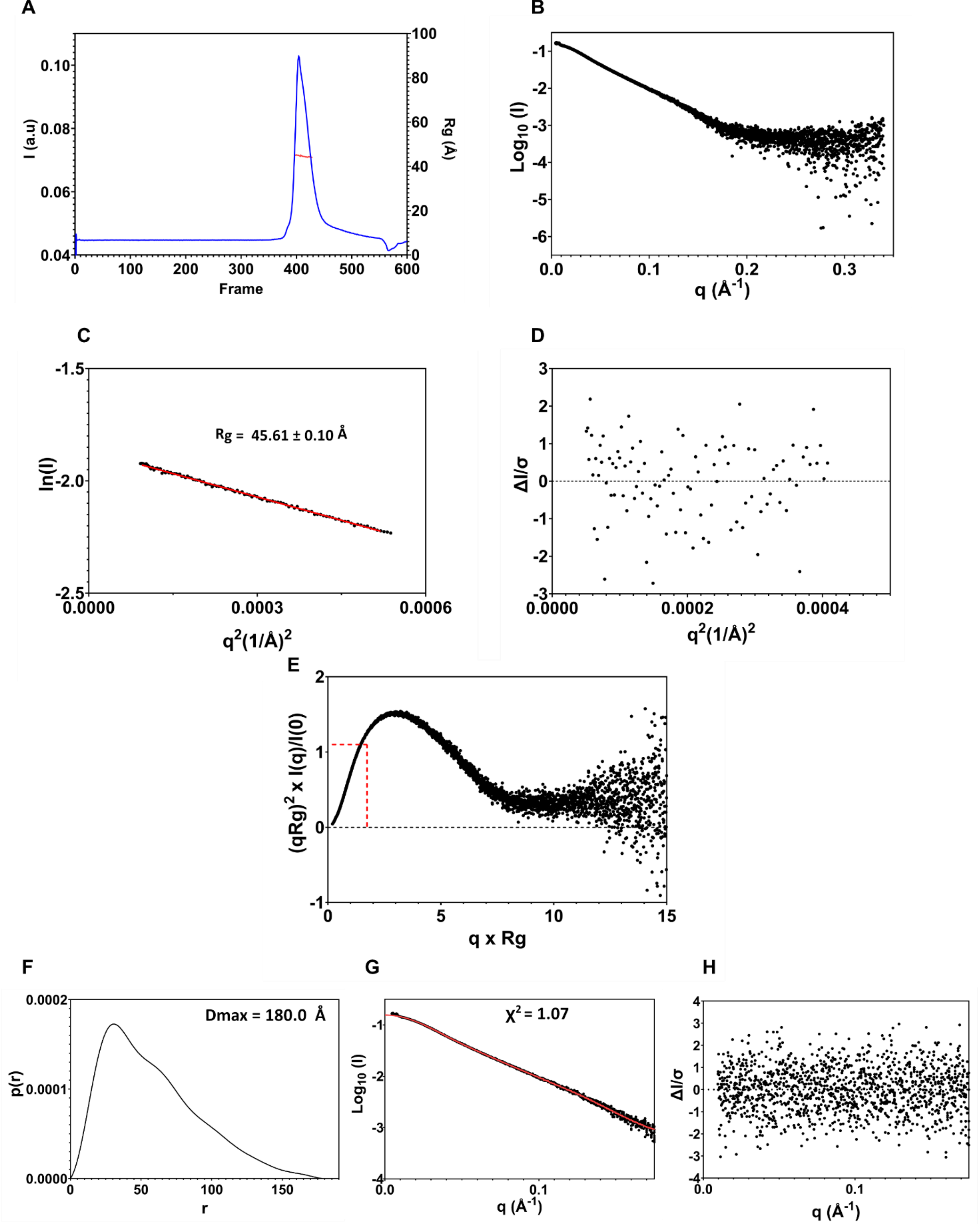


**Supplementary Figure 94 SEC-SAXS analysis of RAD51/BRC3-4 complex A.** SEC - SAXS profile of RAD51 – BRC3-4 complex. The red line estimates the radius of gyration calculated by the selected sample frames through CHROMIXS. **B.** Scatter plot (logarithmic scale) **C.** Guinier analysis of SAXS Profile. **D.** Residuals of Guinier fitting **E.** Dimensionless Kratky plot. Red crosshairs designate the Guinier-Kratky point. The Guinier-Kratky plot curve peak coincides with this specific point when the protein is behaving as a compact and folded system **F.** SAXS p(r) analysis. Left: p(r) plot and calculated $D_{max}$. Middle: p(r) fitting to experimental data. Right: residuals of p(r) fitting.

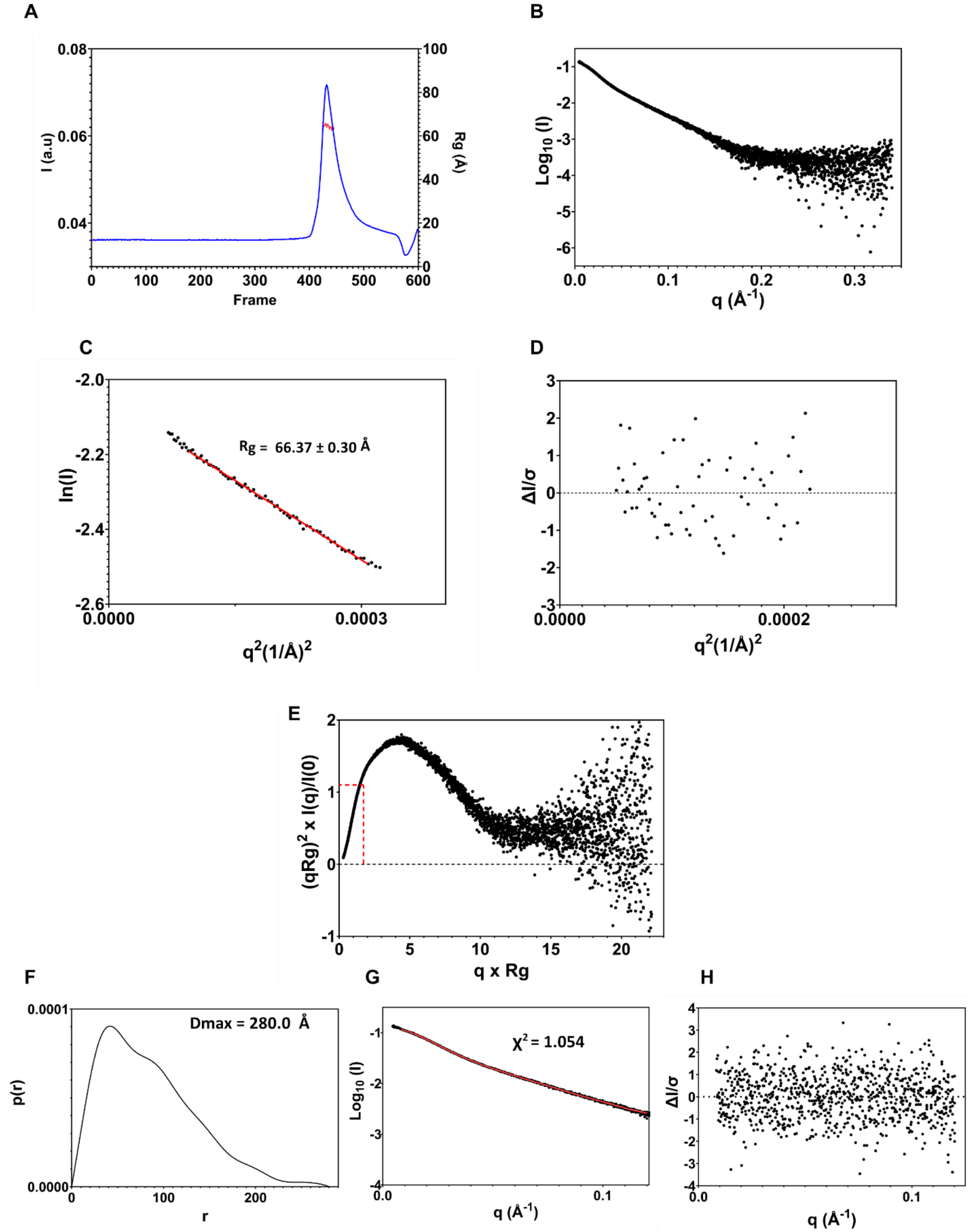


**Supplementary Figure 95 SEC-SAXS analysis of RAD51/BRC2-4 complex A.** SEC - SAXS profile of RAD51 – BRC2-4 complex. The red line estimates the radius of gyration calculated by the selected sample frames through CHROMIXS. **B.** Scatter plot (logarithmic scale) **C.** Guinier analysis of SAXS Profile. **D.** Residuals of Guinier fitting **E.** Dimensionless Kratky plot. Red crosshairs designate the Guinier-Kratky point. The Guinier-Kratky plot curve peak coincides with this specific point when the protein is behaving as a compact and folded system **F.** SAXS p(r) analysis. Left: p(r) plot and calculated $D_{max}$. Middle: p(r) fitting to experimental data. Right: residuals of p(r) fitting.

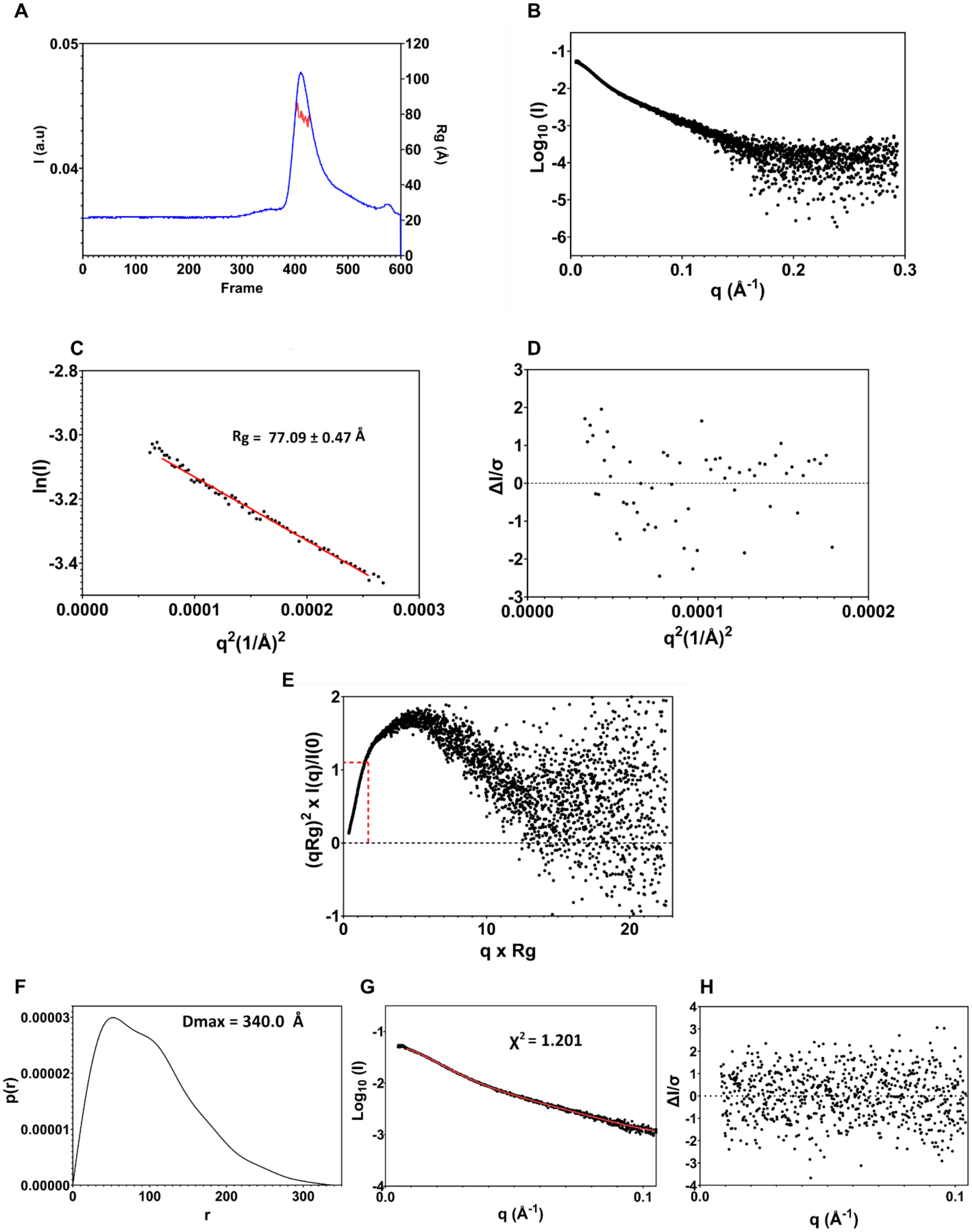


**Supplementary Figure 96 SEC-SAXS analysis of RAD51/BRC1-4 complex A.** SEC - SAXS profile of RAD51 – BRC1-4 complex. The red line estimates the radius of gyration calculated by the selected sample frames through CHROMIXS. **B.** Scatter plot (logarithmic scale) **C.** Guinier analysis of SAXS Profile. **D.** Residuals of Guinier fitting **E.** Dimensionless Kratky plot. Red crosshairs designate the Guinier-Kratky point. The Guinier-Kratky plot curve peak coincides with this specific point when the protein is behaving as a compact and folded system **F.** SAXS p(r) analysis. Left: p(r) plot and calculated $D_{max}$. Middle: p(r) fitting to experimental data. Right: residuals of p(r) fitting.

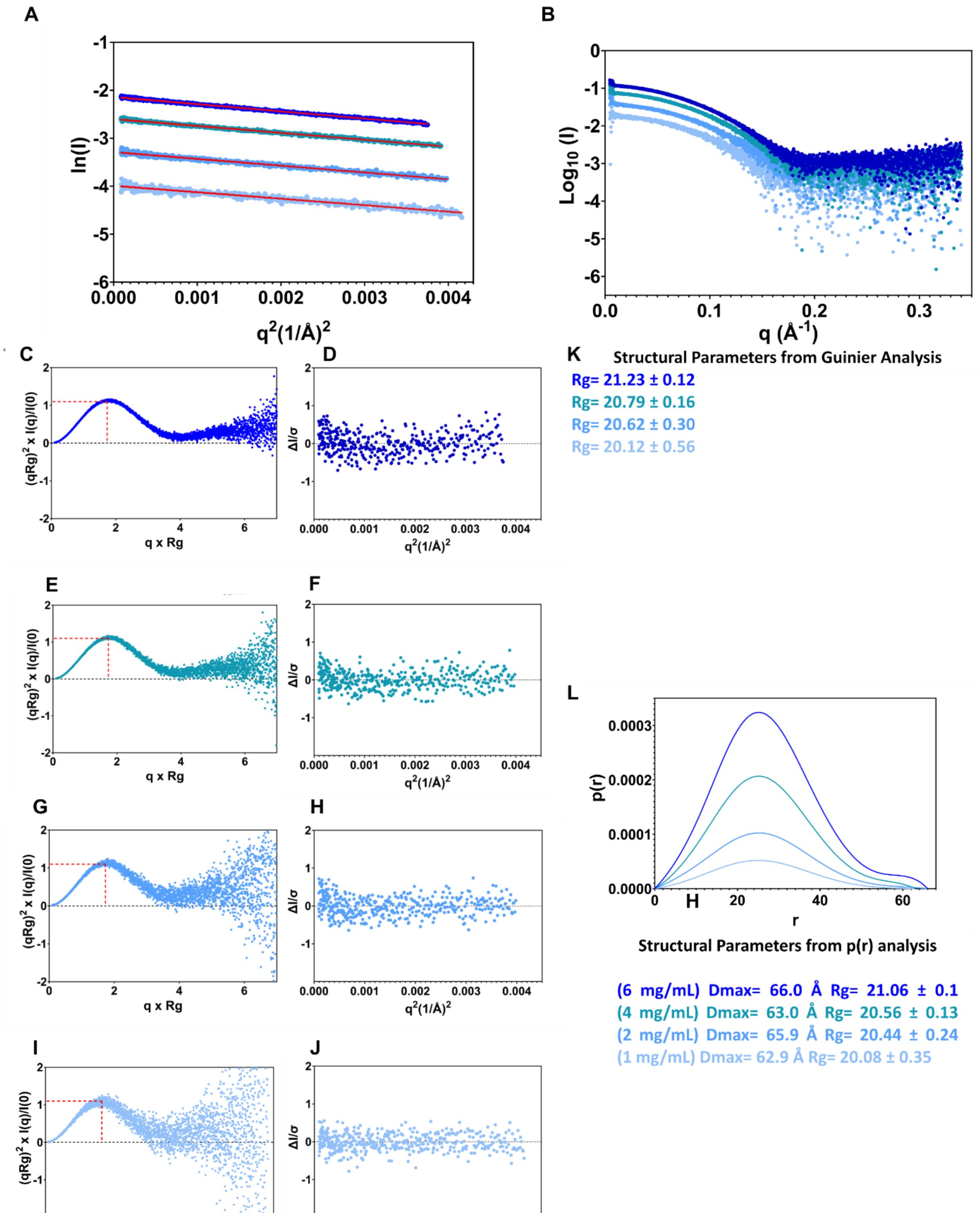


**Supplementary Figure 97 Batch-SAXS analysis of Δ97-RAD51/BRC4 complex A.** Guinier analysis of SAXS Profile at different Δ97-RAD51/BRC4 concentrations **B.** Scatter plot (logarithmic scale) of Δ97-RAD51/BRC4 at different concentrations **C.** Dimensionless Kratky plot at [Δ97-RAD51/BRC4] = 6 mg/mL **D.** Residuals of Guinier fitting at [Δ97-RAD51/BRC4] = 6 mg/mL **E.** Dimensionless Kratky plot at [Δ97-RAD51/BRC4] = 4 mg/mL **F.** Residuals of Guinier fitting at [Δ97-RAD51/BRC4] = 4 mg/mL **G.** Dimensionless Kratky plot at [Δ97-RAD51/BRC4] = 2 mg/mL **H.** Residuals of Guinier fitting at [Δ97-RAD51/BRC4] = 2 mg/mL **I.** Dimensionless Kratky plot at [Δ97-RAD51/BRC4] = 1 mg/mL **J.** Residuals of Guinier fitting [Δ97-RAD51/BRC4] = 1 mg/mL **K.** Rg inferred from Guinier analyses **L.** SAXS p(r) analysis at different concentrations. Bottom Dmax and Rg inferred from p(r) analysis at different Δ97-RAD51/BRC4 concentrations.

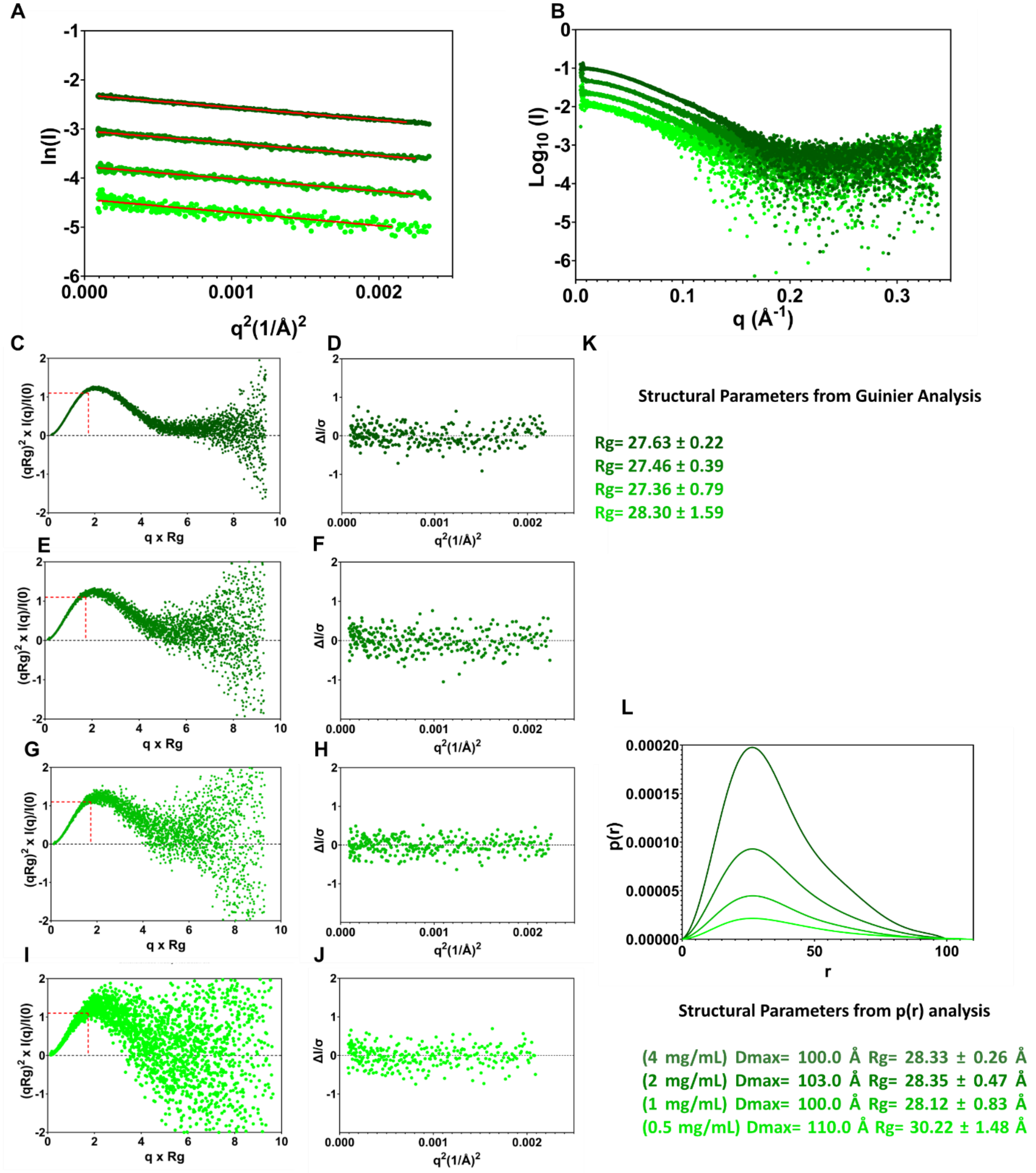


**Supplementary Figure 98 Batch-SAXS analysis of RAD51/BRC4 complex A.** Guinier analysis of SAXS Profile at different RAD51/BRC4 concentrations **B.** Scatter plot (logarithmic scale) of RAD51/BRC4 at different concentrations **C.** Dimensionless Kratky plot at [RAD51/BRC4] = 4 mg/mL **D.** Residuals of Guinier fitting at [RAD51/BRC4] = 4 mg/mL **E.** Dimensionless Kratky plot at [RAD51/BRC4] = 2 mg/mL **F.** Residuals of Guinier fitting at [RAD51/BRC4] = 1 mg/mL **G.** Dimensionless Kratky plot at [RAD51/BRC4] = 1 mg/mL **H.** Residuals of Guinier fitting at [RAD51/BRC4] = 1 mg/mL **I.** Dimensionless Kratky plot at [RAD51/BRC4] = 0.5 mg/mL **J.** Residuals of Guinier fitting [RAD51/BRC4] = 0.5 mg/mL **K.** Rg inferred from Guinier analyses **L.** SAXS p(r) analysis at different concentrations. Bottom Dmax and Rg inferred from p(r) analysis at different RAD51/BRC4 concentrations.

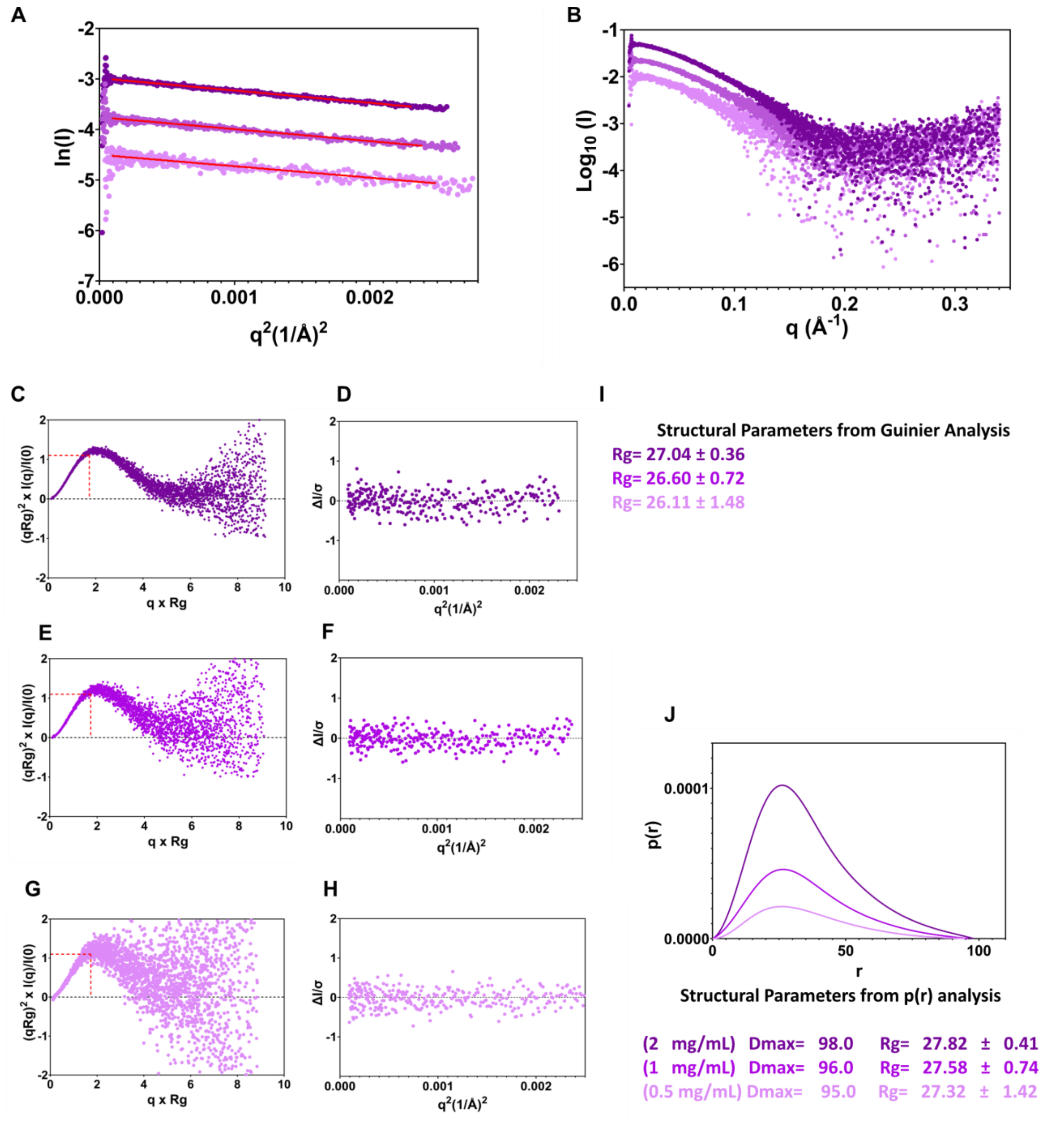


**Supplementary Figure 99 Batch-SAXS analysis of RAD51/BRC2 complex A.** Guinier analysis of SAXS Profile at different RAD51/BRC2 concentrations **B.** Scatter plot (logarithmic scale) of RAD51/BRC2 at different concentrations **C.** Dimensionless Kratky plot at [RAD51/BRC2] = 2 mg/mL **D.** Residuals of Guinier fitting at [RAD51/BRC2] = 2 mg/mL **E.** Dimensionless Kratky plot at [RAD51/BRC2] = 1 mg/mL **F.** Residuals of Guinier fitting at [RAD51/BRC2] = 1 mg/mL **G.** Dimensionless Kratky plot at [RAD51/BRC2] = 0.5 mg/mL **H.** Residuals of Guinier fitting at [RAD51/BRC2] = 0.5 mg/mL **I.** Rg inferred from Guinier analyses **J.** SAXS p(r) analysis at different concentrations. Bottom Dmax and Rg inferred from p(r) analysis at different RAD51/BRC2 concentrations.

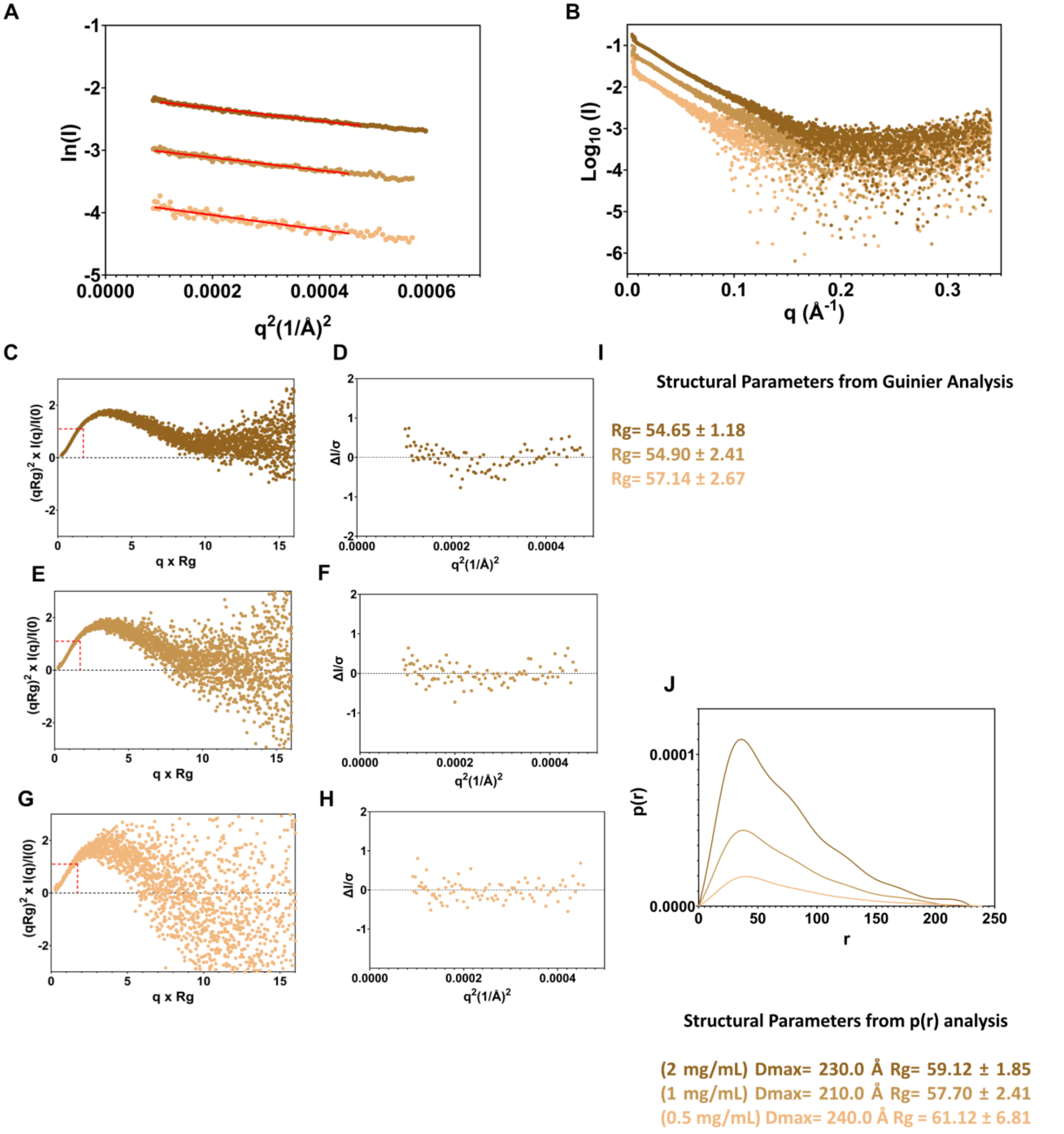


**Supplementary Figure 100 Batch-SAXS analysis of RAD51/BRC1-2 complex A.** Guinier analysis of SAXS Profile at different RAD51/BRC1-2 concentrations **B.** Scatter plot (logarithmic scale) of RAD51/BRC1-2 at different concentrations **C.** Dimensionless Kratky plot at [RAD51/BRC1-2] = 2 mg/mL **D.** Residuals of Guinier fitting at [RAD51/BRC1-2] = 2 mg/mL **E.** Dimensionless Kratky plot at [RAD51/BRC1-2] = 1 mg/mL **F.** Residuals of Guinier fitting at [RAD51/BRC1-2] = 1 mg/mL **G.** Dimensionless Kratky plot at [RAD51/BRC1-2] = 0.5 mg/mL **H.** Residuals of Guinier fitting at [RAD51/BRC1-2] = 0.5 mg/mL **I.** Rg inferred from Guinier analyses **J.** SAXS p(r) analysis at different concentrations. Bottom Dmax and Rg inferred from p(r) analysis at different RAD51/BRC1-2 concentrations.

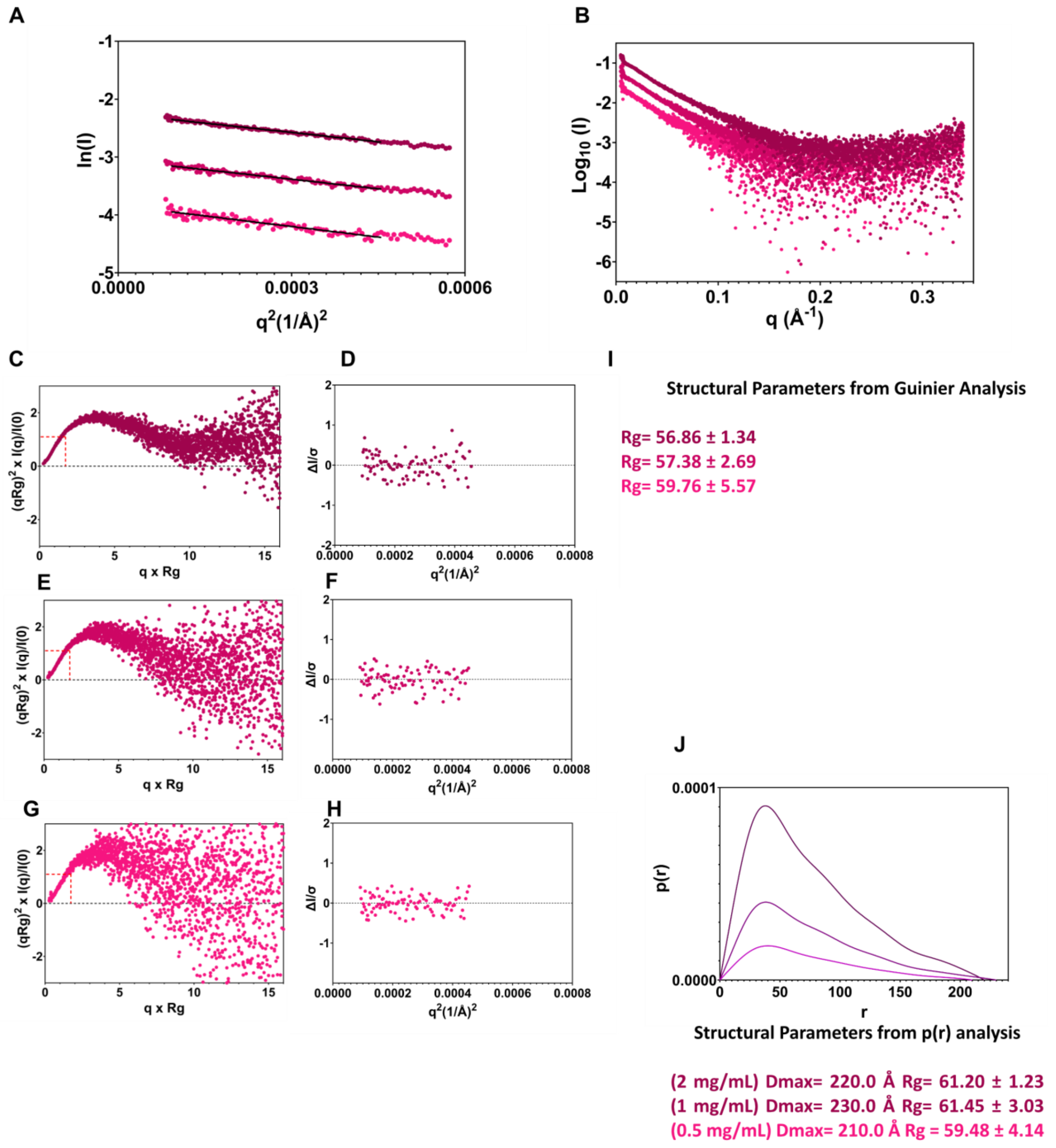


**Supplementary Figure 101 Batch-SAXS analysis of RAD51/BRC2-3 complex A.** Guinier analysis of SAXS Profile at different RAD51/BRC2-3 concentrations **B.** Scatter plot (logarithmic scale) of RAD51/BRC2-3 at different concentrations **C.** Dimensionless Kratky plot at [RAD51/BRC2-3] = 2 mg/mL **D.** Residuals of Guinier fitting at [RAD51/BRC2-3] = 2 mg/mL **E.** Dimensionless Kratky plot at [RAD51/BRC2-3] = 1 mg/mL **F.** Residuals of Guinier fitting at [RAD51/BRC2-3] = 1 mg/mL **G.** Dimensionless Kratky plot at [RAD51/BRC2-3] = 0.5 mg/mL **H.** Residuals of Guinier fitting at [RAD51/BRC2-3] = 0.5 mg/mL **I.** Rg inferred from Guinier analyses **J.** SAXS p(r) analysis at different concentrations. Bottom Dmax and Rg inferred from p(r) analysis at different RAD51/BRC2-3 concentrations.

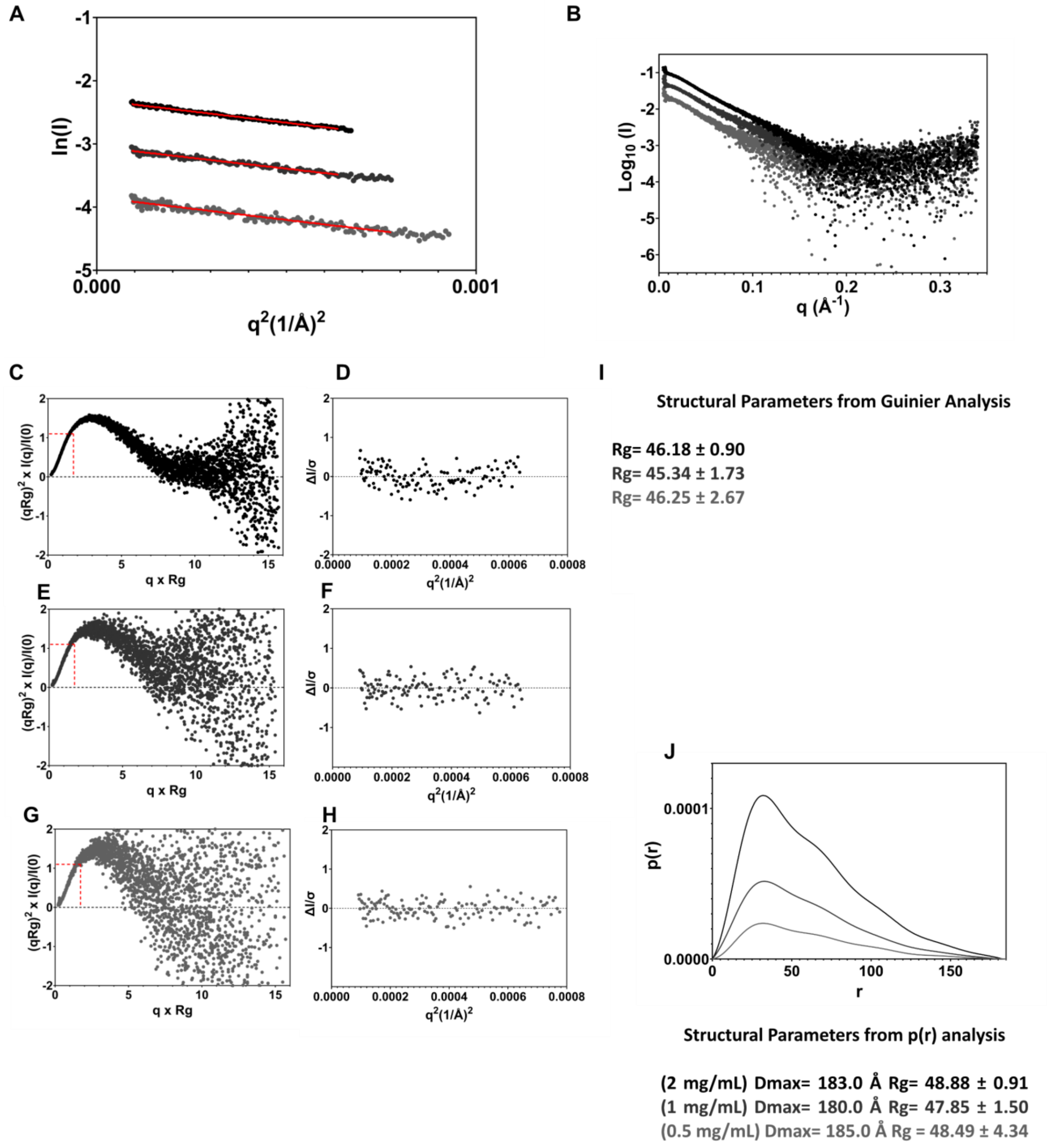


**Supplementary Figure 102 Batch-SAXS analysis of RAD51/BRC3-4 complex A.** Guinier analysis of SAXS Profile at different RAD51/BRC3-4 concentrations **B.** Scatter plot (logarithmic scale) of RAD51/BRC3-4 at different concentrations **C.** Dimensionless Kratky plot at [RAD51/BRC3-4] = 2 mg/mL **D.** Residuals of Guinier fitting at [RAD51/BRC3-4] = 2 mg/mL **E.** Dimensionless Kratky plot at [RAD51/BRC3-4] = 1 mg/mL **F.** Residuals of Guinier fitting at [RAD51/BRC3-4] = 1 mg/mL **G.** Dimensionless Kratky plot at [RAD51/BRC3-4] = 0.5 mg/mL **H.** Residuals of Guinier fitting at [RAD51/BRC3-4] = 0.5 mg/mL **I.** Rg inferred from Guinier analyses **J.** SAXS p(r) analysis at different concentrations. Bottom Dmax and Rg inferred from p(r) analysis at different RAD51/BRC3-4 concentrations.

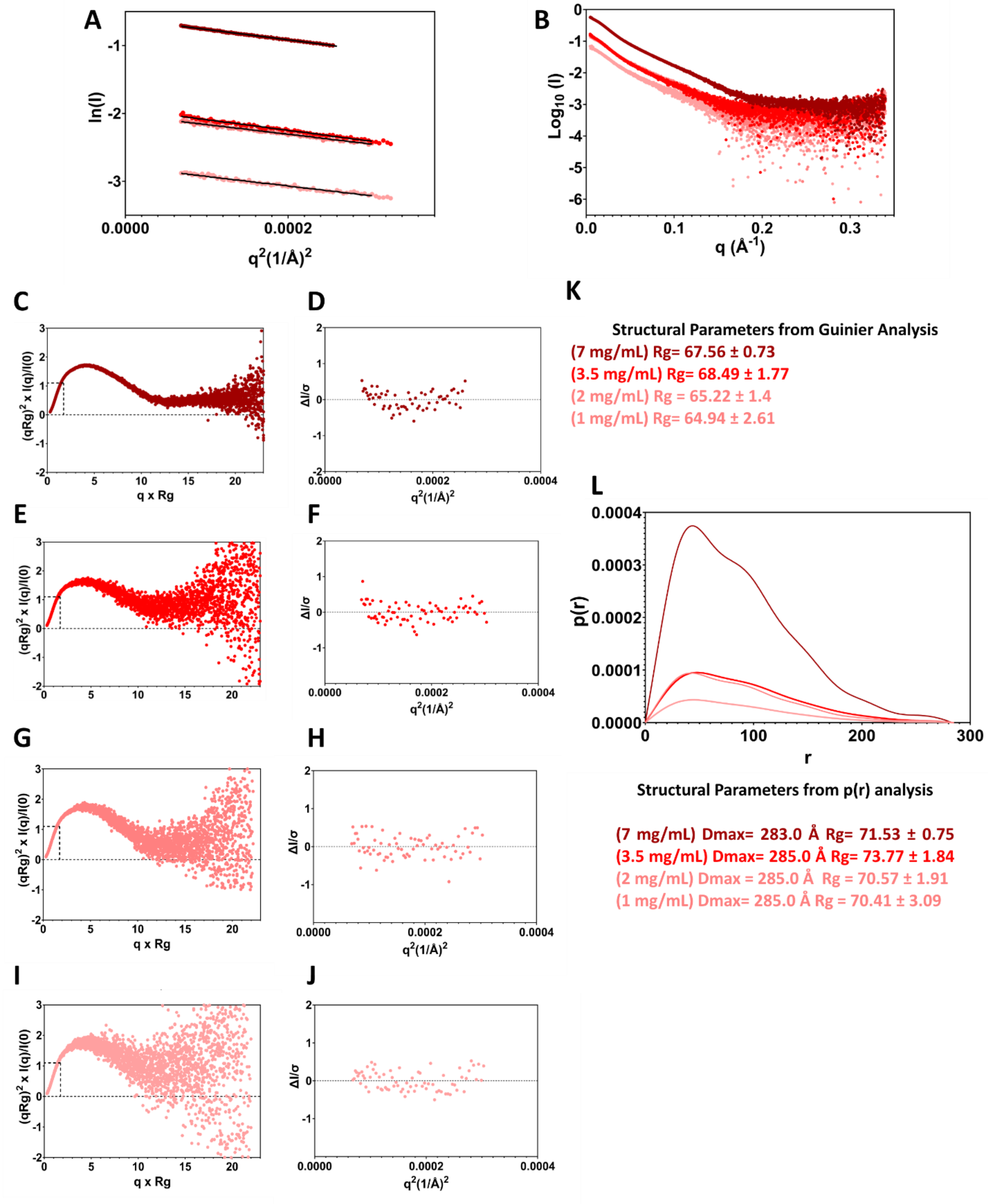


**Supplementary Figure 103 Batch-SAXS analysis of RAD51/BRC2-4 complex A.** Guinier analysis of SAXS Profile at different RAD51/BRC2-4 concentrations **B.** Scatter plot (logarithmic scale) of RAD51/BRC1-4 at different concentrations **C.** Dimensionless Kratky plot at [RAD51/BRC2-4] = 7 mg/mL **D.** Residuals of Guinier fitting at [RAD51/BRC2-4] = 7 mg/mL **E.** Dimensionless Kratky plot at [RAD51/BRC2-4] = 3.5 mg/mL **F.** Residuals of Guinier fitting at [RAD51/BRC2-4] = 3.5 mg/mL **G.** Dimensionless Kratky plot at [RAD51/BRC2-4] = 2 mg/mL **H.** Residuals of Guinier fitting at [RAD51/BRC2-4] = 2 mg/mL **I.** Dimensionless Kratky plot at [RAD51/BRC2-4] = 1 mg/mL **J.** Residuals of Guinier fitting at [RAD51/BRC2-4] = 1 mg/mL **K.** Rg inferred from Guinier analyses **L.** SAXS p(r) analysis at different concentrations. Bottom Dmax and Rg inferred from p(r) analysis at different RAD51/BRC1-4 concentrations.

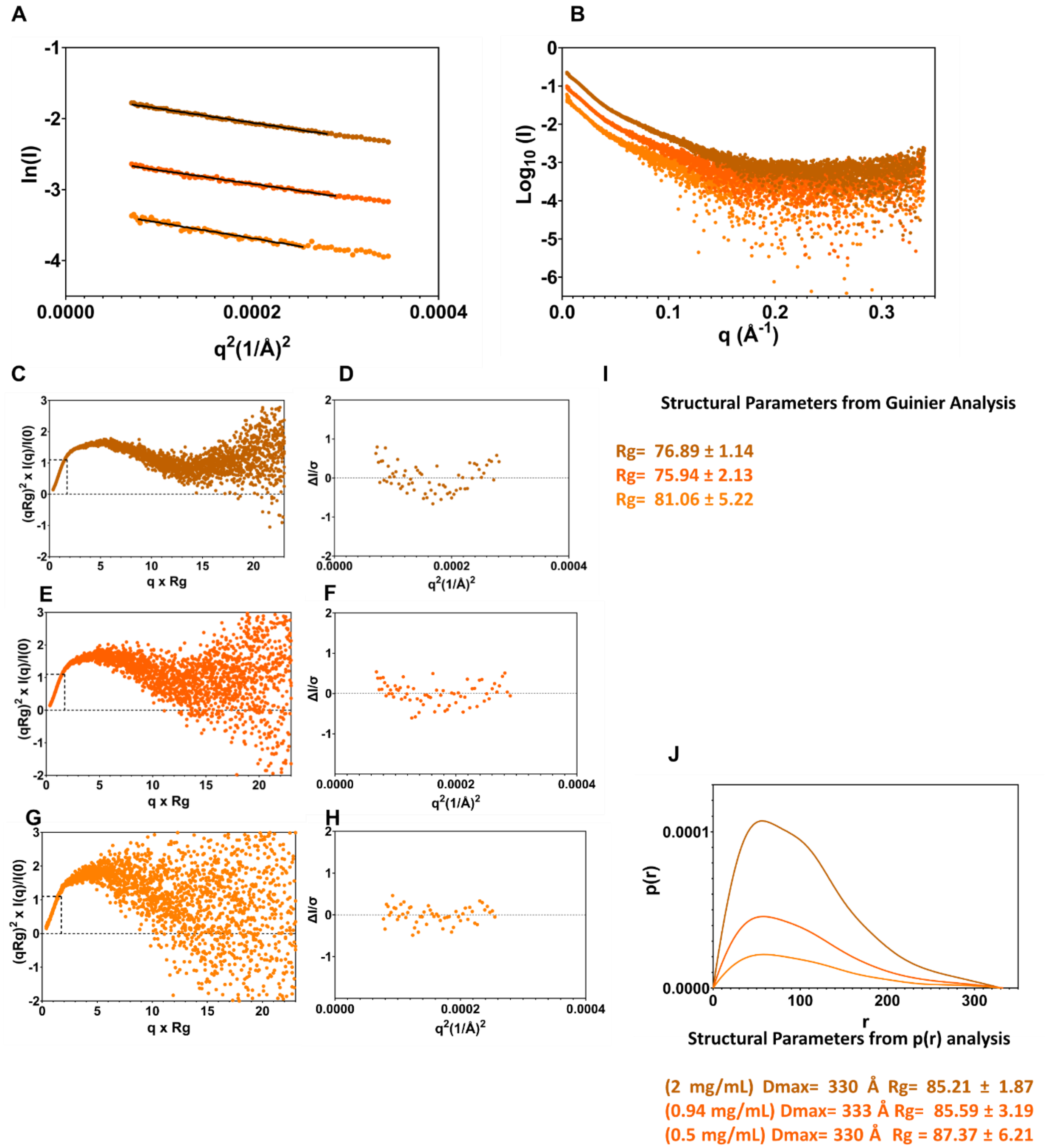


**Supplementary Figure 104 Batch-SAXS analysis of RAD51/BRC1-4 complex A.** Guinier analysis of SAXS Profile at different RAD51/BRC1-4 concentrations **B.** Scatter plot (logarithmic scale) of RAD51/BRC1-4 at different concentrations **C.** Dimensionless Kratky plot at [RAD51/BRC1-4] = 2 mg/mL **D.** Residuals of Guinier fitting at [RAD51/BRC1-4] = 2 mg/mL **E.** Dimensionless Kratky plot at [RAD51/BRC1-4] = 0.94 mg/mL **F.** Residuals of Guinier fitting at [RAD51/BRC1-4] = 0.94 mg/mL **G.** Dimensionless Kratky plot at [RAD51/BRC1-4] = 0.5 mg/mL **H.** Residuals of Guinier fitting at [RAD51/BRC1-4] = 0.5 mg/mL **I.** Rg inferred from Guinier analyses **J.** SAXS p(r) analysis at different concentrations. Bottom Dmax and Rg inferred from p(r) analysis at different RAD51/BRC1-4 concentrations.

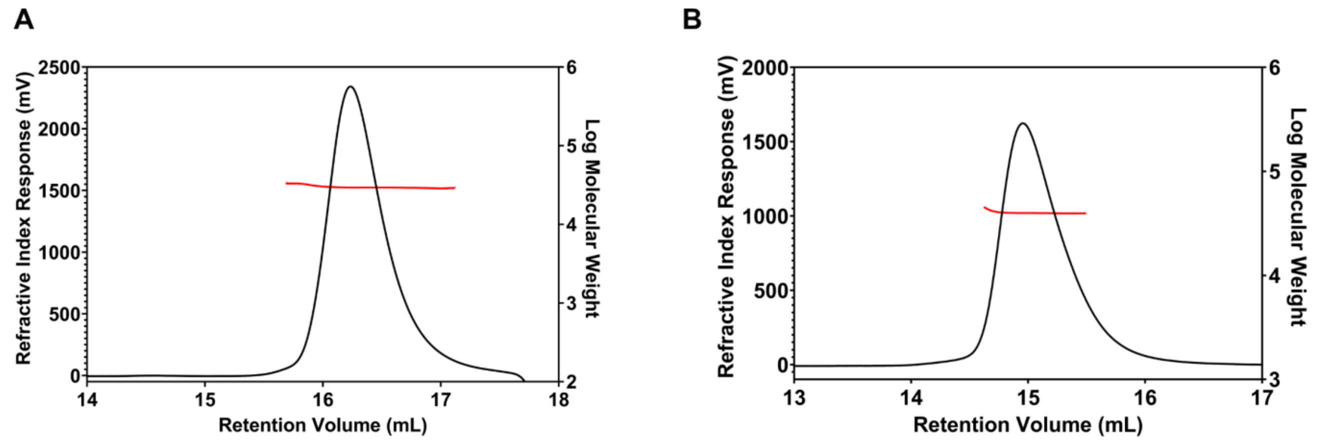


**Supplementary Figure 105 Static light scattering analysis** of **A.** $^{15}$N Δ97-RAD51-BRC4 and **B.** $^{15}$N RAD51-BRC4

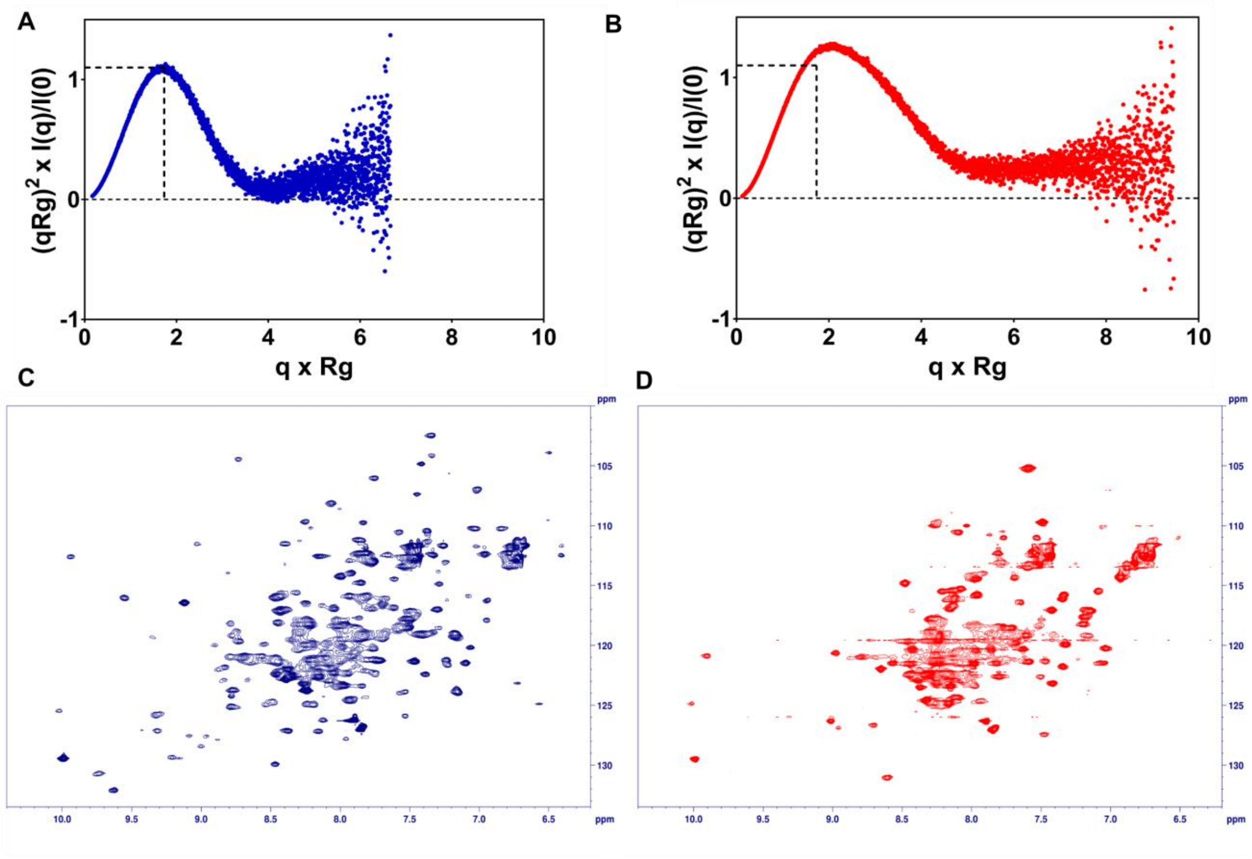


**Supplementary Figure 106 SAXS and NMR studies on Δ97-RAD51/BRC4 and RAD51/BRC4** Dimensionless Kratky Plots of **A.** Δ97-RAD51/BRC4 **B.** RAD51/BRC4, $^{1}$H-$^{15}$N Heteronuclear Single Quantum Coherence (HSQC) spectra of **C.** Δ97-RAD51/BRC4 **D.** RAD51/BRC4

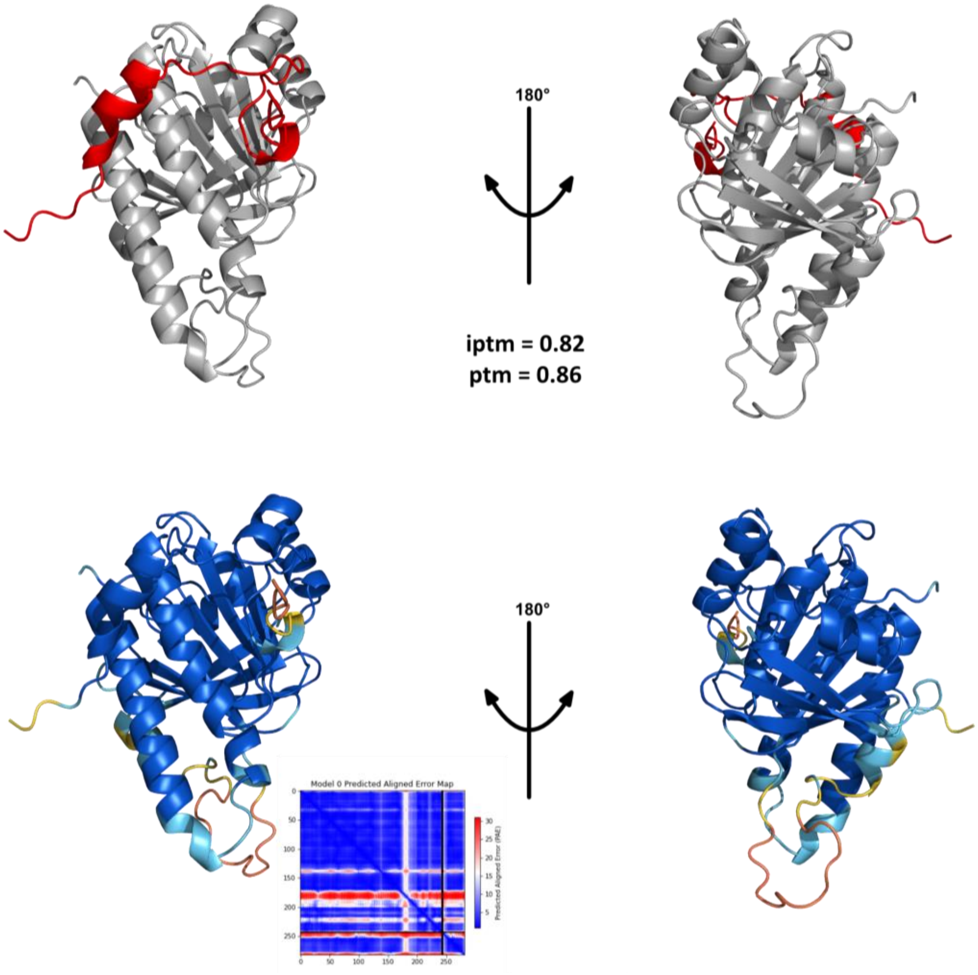


**Supplementary Figure 107 Alphafold model generation of Δ97-RAD51-BRC4** Top: Structure of the generated model where Δ97-RAD51 is grey and the BRC4 peptide is red. Bottom: pLDDT representation and PAE matrix of the shown model.

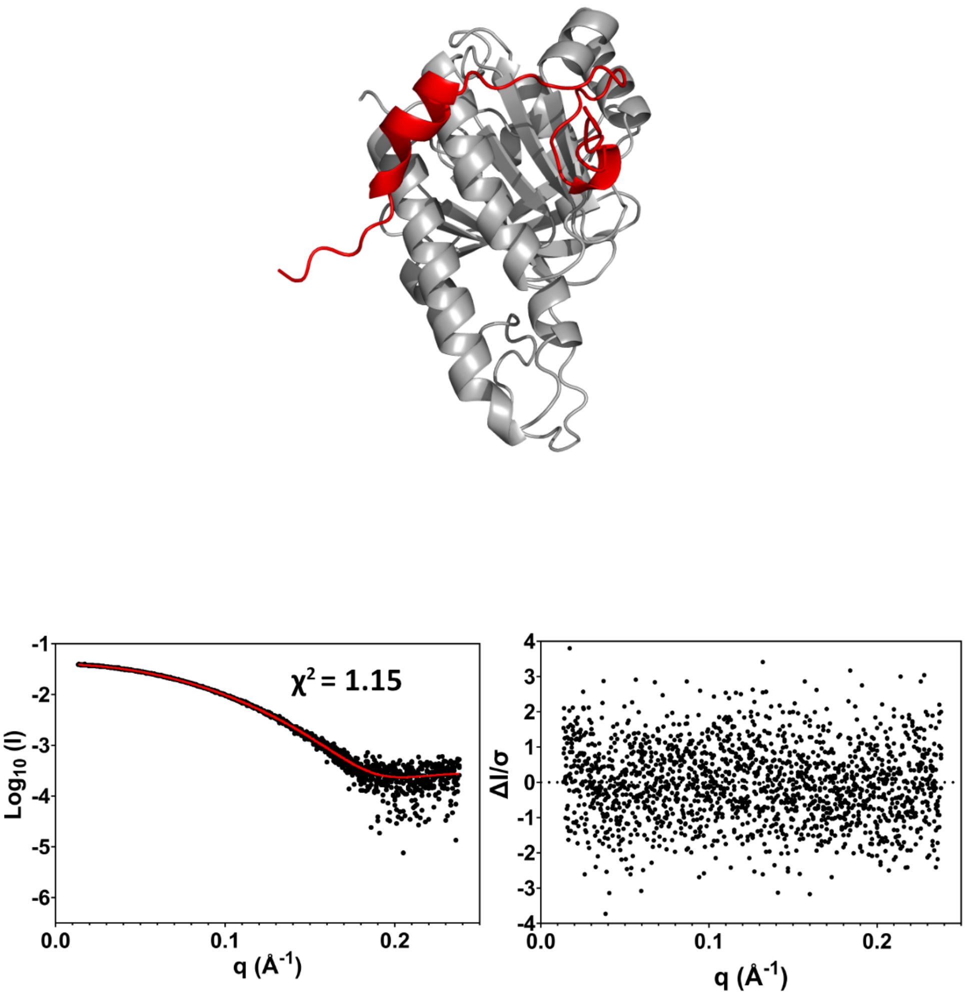


**Supplementary Figure 108 FoXS modeling of Δ97-RAD51/BRC4 AF prediction.** Top: The AF Δ97-RAD51/BRC4 complex prediction. Bottom left: Model fit to experimental SAXS data. Bottom right: model residuals

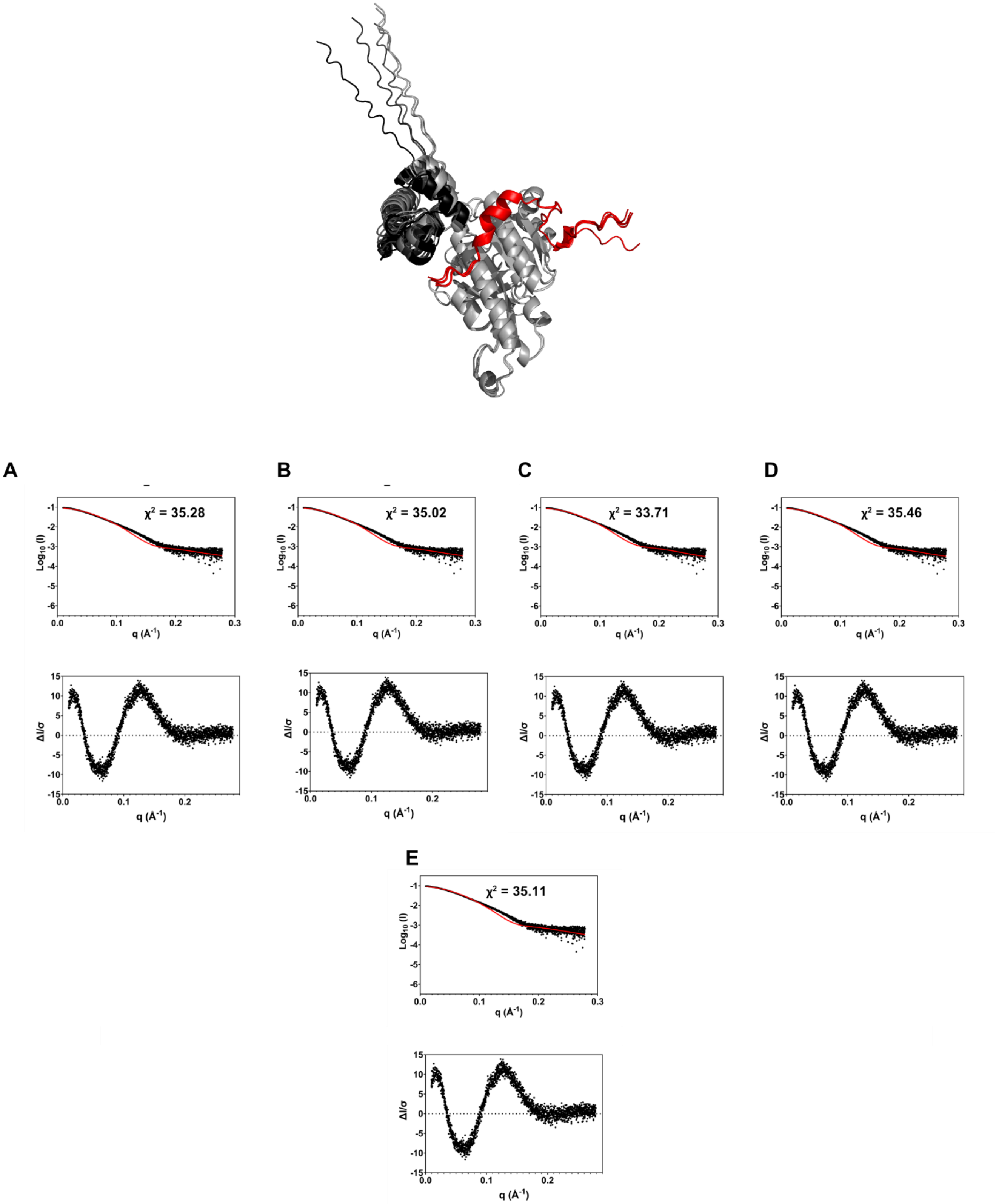


**Supplementary Figure 109** FoXS modelling of the RAD51-BRC4 complex predictions generated through AlphaFold2. Top: comparison between the experimental SAXS data and the input structure (red curve) along with $\chi^2$. Bottom: standardized residuals associated to FoXS modelling. **A.**Model 0 **B.** Model 1 **C.** Model 2 **D.** Model 3 **E.** Model 4

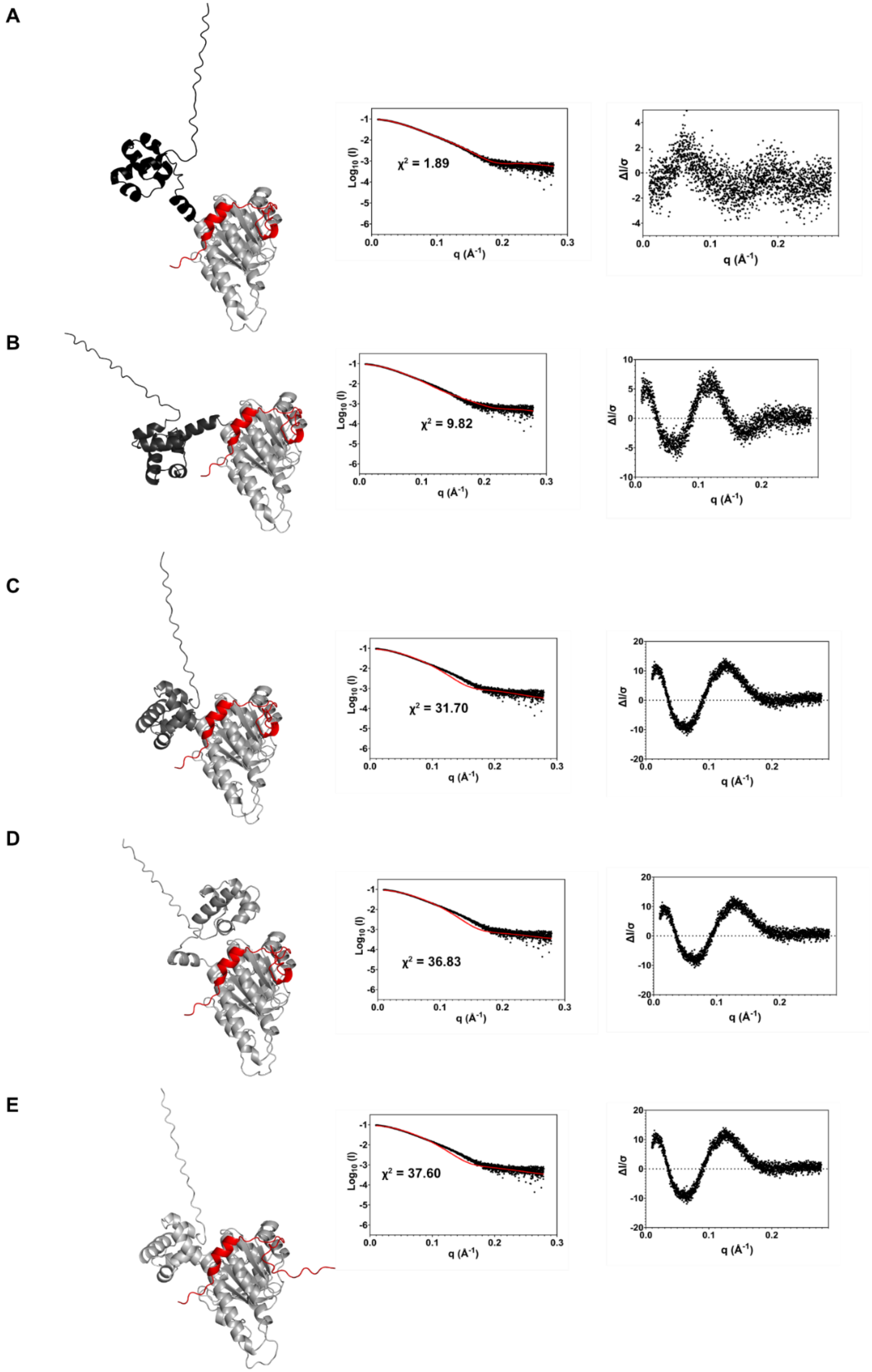


**Supplementary Figure 110** FoXS modelling of the RAD51-BRC4 complex predictions generated through AlphaFold3. Left: the BRC4 peptide is colored red while the N-Terminal domain is displayed in different shades of grey depending on the model. Center: comparison between the experimental SAXS data and the input structure (red curve) along with $\chi^2$. Right: standardized residuals associated with FoXS modelling. **A.** Model 0 **B.** Model 1 **C.** Model 2 **D.** Model 3 **E.** Model 4

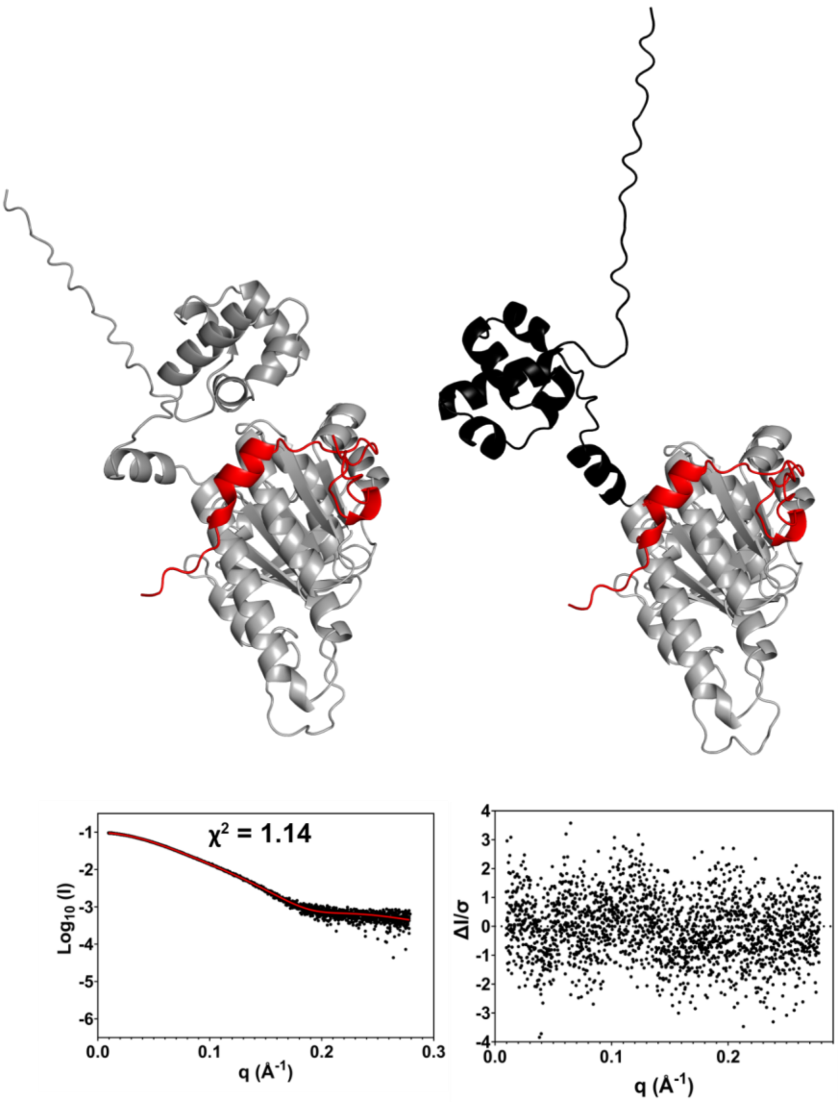


**Supplementary Figure 111 Multi-FoXS modeling of RAD51/BRC4 AF3 prediction.** Top: The two best scoring Multi-FoXS 2-state models for the AF RAD51/BRC4 complex. Bottom left: Model fit to experimental SAXS data. Bottom right: model residuals.

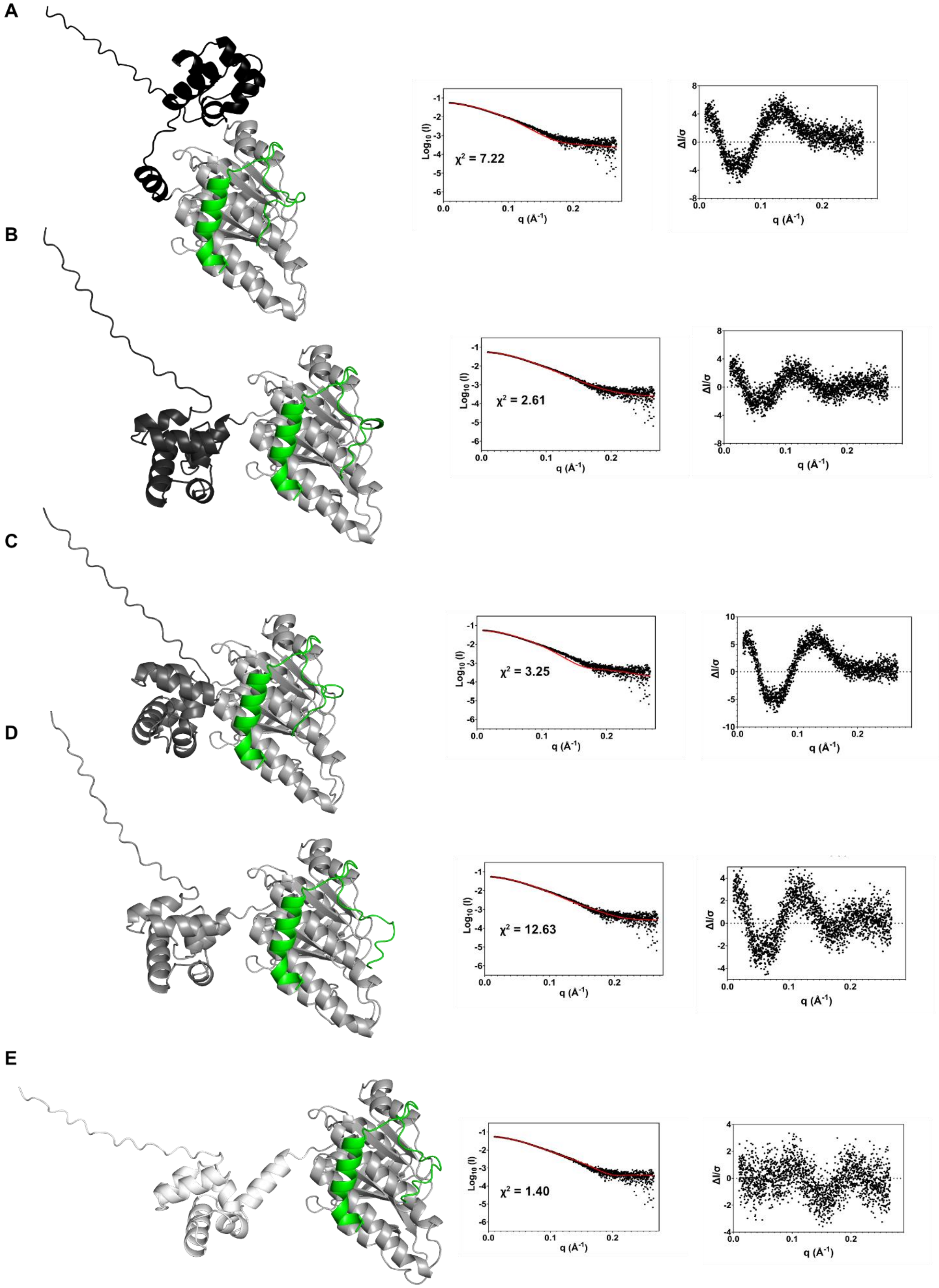


**Supplementary Figure 112** FoXS modelling of the RAD51-BRC2 complex predictions generated through AlphaFold3. Left: the BRC2 peptide is colored green while the N-Terminal domain is displayed in different shades of grey depending on the model. Center: comparison between the experimental SAXS data and the input structure (red curve) along with $\chi^2$. Right: standardized residuals associated with FoXS modelling. **A.**Model 0 **B.** Model 1 **C.** Model 2 **D.** Model 3 **E.** Model 4

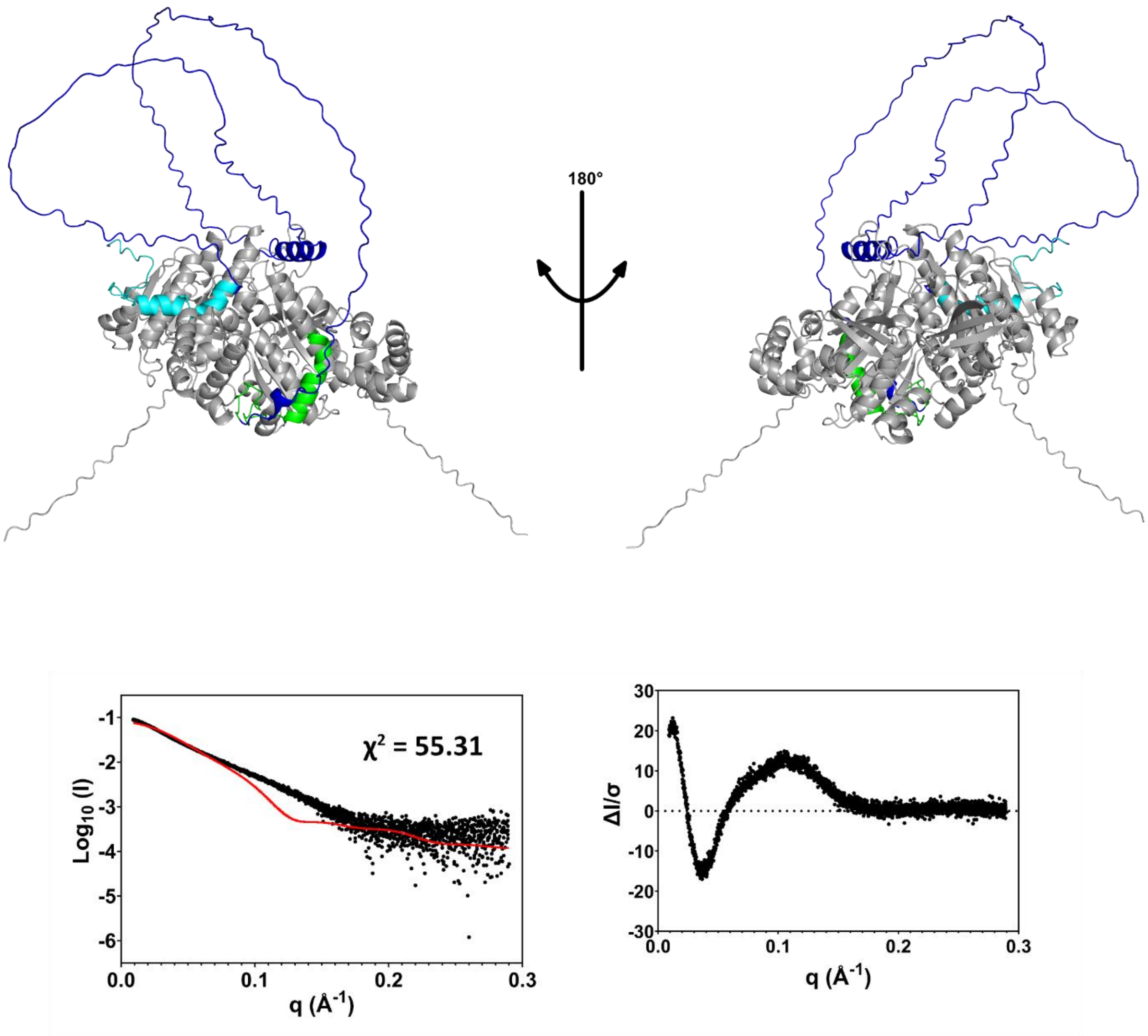


**Supplementary Figure 113 FoXS Modelling of Alphafold prediction of the RAD51/BRC1-2 complex** Top. Structure of the generated model. RAD51 monomers are colored grey, BRC1 is cyan, BRC2 is green and the BRC-repeats connecting loop is colored blue. Bottom left: FoXS modelling for the input structure (red curve) compared to the experimental SAXS data (black dots) Bottom right: Standardized residuals associated to FoXS modelling of the selected structure.

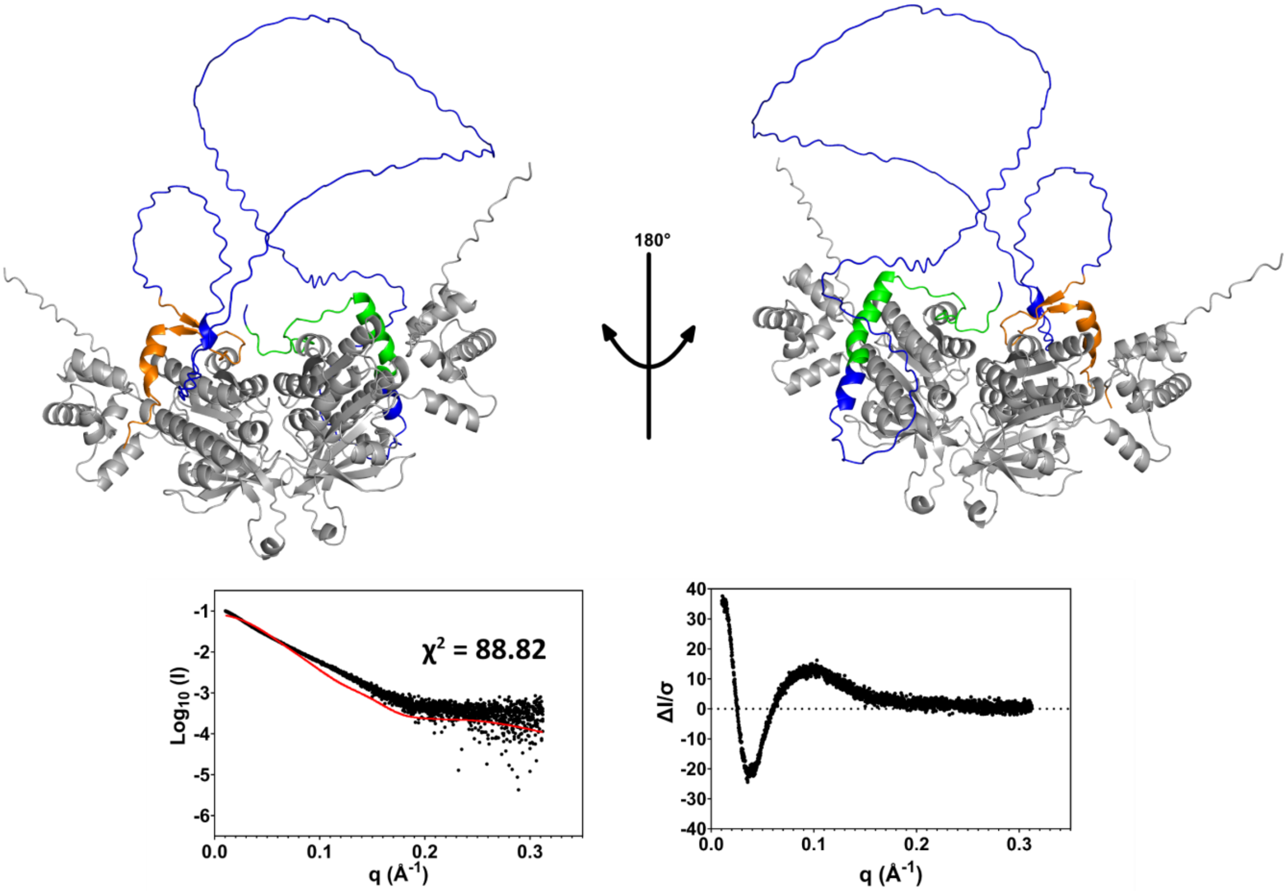


**Supplementary Figure 114 FoXS Modelling of Alphafold prediction of the RAD51/BRC2-3 complex** Top. Structure of the generated model. RAD51 monomers are colored grey, BRC2 is green, BRC3 is orange and the BRC-repeats connecting loop is colored blue. Bottom left: FoXS modelling for the input structure (red curve) compared to the experimental SAXS data (black dots) Bottom right: Standardized residuals associated to FoXS modelling of the selected structure.

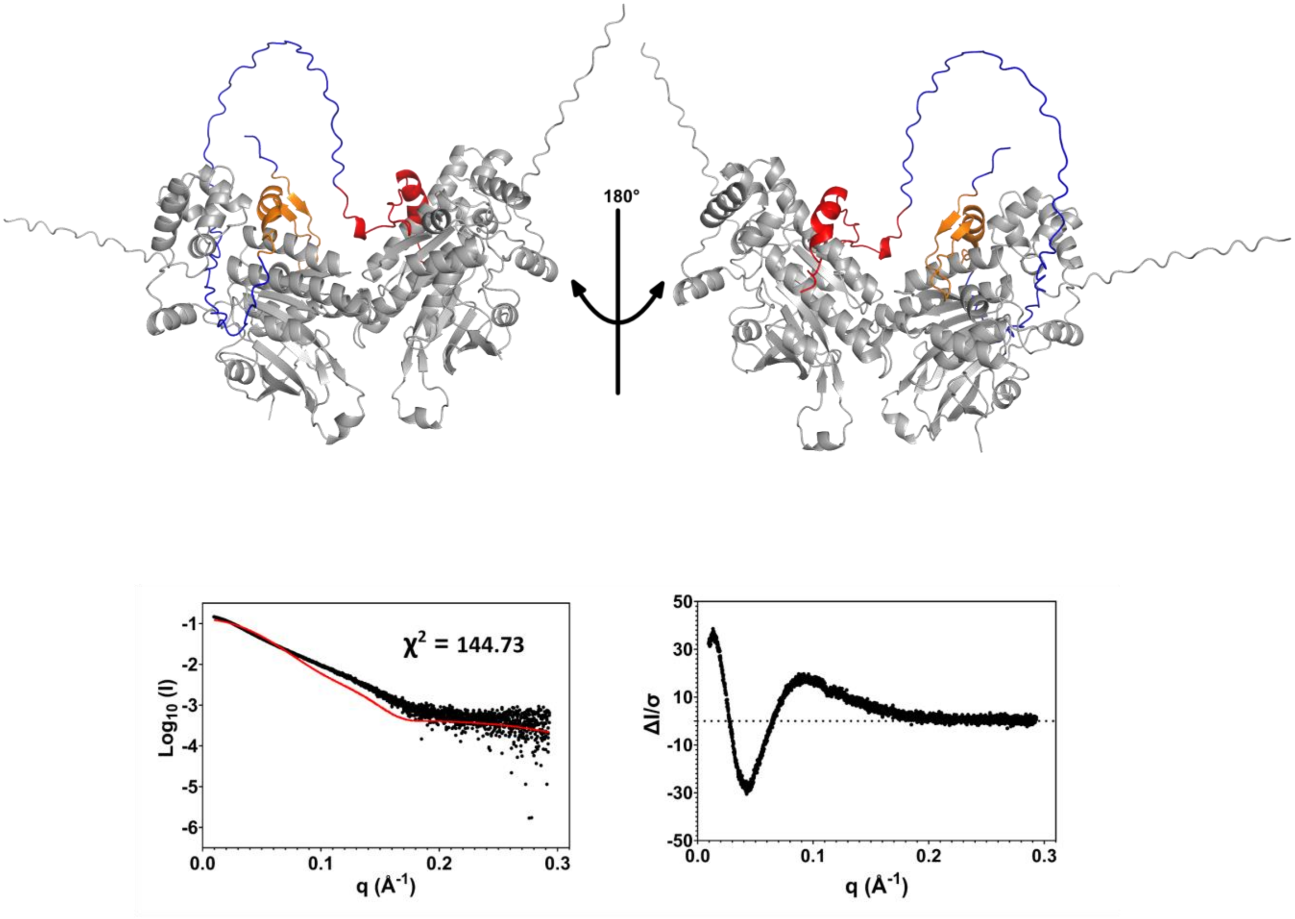


**Supplementary Figure 115 Modelling of Alphafold prediction of the RAD51/BRC3-4 complex** Top. Structure of the generated model. RAD51 monomers are colored grey, BRC3 is orange, BRC4 is red and the BRC-repeats connecting loop is colored blue. Bottom left: FoXS modelling for the input structure (red curve) compared to the experimental SAXS data (black dots) Bottom right: Standardized residuals associated to FoXS modelling of the selected structure.

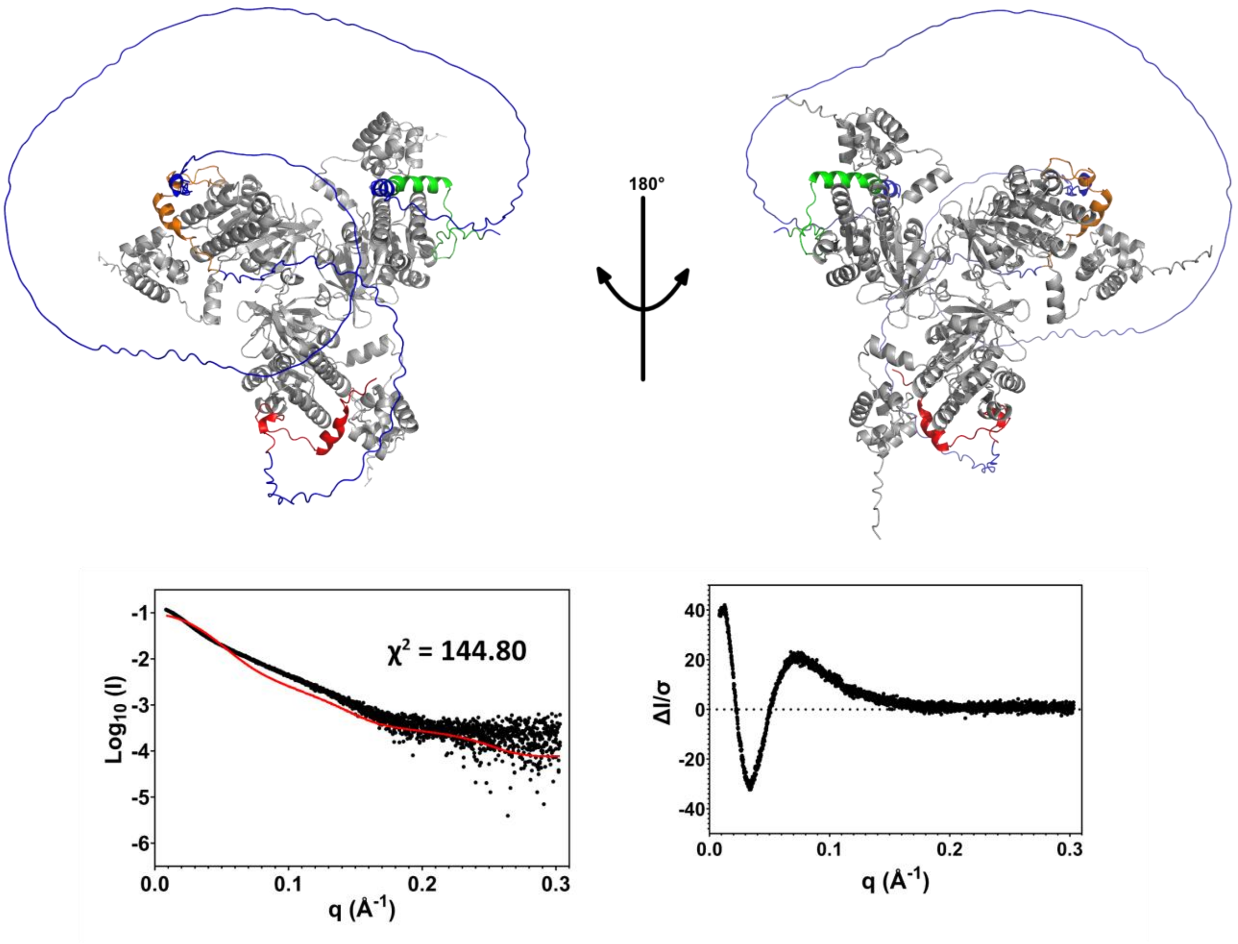


**Supplementary Figure 116 FoXS Modelling of Alphafold prediction of the RAD51/BRC2-4 complex** Top. Structure of the generated model. RAD51 monomers are colored grey, BRC2 is green, BRC3 is orange, BRC4 is red and the BRC-repeats connecting loops are colored blue. Bottom left: FoXS modelling for the input structure (red curve) compared to the experimental SAXS data (black dots) Bottom right: Standardized residuals associated to FoXS modelling of the selected structure.

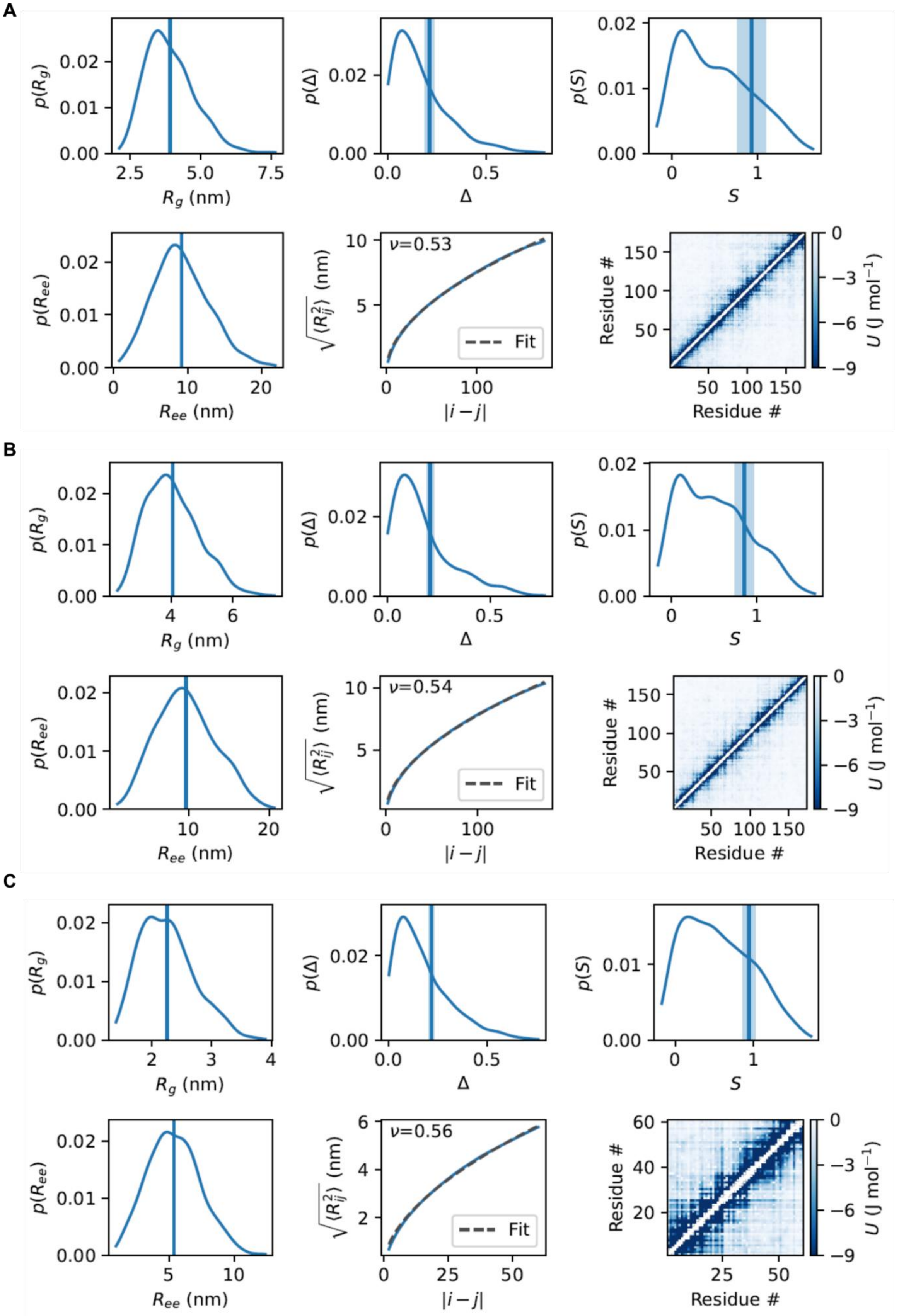


**Supplementary Figure 117 CALVADOS carried out on the the BRC-repeats connecting regions.** Molecular dynamics (MD) simulations of BRC-repeats connecting loops carried out through CALVADOS. This method provides estimation of conformational properties and show the interaction energy maps and distribution of radius of gyration. Through CALVADOS, we were able to estimate the Flory scaling exponent v for each BRC-repeats connecting region. **A**. BRC1-2, **B.** BRC2-3, **C.** BRC3-4

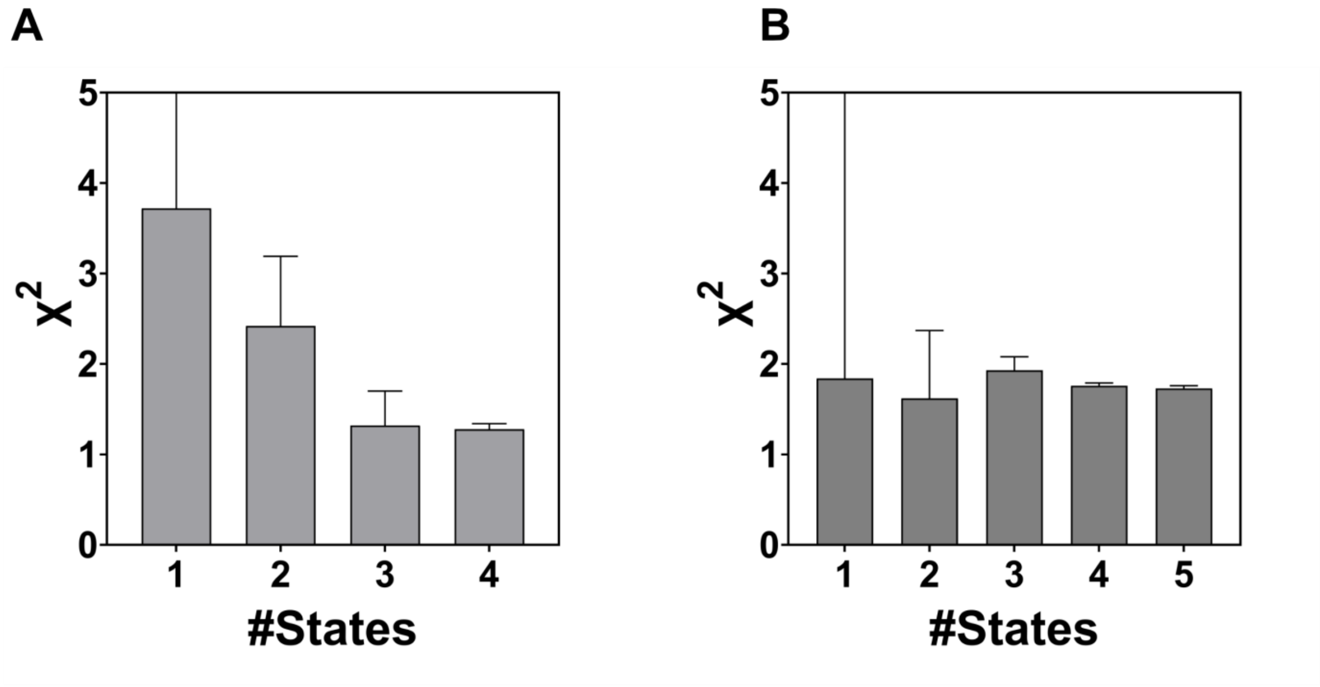


**Supplementary Figure 118 Histograms representation of the χ2 for two replicate of multi-state modelling performed through MultiFoXS using as an input the Alphafold prediction of the RAD51-BRC1-2 complex.**

**A**

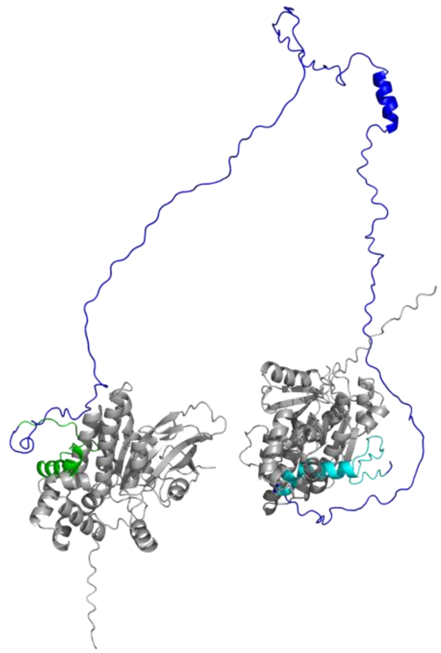

**B**

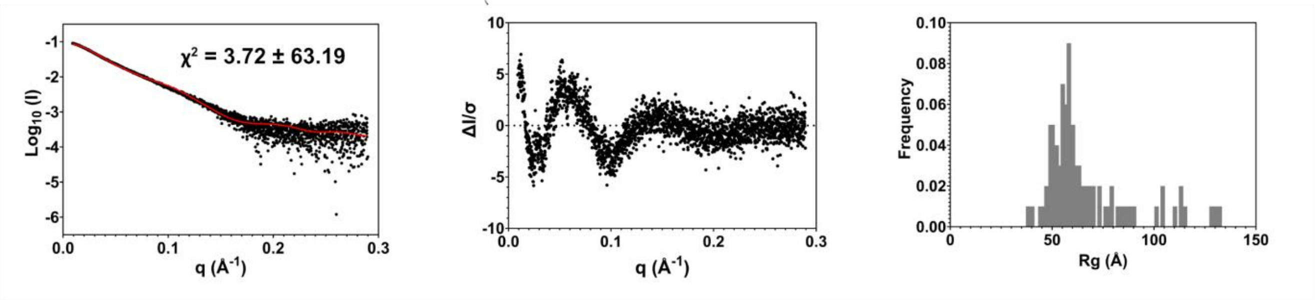


**Supplementary Figure 119 Multi-FoXS Modelling of Alphafold RAD51/BRC1-2 complex prediction model 1-state A.** Generated 1 state-model **B.** Left: fit to experimental SAXS data. Middle: normalized residuals of the reported fitting. Right: distribution of radii of gyration for the best scoring 1 state model

**A**

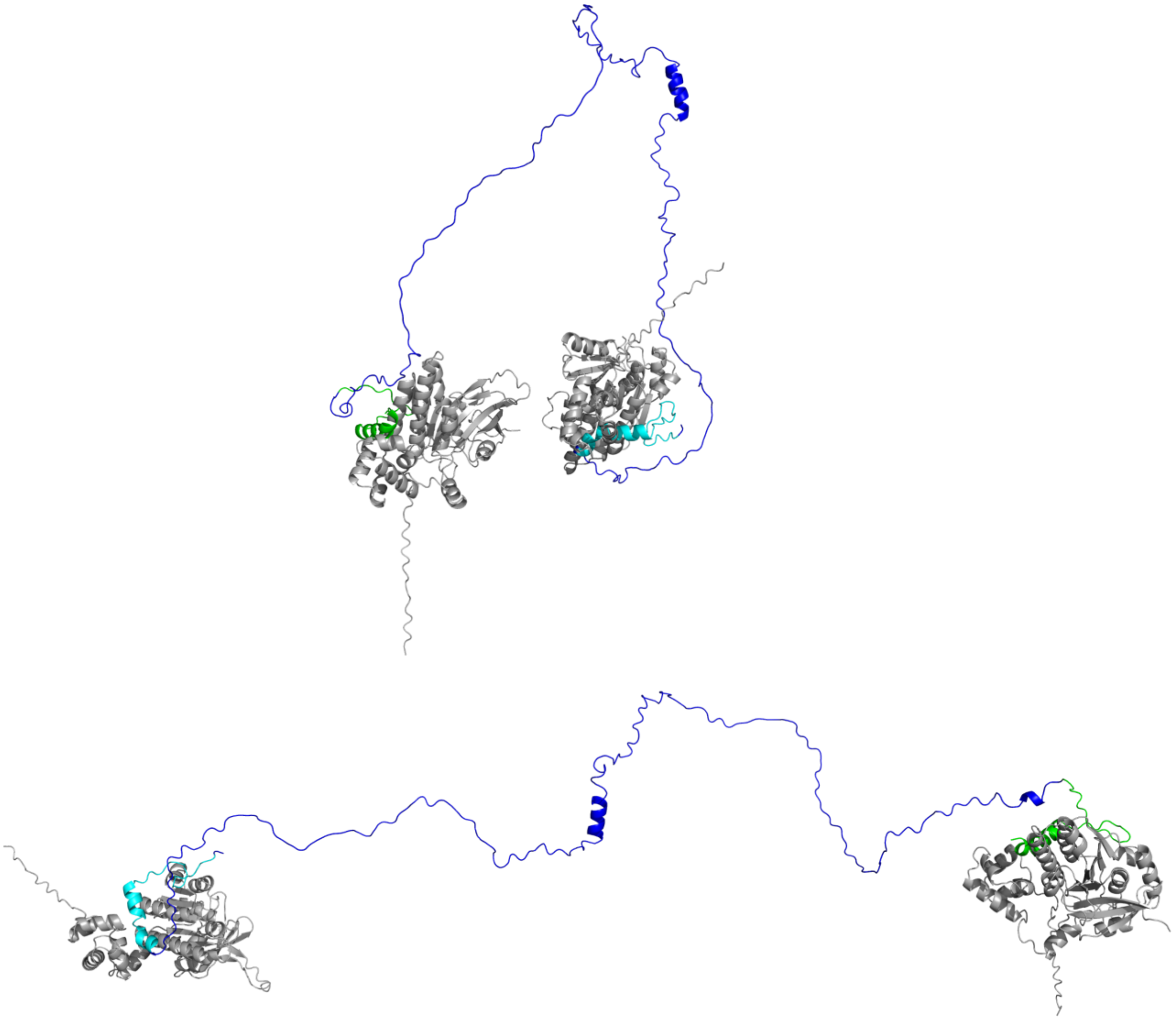

**B**

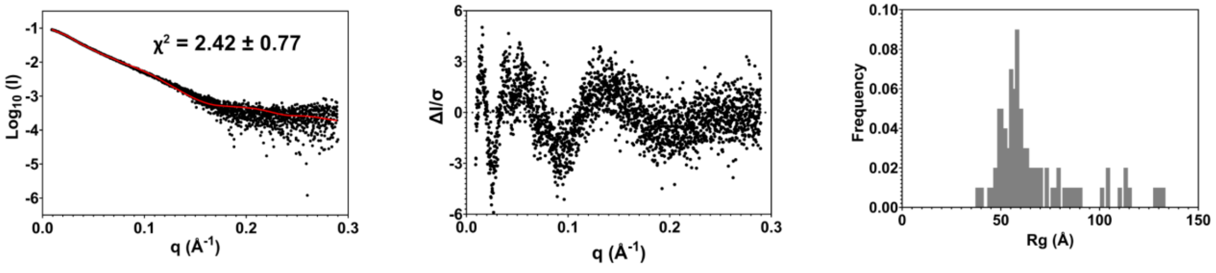


**Supplementary Figure 120 Multi-FoXS Modelling of Alphafold RAD51/BRC1-2 complex prediction model 2-state A.** Generated 2 state-model **B.** Left: fit to experimental SAXS data. Middle: normalized residuals of the reported fitting. Right: distribution of radii of gyration for the best scoring 2 state model

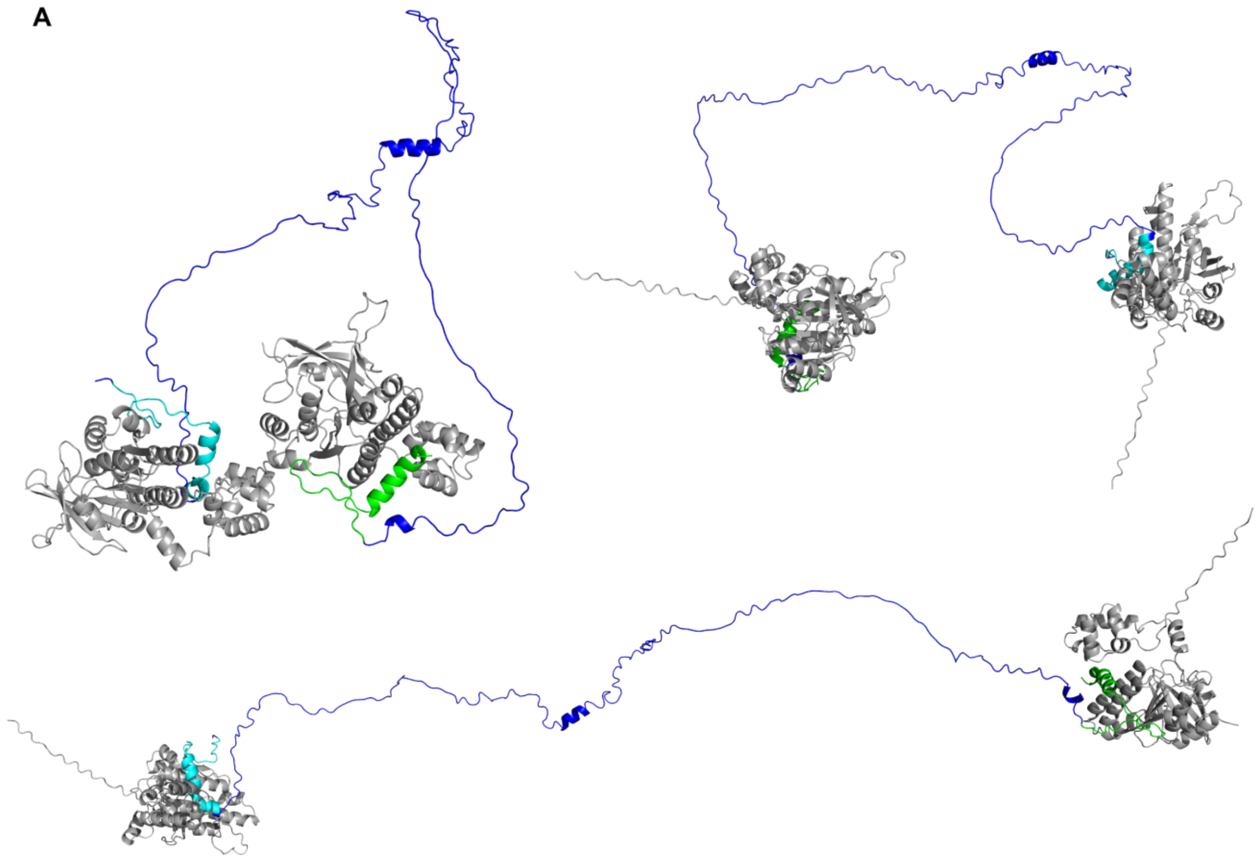


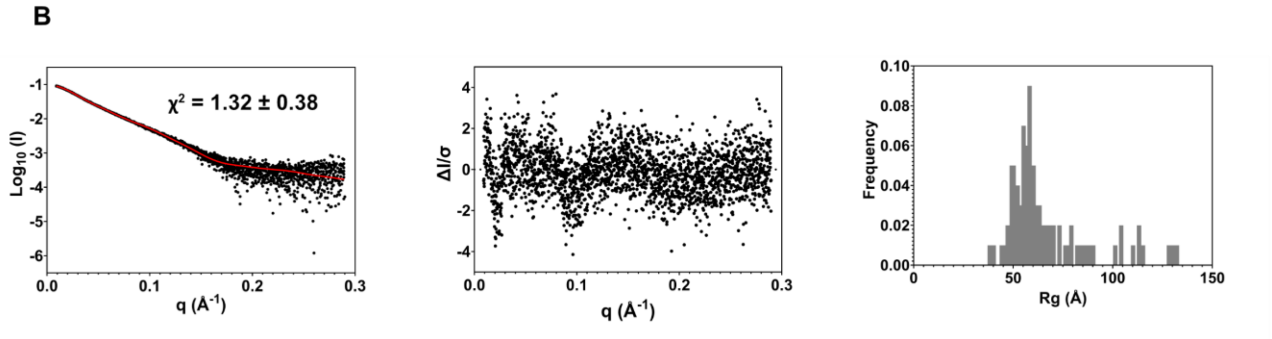


**Supplementary Figure 121 Multi-FoXS Modelling of Alphafold RAD51/BRC1-2 complex prediction model 3-state A.** Generated 3 state-model **B.** Left: fit to experimental SAXS data. Middle: normalized residuals of the reported fitting. Right: distribution of radii of gyration for the best scoring 3 state model

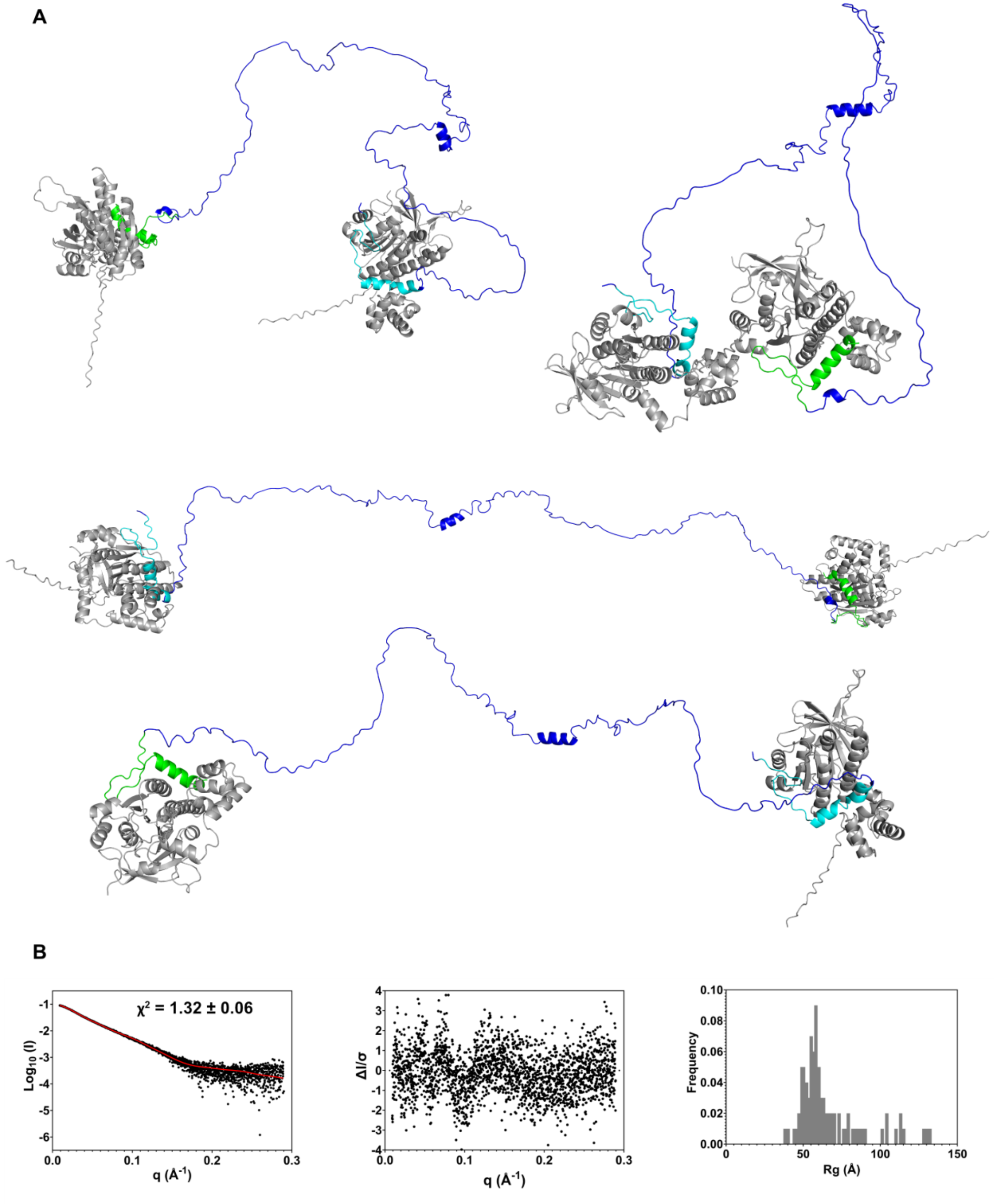


**Supplementary Figure 122 Multi-FoXS Modelling of Alphafold RAD51/BRC1-2 complex prediction model 4-state A.** Generated 4 state-model **B.** Left: fit to experimental SAXS data. Middle: normalized residuals of the reported fitting. Right: distribution of radii of gyration for the best scoring 4 state model

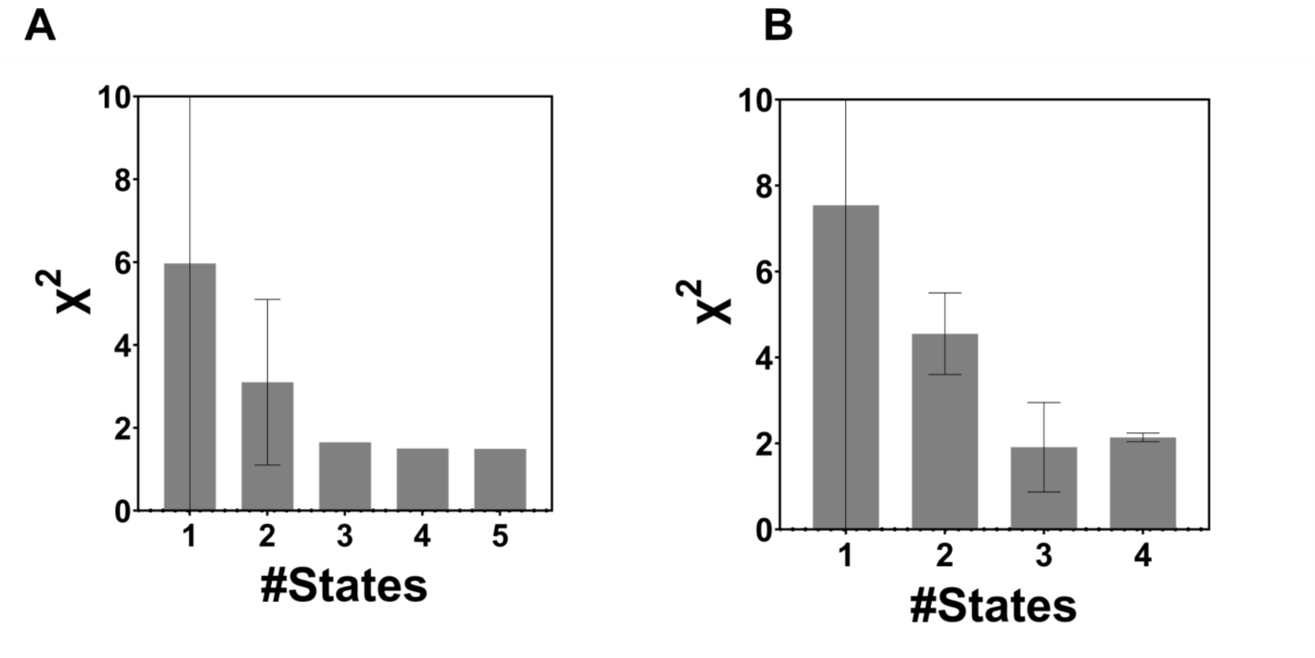


**Supplementary Figure 123 Histograms representation of the χ2 for two replicate of multi-state modelling performed through MultiFoXS using as an input the Alphafold prediction of the RAD51-BRC2-3 complex.**

A

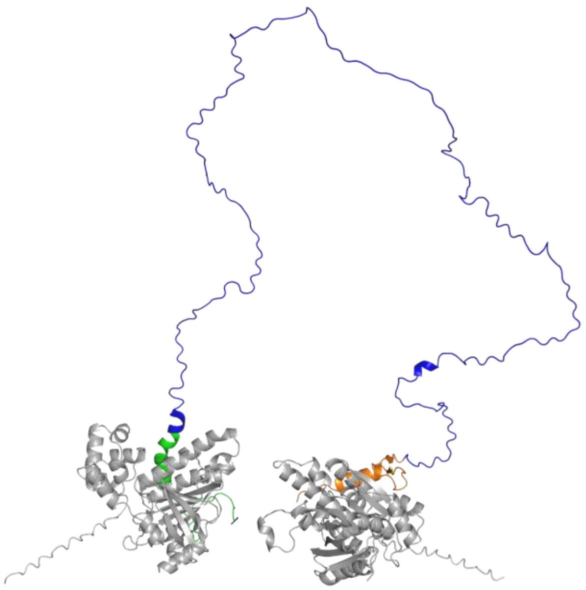

B

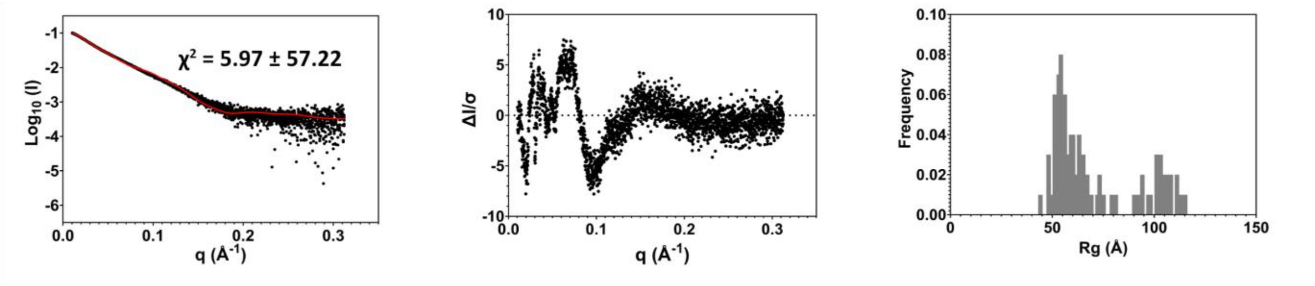


**Supplementary Figure 124 Multi-FoXS Modelling of Alphafold RAD51/BRC2-3 complex prediction model 1-state A.** Generated 1 state-model **B.** Left: fit to experimental SAXS data. Middle: normalized residuals of the reported fitting. Right: distribution of radii of gyration for the best scoring 1 state model

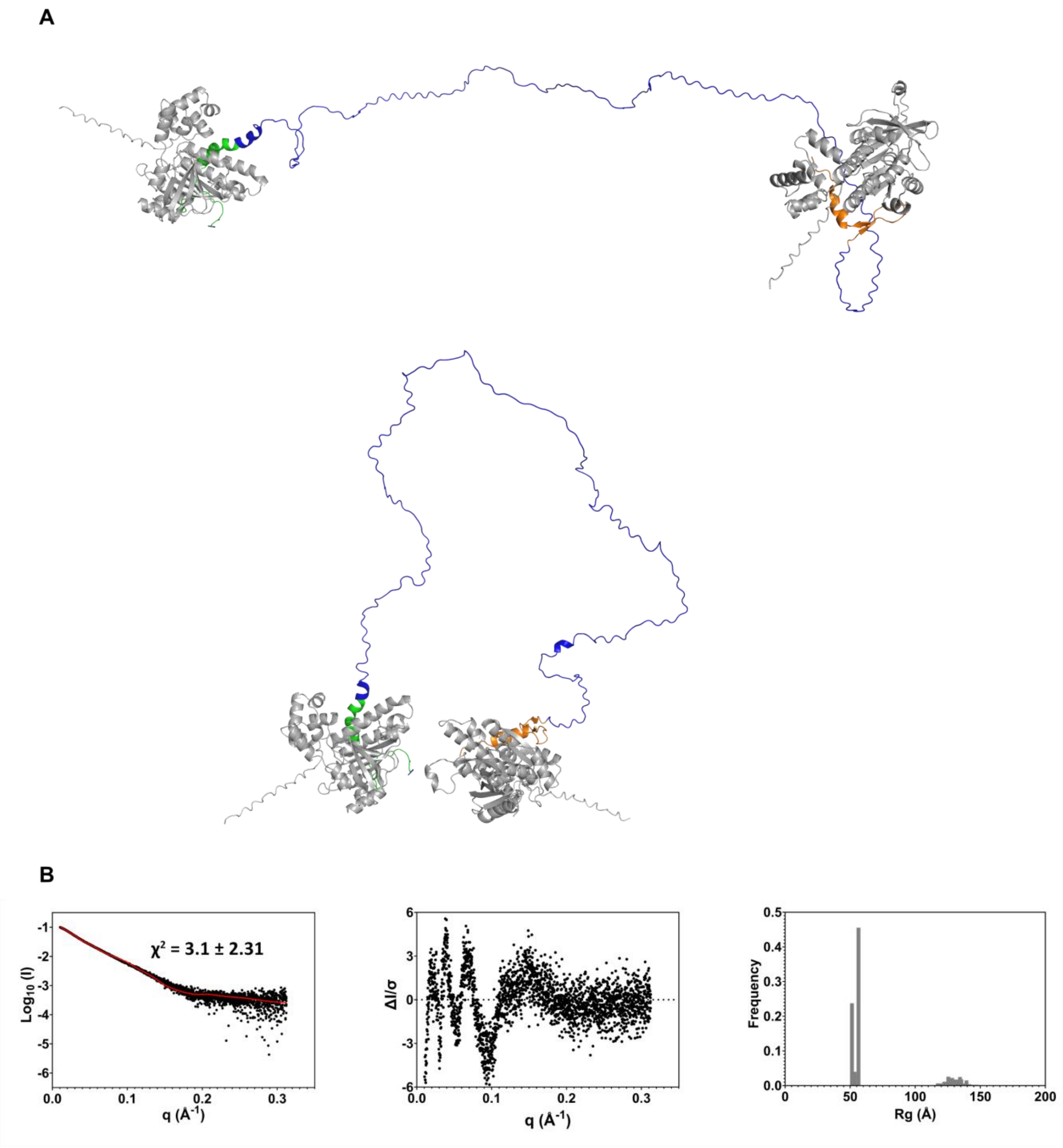


**Supplementary Figure 125 Multi-FoXS Modelling of Alphafold RAD51/BRC2-3 complex prediction model 2-state A.** Generated 2 state-model **B.** Left: fit to experimental SAXS data. Middle: normalized residuals of the reported fitting. Right: distribution of radii of gyration for the best scoring 2 state model

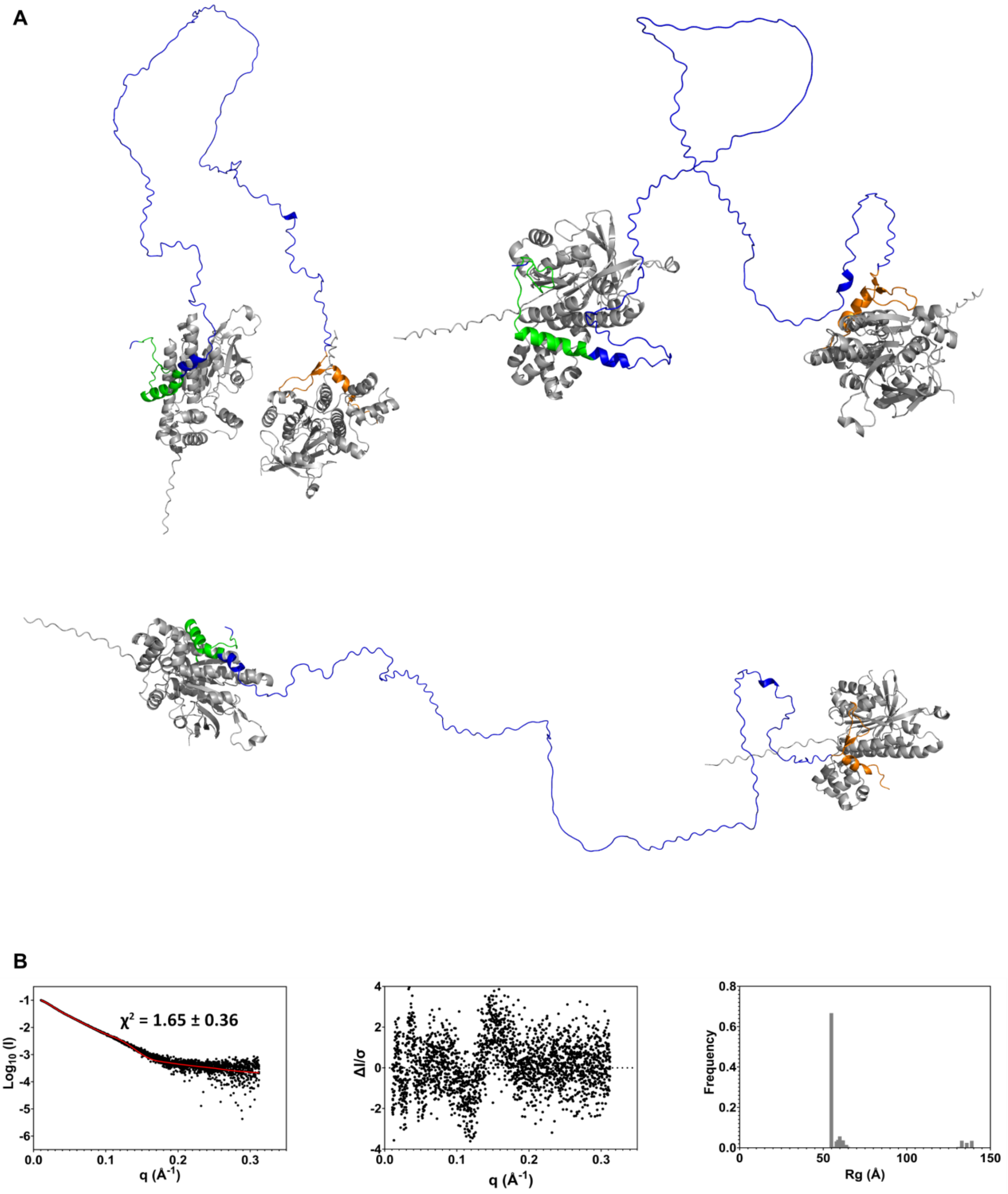


**Supplementary Figure 126 Multi-FoXS Modelling of Alphafold RAD51/BRC2-3 complex prediction model 3-state A.** Generated 3 state-model **B.** Left: fit to experimental SAXS data. Middle: normalized residuals of the reported fitting. Right: distribution of radii of gyration for the best scoring 3 state model

A

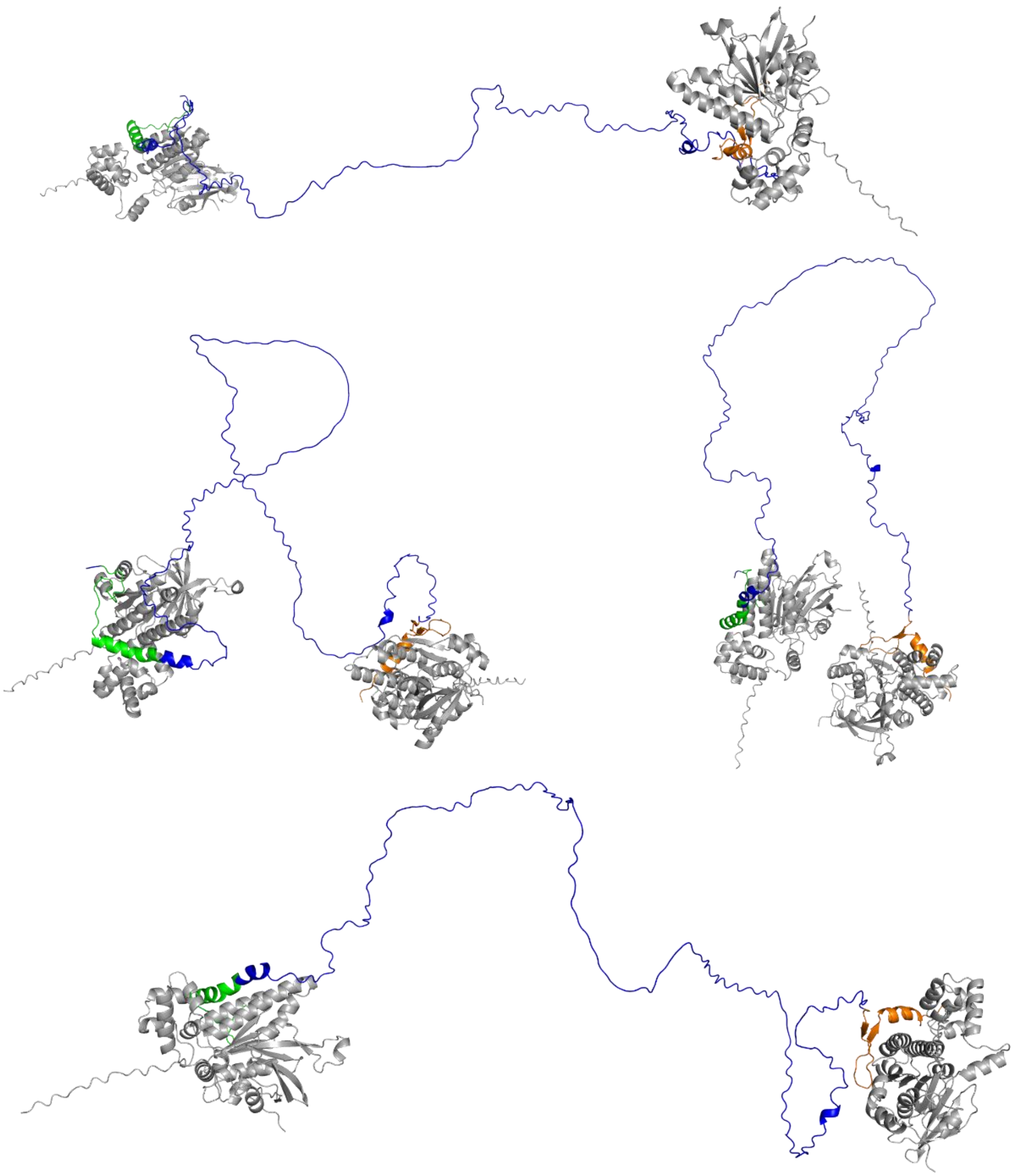

B

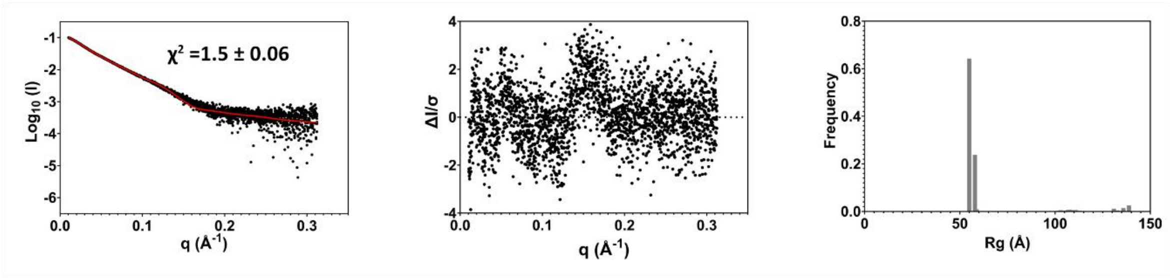


**Supplementary Figure 127 Multi-FoXS Modelling of Alphafold RAD51/BRC2-3 complex prediction model 4-state A.** Generated 4 state-model **B.** Left: fit to experimental SAXS data. Middle: normalized residuals of the reported fitting. Right: distribution of radii of gyration for the best scoring 4 state model

A

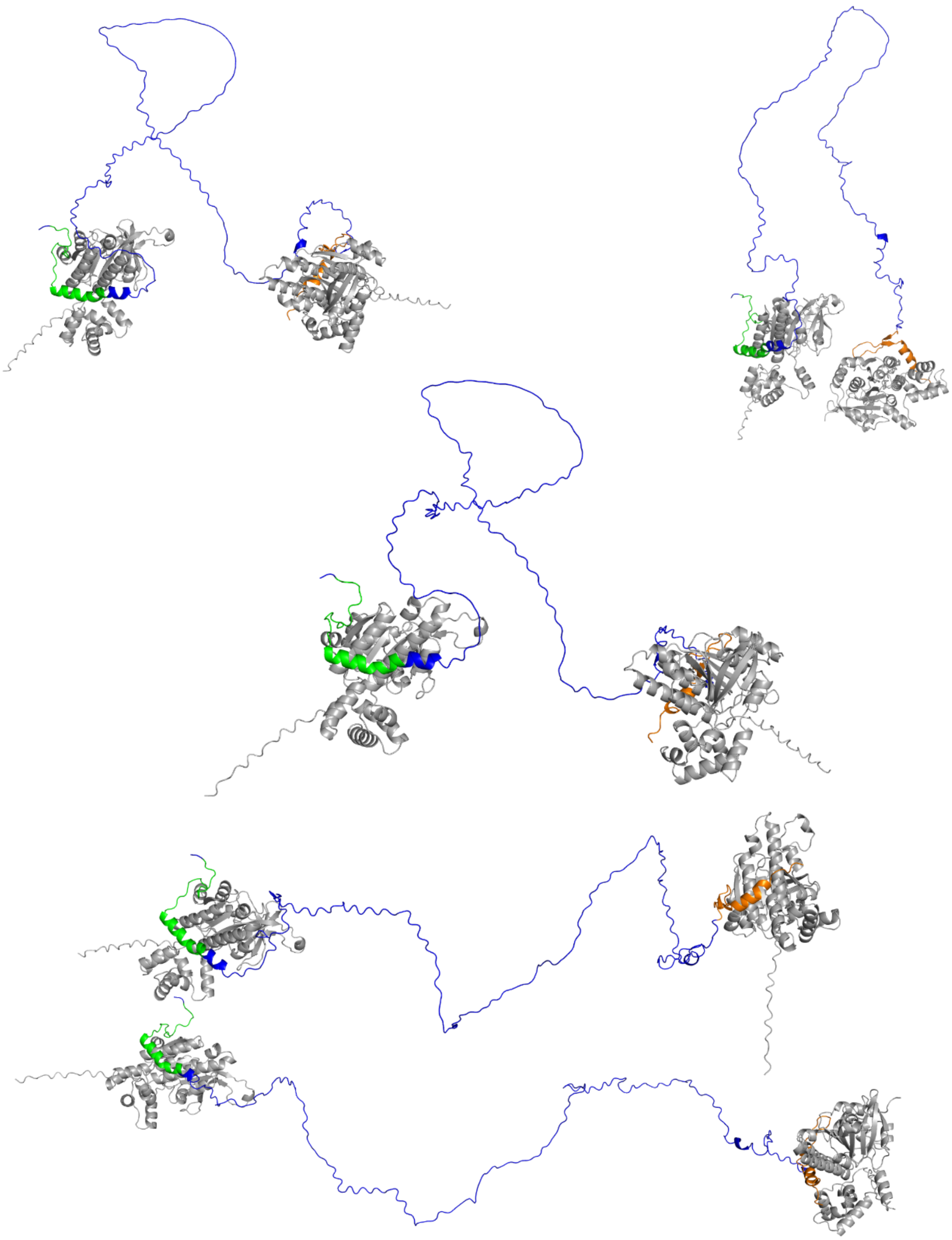

B

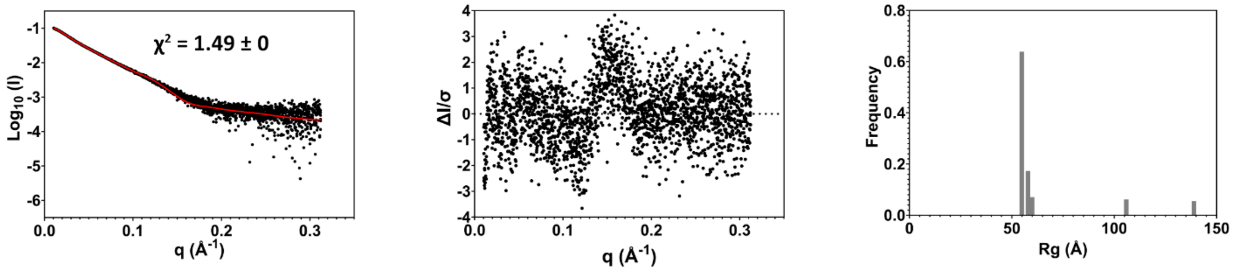


**Supplementary Figure 128 Multi-FoXS Modelling of Alphafold RAD51/BRC2-3 complex prediction model 5-state A.** Generated 5 state-model **B.** Left: fit to experimental SAXS data. Middle: normalized residuals of the reported fitting. Right: distribution of radii of gyration for the best scoring 5 state model

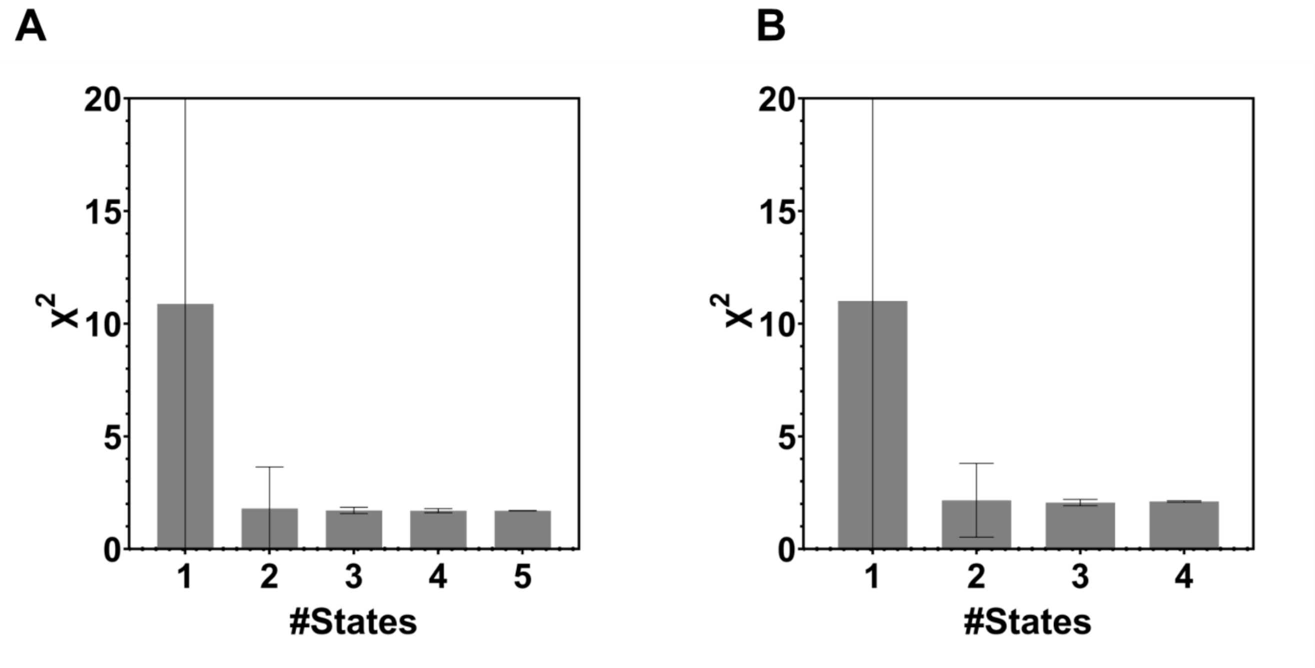


**Supplementary Figure 129 Histograms representation of the χ2 for two replicate of multi-state modelling performed through MultiFoXS using as an input the Alphafold prediction of the RAD51-BRC3-4 complex.**

A

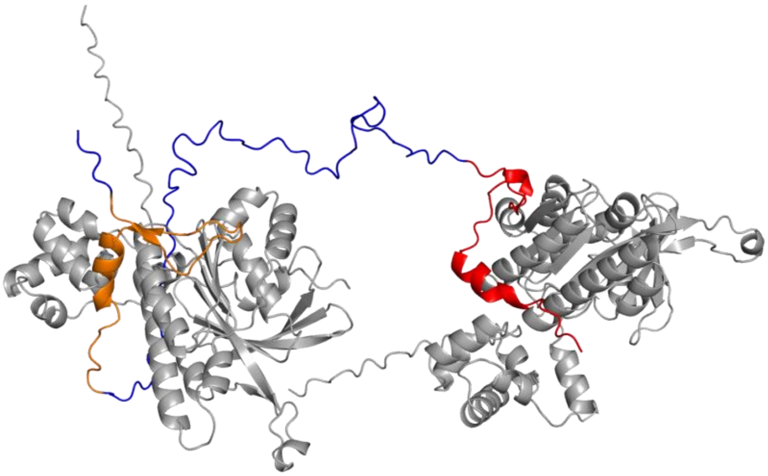

B

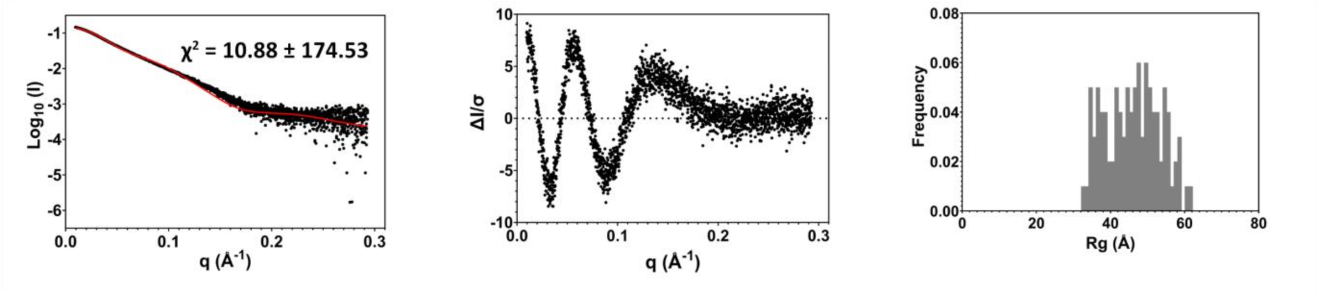


**Supplementary Figure 130 Multi-FoXS Modelling of Alphafold RAD51/BRC3-4 complex prediction model 1-state A.** Generated 1 state-model **B.** Left: fit to experimental SAXS data. Middle: normalized residuals of the reported fitting. Right: distribution of radii of gyration for the best scoring 1 state model

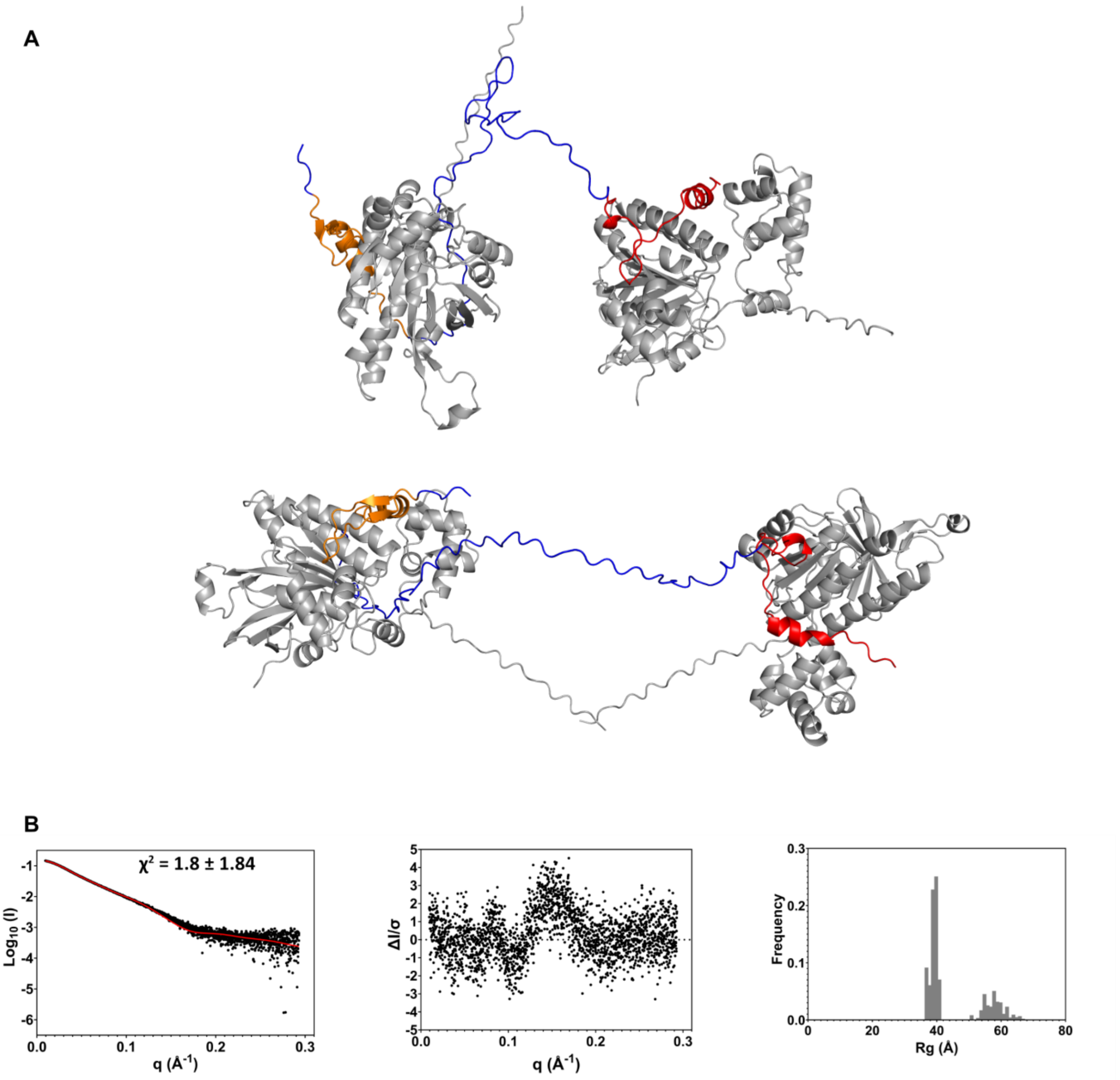


**Supplementary Figure 131 Multi-FoXS Modelling of Alphafold RAD51/BRC3-4 complex prediction model 2-state A.** Generated 2 state-model **B.** Left: fit to experimental SAXS data. Middle: normalized residuals of the reported fitting. Right: distribution of radii of gyration for the best scoring 2 state model

A

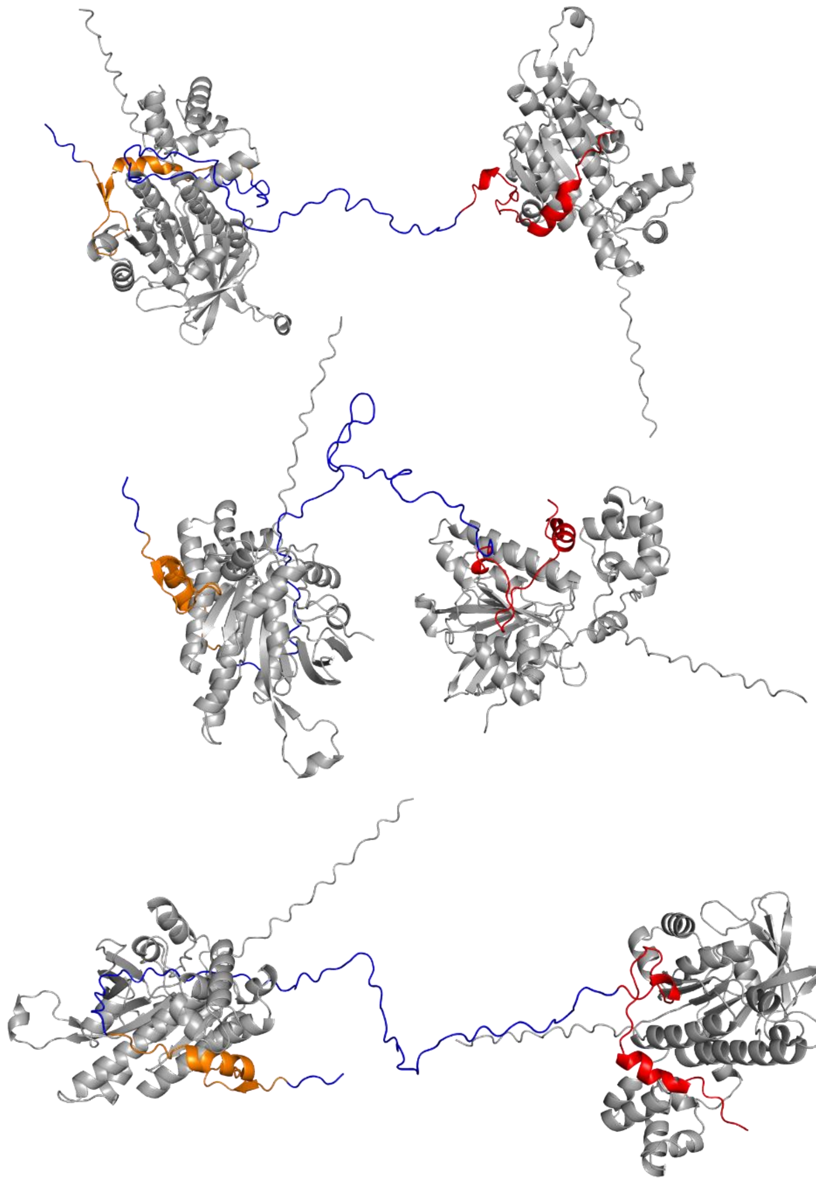

B

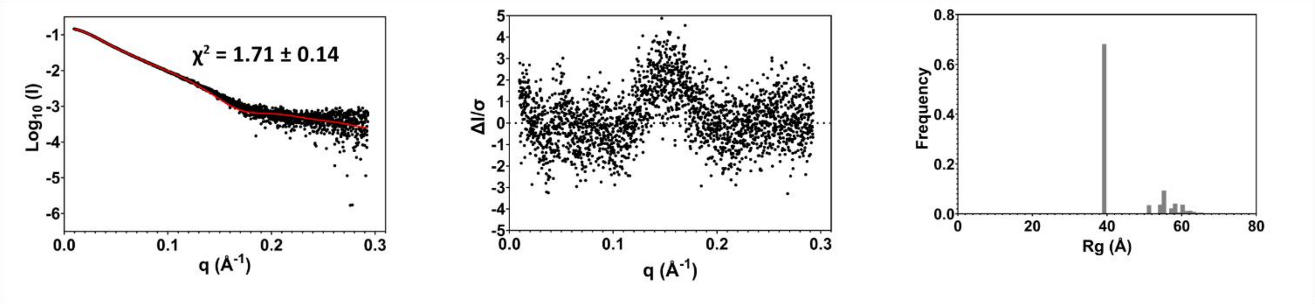


**Supplementary Figure 132 Multi-FoXS Modelling of Alphafold RAD51/BRC3-4 complex prediction model 3-state A.** Generated 3 state-model **B.** Left: fit to experimental SAXS data. Middle: normalized residuals of the reported fitting. Right: distribution of radii of gyration for the best scoring 3 state model

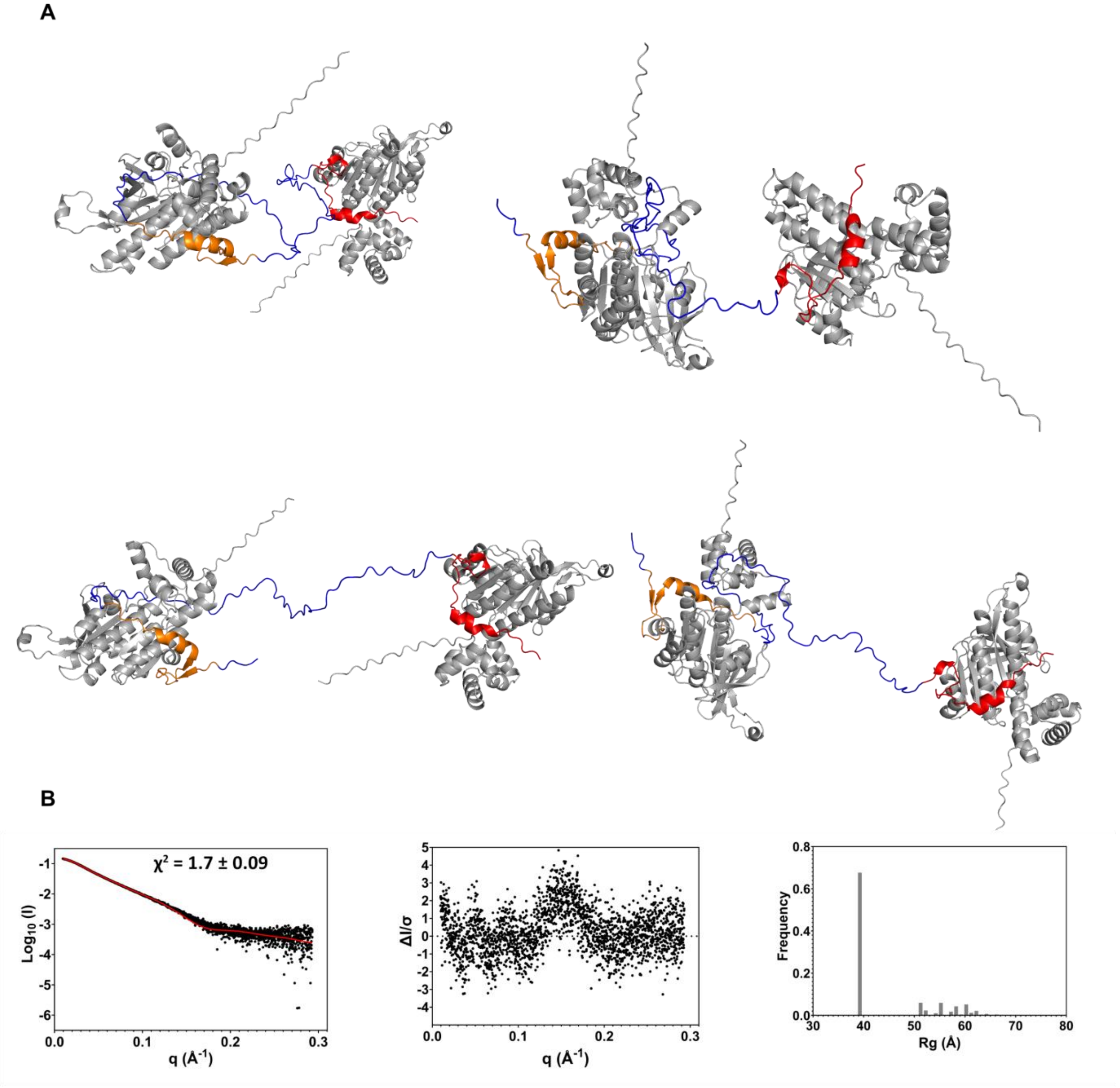


**Supplementary Figure 133 Multi-FoXS Modelling of Alphafold RAD51/BRC3-4 complex prediction model 4-state A.** Generated 4 state-model **B.** Left: fit to experimental SAXS data. Middle: normalized residuals of the reported fitting. Right: distribution of radii of gyration for the best scoring 4 state model

A

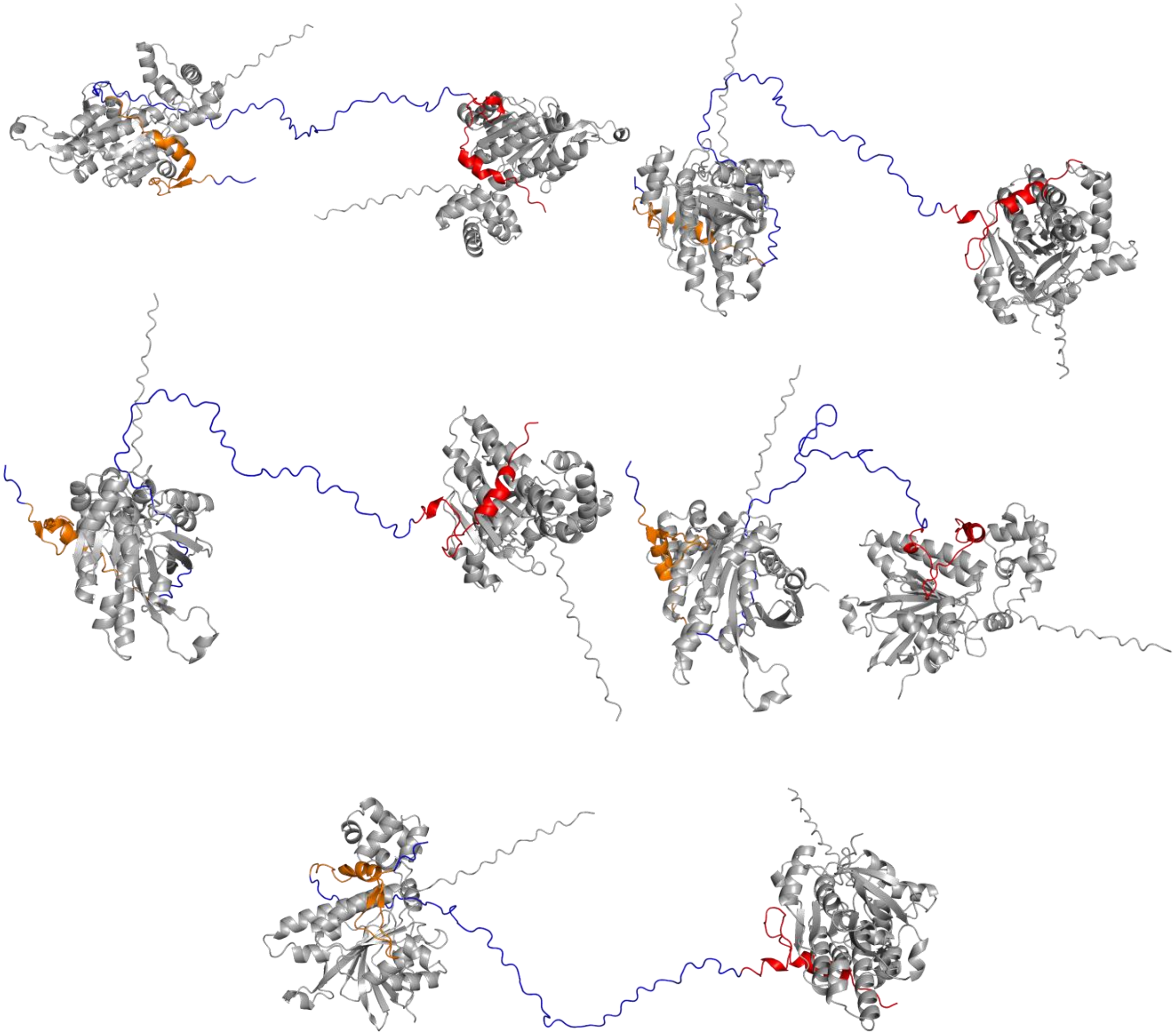

B

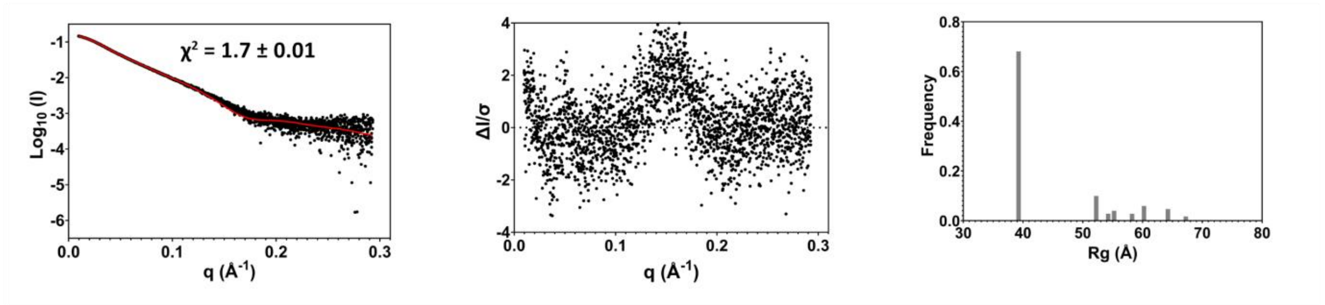


**Supplementary Figure 134 Multi-FoXS Modelling of Alphafold RAD51/BRC3-4 complex prediction model 5-state A.** Generated 5 state-model **B.** Left: fit to experimental SAXS data. Middle: normalized residuals of the reported fitting. Right: distribution of radii of gyration for the best scoring 5 state model

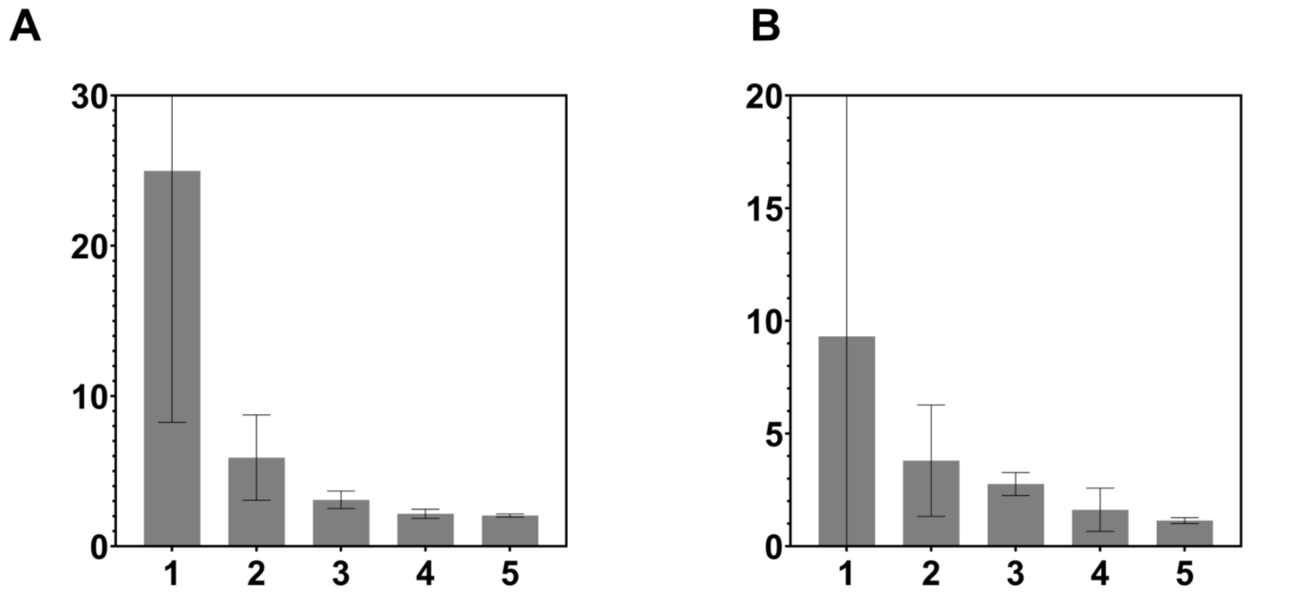


**Supplementary Figure 135** Histograms representation of the χ2 for two replicate of multi-state modelling performed through MultiFoXS using as an input the Alphafold prediction of the RAD51-BRC2-4 complex.

**A**

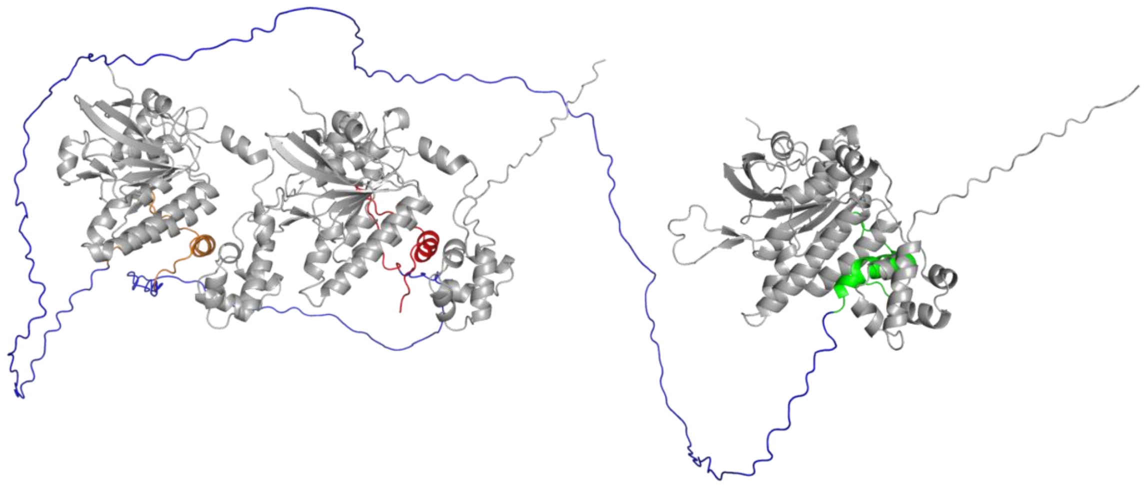

**B**

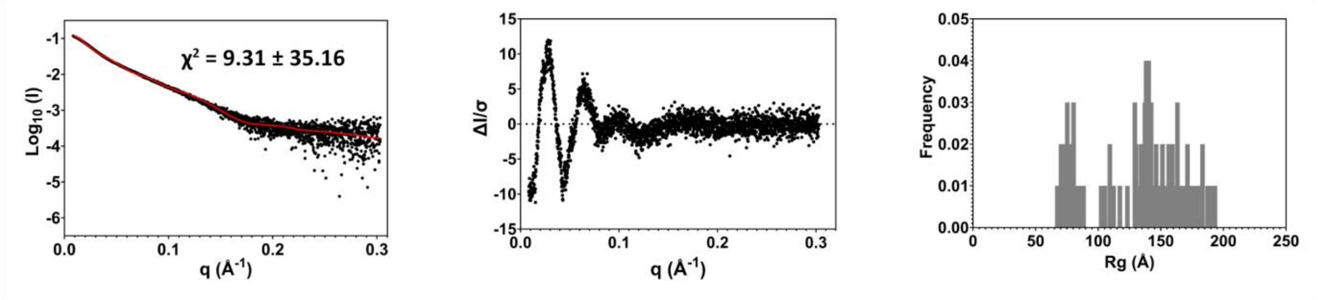


**Supplementary Figure 136 Multi-FoXS Modelling of Alphafold RAD51/BRC2-4 complex prediction model 1-state A.** Generated 1 state-model **B.** Left: fit to experimental SAXS data. Middle: normalized residuals of the reported fitting. Right: distribution of radii of gyration for the best scoring 1 state model

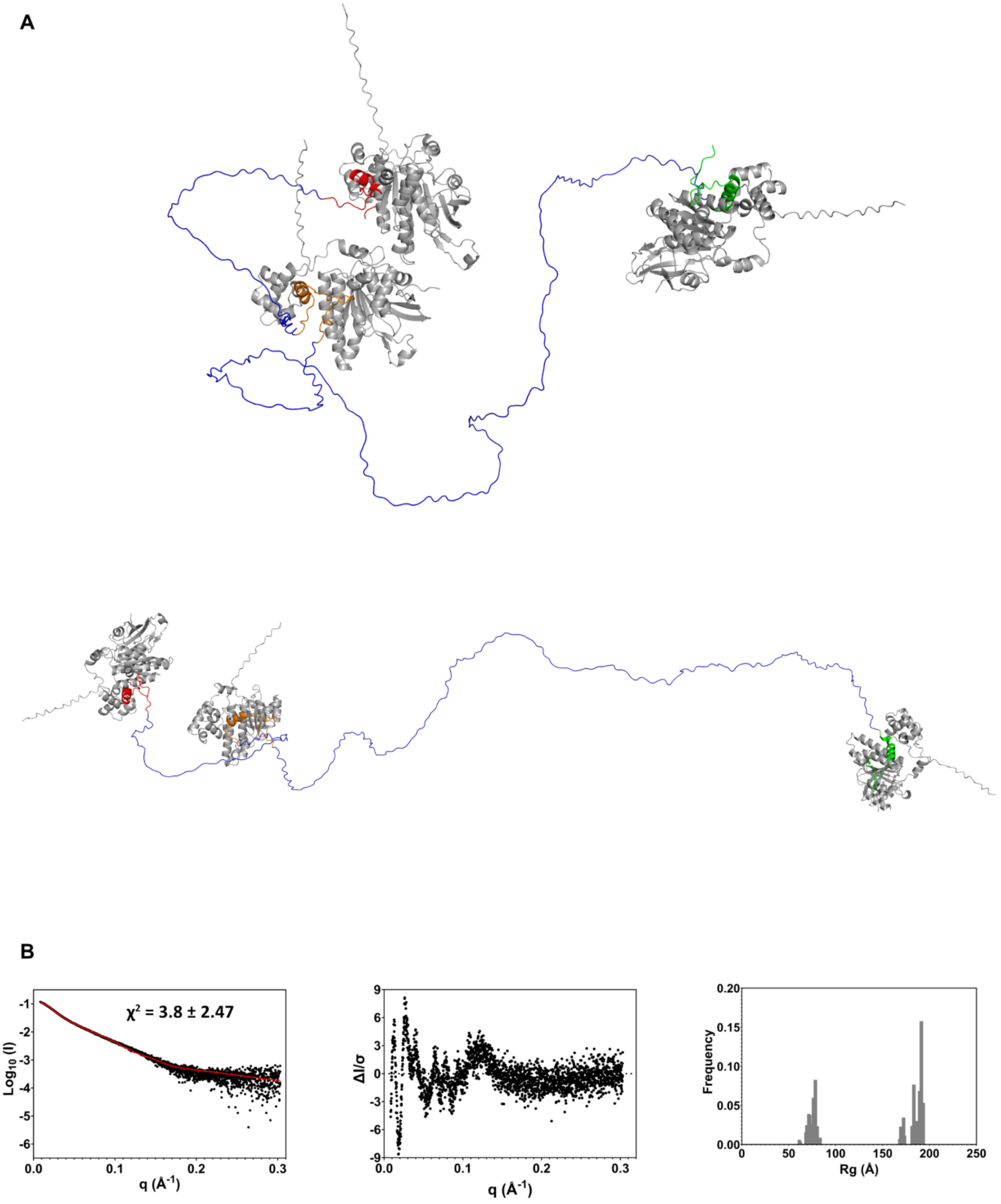


**Supplementary Figure 137 Multi-FoXS Modelling of Alphafold RAD51/BRC2-4 complex prediction model 2-state A.** Generated 2 state-model **B.** Left: fit to experimental SAXS data. Middle: normalized residuals of the reported fitting. Right: distribution of radii of gyration for the best scoring 2 state model

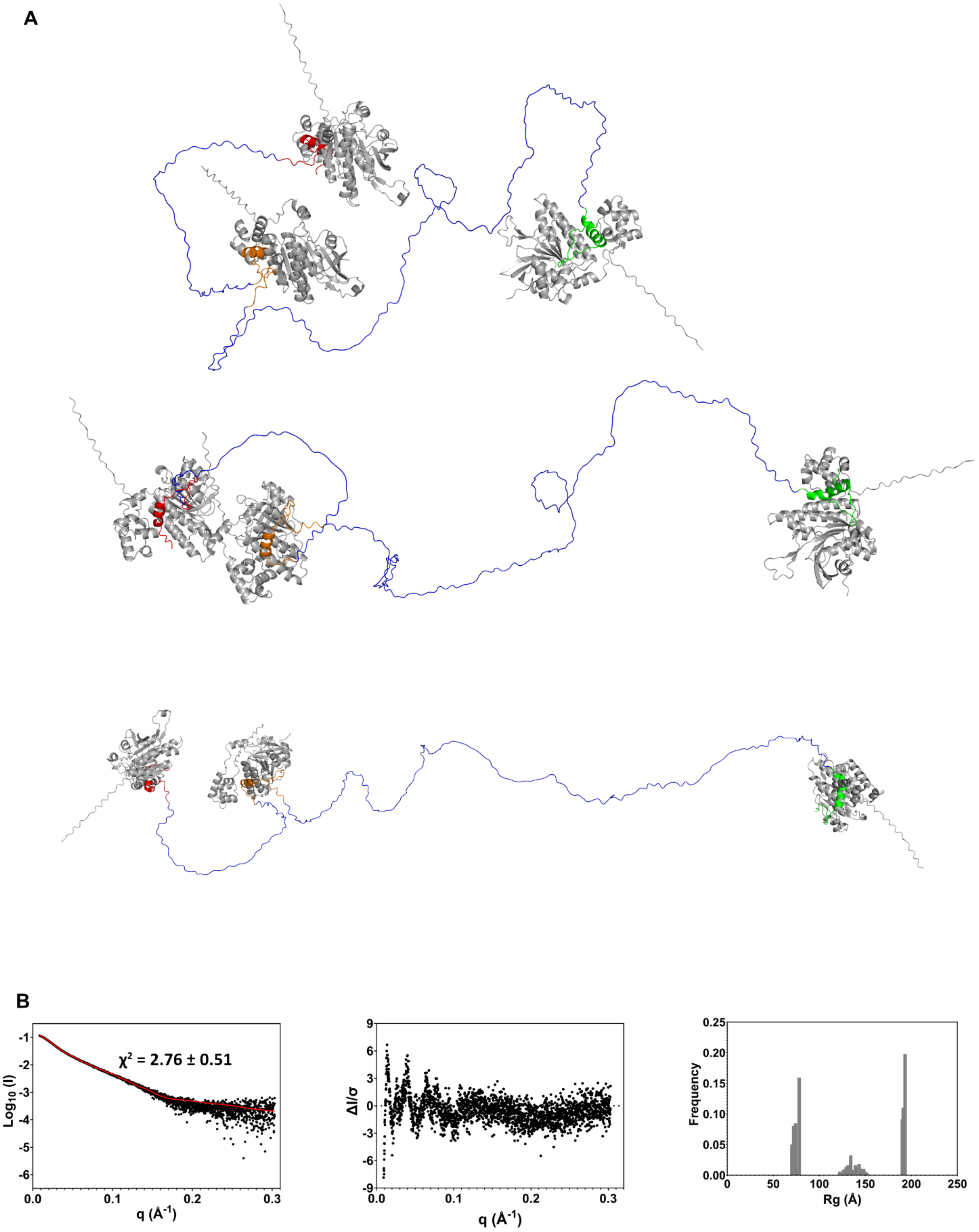


**Supplementary Figure 138 Multi-FoXS Modelling of Alphafold RAD51/BRC2-4 complex prediction model 3-state A.** Generated 3 state-model **B.** Left: fit to experimental SAXS data. Middle: normalized residuals of the reported fitting. Right: distribution of radii of gyration for the best scoring 3 state model

A

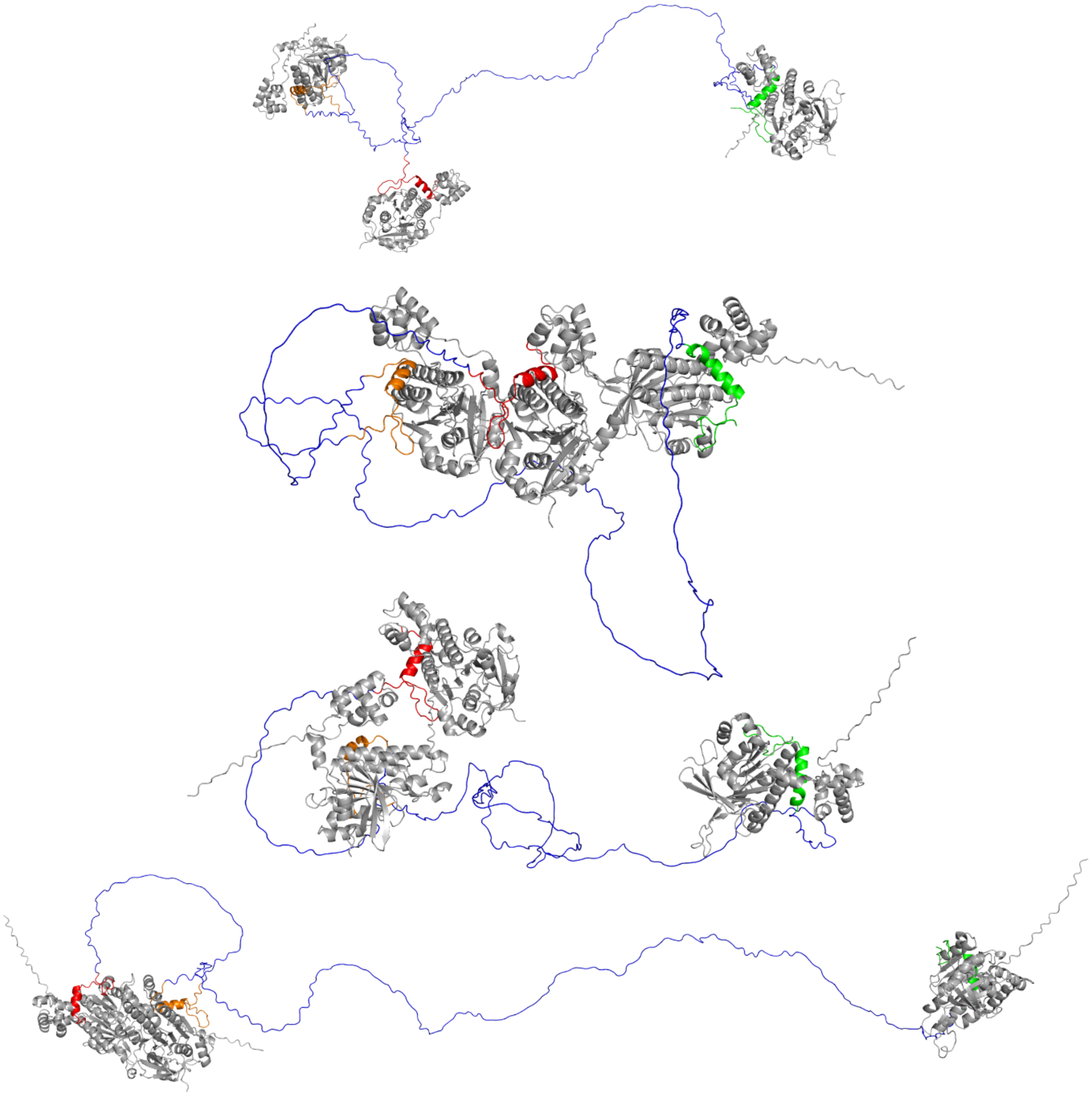

B

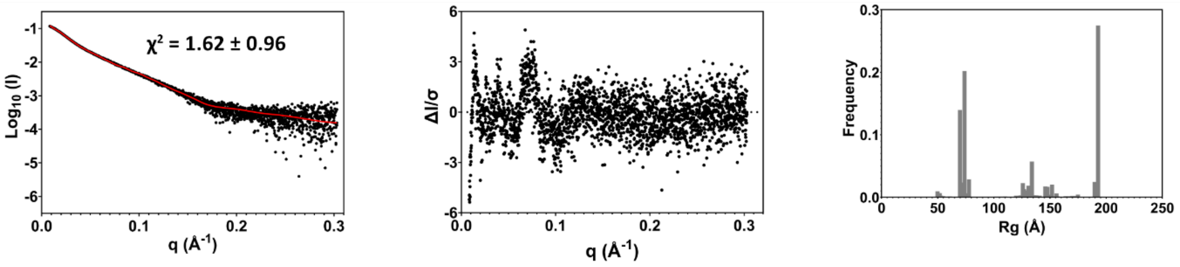


**Supplementary Figure 139 Multi-FoXS Modelling of Alphafold RAD51/BRC2-4 complex prediction model 4-state A.** Generated 4 state-model **B.** Left: fit to experimental SAXS data. Middle: normalized residuals of the reported fitting. Right: distribution of radii of gyration for the best scoring 4 state model

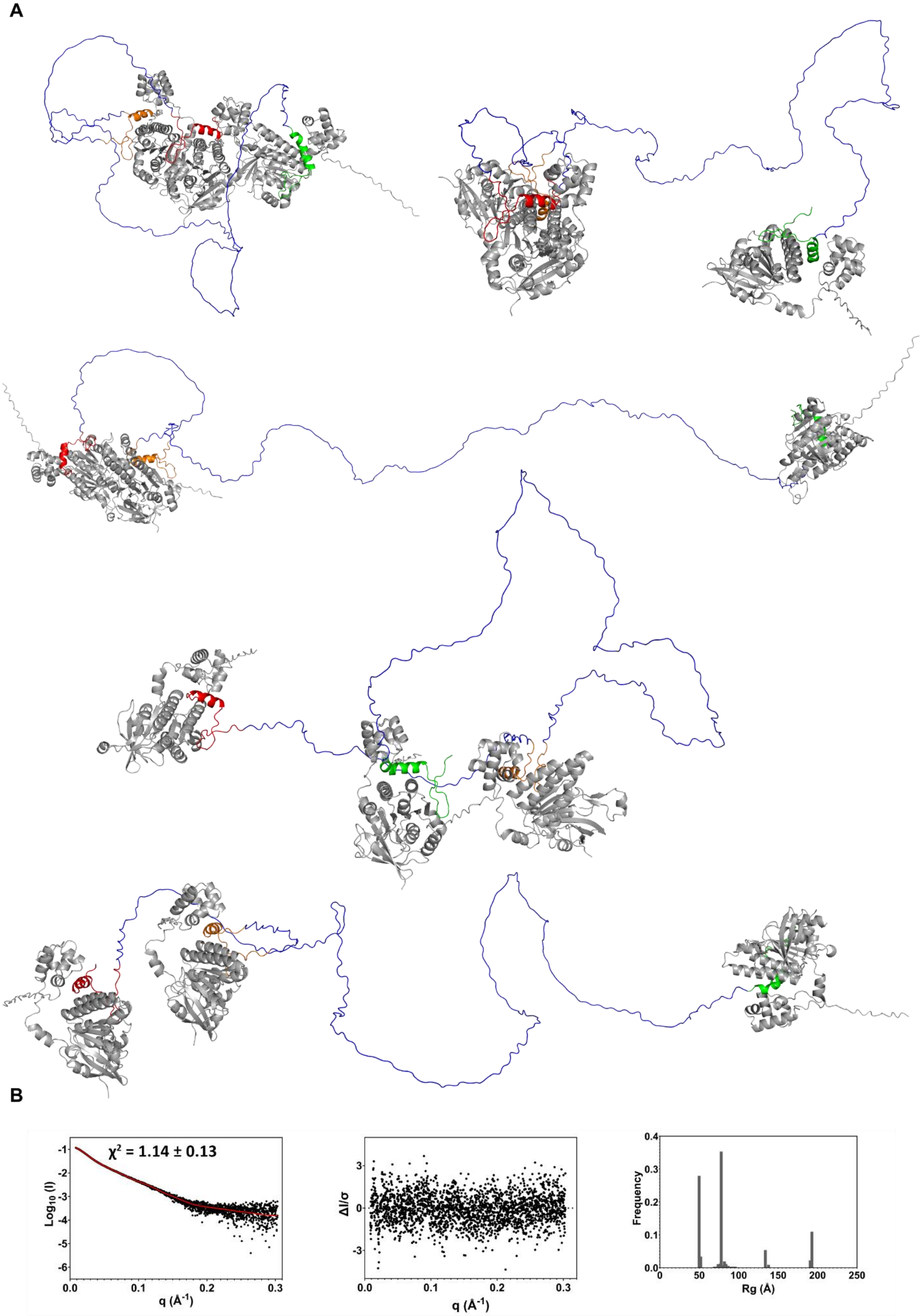


**Supplementary Figure 140 Multi-FoXS Modelling of Alphafold RAD51/BRC2-4 complex prediction model 5-state A.** Generated 5 state-model **B.** Left: fit to experimental SAXS data. Middle: normalized residuals of the reported fitting. Right: distribution of radii of gyration for the best scoring 5 state model

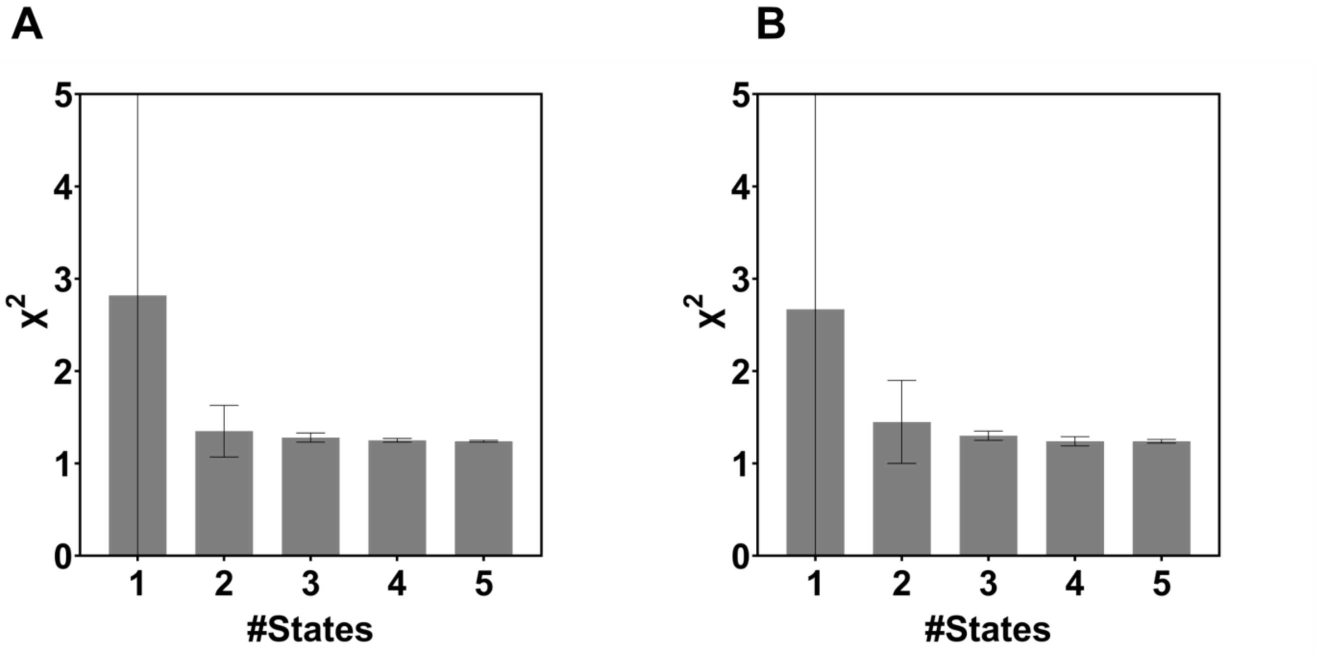


**Supplementary Figure 141 Histograms representation of the χ2 for two replicate of multi-state modelling performed through MultiFoXS using as an input the Alphafold prediction of the RAD51-BRC1-4 complex.**

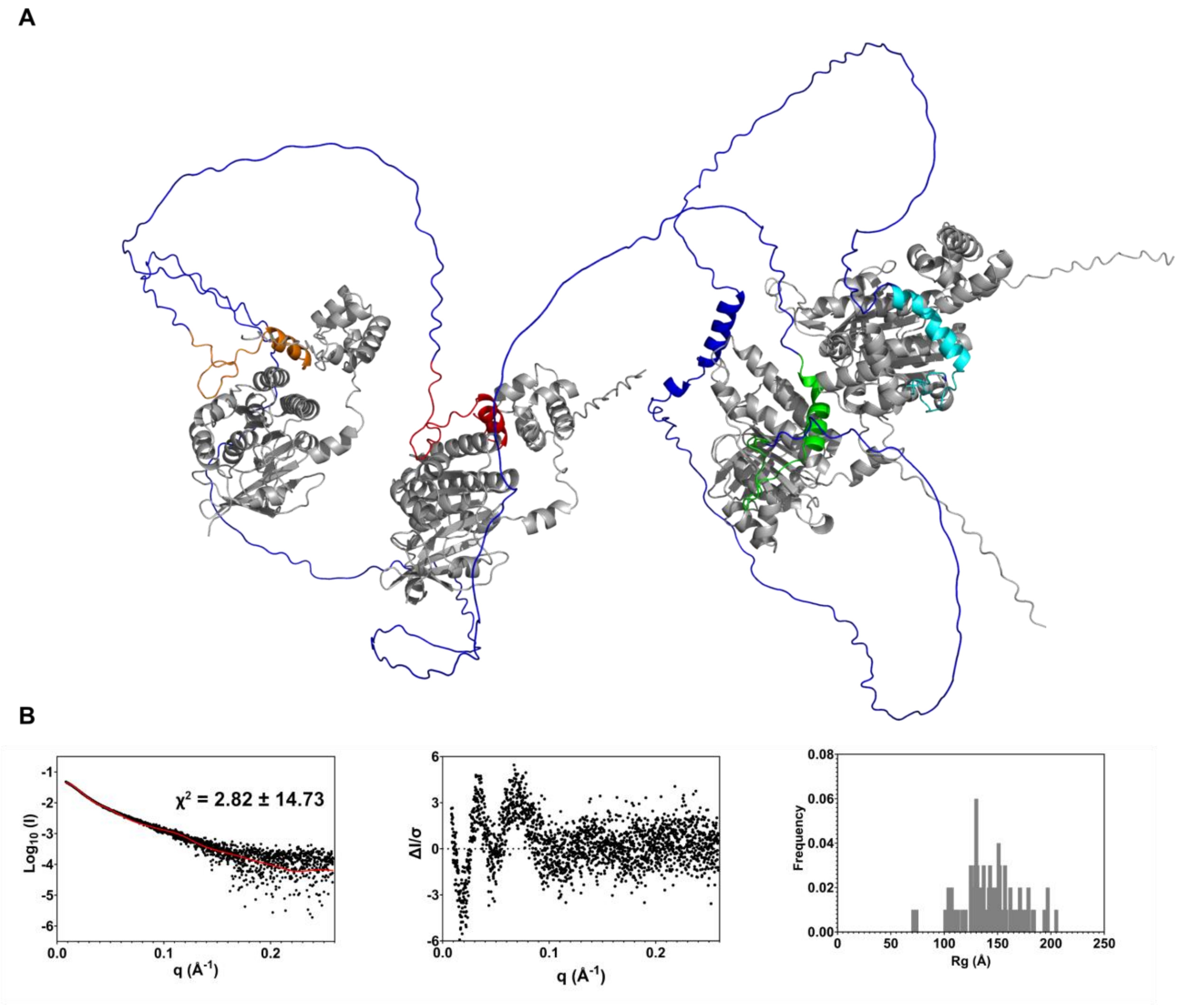


**Supplementary Figure 142 Multi-FoXS Modelling of Alphafold RAD51/BRC1-4 complex prediction model 1-state A.** Generated 1 state-model **B.** Left: fit to experimental SAXS data. Middle: normalized residuals of the reported fitting. Right: distribution of radii of gyration for the best scoring 1 state model

A

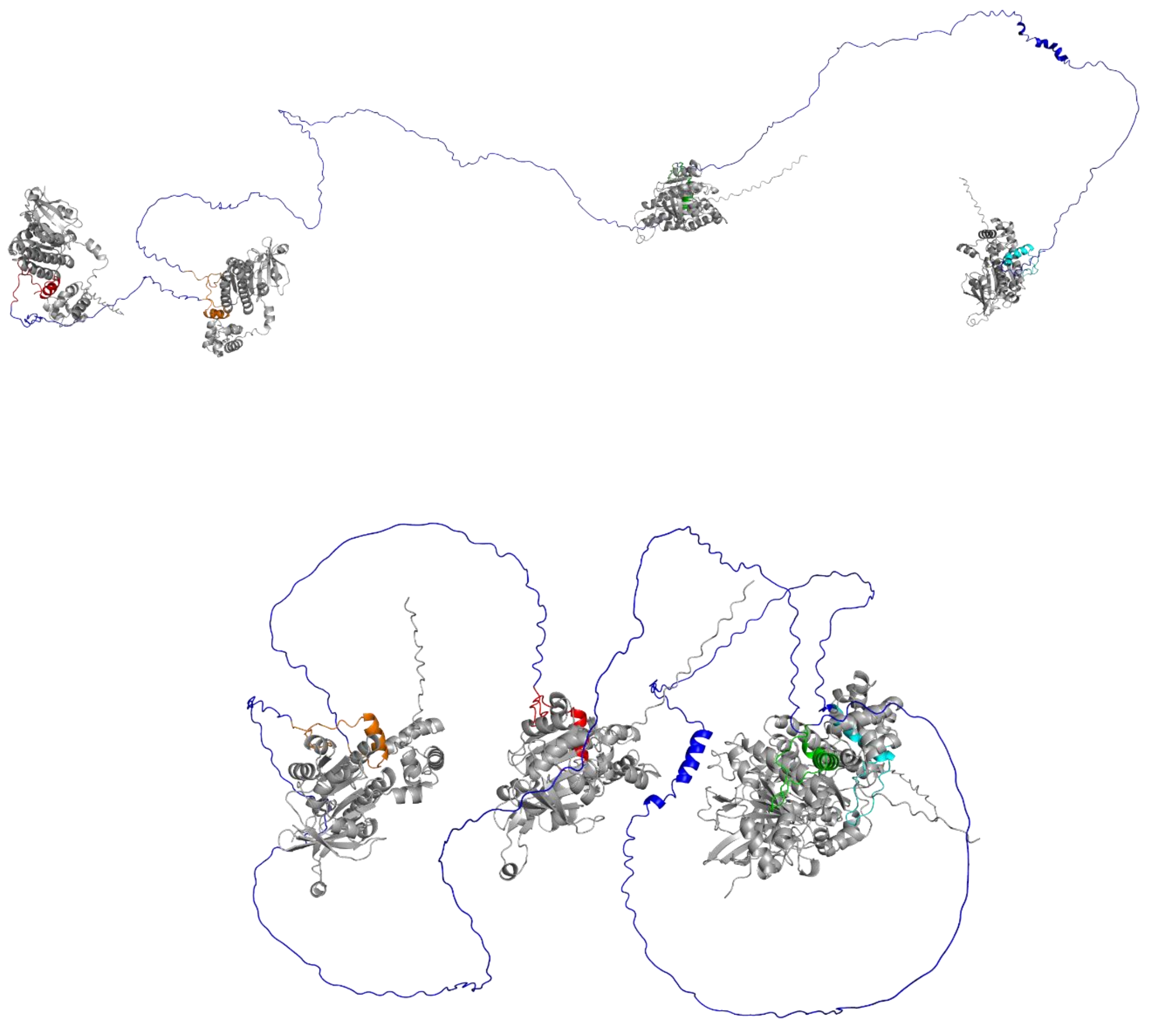

B

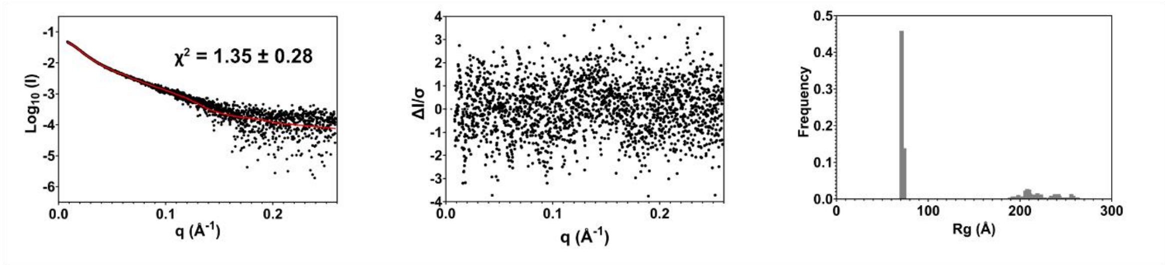


**Supplementary Figure 143 Multi-FoXS Modelling of Alphafold RAD51/BRC1-4 complex prediction model 2-state A.** Generated 2 state-model **B.** Left: fit to experimental SAXS data. Middle: normalized residuals of the reported fitting. Right: distribution of radii of gyration for the best scoring 2 state model

**A**

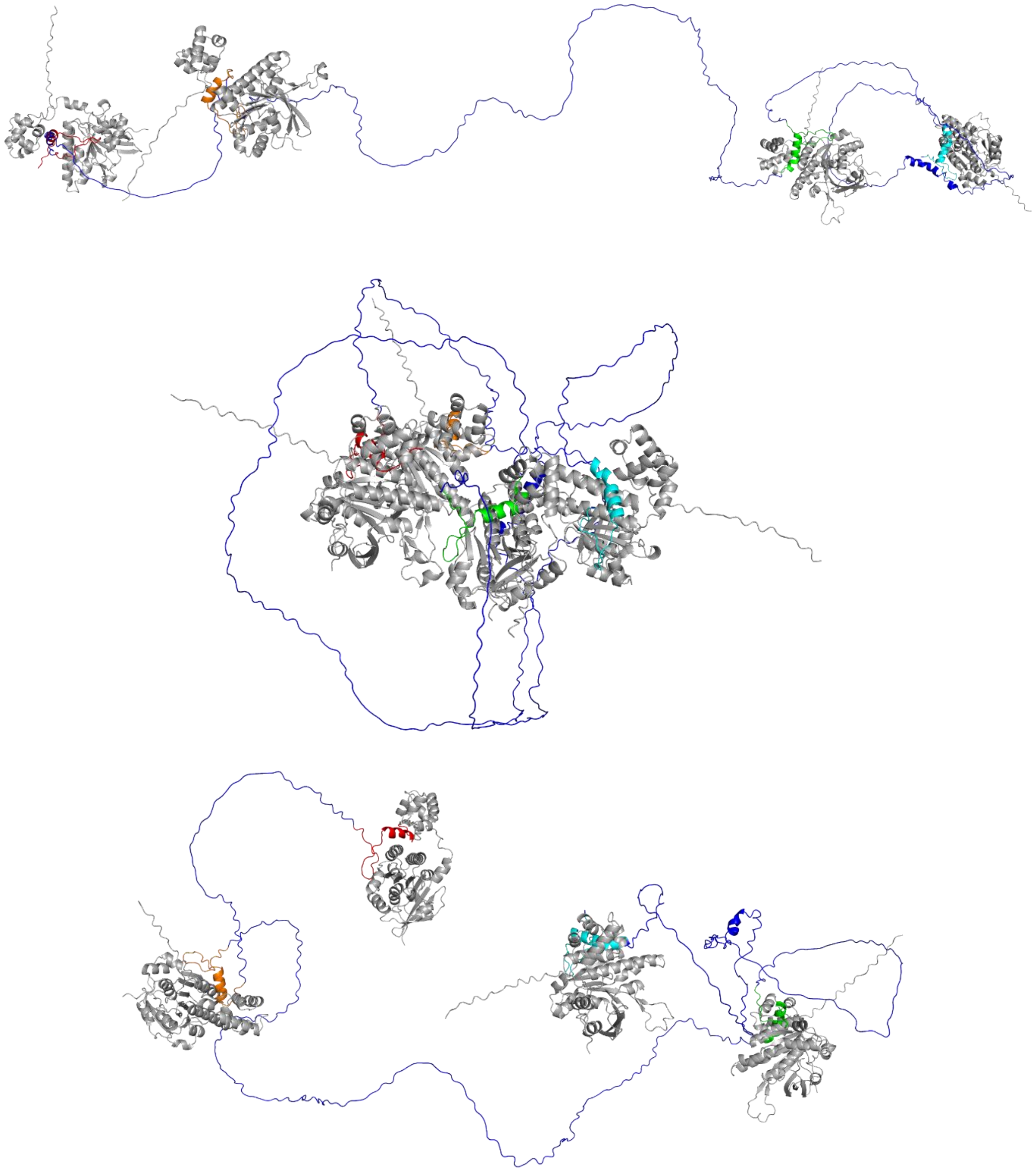

**B**

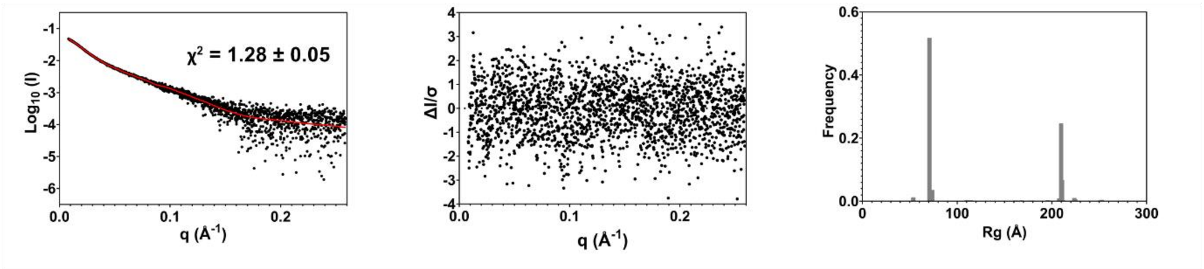


**Supplementary Figure 144 Multi-FoXS Modelling of Alphafold RAD51/BRC1-4 complex prediction model 3-state A.** Generated 3 state-model **B.** Left: fit to experimental SAXS data. Middle: normalized residuals of the reported fitting. Right: distribution of radii of gyration for the best scoring 3 state model

A

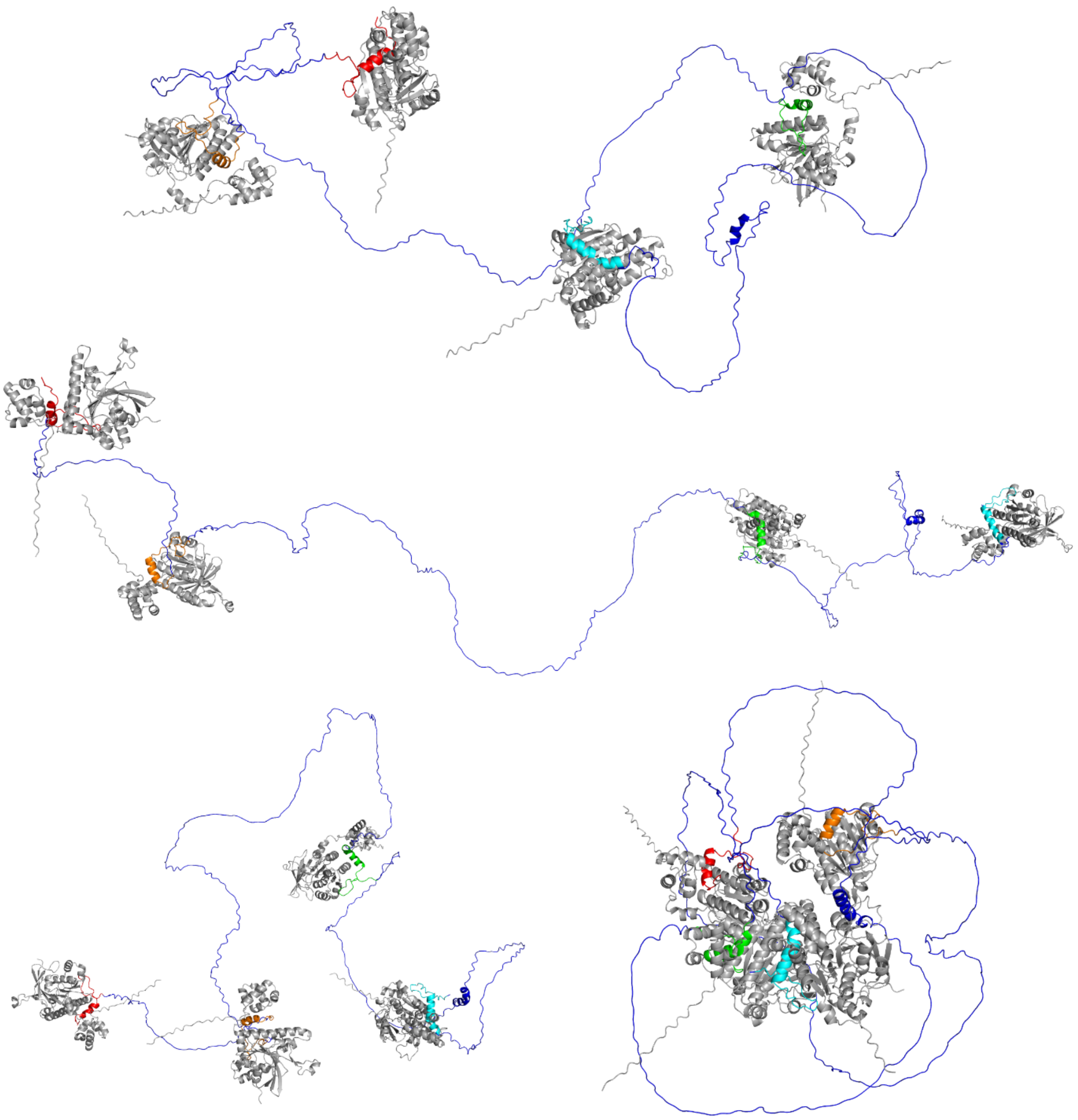

B

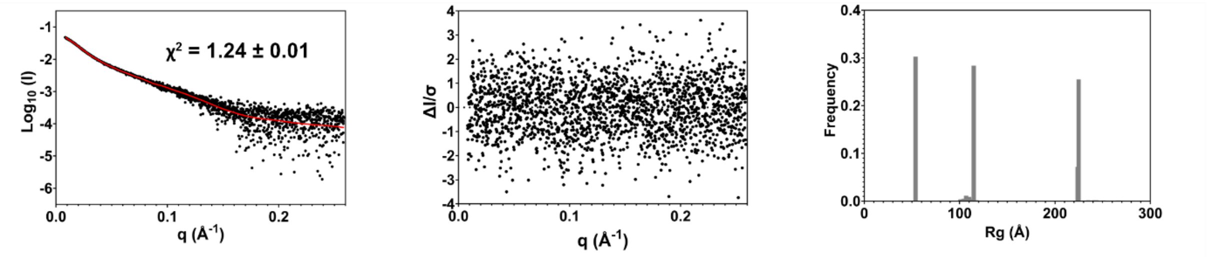


**Supplementary Figure 145 Multi-FoXS Modelling of Alphafold RAD51/BRC1-4 complex prediction model 4-state A.** Generated 3 state-model **B.** Left: fit to experimental SAXS data. Middle: normalized residuals of the reported fitting. Right: distribution of radii of gyration for the best scoring 4 state model

**Supplementary Table 11**

**SAXS Analysis N-Terminal truncated RAD51 (Δ97) in complex with fourth BRC repeat (BRC4)**

(a) Sample details

| | |
|---|---|
| Organism | Homo sapiens (Human) |
| Source | *E. Coli* expressed |
| Uniprot sequence ID | 98-339 Q06609 in complex with 1517-1551 P51587 |
| Extinction coefficient [A280, 0.1%(w/v)] ($M^{-1}$ $cm^{-1}$) | Ext. coefficient 19160<br>Abs 0.1% (=1 g/l) 0.619, assuming all C are cystines<br><br>Ext. coefficient 18910<br>Abs 0.1% (=1 g/l) 0.611, assuming all C are reduced |
| $\bar{v}$ from chemical composition ($cm^3$/ g) | 0.737 |
| Particle contrast from sequence and solvent constituents $\bar{\rho}$ (ρ protein - ρ solvent) $10^{-6}$ $Å^{-2}$ | 2.707 |
| M from chemical composition (Da) | 30945.47 |
| SEC–SAXS column | Superdex 200 Increase 3.2/300 |
| Loading concentration (mg/mL) | 6.6 |
| Injection Volume (mL) | 0.1 |
| Flow rate (mL/min) | 0.075 |
| Solvent (solvent blanks taken from SEC flow-through prior to elution of protein) | 20 mM $K_2HPO_4$/$KH_2PO_4$ pH 8, 100 mM NaCl, 200 mM $Li_2SO_4$, 1 mM DTT, 2% sucrose |

(b) SAXS data-collection parameters

| | |
|---|---|
| Instrument/data processing | Diamond Light Source B21 SAXS Beamline with EigerX 4M (Dectris) detector[1] |
| Wavelength (Å) | 0.9464 |
| Beam size at sample (mm) | 1.0 x 0.25 |
| Beam size at detector (focus) (mm) | 0.05 x 0.05 (FWHM) |
| Camera length (m) | 3.6883 |
| q measurement range (Å) | 0.0045-0.34 |
| Absolute scaling method | Absolute intensity scaled to water scatter at 0.0163 |
| Sample configuration | SEC–SAXS Dual Agilent 1260 HPLC system<br><br>Batch samples are loaded to Sample exposure unit through an Arinax BIOSAXS liquid sample-handling robot and measured in flow mode. |

| | |
|---|---|
| Sample temperature (°C) | 15 |
| Exposure time | SEC-SAXS Continuous 3 seconds exposure for 600 frames<br><br>Batch-SAXS 1 second exposure for 21 frames (sample has been moved during the exposure to attenuate radiation damage) |

(c) Structural parameters **SEC-SAXS**

| | |
|---|---|
| Guinier analysis | |
| I(0) ($cm^{-1}$) | 0.04 ± 2.62e-5 |
| Rg (Å) | 20.30 ± 0.03 |
| qmin ($Å^{-1}$) | 0.01352 |
| qmax ($Å^{-1}$) | 0.0641 |
| qRg min | 0.27 |
| qRg max | 1.3 |
| n min | 40 |
| n max | 427 |
| Coefficient of correlation, $R^2$ | 0.9974 |
| Molecular Weight | |
| $V_c$ (kDa) | 29.6 |
| $V_p$ (kDa) | 31.5 |
| Bayes (kDa) | 28.9 (28.6 – 32) |
| Shape&Size MW | 33.4 |
| Porod volume ($Å^{-3}$) | 19345 |
| P(r) analysis | |
| I(0) ($cm^{-1}$) | 0.04 ± 2.6e-5 |
| Rg (Å) | 20.18 ± 0.02 |
| Dmax (Å) | 66.0 |
| q range ($Å^{-1}$) | 0.013 - 0.328 |
| $\chi^2$ (total estimate from GNOM) | 1.11 |
| GNOM Interpretation | A good solution |
| Ambimeter | |
| Number of compatible shapes | 1 |
| Ambiguity score | 3.603E-02 |
| Ambimeter says | 3D reconstruction is potentially unique |
| Shanum | |
| Dmax (Å) | 65.98 |
| Smax | 0.33 |
| N Shannon channels | 6.89 |
| Optimal number of Shannon channels | 5 |
| Optimal value of q-vector | 0.23 |
| Last optimal point | 1759 |
| Atomistic Modelling | |
| Structure | AlphaFold Model |

| | |
|---|---|
| q range for modelling | 0.0134 - 0.2381 |
| FoXS | |
| $\chi^2$ | 1.15 |
| Predicted Rg (Å) | 18.78 |
| $c_1$, $c_2$ | 1.01, 1.41 |
| SASBDB IDs for data and models | *SASDWT5* |

(d) Structural Parameters **Batch SAXS**

| Protein storage buffer used for subtraction | 20 mM $K_2HPO_4$/$KH_2PO_4$ pH 8, 100 mM NaCl, 200 mM $Li_2SO_4$, 1 mM DTT, 5% Glycerol | | | |
|---|---|---|---|---|
| Protein Concentration | 1 mg/mL | 2 mg/mL | 4 mg/mL | 6 mg/mL |
| Guinier analysis | | | | |
| I(0) ($cm^{-1}$) | 0.02 ± 2.59e-4 | 0.0373 ± 2.75e-4 | 0.0743 ± 3.02e-4 | 0.1178 ± 3.38e-4 |
| Rg (Å) | 20.12 ± 0.56 | 20.62 ± 0.30 | 20.79 ± 0.16 | 21.23 ± 0.12 |
| qmin ($Å^{-1}$) | 0.00972 | 0.00972 | 0.00972 | 0.00972 |
| qmax ($Å^{-1}$) | 0.06463 | 0.0632 | 0.06268 | 0.06137 |
| qRg min | 0.20 | 0.20 | 0.20 | 0.21 |
| qRg max | 1.30 | 1.30 | 1.30 | 1.30 |
| n min | 40 | 40 | 40 | 40 |
| n max | 461 | 450 | 446 | 436 |
| $R^2$ | 0.9458 | 0.9782 | 0.99 | 0.99 |
| Molecular Weight | | | | |
| $V_c$ (kDa) | 31.1 | 28.0 | 24.7 | 29.4 |
| Bayes (kDa) | 28.9 (28.5-32) | 28.9 (25.9-29.2) | 27.6 (25.2-28.6) | 28.9 (27.9-32) |
| P(r) analysis | | | | |
| I(0) ($cm^{-1}$) | 0.02 | 0.04 | 0.07 | 0.12 |
| Rg (Å) | 20.08 ± 0.35 | 20.44 ± 0.24 | 20.56 ± 0.13 | 21.06 ± 0.1 |
| Dmax (Å) | 62.9 | 65.9 | 63.0 | 66.0 |
| q range ($Å^{-1}$) | 0.01-0.34 | 0.01-0.34 | 0.01-0.34 | 0.01-0.34 |
| $\chi^2$ | 0.046 | 0.058 | 0.052 | 0.062 |
| GNOM solution | good | good | good | good |
| SASBDB IDs | *SASDWU5* | *SASDWV5* | *SASDWW5* | *SASDWX5* |

**Aminoacidic Composition**

**>A**

**MEIIQITTGSKELDKLLQGGIETGSITEMFGEFRTGKTQICHTLAVTCQLPIDRGGGEGKAMYIDTEGTFRPERLLAVAER YGLSGSDVLDNVAYARAFNTDHQTQLLYQASAMMVESRYALLIVDSATALYRTDYSGRGELSARQMHLARFLRMLLR LADEFGVAVVITNQVVAQVDGAAMFAADPKKPIGGNIIAHASTTRLYLRKGRGETRICKIYDSPCLPEAEAMFAINAD GVGDAKD**

**>B**

**GWHMKEPTLLGFHTASGKKVKIAKESLDKVKNLFDEKEQ**

**Supplementary Table 12**

**SAXS Analysis RAD51 [F86E, A89E] in complex with fourth BRC repeat (BRC4)**

(a) Sample details

| | |
|---|---|
| Organism | Homo sapiens (Human) |
| Source | *E. Coli* expressed |
| Uniprot sequence ID | Q06609 in complex with 1517-1551 P51587<br>Q06609 mutations in position F86 and A89 with glutamic acid (E) |
| Extinction coefficient [A280, 0.1%(w/v)] | Ext. coefficient = 20650<br>Abs 0.1% (=1 g/l) = 0.497, assuming all C are cystines<br><br>Ext. coefficient 20400<br>Abs 0.1% (=1 g/l) 0.491, assuming all C are reduced |
| $\bar{v}$ from chemical composition of N-terminal domain ($cm^3$/ g) | 0.736 |
| Particle contrast from sequence and solvent constituents<br>$\bar{\rho}$ (ρ protein - ρ solvent) $10^{-6}$ $Å^{-2}$ | 2.734 |
| M from chemical composition (Da) | 41514.37 |
| SEC–SAXS column | Superdex 200 Increase 3.2/300 column |
| Loading concentration (mg/mL) | 11.3 |
| Injection Volume (mL) | 0.1 |
| Flow rate (mL/min) | 0.075 |
| Solvent (solvent blanks taken from SEC flow-through prior to elution of protein) | 20 mM $K_2HPO_4/KH_2PO_4$ pH 8, 100 mM NaCl, 200 mM $Li_2SO_4$, 1 mM DTT, 2% sucrose |

(b) SAXS data-collection parameters

| | |
|---|---|
| Instrument/data processing | Diamond Light Source B21 SAXS Beamline with EigerX 4M (Dectris) detector[1] |
| Wavelength (Å) | 0.9464 |
| Beam size at sample (mm) | 1.0 x 0.25 |
| Beam size at detector (focus) (mm) | 0.05 x 0.05 (FWHM) |
| Camera length (m) | 3.6883 |
| q measurement range (Å) | 0.0045-0.34 |
| Absolute scaling method | Absolute intensity scaled to water scatter at 0.0163 |
| Sample configuration | SEC–SAXS Dual Agilent 1260 HPLC system |
| Sample temperature (°C) | 15 |

| | |
|---|---|
| Exposure time | SEC-SAXS Continuous 3 seconds exposure for 600 frames<br><br>Batch-SAXS 1 second exposure for 21 frames (sample has been moved during the exposure to attenuate radiation damage) |

(c) Structural parameters **SEC SAXS**

| | |
|---|---|
| Guinier analysis | |
| I(0) (cm$^{-1}$) | 0.0972 ± 5.38e-5 |
| Rg (Å) | 27.82 ± 0.03 |
| qmin (Å$^{-1}$) | 0.00973 |
| qmax (Å$^{-1}$) | 0.03979 |
| qRg min | 0.2707 |
| qRg max | 1.1071 |
| n min | 40 |
| n max | 270 |
| Coefficient of correlation, $R^2$ | 0.9984 |
| Molecular Weight | |
| $V_c$ (kDa) | 40.3 |
| $V_p$ (kDa) | 47.6 |
| Bayes (kDa) | 43.8 (41.5 - 46.1) |
| Shape&Size MW | 47.9 |
| Porod volume (Å$^{-3}$) | 64109 |
| P(r) analysis | |
| I(0) (cm$^{-1}$) | 0.1 5.15e-5 |
| Rg (Å) | 28.37 0.04 |
| Dmax (Å) | 105.0 |
| q range (Å$^{-1}$) | 0.01- 0.287 |
| $\chi^2$ (total estimate from GNOM) | 1.09 |
| GNOM Interpretation | A reasonable solution |
| Ambimeter | |
| Number of compatible shapes | 11 |
| Ambiguity score | 1.916E-02 |
| Ambimeter says | 3D reconstruction is potentially unique |
| Shanum | |
| Dmax (Å) | 90.21 |
| Smax | 0.34 |
| N Shannon channels | 9.76 |
| Optimal number of Shannon channels | 8 |
| Optimal value of q-vector | 0.279 |
| Last optimal point | 2098 |
| Atomistic Modelling | |
| Structure | All the AlphaFold3 Models (5) generated through one job |
| q range for modelling | 0.0096 - 0.2381 |
| FoXS | |
| Model_0 | |

| | |
|---|---|
| χ2 | 1.89 |
| Predicted Rg (Å) | 27.42 |
| c1, c2 | 1.03, 0.27 |
| Model_1 | |
| χ2 | 9.82 |
| Predicted Rg (Å) | 25.56 |
| c1, c2 | 0.99,1.40 |
| Model_2 | |
| χ2 | 37.60 |
| Predicted Rg (Å) | 23.17 |
| c1, c2 | 0.99, 3.25 |
| Model_3 | |
| χ2 | 31.70 |
| Predicted Rg (Å) | 23.72 |
| c1, c2 | 0.99, 3.20 |
| Model_4 | |
| χ2 | 36.83 |
| Predicted Rg (Å) | 23.13 |
| c1, c2 | 0.99, 3.20 |
| Multistate/ensemble models | |
| Starting ensemble | All the AlphaFold3 Models (5) generated through one job |
| MultiFoXS | |
| No. of states | 2 |
| χ2 | 1.14 ± 0.22 |
| c1, c2 | 1.01, 1.1 |
| Rg values of each state (Å) | Model_0 (27.42), Model_3 (23.72) |
| Weights $w_n$ | $w_1$ (Model_0) = 0.836 $w_2$ (Model_3) = 0.164 |
| SASBDB IDs for data and models | *SASDWY5* |

(d) Structural parameters **Batch SAXS**

| Protein storage buffer used for subtraction | 20 mM $K_2HPO_4$/$KH_2PO_4$ pH 8, 100 mM NaCl, 200 mM $Li_2SO_4$, 1 mM DTT, 5% Glycerol | | | |
|---|---|---|---|---|
| Protein Concentration | 0.5 mg/mL | 1 mg/mL | 2 mg/mL | 4 mg/mL |
| Guinier analysis | | | | |
| I(0) ($cm^{-1}$) | 0.0119 ± 3.74e-4 | 0.0231 ± 3.72e-4 | 0.0479 ± 3.88e-4 | 0.0996 ± 4.44e-4 |
| Rg (Å) | 28.30 ± 1.59 | 27.36 ± 0.79 | 27.46 ± 0.39 | 27.63 ± 0.22 |
| qmin ($Å^{-1}$) | 0.00972 | 0.00972 | 0.00972 | 0.00972 |
| qmax ($Å^{-1}$) | 0.04585 | 0.04755 | 0.04755 | 0.04689 |
| qRg min | 0.2750 | 0.2658 | 0.2669 | 0.2685 |
| qRg max | 1.2974 | 1.3007 | 1.3058 | 1.2957 |
| n min | 40 | 40 | 40 | 40 |
| n max | 317 | 330 | 330 | 325 |
| $R^2$ | 0.8148 | 0.9630 | 0.9836 | 0.9962 |
| Molecular Weight | | | | |
| $V_c$ (kDa) | 37.2 | 29.5 | 43.5 | 43.7 |
| Bayes (kDa) | 38.5 (35.7-40.6) | 33.8 (32.6-38.1) | 44.7 (43.3-47.2) | 46.6 (45.2-50.3) |

| P(r) analysis | | | | |
|---|---|---|---|---|
| I(0) ($cm^{-1}$) | 0.01 3.63e-4 | 0.02 3.63e-4 | 0.05 4.64e-4 | 0.1 4.35e-4 |
| Rg (Å) | 30.22 ± 1.48 | 28.12 ± 0.83 | 28.35 ± 0.47 | 28.33 ± 0.26 |
| Dmax (Å) | 110.0 | 100.0 | 103.0 | 100.0 |
| q range ($Å^{-1}$) | 0.01- 0.282 | 0.01-0.292 | 0.01- 0.291 | 0.01 0.289 |
| $\chi^2$ | 0.06 | 0.042 | 0.066 | 0.046 |
| GNOM solution | Reasonable | Good | Reasonable | Reasonable |
| SASBDB IDs | *SASDWZ5* | *SASDW26* | *SASDW36* | *SASDW46* |

**Aminoacidic Composition**

**>A**

**GMAMQMQLEANADTSVEEESFGPQPISRLEQCGINANDVKKLEEAGFHTVEAVAYAPKKELINIKGISEAKADKILAE AAKLVPMGETTETEFHQRRSEIIQITTGSKELDKLLQGGIETGSITEMFGEFRTGKTQICHTLAVTCQLPIDRGGGEGKA MYIDTEGTFRPERLLAVAERYG**

**LSGSDVLDNVAYARAFNTDHQTQLLYQASAMMVESRYALLIVDSATALYRTDYSGRGELSARQMHLARFLRMLLRLA DEFGVAVVITNQVVAQVDGAAMFAADPKKPIGGNIIAHASTTRLYLRKGRGETRICKIYDSPCLPEAEAMFAINADGV GDAKD**

**>B**

**GWHMKEPTLLGFHTASGKKVKIAKESLDKVKNLFDEKEQ**

**Supplementary Table 13**

**SAXS Analysis RAD51 [F86E, A89E] in complex with second BRC repeat (BRC2)**

(a) Sample details

| | |
|---|---|
| Organism | Homo sapiens (Human) |
| Source | *E. Coli* expressed |
| Uniprot sequence ID | Q06609 in complex with 1212-1246 P51587<br>Q06609 mutations in position F86 and A89 with glutamic acid (E) |
| Extinction coefficient [$A_{280}$, 0.1%(w/v)] | Ext. coefficient = 22140<br>Abs 0.1% (=1 g/l) 0.534, assuming all C are cystines<br><br>Ext. coefficient = 21890<br>Abs 0.1% (=1 g/l) 0.528, assuming all C are reduced |
| $\bar{v}$ from chemical composition of N-terminal domain ($cm^3$/ g) | 0.734 |
| Particle contrast from sequence and solvent constituents<br>$\bar{\rho}$ (ρ protein - ρ solvent) $10^{-6}$ $Å^{-2}$ | 2.727 |
| M from chemical composition (Da) | 41428.11 |
| SEC–SAXS column | Superdex 200 Increase 3.2/300 column |
| Loading concentration (mg/mL) | 6.1 |

| | |
|---|---|
| Injection Volume (mL) | 0.1 |
| Flow rate (mL/min) | 0.075 |
| Solvent (solvent blanks taken from SEC flow-through prior to elution of protein) | 20 mM $K_2HPO_4/KH_2PO_4$ pH 8, 100 mM NaCl, 200 mM $Li_2SO_4$, 1 mM DTT, 2% sucrose |

(b) SAXS data-collection parameters

| | |
|---|---|
| Instrument/data processing | Diamond Light Source B21 SAXS Beamline with EigerX 4M (Dectris) detector[1] |
| Wavelength (Å) | 0.9464 |
| Beam size at sample (mm) | 1.0 x 0.25 |
| Beam size at detector (focus) (mm) | 0.05 x 0.05 (FWHM) |
| Camera length (m) | 3.6883 |
| q measurement range (Å) | 0.0045-0.34 |
| Absolute scaling method | Absolute intensity scaled to water scatter at 0.0163 |
| Sample configuration | SEC–SAXS Dual Agilent 1260 HPLC system |
| Sample temperature (°C) | 15 |
| Exposure time | SEC-SAXS Continuous 3 seconds exposure for 600 frames<br><br>Batch-SAXS 1 second exposure for 21 frames (sample has been moved during the exposure to attenuate radiation damage) |

(c) Structural parameters **SEC SAXS**

| | |
|---|---|
| Guinier analysis | |
| I(0) ($cm^{-1}$) | 0.0561 ± 5.36e-5 |
| $R_g$ (Å) | 27.2892 ± 0.07 |
| qmin ($Å^{-1}$) | 0.00973 |
| qmax ($Å^{-1}$) | 0.03757 |
| $qR_g$ min | 0.2655 |
| $qR_g$ max | 1.0252 |
| n min | 40 |
| n max | 253 |
| Coefficient of correlation, $R^2$ | 0.9943 |
| Molecular Weight | |
| $V_c$ (kDa) | 37.9 |
| $V_p$ (kDa) | 44.5 |
| Bayes (kDa) | 42.0 (39.8 - 44.2) |
| Shape&Size MW | 43.7 |
| Porod volume ($Å^{-3}$) | 51288 |
| P(r) analysis | |
| I(0) ($cm^{-1}$) | 0.06 ± 4.31e-5 |
| $R_g$ (Å) | 27.51 ± 0.04 |

| | |
|---|---|
| Dmax (Å) | 96.0 |
| q range (Å$^{-1}$) | 0.01-0.293 |
| $\chi^2$ (total estimate from GNOM) | 1.026 |
| GNOM Interpretation | A reasonable solution |
| Ambimeter | |
| Number of compatible shapes | 5 |
| Ambiguity score | 0.6990 |
| Ambimeter says | 3D reconstruction is potentially unique |
| Shanum | |
| Dmax (Å) | 82.37 |
| Smax | 0.34 |
| N Shannon channels | 8.91 |
| Optimal number of Shannon channels | 7 |
| Optimal value of q-vector | 0.27 |
| Last optimal point | 2009 |
| Atomistic Modelling | |
| Structure | All the AlphaFold3 Models (5) generated through one job |
| q range for modelling | 0.0096 - 0.2669 |
| FoXS | |
| Model_0 | |
| χ2 | 7.22 |
| Predicted Rg (Å) | 23.90 |
| c1, c2 | 0.99, 2.59 |
| Model_1 | |
| χ2 | 2.61 |
| Predicted Rg (Å) | 25.41 |
| c1, c2 | 1.00, 0.58 |
| Model_2 | |
| χ2 | 12.63 |
| Predicted Rg (Å) | 22.95 |
| c1, c2 | 0.99, 2.72 |
| Model_3 | |
| χ2 | 3.25 |
| Predicted Rg (Å) | 25.24 |
| c1, c2 | 0.99, 0.67 |
| Model_4 | |
| χ2 | 1.40 |
| Predicted Rg (Å) | 26.91 |
| c1, c2 | 1.04, -0.41 |
| Multistate/ensemble models | |
| Starting ensemble | All the AlphaFold3 Models (5) generated through one job |
| MultiFoXS | |
| No. of states | 2 |
| χ2 | 1.29 ± 0.02 |
| c1, c2 | 1.04, -0.4 |
| Rg values of each state (Å) | 26.91, 23.90 |

| | |
|---|---|
| Weights $w_n$ | $w_1$ (Model_4) = 0.924 $w_2$ (Model_0) = 0.076 |
| SASBDB IDs for data and models | *SASDW56* |

(d) Structural parameters **Batch SAXS**

| Protein storage buffer used for subtraction | | 20 mM $K_2HPO_4$/$KH_2PO_4$ pH 8, 100 mM NaCl, 200 mM $Li_2SO_4$, 1 mM DTT, 5% Glycerol | |
|---|---|---|---|
| Protein Concentration | 0.5 mg/mL | 1 mg/mL | 2 mg/mL |
| Guinier analysis | | | |
| I(0) ($cm^{-1}$) | 0.0107 ± 3.36e-4 | 0.0233 ± 3.53e-4 | 0.0505 ± 3.83e-4 |
| Rg (Å) | 26.1194 ± 1.48 | 26.6046 ± 0.72 | 27.0352 ± 0.36 |
| qmin ($Å^{-1}$) | 0.00972 | 0.00972 | 0.00972 |
| qmax ($Å^{-1}$) | 0.05002 | 0.04898 | 0.0482 |
| qRg min | 0.2538 | 0.2585 | 0.2627 |
| qRg max | 1.3066 | 1.3031 | 1.3031 |
| n min | 40 | 40 | 40 |
| n max | 349 | 341 | 335 |
| $R^2$ | 0.8526 | 0.9631 | 0.9881 |
| Molecular Weight | | | |
| $V_c$ (kDa) | 51.7 | 38.1 | 42.7 |
| Bayes (kDa) | 49.8 (49.2, 60.2) | 39.4 (37.3-41.5) | 47.7 (44.2-48.2) |
| P(r) analysis | | | |
| I(0) ($cm^{-1}$) | 0.01 ± 3.67e-4 | 0.02 ± 3.96e-4 | 0.05 ± 0.41 |
| Rg (Å) | 27.32 ± 1.42 | 27.58 ± 0.74 | 27.82 ± 0.41 |
| Dmax (Å) | 95.0 | 96.0 | 98.0 |
| q range ($Å^{-1}$) | 0.01 - 0.306 | 0.01 – 0.3 | 0.01 – 0.296 |
| $\chi^2$ | 0.044 | 0.04 | 0.05 |
| GNOM solution | Good | Good | Reasonable |
| SASBDB IDs | *SASDW66* | *SASDW76* | *SASDW86* |

**Aminoacidic Composition**

**>A**

**GMAMQMQLEANADTSVEEESFGPQPISRLEQCGINANDVKKLEEAGFHTVEAVAYAPKKELINIKGISEAKADKILAE AAKLVPMGETTETEFHQRRSEIIQITTGSKELDKLLQGGIETGSITEMFGEFRTGKTQICHTLAVTCQLPIDRGGGEGKA MYIDTEGTFRPERLLAVAERYGLSGSDVLDNVAYARAFNTDHQTQLLYQASAMMVESRYALLIVDSATALYRTDYSGR GELSARQMHLARFLRMLLRLADEFGVAVVITNQVVAQVDGAAMFAADPKKPIGGNIIAHASTTRLYLRKGRGETRIC KIYDSPCLPEAEAMFAINADGVGDAKD**

**>B**

**GWHMNEVGFRGFYSAHGTKLNVSTEALQKAVKLFSDIEN**

**Supplementary Table 14**

**SAXS Analysis RAD51 [F86E, A89E] in complex with BRCA2 truncation containing the first and the second BRC repeats (BRC1-2)**

(a) Sample details

| | |
|---|---|
| Organism | Homo sapiens (Human) |
| Source | *E. Coli* expressed |
| Uniprot sequence ID | Q06609 in complex with 1002-1246 P51587<br>Q06609 mutations in position F86 and A89 with glutamic acid (E) |
| Extinction coefficient [A280, 0.1%(w/v)] | Ext. coefficient = 36635<br>Abs 0.1% (=1 g/l) 0.360, assuming all C form cystines<br><br>Ext. coefficient = 35760<br>Abs 0.1% (=1 g/l) 0.352, assuming all C are reduced |
| $\bar{v}$ from chemical composition of N-terminal domain ($cm^3$/ g) | 0.733 |
| Particle contrast from sequence and solvent constituents<br>$\bar{\rho}$ (ρ protein - ρ solvent) $10^{-6}$ $Å^{-2}$ | 2.763 |
| M from chemical composition (Da) | 101721.46 |
| SEC–SAXS column | Superdex 200 Increase 3.2/300 column |
| Loading concentration (mg/mL) | 6.3 |
| Injection Volume (mL) | 0.1 |
| Flow rate (mL/min) | 0.075 |
| Solvent (solvent blanks taken from SEC flow-through prior to elution of protein) | 20 mM $K_2HPO_4$/$KH_2PO_4$ pH 8, 100 mM NaCl, 200 mM $Li_2SO_4$, 1 mM DTT, 2% sucrose |

(b) SAXS data-collection parameters

| | |
|---|---|
| Instrument/data processing | Diamond Light Source B21 SAXS Beamline with EigerX 4M (Dectris) detector[1] |
| Wavelength (Å) | 0.9464 |
| Beam size at sample (mm) | 1.0 x 0.25 |
| Beam size at detector (focus) (mm) | 0.05 x 0.05 (FWHM) |
| Camera length (m) | 3.6883 |
| q measurement range (Å) | 0.0045-0.34 |
| Absolute scaling method | Absolute intensity scaled to water scatter at 0.0163 |
| Sample configuration | SEC–SAXS Dual Agilent 1260 HPLC system |
| Sample temperature (°C) | 15 |

| | |
|---|---|
| Exposure time | SEC-SAXS Continuous 3 seconds exposure for 600 frames<br><br>Batch-SAXS 1 second exposure for 21 frames (sample has been moved during the exposure to attenuate radiation damage) |

(c) Structural parameters **SEC SAXS**

| | |
|---|---|
| Guinier analysis | |
| I(0) ($cm^{-1}$) | 0.1 ± 1.82e-4 |
| Rg (Å) | 52.54 ± 0.2 |
| qmin ($Å^{-1}$) | 0.00973 |
| qmax ($Å^{-1}$) | 0.02058 |
| qRg min | 0.5111 |
| qRg max | 1.0811 |
| n min | 40 |
| n max | 123 |
| Coefficient of correlation, $R^2$ | 1.0 |
| Molecular Weight | |
| $V_c$ (kDa) | 98.9 |
| $V_p$ (kDa) | 137.3 |
| Bayes (kDa) | 113.7 (106.9-121.5) |
| Shape&Size MW | 124.6 |
| Porod volume ($Å^{-3}$) | 243950 |
| P(r) analysis | |
| I(0) ($cm^{-1}$) | 0.1 ± 1.69e-4 |
| Rg (Å) | 55.3 ± 0.2 |
| Dmax (Å) | 210.0 |
| q range ($Å^{-1}$) | 0.01 0.152 |
| $\chi^2$ (total estimate from GNOM) | 1.05 |
| GNOM Interpretation | A reasonable solution |
| Ambimeter | |
| Number of compatible shapes | 33 |
| Ambiguity score | 1.519 |
| Ambimeter says | 3D reconstruction might be ambiguous |
| Shanum | |
| Dmax (Å) | 184.44 |
| Smax | 0.34 |
| N Shannon channels | 19.96 |
| Optimal number of Shannon channels | 17 |
| Optimal value of q-vector | 0.289 |
| Last optimal point | 2182 |
| Atomistic Modelling | |
| Structure | AlphaFold 2.3 |
| q range for modelling | 0.0091 - 0.2896 |
| FoXS | |
| $\chi^2$ | 55.31 |

| | |
|---|---|
| Predicted Rg (Å) | 36.37 |
| $c_1$, $c_2$ | 0.99, 3.12 |
| Multistate/ensemble models | |
| Flexible residues | Chain A 40 - 211 |
| Rigid bodies connections | 220-223 A 191-194 B, 10-13A 191-194 C |
| MultiFoXS | Trial 1 |
| No. of states | 1 |
| χ2 | 1.84 ± 60.38 |
| $c_1$, $c_2$ | 1.84, -0.23 |
| Rg values of each state (Å) | 52.07 |
| Weights wn | $w_1$ = 1.0 |
| No. of states | 2 |
| χ2 | 1.62 ± 0.75 |
| $c_1$, $c_2$ | 1.01, 0.24 |
| Rg values of each state (Å) | 52.06, 149.364 |
| Weights wn | $w_1$ = 0.913, $w_2$ = 0.087 |
| No. of states | 3 |
| χ2 | 1.93 ± 0.15 |
| $c_1$, $c_2$ | 1.04, 0.18 |
| Rg values of each state (Å) | 51.38, 149.13, 64.50 |
| Weights $w_n$ | $w_1$ = 0.708, $w_2$ = 0.146, $w_3$ = 0.145 |
| No. of states | 4 |
| χ2 | 1.76 ± 0.03 |
| $c_1$, $c_2$ | 1.04, 0.27 |
| Rg values of each state (Å) | 51.38, 149.13, 64.50, 120.72 |
| Weights wn | $w_1$ = 0.696, $w_2$ = 0.089, $w_3$ = 0.139, $w_4$ = 0.075 |
| No. of states | 5 |
| χ2 | 1.73 ± 0.03 |
| $c_1$, $c_2$ | 1.04, 0.45 |
| Rg values of each state (Å) | 51.38, 167.19, 61.39, 136.11, 97.50 |
| Weights wn | $w_1$ = 0.666, $w_2$ = 0.079, $w_3$ = 0.128, $w_4$ = 0.074, $w_5$ = 0.053 |
| MultiFoXS | Trial 2 |
| No. of states | 1 |
| χ2 | 3.72 ± 63.19 |
| $c_1$, $c_2$ | 1.03, 0.46 |
| Rg values of each state (Å) | 50.697 |
| Weights wn | $w_1$ = 1 |
| No. of states | 2 |
| χ2 | 2.42 ± 0.77 |
| $c_1$, $c_2$ | 1.02, 0.76 |
| Rg values of each state (Å) | 50.697, 141.86 |
| Weights wn | $w_1$ = 0.807, $w_2$ = 0.193 |
| No. of states | 3 |
| χ2 | 1.32 ± 0.38 |
| c1, c2 | 1.01, 0.31 |
| Rg values of each state (Å) | 49.45, 171.60, 71.40 |

| | |
|---|---|
| Weights wn | $w_1$ = 0.814, $w_2$ = 0.055, $w_3$ = 0.131 |
| No. of states | 4 |
| χ2 | 1.28 ± 0.06 |
| c1, c2 | 1.0, 0.76 |
| Rg values of each state (Å) | 49.45, 171.60, 69.14, 118.75 |
| Weights wn | $w_1$ = 0.757, $w_2$ = 0.052, $w_3$ = 0.133, $w_4$ = 0.058 |
| SASBDB IDs for data and models | *SASDWJ6* |

(d) Structural parameters **Batch SAXS**

| Solvent | 20 mM $K_2HPO_4$/$KH_2PO_4$ pH 8, 100 mM NaCl, 200 mM $Li_2SO_4$, 1 mM DTT, 5% Glycerol | | |
|---|---|---|---|
| Protein Concentration | 0.5 mg/mL | 1 mg/mL | 2 mg/mL |
| Guinier analysis | | | |
| I(0) ($cm^{-1}$) | 0.0214 ± 1.33e-3 | 0.0541 ± 1.39e-3 | 0.1189 ± 1.56e-3 |
| Rg (Å) | 57.1406 ± 5.64 | 54.9040 ± 2.41 | 54.6535 ± 1.18 |
| qmin ($Å^{-1}$) | 0.00972 | 0.00972 | 0.01024 |
| qmax ($Å^{-1}$) | 0.02146 | 0.02146 | 0.02198 |
| qRg min | 0.5553 | 0.5335 | 0.5596 |
| qRg max | 1.2261 | 1.1781 | 1.2013 |
| n min | 40 | 40 | 44 |
| n max | 130 | 130 | 134 |
| $R^2$ | 0.84 | 0.96 | 0.99 |
| Molecular Weight | | | |
| $V_c$ (kDa) | 320.4 | 119.1 | 101.0 |
| Bayes (kDa) | 157.1 (151.4-194.9) | 130.9 (121.5-142.2) | 113.7 (106.9-121.5) |
| P(r) analysis | | | |
| I(0) ($cm^{-1}$) | 0.02 ± 1.39e-3 | 0.05 ± 1.45e-3 | 0.12 ± 1.88e-3 |
| Rg (Å) | 61.12 ± 6.81 | 57.7 ± 2.41 | 59.12 ± 1.85 |
| Dmax (Å) | 240.0 | 210.0 | 230.0 |
| q range ($Å^{-1}$) | 0.01 - 0.139 | 0.01 - 0.145 | 0.01 - 0.146 |
| $\chi^2$ | 0.045 | 0.05 | 0.048 |
| GNOM solution | Reasonable | Reasonable | Reasonable |
| SASBDB IDs | *SASDWK6* | *SASDWL6* | *SASDWM6* |

**Aminoacidic Composition**

**>A**

**GMAMQMQLEANADTSVEEESFGPQPISRLEQCGINANDVKKLEEAGFHTVEAVAYAPKKELINIKGISEAKADKILAE AAKLVPMGETTETEFHQRRSEIIQITTGSKELDKLLQGGIETGSITEMFGEFRTGKTQICHTLAVTCQLPIDRGGGEGKA MYIDTEGTFRPERLLAVAERYGLSGSDVLDNVAYARAFNTDHQTQLLYQASAMMVESRYALLIVDSATALYRTDYSGR GELSARQMHLARFLRMLLRLADEFGVAVVITNQVVAQVDGAAMFAADPKKPIGGNIIAHASTTRLYLRKGRGETRIC KIYDSPCLPEAEAMFAINADGVGDAKD**

**>B**

**GMAMQMQLEANADTSVEEESFGPQPISRLEQCGINANDVKKLEEAGFHTVEAVAYAPKKELINIKGISEAKADKILAE AAKLVPMGETTETEFHQRRSEIIQITTGSKELDKLLQGGIETGSITEMFGEFRTGKTQICHTLAVTCQLPIDRGGGEGKA MYIDTEGTFRPERLLAVAERYGLSGSDVLDNVAYARAFNTDHQTQLLYQASAMMVESRYALLIVDSATALYRTDYSGR GELSARQMHLARFLRMLLRLADEFGVAVVITNQVVAQVDGAAMFAADPKKPIGGNIIAHASTTRLYLRKGRGETRIC KIYDSPCLPEAEAMFAINADGVGDAKD**

**>C**

**MANEVGFRGFYSAHGTKLNVSTEALQKAVKLFSDIENISEETSAEVHPISLSSSKCHDSVVSMFKIENHNDKTVSEKNN KCQLILQNNIEMTTGTFVEEITENYKRNTENEDNKYTAASRNSHNLEFDGSDSSKNDTVCIHKDETDLLFTDQHNICLK LSGQFMKEGNTQIKEDLSDLTFLEVAKAQEACHGNTSNKEQLTATKTEQNIKDFETSDTFFQTASGKNISVAKESFNKIV NFFDQKPE**

**Supplementary Table 15**

**SAXS Analysis RAD51 [F86E, A89E] in complex with BRCA2 truncation containing the first and the second BRC repeats (BRC2-3)**

(a) Sample details

| | |
|---|---|
| Organism | Homo sapiens (Human) |
| Source | *E. Coli* expressed |
| Uniprot sequence ID | Q06609 in complex with 1212-1455 P51587<br>Q06609 mutations in position F86 and A89 with glutamic acid (E) |
| Extinction coefficient [A280, 0.1%(w/v)] | Ext. coefficient = 35145<br>Abs 0.1% (=1 g/l) 0.345, assuming C form cystines<br><br>Ext. coefficient 34270<br>Abs 0.1% (=1 g/l) 0.337, assuming all C are reduced |
| $\bar{v}$ from chemical composition of N-terminal domain ($cm^3/g$) | 0.73 |
| Particle contrast from sequence and solvent constituents<br>$\bar{\rho}$ (ρ protein - ρ solvent) $10^{-6}$ Å$^{-2}$ | 2.855 |
| M from chemical composition (Da) | 101765.85 |
| SEC–SAXS column | Superose 6 Increase 3.2/300 |
| Loading concentration (mg/mL) | 4.9 |
| Injection Volume (mL) | 0.1 |
| Flow rate (mL/min) | 0.075 |
| Solvent (solvent blanks taken from SEC flow-through prior to elution of protein) | 20 mM $K_2HPO_4$/$KH_2PO_4$ pH 8, 100 mM NaCl, 100 mM $Li_2SO_4$, 1 mM DTT, 1% sucrose |

(b) SAXS data-collection parameters

| | |
|---|---|
| Instrument/data processing | Diamond Light Source B21 SAXS Beamline with EigerX 4M (Dectris) detector[1] |
| Wavelength (Å) | 0.9464 |
| Beam size at sample (mm) | 1.0 x 0.25 |
| Beam size at detector (focus) (mm) | 0.05 x 0.05 (FWHM) |
| Camera length (m) | 3.6883 |
| q measurement range (Å) | 0.0045-0.34 |
| Absolute scaling method | Absolute intensity scaled to water scatter at 0.0163 |
| Sample configuration | SEC–SAXS Dual Agilent 1260 HPLC system |
| Sample temperature (°C) | 15 |
| Exposure time | SEC-SAXS Continuous 3 seconds exposure for 600 frames<br><br>Batch-SAXS 1 second exposure for 21 frames (sample has been moved during the exposure to attenuate radiation damage) |

(c) Structural parameters **SEC SAXS**

| | |
|---|---|
| Guinier analysis | |
| I(0) ($cm^{-1}$) | 0.11 ± 2.02e-4 |
| Rg (Å) | 54.39 ± 0.19 |
| qmin ($Å^{-1}$) | 0.0105 |
| qmax ($Å^{-1}$) | 0.02002 |
| qRg min | 0.5711 |
| qRg max | 1.0891 |
| n min | 46 |
| n max | 119 |
| Coefficient of correlation, $R^2$ | 1.0 |
| Molecular Weight | |
| $V_c$ (kDa) | 93.9 |
| $V_p$ (kDa) | 138.4 |
| Bayes (kDa) | 109.1 (102.9-116.0) |
| Shape&Size MW | 123.9 |
| Porod volume ($Å^{-3}$) | 282933 |
| P(r) analysis | |
| I(0) ($cm^{-1}$) | 0.11 ± 2.11e-4 |
| Rg (Å) | 58.82 0.22 |
| Dmax (Å) | 230.0 |
| q range ($Å^{-1}$) | 0.01-0.147 |
| $\chi^2$ (total estimate from GNOM) | 1.065 |
| GNOM Interpretation | A reasonable solution |
| Ambimeter | |
| Number of compatible shapes | 4 |
| Ambiguity score | 0.6021 |
| Ambimeter says | 3D reconstruction is potentially unique |

| | |
|---|---|
| Shanum | |
| Dmax (Å) | 281.59 |
| Smax | 0.34 |
| N Shannon channels | 30.48 |
| Optimal number of Shannon channels | 28 |
| Optimal value of q-vector | 0.31 |
| Last optimal point | 2361 |
| Atomistic Modelling | |
| Structure | AlphaFold3 Model |
| q range for modelling | 0.0104 - 0.3123 |
| FoXS | |
| $\chi^2$ | 88.82 |
| Predicted Rg (Å) | 37.25 |
| $c_1$, $c_2$ | 0.99, -0.32 |
| Multistate/ensemble models | |
| Flexible residues | Chain C 40-211 |
| Rigid bodies connections | 220-223 C 191-194 B, 10-13 C 191-194 A |
| MultiFoXS | Trial 1 |
| No. of states | 1 |
| χ2 | 5.97 ± 57.22 |
| c1, c2 | 1.05, -0.5 |
| Rg values of each state (Å) | 56.68 |
| Weights wn | $w_1 = 1$ |
| No. of states | 2 |
| χ2 | 3.1 ± 2.31 |
| c1, c2 | 1.03, 0.53 |
| Rg values of each state (Å) | 56.675, 126.927 |
| Weights wn | $w_1 = 0.79$, $w_2 = 0.21$ |
| No. of states | 3 |
| χ2 | 1.65 ± 0.36 |
| c1, c2 | 1.02, 0.27 |
| Rg values of each state (Å) | 54.91, 133.131, 60.03, |
| Weights wn | $w_1 = 0.655$, $w_2 = 0.106$, $w_3 = 0.238$ |
| No. of states | 4 |
| χ2 | 1.5 ± 0.06 |
| c1, c2 | 1.02, 0.17 |
| Rg values of each state (Å) | 54.91, 139.474, 58.02, 106.472 |
| Weights wn | $w_1 = 0.637$, $w_2 = 0.05$, $w_3 = 0.247$, $w_4 = 0.065$ |
| No. of states | 5 |
| χ2 | 1.49 ± 0 |
| c1, c2 | 1.02, 0.27 |
| Rg values of each state (Å) | 54.91, 139.47, 58.02, 106.47, 60.03 |
| Weights wn | $w_1 = 0.639$, $w_2 = 0.055$, $w_3 = 0.173$, $w_4 = 0.062$, $w_5 = 0.071$ |
| MultiFoXS | Trial 2 |
| No. of states | 1 |
| χ2 | 7.54 ± 61.61 |

| | |
|---|---|
| c1, c2 | 1.02, -0.5 |
| Rg values of each state (Å) | 51.62 |
| Weights wn | $w_1$ = 1.0 |
| No. of states | 2 |
| χ2 | 4.55 ± 0.95 |
| c1, c2 | 0.99, 1.84 |
| Rg values of each state (Å) | 51.62, 125.99 |
| Weights wn | $w_1$ = 0.762, $w_2$ = 0.238 |
| No. of states | 3 |
| χ2 | 1.91 ± 1.04 |
| c1, c2 | 0.99, 0.76 |
| Rg values of each state (Å) | 48.62, 175.50, 83.31 |
| Weights wn | $w_1$ = 0.744, $w_2$ = 0.052, $w_3$ = 0.205 |
| No. of states | 4 |
| χ2 | 2.14 ± 0.1 |
| c1, c2 | 0.99, 1.98 |
| Rg values of each state (Å) | 51.18, 175.50, 66.09, 106.47 |
| Weights wn | $w_1$ = 0.589, $w_2$ = 0.141, $w_3$ = 0.173, $w_4$ = 0.097 |
| SASBDB IDs for data and models | *SASDW96* |

(d) Structural parameters **Batch SAXS**

| Protein storage buffer used for subtraction | 20 mM $K_2HPO_4$/$KH_2PO_4$ pH 8, 100 mM NaCl, 200 mM $Li_2SO_4$, 1 mM DTT, 5% Glycerol | | |
|---|---|---|---|
| Protein Concentration | 0.5 mg/mL | 1 mg/mL | 2 mg/mL |
| Guinier analysis | | | |
| I(0) ($cm^{-1}$) | 0.02 ± 1.36e-3 | 0.0471 ± 1.4e-3 | 0.1053 ± 1.55e-3 |
| Rg (Å) | 59.76 ± 5.57 | 57.38 ± 2.69 | 56.87 ± 1.34 |
| qmin ($Å^{-1}$) | 0.00972 | 0.00972 | 0.0105 |
| qmax ($Å^{-1}$) | 0.02146 | 0.02146 | 0.02146 |
| qRg min | 0.5807 | 0.5576 | 0.5526 |
| qRg max | 1.2823 | 1.2312 | 1.2202 |
| n min | 40 | 40 | 40 |
| n max | 130 | 130 | 130 |
| $R^2$ | 0.8797 | 0.9511 | 0.9846 |
| Molecular Weight | | | |
| $V_c$ (kDa) | 108.9 | 115.1 | 89.7 |
| Bayes (kDa) | 130.9 (121.5-134.3) | 130.9 (121.5-142.2) | 94.2 (92.7-106.9) |
| P(r) analysis | | | |
| I(0) ($cm^{-1}$) | 0.02 ± 1.06e-3 | 0.05 ± 1.51e-3 | 0.11 ± 1.46e-3 |
| Rg (Å) | 59.48 ± 4.14 | 61.45 ± 3.03 | 61.2 ± 1.23 |
| Dmax (Å) | 210.0 | 230.0 | 220.0 |
| q range ($Å^{-1}$) | 0.01-0.132 | 0.01-0.139 | 0.009-0.135 |
| $\chi^2$ | 0.047 | 0.065 | 0.077 |
| GNOM solution | Reasonable | Reasonable | Reasonable |
| SASBDB IDs | *SASDWA6* | *SASDWB6* | *SASDWC6* |

**Aminoacidic Composition**

**>A**
**GMAMQMQLEANADTSVEEESFGPQPISRLEQCGINANDVKKLEEAGFHTVEAVAYAPKKELINIKGISEAKADKILAE AAKLVPMGETTETEFHQRRSEIIQITTGSKELDKLLQGGIETGSITEMFGEFRTGKTQICHTLAVTCQLPIDRGGGEGKA MYIDTEGTFRPERLLAVAERYGLSGSDVLDNVAYARAFNTDHQTQLLYQASAMMVESRYALLIVDSATALYRTDYSGR GELSARQMHLARFLRMLLRLADEFGVAVVITNQVVAQVDGAAMFAADPKKPIGGNIIAHASTTRLYLRKGRGETRIC KIYDSPCLPEAEAMFAINADGVGDAKD**

**>B**

**GMAMQMQLEANADTSVEEESFGPQPISRLEQCGINANDVKKLEEAGFHTVEAVAYAPKKELINIKGISEAKADKILAE AAKLVPMGETTETEFHQRRSEIIQITTGSKELDKLLQGGIETGSITEMFGEFRTGKTQICHTLAVTCQLPIDRGGGEGKA MYIDTEGTFRPERLLAVAERYGLSGSDVLDNVAYARAFNTDHQTQLLYQASAMMVESRYALLIVDSATALYRTDYSGR GELSARQMHLARFLRMLLRLADEFGVAVVITNQVVAQVDGAAMFAADPKKPIGGNIIAHASTTRLYLRKGRGETRIC KIYDSPCLPEAEAMFAINADGVGDAKD**

**>C**

**MANHSFGGSFRTASNKEIKLSEHNIKKSKMFFKDIEEQYPTSLACVEIVNTLALDNQKKLSKPQSINTVSAHLQSSVVV SDCKNSHITPQMLFSKQDFNSNHNLTPSQKAEITELSTILEESGSQFEFTQFRKPSYILQKSTFEVPENQMTILKTTSEEC RDADLHVIMNAPSIGQVDSSKQFEGTVEIKRKFAGLLKNDCNKSASGYLTDENEVGFRGFYSAHGTKLNVSTEALQKA VKLFSDIEN**

**Supplementary Table 16**

**SAXS Analysis RAD51 [F86E, A89E] in complex with BRCA2 truncation containing the third and fourth BRC repeats (BRC3-4)**

(a) Sample details

| | |
|---|---|
| Organism | Homo sapiens (Human) |
| Source | *E. Coli* expressed |
| Uniprot sequence ID | Q06609 in complex with 1421-1551 P51587<br>Q06609 mutations in position F86 and A89 with glutamic acid (E) |
| Extinction coefficient [A280, 0.1%(w/v)] | Ext. coefficient = 37415<br>Abs 0.1% (=1 g/l) 0.417, assuming C are cystines<br><br>Ext. coefficient 36790<br>Abs 0.1% (=1 g/l) 0.410, assuming all C are reduced |
| v̄ from chemical composition (cm$^3$/ g) | 0.735 |
| Particle contrast from sequence and solvent constituents<br>ρ̄ (ρ protein - ρ solvent) $10^{-6}$ Å$^{-2}$ | 2.745 |

| | |
|---|---|
| M from chemical composition (Da) | 89663.96 |
| SEC–SAXS column | Superdex 200 Increase 3.2/300 column |
| Loading concentration (mg/mL) | 7.7 mg/ml |
| Injection Volume (mL) | 0.1 |
| Flow rate (mL/min) | 0.075 |
| Solvent (solvent blanks taken from SEC flow-through prior to elution of protein) | 20 mM $K_2HPO_4/KH_2PO_4$ pH 8, 100 mM NaCl, 200 mM $Li_2SO_4$, 1 mM DTT, 2% sucrose |

(b) SAXS data-collection parameters

| | |
|---|---|
| Instrument/data processing | Diamond Light Source B21 SAXS Beamline with EigerX 4M (Dectris) detector[1] |
| Wavelength (Å) | 0.9464 |
| Beam size at sample (mm) | 1.0 x 0.25 |
| Beam size at detector (focus) (mm) | 0.05 x 0.05 (FWHM) |
| Camera length (m) | 3.6883 |
| q measurement range (Å) | 0.0045-0.34 |
| Absolute scaling method | Absolute intensity scaled to water scatter at 0.0163 |
| Sample configuration | SEC–SAXS Dual Agilent 1260 HPLC system |
| Sample temperature (°C) | 15 |
| Exposure time | SEC-SAXS Continuous 3 seconds exposure for 600 frames<br><br>Batch-SAXS 1 second exposure for 21 frames (sample has been moved during the exposure to attenuate radiation damage) |

(c) Structural parameters **SEC-SAXS**

| | |
|---|---|
| Guinier analysis | |
| I(0) ($cm^{-1}$) | 0.1551 ± 1.54e-4 |
| Rg (Å) | 45.61 ± 0.10 |
| qmin ($Å^{-1}$) | 0.00973 |
| qmax ($Å^{-1}$) | 0.0228 |
| qRg min | 0.4437 |
| qRg max | 1.0398 |
| n min | 40 |
| n max | 140 |
| Coefficient of correlation, $R^2$ | 0.9979 |
| Molecular Weight | |
| $V_c$ (kDa) | 85.5 |
| $V_p$ (kDa) | 112.9 |
| Bayes (kDa) | 94.2 (89.7-95.8) |
| Shape&Size MW | 94.6 |

| | |
|---|---|
| Porod volume (Å$^{-3}$) | 194506 |
| P(r) analysis | |
| I(0) (cm$^{-1}$) | 0.16 |
| Rg (Å) | 47.51 0.09 |
| Dmax (Å) | 180.0 |
| q range (Å$^{-1}$) | 0.01 - 0.175 |
| $\chi^2$ (total estimate from GNOM) | 1.07 |
| GNOM Interpretation | A reasonable solution |
| Ambimeter | |
| Number of compatible shapes | 144 |
| Ambiguity score | 2.158 |
| Ambimeter says | 3D reconstruction might be ambiguous |
| Shanum | |
| Dmax (Å) | 182.31 |
| Smax | 0.34 |
| N Shannon channels | 19.73 |
| Optimal number of Shannon channels | 17 |
| Optimal value of q-vector | 0.292 |
| Last optimal point | 2208 |
| Atomistic Modelling | |
| Structure | AlphaFold3 Model |
| q range for modelling | 0.0096 - 0.2929 |
| FoXS | |
| $\chi^2$ | 144.73 |
| Predicted Rg (Å) | 33.01 |
| $c_1$, $c_2$ | 0.99, 2.03 |
| Multistate/ensemble models | |
| Flexible residues | Chain C residues 40 – 101 |
| Rigid bodies connections | |
| MultiFoXS | Trial 1 |
| No. of states | 1 |
| χ2 | 10.88 ± 174.53 |
| c1, c2 | 0.99, 1.72 |
| Rg values of each state (Å) | 41.94 |
| Weights wn | $w_1$ = 1.00 |
| No. of states | 2 |
| χ2 | 1.8 ± 1.84 |
| c1, c2 | 0.99, 1.98 |
| Rg values of each state (Å) | 39.24, 60.73 |
| Weights wn | $w_1$ = 0.676, $w_2$ = 0.206 |
| No. of states | 3 |
| χ2 | 1.71 ± 0.14 |
| c1, c2 | 0.99, 1.98 |
| Rg values of each state (Å) | 39.24, 60.73, 52.52 |
| Weights wn | $w_1$ = 0.676, $w_2$ = 0.206, $w_3$ = 0.118 |
| No. of states | 4 |

| | |
|---|---|
| χ2 | 1.7 ± 0.09 |
| c1, c2 | 0.99, 1.93 |
| Rg values of each state (Å) | 39.24, 60.73, 52.52, 58.70 |
| Weights wn | $w_1$ = 0.682, $w_2$ = 0.14, $w_3$ = 0.108, $w_4$ = 0.07 |
| No. of states | 5 |
| χ2 | 1.7 ± 0.01 |
| c1, c2 | 0.99, 1.96 |
| Rg values of each state (Å) | 39.24, 60.73, 52.52, 55.99, 67.83 |
| Weights wn | $w_1$ = 0.682, $w_2$ = 0.076, $w_3$ = 0.121, $w_4$ = 0.069 $w_5$ = 0.051 |
| MultiFoXS | Trial 2 |
| No. of states | 1 |
| χ2 | 11.01 ± 193.12 |
| c1, c2 | 0.99, 0.94 |
| Rg values of each state (Å) | 41.05 |
| Weights wn | $w_1$ = 1.0 |
| No. of states | 2 |
| χ2 | 2.16 ± 1.64 |
| c1, c2 | 0.99, 2.02 |
| Rg values of each state (Å) | 39.39, 59.77 |
| Weights wn | $w_1$ = 0.733, $w_2$ = 0.267 |
| No. of states | 3 |
| χ2 | 2.06 ± 0.14 |
| c1, c2 | 0.99, 2.02 |
| Rg values of each state (Å) | 39.39, 56.66, 65.27 |
| Weights wn | $w_1$ = 0.71, $w_2$ = 0.23, $w_3$ = 0.06 |
| No. of states | 4 |
| χ2 | 2.11 ± 0.03 |
| c1, c2 | 0.99, 2.02 |
| Rg values of each state (Å) | 39.39, 53.29, 63.42, 58.20 |
| Weights wn | $w_1$ =0.707, $w_2$ =0.096, $w_3$ =0.072, $w_4$ =0.125 |
| SASBDB IDs for data and models | *SASDWE6* |

(d) Structural parameters **Batch SAXS**

| Protein storage buffer used for subtraction | 20 mM $K_2HPO_4$/$KH_2PO_4$ pH 8, 100 mM NaCl, 200 mM $Li_2SO_4$, 1 mM DTT, 5% Glycerol | | |
|---|---|---|---|
| Protein Concentration | 0.5 mg/mL | 1 mg/mL | 2 mg/mL |
| Guinier analysis | | | |
| I(0) ($cm^{-1}$) | 0.02 ± 7.92e-4 | 0.0474 ± 9.63e-4 | 0.0995 ± 1.07e-3 |
| Rg (Å) | 46.25 ± 2.67 | 45.3366 ± 1.73 | 46.1781 ± 0.9 |
| qmin ($Å^{-1}$) | 0.00972 | 0.00972 | 0.00972 |
| qmax ($Å^{-1}$) | 0.02798 | 0.02537 | 0.02537 |
| qRg min | 0.4494 | 0.4406 | 0.4487 |
| qRg max | 1.2940 | 1.1502 | 1.1716 |
| n min | 40 | 40 | 40 |
| n max | 180 | 160 | 160 |

| | | | |
|---|---|---|---|
| $R^2$ | 0.9216 | 0.9589 | 0.9880 |
| Molecular Weight | | | |
| $V_c$ (kDa) | 122.4 | 79.3 | 97.5 |
| Bayes (kDa) | 130.9 (111.2-134.3) | 85.7 (81.9-95.8) | 101.0 (95.8-111.2) |
| P(r) analysis | | | |
| I(0) ($cm^{-1}$) | 0.02 ± 9.56e-4 | 0.05 ± 8.56e-4 | 0.1 ± 1.06e-3 |
| Rg (Å) | 48.49 ± 4.34 | 47.85 ± 1.5 | 48.88 ± 0.91 |
| Dmax (Å) | 185.0 | 180.0 | 183.0 |
| q range ($Å^{-1}$) | 0.01 - 0.173 | 0.01 - 0.176 | 0.01 - 0.173 |
| $\chi^2$ | 0.04 | 0.065 | 0.04 |
| GNOM solution | Reasonable | Reasonable | Reasonable |
| SASBDB IDs | *SASDWF6* | *SASDWG6* | *SASDWH6* |

**Aminoacidic Composition**

**>A**

**GMAMQMQLEANADTSVEEESFGPQPISRLEQCGINANDVKKLEEAGFHTVEAVAYAPKKELINIKGISEAKADKILAE AAKLVPMGETTETEFHQRRSEIIQITTGSKELDKLLQGGIETGSITEMFGEFRTGKTQICHTLAVTCQLPIDRGGGEGKA MYIDTEGTFRPERLLAVAERYGLSGSDVLDNVAYARAFNTDHQTQLLYQASAMMVESRYALLIVDSATALYRTDYSGR GELSARQMHLARFLRMLLRLADEFGVAVVITNQVVAQVDGAAMFAADPKKPIGGNIIAHASTTRLYLRKGRGETRIC KIYDSPCLPEAEAMFAINADGVGDAKD**

**>B**

**GMAMQMQLEANADTSVEEESFGPQPISRLEQCGINANDVKKLEEAGFHTVEAVAYAPKKELINIKGISEAKADKILAE AAKLVPMGETTETEFHQRRSEIIQITTGSKELDKLLQGGIETGSITEMFGEFRTGKTQICHTLAVTCQLPIDRGGGEGKA MYIDTEGTFRPERLLAVAERYGLSGSDVLDNVAYARAFNTDHQTQLLYQASAMMVESRYALLIVDSATALYRTDYSGR GELSARQMHLARFLRMLLRLADEFGVAVVITNQVVAQVDGAAMFAADPKKPIGGNIIAHASTTRLYLRKGRGETRIC KIYDSPCLPEAEAMFAINADGVGDAKD**

**>C**

**GWIKDFETSDTFFQTASGKNISVAKESFNKIVNFFDQKPEELHNFSLNSELHSDIRKNKMDILSYEETDIVKHKILKESVP VGTGNQLVTFQGQPERDEKIKEPTLLGFHTASGKKVKIAKESLDKVKNLFDEKEQ**

**Supplementary Table 17**

**SAXS Analysis RAD51 [F86E, A89E] in complex with BRCA2 truncation containing the second, the third and the fourth BRC repeats (BRC2-4)**

(a) Sample details

| | |
|---|---|
| Organism | Homo sapiens (Human) |
| Source | *E. Coli* expressed |
| Uniprot sequence ID | Q06609 in complex with 1212-1551 P51587<br>Q06609 mutations in position F86 and A89 with glutamic acid (E) |

| | |
|---|---|
| Extinction coefficient [A280, 0.1%(w/v)] | Ext. coefficient = 51910<br>Abs 0.1% (=1 g/l) 0.347, assuming all C form cystines<br><br>Ext. coefficient = 50660<br>Abs 0.1% (=1 g/l) 0.338, assuming all C are reduced |
| $\bar{v}$ ($cm^3$/ g) | 0.732 |
| Particle contrast from sequence and solvent constituents<br>$\bar{\rho}$ (ρ protein - ρ solvent) $10^{-6}$ $Å^{-2}$ | 2.838 |
| M from chemical composition (Da) | 149782.52 |
| SEC–SAXS column | Superose 6 Increase 3.2/300 column |
| Loading concentration (mg/mL) | 7 |
| Injection Volume (mL) | 0.05 |
| Flow rate (mL/min) | 0.075 |
| Solvent (solvent blanks taken from SEC flow-through prior to elution of protein) | 20 mM $K_2HPO_4$/$KH_2PO_4$ pH 8, 100 mM NaCl, 100 mM $Li_2SO_4$, 1 mM DTT, 1% sucrose |

(b) SAXS data-collection parameters

| | |
|---|---|
| Instrument/data processing | Diamond Light Source B21 SAXS Beamline with EigerX 4M (Dectris) detector[1] |
| Wavelength (Å) | 0.9464 |
| Beam size at sample (mm) | 1.0 x 0.25 |
| Beam size at detector (focus) (mm) | 0.05 x 0.05 (FWHM) |
| Camera length (m) | 3.6883 |
| q measurement range (Å) | 0.0045-0.34 |
| Absolute scaling method | Absolute intensity scaled to water scatter at 0.0163 |
| Sample configuration | SEC–SAXS Dual Agilent 1260 HPLC system |
| Sample temperature (°C) | 15 |
| Exposure time | SEC-SAXS Continuous 3 seconds exposure for 600 frames<br><br>Batch-SAXS 1 second exposure for 21 frames (sample has been moved during the exposure to attenuate radiation damage) |

(c) Structural parameters **SEC SAXS**

| | |
|---|---|
| Guinier analysis | |
| I(0) ($cm^{-1}$) | 0.13 ± 2.99e-4 |
| Rg (Å) | 66.37 ± 0.3 |
| qmin ($Å^{-1}$) | 0.00854 |
| qmax ($Å^{-1}$) | 0.01572 |

| | |
|---|---|
| qRg min | 0.5670 |
| qRg max | 1.0431 |
| n min | 31 |
| n max | 86 |
| Coefficient of correlation, $R^2$ | 0.9949 |
| Molecular Weight | |
| $V_c$ (kDa) | 152.1 |
| $V_p$ (kDa) | 228.0 |
| Bayes (kDa) | 185.8 (162.7- 194.9) |
| Shape&Size MW | N/A |
| Porod volume ($Å^{-3}$) | 560738 |
| P(r) analysis | |
| I(0) ($cm^{-1}$) | 0.13 ± 2.46e-4 |
| Rg (Å) | 69.87 ± 0.26 |
| Dmax (Å) | 280.0 |
| q range ($Å^{-1}$) | 0.008 - 0.12 |
| $\chi^2$ (total estimate from GNOM) | 1.054 |
| GNOM Interpretation | A reasonable solution |
| Ambimeter | |
| Number of compatible shapes | 154 |
| Ambiguity score | 2.188 |
| Ambimeter says | 3D reconstruction might be ambiguous |
| Shanum | |
| Dmax (Å) | 280 |
| Smax | 0.34 |
| N Shannon channels | 30.30 |
| Optimal number of Shannon channels | 27 |
| Optimal value of q-vector | 0.30 |
| Last optimal point | 2289 |
| Atomistic Modelling | |
| Structure | AlphaFold3 Model |
| q range for modelling | 0.0084-0.303 |
| FoXS | |
| $\chi^2$ | 144.80 |
| Predicted Rg (Å) | 44.42 |
| $c_1$, $c_2$ | 0.99, -0.29 |
| Multistate/ensemble models | |
| Starting Model | MultiFoXS modelled AlphaFold2 BRC1-4 modified to obtain the same aminoacidic composition of BRC2-4 |
| Flexible residues | Chain A 247-420, 456-516 |
| Rigid bodies connections | 219-222 A 191-194 D, 428-431 A 191-194 B<br>524-527 A 191-194 C |
| MultiFoXS | First Trial |
| No. of states | 1 |
| χ2 | 9.31 ± 35.16 |
| $c_1$, $c_2$ | 1.03, 0.05 |

| | |
|---|---|
| Rg values of each state (Å) | 74.32 |
| Weights wn | $w_1$ =1.0 |
| No. of states | 2 |
| χ2 | 3.8 ± 2.47 |
| $c_1$, $c_2$ | 1.0, 4.0 |
| Rg values of each state (Å) | 70.21, 193.396 |
| Weights wn | $w_1$ =0.487, $w_2$ =0.513 |
| No. of states | 3 |
| χ2 | 2.76 ± 0.51 |
| $c_1$, $c_2$ | 1.01, 4.0 |
| Rg values of each state (Å) | 193.40, 70.60, 134.76 |
| Weights wn | $w_1$ =0.389, $w_2$ =0.352, $w_3$ =0.259 |
| No. of states | 4 |
| χ2 | 1.62 ± 0.96 |
| $c_1$, $c_2$ | 0.99, 2.11 |
| Rg values of each state (Å) | 73.33, 193.40, 134.67, 52.15 |
| Weights wn | $w_1$ =0.707, $w_2$ =0.096, $w_3$ =0.072, $w_4$ =0.125 |
| No. of states | 5 |
| χ2 | 1.14 ± 0.13 |
| $c_1$, $c_2$ | 1.0, 0.49 |
| Rg values of each state (Å) | 139.40, 78.26, 134.76, 49.84, 80.65 |
| Weights wn | $w_1$ =0.121, $w_2$ =0.341, $w_3$ =0.065, $w_4$ =0.323, $w_5$ =0.15 |
| MultiFoXS | Second Trial |
| No. of states | 1 |
| χ2 | 24.98 ± 16.74 |
| $c_1$, $c_2$ | 1.05, 4.0 |
| Rg values of each state (Å) | 143.14 |
| Weights wn | $w_1$ = 1.0 |
| No. of states | 2 |
| χ2 | 5.9 ± 2.84 |
| $c_1$, $c_2$ | 1.02, 4.0 0.58 |
| Rg values of each state (Å) | 178.46, 54.60 |
| Weights wn | $w_1$ =0.704, $w_2$ =0.296 |
| No. of states | 3 |
| χ2 | 3.09 ± 0.3 |
| $c_1$, $c_2$ | 1.01, 3.91 |
| Rg values of each state (Å) | 178.51, 54.19, 116.867 |
| Weights wn | $w_1$ =0.459, $w_2$ =0.334, $w_3$ =0.206 |
| No. of states | 4 |
| χ2 | 2.16 ± 0.1 |
| $c_1$, $c_2$ | 1.03, 0.94 |
| Rg values of each state (Å) | 177.96, 53.94, 169.29, 84.84 |
| Weights wn | $w_1$ =0.1, $w_2$ =0.441, $w_3$ =0.269, $w_4$ =0.19 |
| No. of states | 5 |
| χ2 | 2.05 ± |
| $c_1$, $c_2$ | 1.03, 0.69 |

| | |
|---|---|
| Rg values of each state (Å) | 177.96, 53.94, 165.57, 86.81, 78.59 |
| Weights wn | $w_1$ =0.088, $w_2$ =0.459, $w_3$ =0.25, $w_4$ =0.13, $w_5$ =0.072 |
| SASBDB IDs for data and models | *SASDWD6* |

(d) Structural parameters **Batch SAXS**

| Protein storage buffer used for subtraction | 20 mM $K_2HPO_4$/$KH_2PO_4$ pH 8, 100 mM NaCl, 200 mM $Li_2SO_4$, 1 mM DTT, 5% Glycerol | | 20 mM Hepes pH 7.5, 350 mM KCl, 1 mM DTT, 5% Glycerol | |
|---|---|---|---|---|
| Protein Concentration | 1 mg/mL | 2 mg/mL | 3.5 mg/mL | 7 mg/mL |
| Guinier analysis | | | | |
| I(0) ($cm^{-1}$) | 0.06 ± 1.42e-3 | 0.13 ± 1.65e-3 | 0.14 ± 2.36e-3 | 0.54 ± 3.2e-3 |
| Rg (Å) | 64.94 ± 2.61 | 65.22 ± 1.4 | 68.49 ± 1.77 | 67.56 ± 0.73 |
| qmin ($Å^{-1}$) | 0.00841 | 0.00841 | 0.00841 | 0.00841 |
| qmax ($Å^{-1}$) | 0.01754 | 0.01754 | 0.01754 | 0.01624 |
| qRg min | 0.5464 | 0.5487 | 0.5763 | 0.5684 |
| qRg max | 1.1394 | 1.1442 | 1.2017 | 1.0971 |
| n min | 30 | 30 | 30 | 30 |
| n max | 100 | 100 | 100 | 90 |
| $R^2$ | 0.9736 | 0.9880 | 0.9871 | 0.9983 |
| Molecular Weight | | | | |
| $V_c$ (kDa) | 116.9 | 139.1 | 153.8 | 164.0 |
| Bayes (kDa) | 118.8 (116.0-142.2) | 169.2 (151.4-176.6) | 185.8 (151.4-194.9) | 185.8 (176.6-221.1) |
| P(r) analysis | | | | |
| I(0) ($cm^{-1}$) | 0.06 ± 1.55e-3 | 0.13 ± 1.87e-3 | 0.15 ± 2.22e-3 | 0.55 ± 3.26e-3 |
| Rg (Å) | 70.41 ± 3.09 | 70.57 ± 1.91 | 73.77 ± 1.84 | 71.53 ± 0.75 |
| Dmax (Å) | 285.0 | 285.0 | 285.0 | 283.0 |
| q range ($Å^{-1}$) | 0.008 - 0.123 | 0.008 - 0.122 | 0.008 - 0.117 | 0.008 - 0.118 |
| $\chi^2$ | 0.057 | 0.066 | 0.068 | 0.053 |
| GNOM | Reasonable | Reasonable | Reasonable | Reasonable |
| SASBDB IDs | SASDX95 | SASDXA5 | SASDXB5 | SASDXC5 |

**Aminoacidic Composition**

**>A**

**GMAMQMQLEANADTSVEEESFGPQPISRLEQCGINANDVKKLEEAGFHTVEAVAYAPKKELINIKGISEAKADKILAE AAKLVPMGETTETEFHQRRSEIIQITTGSKELDKLLQGGIETGSITEMFGEFRTGKTQICHTLAVTCQLPIDRGGGEGKA MYIDTEGTFRPERLLAVAERYGLSGSDVLDNVAYARAFNTDHQTQLLYQASAMMVESRYALLIVDSATALYRTDYSGR GELSARQMHLARFLRMLLRLADEFGVAVVITNQVVAQVDGAAMFAADPKKPIGGNIIAHASTTRLYLRKGRGETRIC KIYDSPCLPEAEAMFAINADGVGDAKD**

**>B**

**GMAMQMQLEANADTSVEEESFGPQPISRLEQCGINANDVKKLEEAGFHTVEAVAYAPKKELINIKGISEAKADKILAE AAKLVPMGETTETEFHQRRSEIIQITTGSKELDKLLQGGIETGSITEMFGEFRTGKTQICHTLAVTCQLPIDRGGGEGKA MYIDTEGTFRPERLLAVAERYGLSGSDVLDNVAYARAFNTDHQTQLLYQASAMMVESRYALLIVDSATALYRTDYSGR**

**GELSARQMHLARFLRMLLRLADEFGVAVVITNQVVAQVDGAAMFAADPKKPIGGNIIAHASTTRLYLRKGRGETRIC KIYDSPCLPEAEAMFAINADGVGDAKD**

**>C**

**GMAMQMQLEANADTSVEEESFGPQPISRLEQCGINANDVKKLEEAGFHTVEAVAYAPKKELINIKGISEAKADKILAE AAKLVPMGETTETEFHQRRSEIIQITTGSKELDKLLQGGIETGSITEMFGEFRTGKTQICHTLAVTCQLPIDRGGGEGKA MYIDTEGTFRPERLLAVAERYGLSGSDVLDNVAYARAFNTDHQTQLLYQASAMMVESRYALLIVDSATALYRTDYSGR GELSARQMHLARFLRMLLRLADEFGVAVVITNQVVAQVDGAAMFAADPKKPIGGNIIAHASTTRLYLRKGRGETRIC KIYDSPCLPEAEAMFAINADGVGDAKD**

**>D**

**MANEVGFRGFYSAHGTKLNVSTEALQKAVKLFSDIENISEETSAEVHPISLSSSKCHDSVVSMFKIENHNDKTVSEKNN KCQLILQNNIEMTTGTFVEEITENYKRNTENEDNKYTAASRNSHNLEFDGSDSSKNDTVCIHKDETDLLFTDQHNICLK LSGQFMKEGNTQIKEDLSDLTFLEVAKAQEACHGNTSNKEQLTATKTEQNIKDFETSDTFFQTASGKNISVAKESFNKIV NFFDQKPEELHNFSLNSELHSDIRKNKMDILSYEETDIVKHKILKESVPVGTGNQLVTFQGQPERDEKIKEPTLLGFHTA SGKKVKIAKESLDKVKNLFDEKEQ**

**Supplementary Table 18**

**SAXS Analysis RAD51 [F86E, A89E] in complex with BRCA2 truncation containing the first, the second, the third and the fourth BRC repeats (BRC1-4)**

(a) Sample details

| | |
|---|---|
| Organism | Homo sapiens (Human) |
| Source | *E. Coli* expressed |
| Uniprot sequence ID | Q06609 in complex with 1002-1551 P51587<br>Q06609 mutations in position F86 and A89 with glutamic acid (E) |
| Extinction coefficient [$A_{280}$, 0.1%(w/v)] | Ext. coefficient = 71780<br>Abs 0.1% (=1 g/l) assuming all pairs of C form cystines<br><br>Ext. coefficient = 70030<br>Abs 0.1% (=1 g/l) 0.333, assuming all C are reduced |
| $\bar{v}$ from chemical composition of N-terminal domain ($cm^3$/ g) | 0.732 |
| Particle contrast from sequence and solvent constituents<br>$\bar{\rho}$ (ρ protein - ρ solvent) $10^{-6}$ $Å^{-2}$ | 2.839 |
| M from chemical composition (Da) | 210239.98 |
| SEC–SAXS column | Superose 6 Increase 3.2/300 |
| Loading concentration (mg/mL) | 3.7 |
| Injection Volume (mL) | 55 |
| Flow rate (mL/min) | 0.075 |

| | |
|---|---|
| Solvent (solvent blanks taken from SEC flow-through prior to elution of protein) | 20 mM $K_2HPO_4$/$KH_2PO_4$ pH 8 100 mM NaCl, 100 mM $Li_2SO_4$, 1 mM DTT, 1% sucrose |

(b) SAXS data-collection parameters

| | |
|---|---|
| Instrument/data processing | Diamond Light Source B21 SAXS Beamline with EigerX 4M (Dectris) detector[1] |
| Wavelength (Å) | 0.9464 |
| Beam size at sample (mm) | 1.0 x 0.25 |
| Beam size at detector (focus) (mm) | 0.05 x 0.05 (FWHM) |
| Camera length (m) | 3.6883 |
| q measurement range (Å) | 0.0045-0.34 |
| Absolute scaling method | Absolute intensity scaled to water scatter at 0.0163 |
| Sample configuration | SEC–SAXS Dual Agilent 1260 HPLC system |
| Sample temperature (°C) | 15 |
| Exposure time | SEC-SAXS Continuous 3 seconds exposure for 600 frames<br><br>Batch-SAXS 1 second exposure for 21 frames (sample has been moved during the exposure to attenuate radiation damage) |

(c) Structural parameters **SEC SAXS**

| | |
|---|---|
| Guinier analysis | |
| I(0) ($cm^{-1}$) | 0.0532 ± 2.23e-4 |
| Rg (Å) | 77.09 ± 0.47 |
| qmin ($Å^{-1}$) | 0.00841 |
| qmax ($Å^{-1}$) | 0.01598 |
| qRg min | 0.6486 |
| qRg max | 1.2318 |
| n min | 27 |
| n max | 85 |
| Coefficient of correlation, $R^2$ | 0.9906 |
| Molecular Weight | |
| $V_c$ (kDa) | 220.9 |
| $V_p$ (kDa) | 349.9 |
| Bayes (kDa) | 242.6 (221.1-372.7) |
| Shape&Size MW | N/A |
| Porod volume ($Å^{-3}$) | 876750 |
| P(r) analysis | |
| I(0) ($cm^{-1}$) | 0.05 ± 3.02e-4 |
| Rg (Å) | 84.26 0.78 |
| Dmax (Å) | 340.0 |
| q range ($Å^{-1}$) | 0.008 - 0.104 |
| $\chi^2$ (total estimate from GNOM) | 1.201 |

| GNOM Interpretation | A reasonable solution |
|---|---|
| Ambimeter | |
| Number of compatible shapes | 168 |
| Ambiguity score | 2.225 |
| Ambimeter says | 3D reconstruction might be ambiguous |
| Shanum | |
| Dmax (Å) | 340 |
| Smax | 0.293 |
| N Shannon channels | 31.71 |
| Optimal number of Shannon channels | 28 |
| Optimal value of q-vector | 0.2587 |
| Last optimal point | 1947 |
| Atomistic Modelling | |
| Structure | AlphaFold2.3 Model Best Ranked Model |
| q range for modelling | 0.0083-0.2587 |
| FoXS | |
| $\chi 2$ | 40.73 |
| Predicted Rg (Å) | 49.19 |
| c1, c2 | 0.99, 1.82 |
| Structure | AlphaFold2.3 Model Compact Conformation |
| q range for modelling | 0.0083-0.2587 |
| FoXS | |
| $\chi 2$ | 51.34 |
| Predicted Rg (Å) | 46.77 |
| c1, c2 | 0.99, 2.94 |
| Multistate/ensemble models | |
| Starting Model | AlphaFold2.3 Model Best Ranked Model |
| Flexible residues | Chain A 38-211, 247-420, 456-516 |
| Rigid bodies connections | 219-222 A 191-194 D, 428-431 A 191-194 B<br>524-527 A 191-194 C, 10-13 A 191-194 E |
| MultiFoXS | First Trial |
| No. of states | 1 |
| $\chi 2$ | 2.82 ± 14.73 |
| $c_1$, $c_2$ | 0.99, -0.5 |
| Rg values of each state (Å) | 74.76 |
| Weights wn | $w_1$ = 1 |
| No. of states | 2 |
| $\chi 2$ | 1.35 ± 0.28 |
| $c_1$, $c_2$ | 0.99, 2.74 |
| Rg values of each state (Å) | 71.01, 210.58 |
| Weights wn | $w_1$ = 0.558, $w_2$ =0.442 |
| No. of states | 3 |
| $\chi 2$ | 1.28 ± 0.05 |
| $c_1$, $c_2$ | 1.0, 2.56 |
| Rg values of each state (Å) | 225.727, 53.78, 115.13 |
| Weights wn | $w_1$ = 0.37, $w_2$ =0.289, $w_3$ =0.341 |

| | |
|---|---|
| No. of states | 4 |
| χ2 | 1.25 ± 0.02 |
| $c_1$, $c_2$ | 1.0, 1.66 |
| Rg values of each state (Å) | 225.73, 53.78, 115.13, 106.95 |
| Weights wn | $w_1$ = 0.3, $w_2$ =0.33, $w_3$ =0.224, $w_4$ =0.146, $w_5$ =0.072 |
| No. of states | 5 |
| χ2 | 1.24 ± 0.01 |
| $c_1$, $c_2$ | 0.99, 2.11 |
| Rg values of each state (Å) | 225.73, 53.78, 115.13, 241.156, 94.76 |
| Weights wn | $w_1$ = 0.198, $w_2$ =0.307, $w_3$ =0.195, $w_4$ =0.141, $w_5$ =0.16 |
| MultiFoXS | Second Trial |
| No. of states | 1 |
| χ2 | 2.67 ± 10.05 |
| $c_1$, $c_2$ | 1.04, -0.5 |
| Rg values of each state (Å) | 83.42 |
| Weights wn | $w_1$ = 1 |
| No. of states | 2 |
| χ2 | 1.45 ± 0.45 |
| $c_1$, $c_2$ | 0.99, 4.0 |
| Rg values of each state (Å) | 74.52, 208.255 |
| Weights wn | $w_1$ = 0.499, $w_2$ =0.501 |
| No. of states | 3 |
| χ2 | 1.3 ± 0.05 |
| $c_1$, $c_2$ | 0.99, 2.56 |
| Rg values of each state (Å) | 74.52, 226.13, 115.958 |
| Weights wn | $w_1$ = 0.198, $w_2$ =0.307, $w_3$ =0.195, $w_4$ =0.144 |
| No. of states | 4 |
| χ2 | 1.24 ± 0.05 |
| $c_1$, $c_2$ | 0.99, 1.03 |
| Rg values of each state (Å) | 231.701, 55.69, 124.11, 94.20 |
| Weights wn | $w_1$ = 0.212, $w_2$ =0.393, $w_3$ =0.161, $w_4$ =0.234 |
| No. of states | 5 |
| χ2 | 1.24 ± 0.02 |
| $c_1$, $c_2$ | 1.0, 2.11 |
| Rg values of each state (Å) | 231.71, 57.76, 137.14, 243.79, 103.96 |
| Weights wn | $w_1$ = 0.219, $w_2$ =0.358, $w_3$ =0.095, $w_4$ =0.153, $w_5$ =0.175 |
| SASBDB IDs for data and models | *SASDWN6* |

(d) Structural parameters **Batch SAXS**

| Protein storage buffer used for subtraction | 20 mM $K_2HPO_4/KH_2PO_4$ pH 8, 100 mM NaCl, 200 mM $Li_2SO_4$, 1 mM DTT, 5% Glycerol | | |
|---|---|---|---|
| Protein Concentration | 0.5 mg/mL | 0.94 mg/mL | 2 mg/mL |
| Guinier analysis | | | |
| I(0) ($cm^{-1}$) | 0.04 ± 1.96e-3 | 0.08 ± 1.66e-3 | 0.19 ± 2.11e-3 |
| Rg (Å) | 81.06 | 75.94 ± 2.13 | 76.89 ± 1.14 |

| | | | |
|---|---|---|---|
| qmin (Å$^{-1}$) | 0.00894 | 0.00841 | 0.00854 |
| qmax (Å$^{-1}$) | 0.01611 | 0.01715 | 0.01689 |
| qRg min | 0.7243 | 0.6389 | 0.6569 |
| qRg max | 1.3058 | 1.3027 | 1.2988 |
| n min | 34 | 30 | 31 |
| n max | 89 | 97 | 95 |
| $R^2$ | 0.9615 | 0.9864 | 0.9932 |
| Molecular Weight | | | |
| $V_c$ (kDa) | 202.4 | 183.3 | 189.8 |
| Bayes (kDa) | 242.6 (194.9-264.2) | 208.0 (176.6-264.2) | 208.0 (176.6-264.2) |
| P(r) analysis | | | |
| I(0) (cm$^{-1}$) | 0.04 ± 2.01e-3 | 0.08 ± 2.12e-3 | 0.2 ± 2.88e-3 |
| Rg (Å) | 87.37 ± 6.21 | 85.59 ± 3.19 | 85.21 ± 1.87 |
| Dmax (Å) | 330.0 | 333.0 | 330.0 |
| q range (Å$^{-1}$) | 0.009-0.098 | 0.008-0.105 | 0.008 0.104 |
| $\chi^2$ | 0.048 | 0.06 | 0.059 |
| GNOM solution | Reasonable | Reasonable | Reasonable |
| SASBDB IDs | *SASDWP6* | *SASDWQ6* | *SASDWR6* |

**Aminoacidic Composition**

**>A**

**GMAMQMQLEANADTSVEEESFGPQPISRLEQCGINANDVKKLEEAGFHTVEAVAYAPKKELINIKGISEAKADKILAE AAKLVPMGETTETEFHQRRSEIIQITTGSKELDKLLQGGIETGSITEMFGEFRTGKTQICHTLAVTCQLPIDRGGGEGKA MYIDTEGTFRPERLLAVAERYGLSGSDVLDNVAYARAFNTDHQTQLLYQASAMMVESRYALLIVDSATALYRTDYSGR GELSARQMHLARFLRMLLRLADEFGVAVVITNQVVAQVDGAAMFAADPKKPIGGNIIAHASTTRLYLRKGRGETRIC KIYDSPCLPEAEAMFAINADGVGDAKD**

**>B**

**GMAMQMQLEANADTSVEEESFGPQPISRLEQCGINANDVKKLEEAGFHTVEAVAYAPKKELINIKGISEAKADKILAE AAKLVPMGETTETEFHQRRSEIIQITTGSKELDKLLQGGIETGSITEMFGEFRTGKTQICHTLAVTCQLPIDRGGGEGKA MYIDTEGTFRPERLLAVAERYGLSGSDVLDNVAYARAFNTDHQTQLLYQASAMMVESRYALLIVDSATALYRTDYSGR GELSARQMHLARFLRMLLRLADEFGVAVVITNQVVAQVDGAAMFAADPKKPIGGNIIAHASTTRLYLRKGRGETRIC KIYDSPCLPEAEAMFAINADGVGDAKD**

**>C**

**GMAMQMQLEANADTSVEEESFGPQPISRLEQCGINANDVKKLEEAGFHTVEAVAYAPKKELINIKGISEAKADKILAE AAKLVPMGETTETEFHQRRSEIIQITTGSKELDKLLQGGIETGSITEMFGEFRTGKTQICHTLAVTCQLPIDRGGGEGKA MYIDTEGTFRPERLLAVAERYGLSGSDVLDNVAYARAFNTDHQTQLLYQASAMMVESRYALLIVDSATALYRTDYSGR GELSARQMHLARFLRMLLRLADEFGVAVVITNQVVAQVDGAAMFAADPKKPIGGNIIAHASTTRLYLRKGRGETRIC KIYDSPCLPEAEAMFAINADGVGDAKD**

**>D**

**GMAMQMQLEANADTSVEEESFGPQPISRLEQCGINANDVKKLEEAGFHTVEAVAYAPKKELINIKGISEAKADKILAE AAKLVPMGETTETEFHQRRSEIIQITTGSKELDKLLQGGIETGSITEMFGEFRTGKTQICHTLAVTCQLPIDRGGGEGKA**

**MYIDTEGTFRPERLLAVAERYGLSGSDVLDNVAYARAFNTDHQTQLLYQASAMMVESRYALLIVDSATALYRTDYSGR GELSARQMHLARFLRMLLRLADEFGVAVVITNQVVAQVDGAAMFAADPKKPIGGNIIAHASTTRLYLRKGRGETRIC KIYDSPCLPEAEAMFAINADGVGDAKD**

**>E**

**GNHSFGGSFRTASNKEIKLSEHNIKKSKMFFKDIEEQYPTSLACVEIVNTLALDNQKKLSKPQSINTVSAHLQSSVVVSD CKNSHITPQMLFSKQDFNSNHNLTPSQKAEITELSTILEESGSQFEFTQFRKPSYILQKSTFEVPENQMTILKTTSEECRD ADLHVIMNAPSIGQVDSSKQFEGTVEIKRKFAGLLKNDCNKSASGYLTDENEVGFRGFYSAHGTKLNVSTEALQKAVK LFSDIENISEETSAEVHPISLSSSKCHDSVVSMFKIENHNDKTVSEKNNKCQLILQNNIEMTTGTFVEEITENYKRNTENE DNKYTAASRNSHNLEFDGSDSSKNDTVCIHKDETDLLFTDQHNICLKLSGQFMKEGNTQIKEDLSDLTFLEVAKAQEA CHGNTSNKEQLTATKTEQNIKDFETSDTFFQTASGKNISVAKESFNKIVNFFDQKPEELHNFSLNSELHSDIRKNKMDIL SYEETDIVKHKILKESVPVGTGNQLVTFQGQPERDEKIKEPTLLGFHTASGKKVKIAKESLDKVKNLFDEKEQ**

**Supplementary Table 19**: software employed for SAXS data reduction, analysis and interpretation

| | |
|---|---|
| SAXS data reduction | Diamond Light Source Generic Data Acquisition, software. NXS files are automatically normalized and reduced using routines provided in the Diamond Light Source developed Data Analysis WorkbeNch (DAWN) The output is azimuthally averaged. Normalized one-dimensional data are stored in a three column ASCII file (*.dat). Solvent subtraction and frame selection were performed using ScÅtter for SEC-SAXS experiments. [2] |
| Extinction coefficient estimate | Expasy ProtParam tool[3] |
| Calculation of $\bar{v}$ and $\bar{\rho}$ values | MULCh: ModULes for the analysis of Contrast variation data (v1.1.1 - 2017) [4], US-SOMO Beta 0.1[5] |
| Basic analyses: Guinier, P(r), Mw, VP | BioXTAS RAW[6, 7, 8, 9, 10, 11], GNOM[12], PRIMUS[13], Shanum[14], Ambimeter[15] |
| Atomic structure modelling | FoXS via web server (https://modbase.compbio.ucsf.edu/foxs) [16, 17]<br><br>MultiFoXS via web server (https://modbase.compbio.ucsf.edu/multifoxs/) [16, 17] |
| 3D Models generation | AlphaFold2.3 or AlphaFold3[18, 19] |

**Supplementary Methods**

**Supplementary Table 20**

| | Aminoacidic composition | C-term | Manufacturer |
|---|---|---|---|
| BRC1 | NHSFGGSFRTASNKEIKLSEHNIKKSKMFFKDIEE | Amide | ThermoFisher |
| BRC2 | NEVGFRGFYSAHGTKLNVSTEALQKAVKLFSDIEN | Amide | ThermoFisher |
| BRC3 | FETSDTFFQTASGKNISVAKESFNKIVNFFDQKPE | Amide | ThermoFisher |
| BRC5 | IENSALAFYTSCSRKTSVSQTSLLEAKKWLREGIF | $-NH_2$ | Peptide P. Research Ltd. |
| BRC6 | FEVGPPAFRIASGKIVCVSHETIKKVKDIFTDSFS | $-NH_2$ | Peptide P. Research Ltd. |
| BRC7 | SANTCGIFSTASGKSVQVSDASLQNARQVFSEIED | Amide | ThermoFisher |
| BRC8 | NSSAFSGFSTASGKQVSILESSLHKVKGVLEEFDL | Amide | ThermoFisher |
| $^{19}$F-FXXA-BRC1 | F(4-F)-GGSFRTASNKEI | COOH | Peptide P. Research Ltd. |
| $^{19}$F-FXXA-BRC2 | F(4-F)-FRGFYSAHGTKL | COOH | Peptide P. Research Ltd. |
| $^{19}$F-FXXA-BRC3 | F(4-F)-DTFFQTASGKNI | COOH | Peptide P. Research Ltd. |
| $^{19}$F-FXXA-BRC7 | F(4-F)-CGIFSTASGKSV | COOH | Peptide P. Research Ltd. |
| $^{19}$F-FXXA-BRC8 | F(4-F)-FSGFSTASGKQV | COOH | Peptide P. Research Ltd. |
| Bio-BRC1 | Bio-Ahx-NHSFGGSFRTASNKEIKLSEHNIKKSKMFFKDIEE | COOH | Peptide P. Research Ltd. |
| Bio-BRC2 | Bio-Ahx-NEVGFRGFYSAHGTKLNVSTEALQKAVKLFSDIEN | COOH | Peptide P. Research Ltd. |
| Bio-BRC3 | Bio-Ahx-FETSDTFFQTASGKNISVAKESFNKIVNFFDQKPE | COOH | Peptide P. Research Ltd. |
| Bio-BRC4 | Bio-Ahx-KEPTLLGFHTASGKKVKIAKESLDKVKNLFDEKEQ | COOH | Peptide P. Research Ltd. |
| Bio-BRC7 | Bio-Ahx-SANTCGIFSTASGKSVQVSDASLQNARQVFSEIED | COOH | Peptide P. Research Ltd. |
| Bio-BRC8 | Bio-Ahx-NSSAFSGFSTASGKQVSILESSLHKVKGVLEEFDL | COOH | Peptide P. Research Ltd. |

Bio-Ahx Biotin– Amino hexanoic acid

F(4-F)- 4-fluorophenylalanine (para)
